# A Luminal–Basal Stratification of the Native Human Pancreatic Duct is Differentially Represented in Pancreatic Cancers

**DOI:** 10.1101/2025.01.14.632911

**Authors:** JL Van den Bossche, M Van der Vliet, E Michiels, O Azurmendi Senar, S Van Lint, Z Madran, K Coolens, J Brons, M Nacher, T Arsenijevic, N Messaoudi, P Lefesvre, C Bouchart, L Verset, J Navez, J Fallas, N Dusetti, E Montanya, M Rovira, J Van Laethem, I Houbracken, J Baldan, I Rooman

**Affiliations:** Vrije Universiteit Brussel (VUB), Translational Oncology Research Centre, Laboratory for Medical and Molecular Oncology, Brussels, Belgium; BruPACT, Brussels, Belgium; Laboratory of Experimental Gastroenterology, Faculty of Medicine, Université Libre de Bruxelles (ULB), Brussels, Belgium; CIBER of Diabetes and Associated Metabolic Diseases (CIBERDEM), Hospital Universitari Bellvitge, and Bellvitge Biomedical Research Institute (IDIBELL), Barcelona, Spain; Department of Gastroenterology, Hepatology and Digestive Oncology, HUB Bordet Erasme Hospital, Université Libre de Bruxelles, Brussels, Belgium; Department of Hepatopancreatobiliary Surgery, Vrije Universiteit Brussel, Universitair Ziekenhuis Brussel and Europe Hospitals, Brussels, Belgium; Department of Pathology, UZ Brussel, Brussels, Belgium; Department of Radiation Oncology, HUB Institut Jules Bordet, Université Libre de Bruxelles, Belgium; Cancer Research Center of Marseille (CRCM), INSERM, CNRS, Institut Paoli-Calmettes, Aix-Marseille University, Marseille, France; Department of Clinical Sciences, University of Barcelona, Barcelona, Spain

**Keywords:** Spatial transcriptomics, pancreatic cancer, basal cells, heterogeneity, squamous

## Abstract

**Background:** A resemblance of pancreatic tumors to their native tissue architecture remains largely unexplored, while it may reveal novel insights into the healthy and diseased tissue.

**Objective:** This study aims at generating a spatially resolved map of human pancreatic duct cell populations in the native tissue and in tumors, i.e. pancreatic ductal adenocarcinoma (PDAC) and adenosquamous cancer of the pancreas (ASCP).

**Design:** New datasets were acquired with several spatial transcriptomics platforms and were integrated with public single-cell RNAseq datasets and validated by multiplex immunofluorescence. Cell lines and primary human cell cultures were genetically manipulated.

**Results:** Groups of Keratin-5^+^ cells in larger ducts have a gene signature reminiscent of stem cells and (supra)basal cells from other tissues. At single cell resolution, this group comprises ΔNp63^+^ basal cells (BAS) and ΔNp63^-^ supra-basally residing luminal-B cells (LUM-B). The latter express previously unreported MUC4 and MUC16. Additionally, we identified three other luminal cell populations in the ducts (LUM-A, -C, -D). In cancer, BAS and LUM-B signatures associate with basal-like (BL) PDAC, and correlate with lower survival. However, PDAC exhibits a random spatial pattern and fragmented native expression programs while ASCP preserve the identity of LUM-B and BAS in a spatially unmixed pattern. ΔNp63 drives cell plasticity to BAS, conserved from the native tissue to cancer.

**Conclusion:** Spatially distinct duct cell populations are revealed, and the extent of preservation of the native cell identities in pancreatic cancer underpins distinct tumor identities. This warrants a separate consideration in research and therapy.

**What is already known on this topic:** While tumors in the pancreas can exhibit basal-like and squamous features resembling tumors of other tissues, their resemblance to the native duct cells, and possible subpopulations thereof, had not been studied in detail.

**What this study adds:** One basal and four luminal spatially distinct cell populations exist in human pancreatic ducts. These are best conserved in ASCP while PDAC shows loss of the native cell identity program.

**How this study might affect research, practice or policy:** Our study demonstrates fundamental differences between the tumor cells of the BL PDAC versus the rare ASCP, highlighting the necessity for a clear distinction between them in research and in tumor-specific treatments. Meanwhile, the research community should acknowledge the spatial and functional heterogeneity of human duct cells, including in their experimental models.

## Introduction

Organs are organized into specialized cell types, with subpopulations and distinct cell states. In cancer, these native populations are often retained, superimposed with recurrent meta-programs, such as hypoxia or proliferation(1). Such relationships remain largely unexplored for the human exocrine pancreas and its associated malignancies.

The normal exocrine pancreas (NP) comprises digestive enzyme-producing acini and ducts that collect and transport the enzymes to the duodenum(2). Pancreatic Ductal Adenocarcinoma (PDAC), the most prevalent pancreatic cancer, exhibits a ductal histology. Bulk transcriptomic profiling revealed molecular subtypes(3–7) and, more recently, single-cell sequencing studies provided insights with higher resolution(8,9). These studies distinguished classical (CL) and basal-like (BL) subtypes and cell states, with BL tumors characterized by expression of KRT5 and KRT17, a ΔNp63-driven signature and linked to poorer outcomes(10,11). The term ‘BL’ refers to parallels with basal subtypes in other cancers such as lung, breast, and prostate(12–14). At single duct and single cell level, transitions of CL and BL states were documented(15,16). Also basaloid, squamoid and mesenchymal states were discriminated, all resorting under the denominator ‘BL’(8,17). Notably, we reported before that ΔNp63 expression in BL PDAC is heterogeneous, and not all tumors or cell lines display it(17,18). Adenosquamous carcinoma of the pancreas (ASCP), a rarer and more aggressive subtype, is histologically defined by ≥30% ΔNp63⁺ cells(19–22). Literature suggests that up to 10% of pancreatic tumors can be ASCP, while often diagnosed, and always treated as PDAC(19,20). Comparisons of BL PDAC and ASCP beyond genomic studies(23,24) remain limited.

We lack basic insights into these different pancreatic tumor (sub)types as well as into the human ductal system that has generally been considered a simple epithelium of one cell type. Few studies, most in mice, have started describing duct cell heterogeneity(25–27). Our group reported rare ΔNp63^+^KRT5^+^ cells situated on the basal lamina of human pancreatic ducts(18), thus reminiscent of basal cells. These cells are enriched in chronic pancreatitis (CP), a risk factor for cancer(18). Basal cells have been described in other organs, where they function as stem cells residing on the basal lamina, not facing a lumen. They can give rise to suprabasal cells that line the lumen, i.e. luminal cells, for example airway basal cells giving rise to club cells that secrete secretoglobins into the lumen of the airways(28).

Given this gap of knowledge on the native tissue and the fact that tumors across many organs recapitulate native epithelial hierarchies(29–31), we leveraged newly available spatial transcriptomics tools to investigate the non-neoplastic pancreatic duct cells and projected these findings onto PDAC and ASCP, unveiling a luminal-basal stratification in the native duct system that is best preserved in ASCP.

## Methods

### Human material

Samples from pancreas of multiorgan donors without history of pancreatic disease were received from Hospital Universitari Bellvitge-IDIBELL (Barcelona, Spain) and Betacelbank Universitair Ziekenhuis Brussel (UZB). Tumor resections were obtained from the Department of Hepatopancreatobiliary Medicine and Pathology UZB. ASCP and CP samples were obtained from Université Libre de Bruxelles through the BruPaCT (Brussels Pancreatic Cancer Team) initiative. The use of clinical samples conformed with guidelines and regulations in Belgium with ethical approval of UZ Brussel (B.U.N. 1432022000112). H&E and IHC images of all human samples are included in Supplemental File 1.

### Spatial transcriptomics

GeoMx Digital Spatial Profiler® (NanoString, Washington, USA; Supplemental Figure 1) discovery analysis was performed on non-neoplastic pancreas (n=3, on 2 slides) with regions of analysis selected for the presence of keratin (KRT)5^+^ regions within the Pan-cytokeratin (PanCK) ductal system. Targeted UV-cleavable probes were collected after hybridization. Expression data were generated by NanoString nCounter® tag counting, processed using the DSP analysis platform, and analyzed with a linear mixed model (LMM) with Benjamini–Hochberg (BH) correction. The Resolve Molecular Cartography® (MC)(Resolve Biosciences, Monheim am Rhein, Germany) study used a custom-designed 100-gene panel, selected based on our GeoMx® experiment (Supplemental Table 1, Supplemental Figure 1, on non-neoplastic pancreas (n=4), CL PDAC (n=3), BL PDAC (n=3) and ASCP (n=2). Transcriptomic PDAC subtype assignment was done with the Chan-Seng-Yue classification(7).

**Validation studies on larger cohorts and other methods are described in Supplemental Methods and primary antibodies in Supplemental Table 2.**

## Results

### Human pancreatic KRT5^+^ cells have a transcriptomic profile consistent with (supra)basal cells from other tissues

Given our previous report of cells with basal cell markers (including KRT5) in the human pancreas(18), we first engaged in a discovery approach profiling areas of PanCK^+^KRT5^-^ and PanCK^+^KRT5^+^ in non-neoplastic pancreas (n=3, 24 ROIs, 6 mm^2^) using GeoMx® (Figure 1A, Supplemental Figure 1). After obtaining on average 858.000 unique reads per Region Of Interest (ROI) (Figure 1B), comparative transcriptomic analysis demonstrated typical duct cell markers (HNF1B, ONECUT1/2) in the KRT5^-^ population and typical basal cell markers from other organs(28,32) (KRT15, KRT17, KRT13, TP63, TRIM29) in the KRT5^+^ subset of cells (Figure 1C, Supplemental file 2).

**Figure 1:**
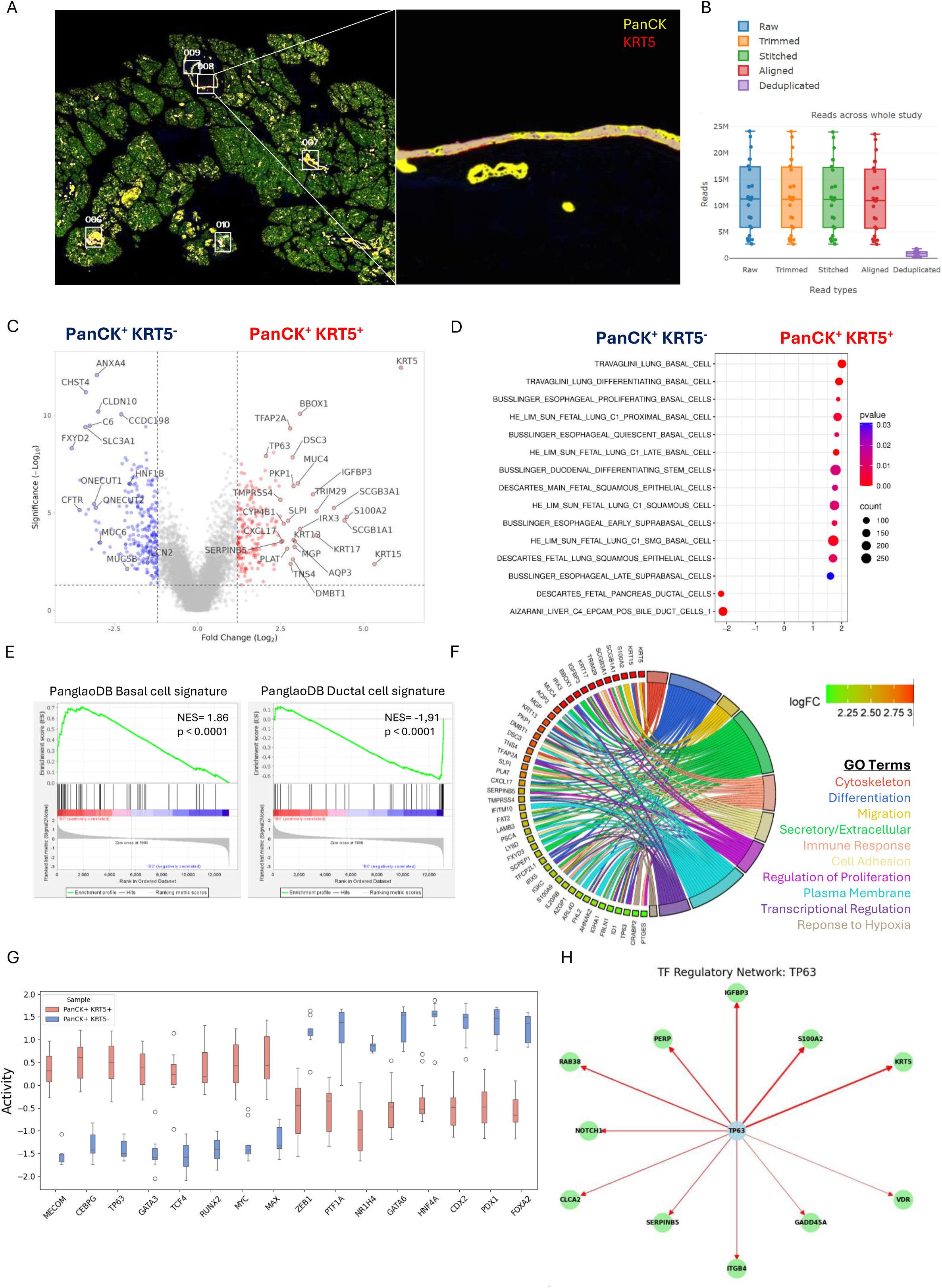
Non-neoplastic human pancreatic KRT5^+^ cells have a transcriptomic profile consistent with (supra)basal cells from other tissues. (A) Representation of GeoMx® slide with regions of interest. (B) Quality control of reads of GeoMx® experiment. (C) Volcano plot of differentially expressed genes between Pan-cytokeratin (PanCK)^+^ Keratin-5 (KRT5)^-^ and PanCK^+^KRT5^+^ cells in non-neoplastic human pancreas (n=3). Analysis was performed using GeoMx® DSP. Top expressed genes and known basal/duct markers are shown. (D) Gene Set Enrichment Analysis (GSEA) applied to data from panel A with cell type genesets (c8.all.v2024.1.Hs.symbols.gmt). Number of permutations = 1000, normalization performed using meandiv. (E) Enrichment plots using PanglaoDB cell profiles applied to the genes from panel C. NES = 1.86 (basal profile) and -1.91 (ductal profile). (F) Chord plot of PanCK^+^KRT5^+^-specific genes expressed in non-neoplastic pancreas with the associated Gene Ontology Term. The absolute Log_2_Fold Change values are represented with a color scheme. (G) Boxplot showing activity scores for top differentially expressed transcription factors in two populations. Analysis performed with Priori package. (H) Transcriptional network for TP63 created with Priori. For panel C, statistical analysis was performed with a linear mixed model (LMM) with Benjamini-Hochberg (BH) correction. Genes with p-value < 0.05 were considered differentially expressed.

Gene set enrichment analysis (GSEA’) substantiated the identity of the KRT5^+^ cells; top C8 cell type enriched signatures included basal and stem cell types from organs such as lung and esophagus (Figure 1D) and pointed to diverse positions within the epithelial layer (basal/suprabasal) and distinct activation states (differentiating/proliferating/quiescent). Using the Panglao Database(33) with single cell profiles from airways, prostate and breast, the basal cell signature was strongly enriched in the pancreatic KRT5^+^ cells, whereas the ductal cell signature was negatively enriched (Figure 1E). The top differentially expressed genes of KRT5^+^ cells, when annotated in Gene Ontology (Figure 1F), featured secreted proteins such as those from lung club cells, i.e. secretoglobins SCGB1A1 and SCGB3A1(34), which are also generated by basal cells in the airways(28). We also note Mucin-4 (MUC4), that is extensively described as undetectable in NP and CP(35–37) and highly expressed in PDAC(38), being enriched in the KRT5^+^ regions. Under ‘Transcriptional Regulation’, the basal cell transcription factor TP63 featured, along with a reported coregulator TFAP2A(39), and other transcription factors such as IRX3, IRX5 or TFCP2L1 that are involved in development, cell fate decisions and pluripotency of stem cells(40,41). Other interesting properties included immune regulation, differentiation, proliferation and migration (Figure 1F). Normalized counts for all genes of interest are provided (Supplemental Figure 2A).

Analysis with the Priori package(42) (Figure 1G), which infers transcription factor activity by combining gene expression data of bulk RNAseq with prior transcription factor-target knowledge, demonstrated that those with highest variance between KRT5^+^ and KRT5^-^ cells belonged to lineage specification and oncogenic reprogramming (CEBPG, TP63, MYC, MECOM) for the KRT5^+^ cells while it are the typical pancreatic identity genes (PTF1a, PDX1, GATA6) in the KRT5^-^ cells. Indeed, in the KRT5^+^ cells, CEBPG, whose expression levels are regulated by TP63 for maintaining basal cell identity(43), had the highest score, next to TP63 itself, reinforcing the role of TP63. MYC, also regulated by TP63, is involved in maintaining the undifferentiated state of basal cells and in their self-renewal capacity(44–46). Gene network analysis identified TP63-interacting genes expressed in KRT5^+^ cells, among which NOTCH1 that is described to regulate differentiation of basal cells(47), hinting towards a similar mechanism in pancreas (Figure 1H). The gene network of MYC is shown in Supplemental Figure 2B.

Collectively, this first time full transcriptomic profiling supported the existence of KRT5^+^TP63^+^ pancreatic basal cells, with a gene signature as reported in (supra)basal cells from other epithelia.

### The human pancreatic duct system comprises a basal and four luminal populations

Based on the above GeoMx® profiling and murine pancreatic ductal gene signatures (27), we designed a 100-gene panel for obtaining single-cell resolution transcript detection using Resolve MC®. A first assessment of n=2 human non-neoplastic pancreas (tail regions), resulted in 280.000 counts per ROI, segmented by Cellpose. We developed an R function to import segmentation output into a Seurat object, enabling downstream spatial transcriptomics analysis with Seurat framework(48). Additional functions for data handling are included in an R package (manuscript under development). Figure 2A illustrates the 100-gene spatial map. To avoid optical crowding from KRT19, SOX9 was used as duct cell marker, covering small to large ducts(27). Subclustering of SOX9^+^ annotated ductal clusters resulted in five subclusters (0–4)(Figure 2A-C) that were mapped back to the tissue (Figure 2D). Cluster 0, expressing high CFTR, mapped to small collecting duct cells between acini, similar to murine pancreas(27). Clusters 1 and 3 —both high in KRT5— mapped to large ducts while cluster 2 made up the rest of the duct. The small cluster 4, along with cells from clusters 0 and 2, localized to structures reminiscent of ductal invaginations or glands. The PanCK^+^ KRT5^+^ (supra)basal cell signature (Figure 1) was confined to clusters 1 and 3, albeit with distinct levels of expression; cluster 3 showed highest TP63 and traditional basal markers (KRT15, S100A2, KRT5, AHNAK2, MYC, CSPG4, SERPINB5), while cluster 1 expressed MUC4, MUC16 and elevated ductal markers LCN2 and SOX9 (Figure 2E)(27). In the spatial plot, cluster 3 KRT5^+^KRT15^+^TP63^+^ cells were residing at the basal side of the duct, while the cluster 1 MUC4^+^ cells faced the lumen (indicated by asterisk, Figure 2F). Hence, from here on, we called the TP63^+^ basally positioned cell cluster 3 “BAS” and the cells from cluster 1 “LUM-B” (Luminal-B, with B referring to supra-<u>B</u>asal). Indeed, by immunostaining for MUC4 and ΔNp63, as respective LUM-B and BAS markers, LUM-B were always apically of the BAS, being less than 50 micrometer apart (Figure 2G). Most luminal cells in the duct (cluster 2) were called “LUM-A”. Cells from <u>c</u>ollecting ducts (cluster 0) were called LUM-<u>C</u> and cells unique to the sporadic ductal invaginations or ductal glands, LUM-D (cluster 4, Figure 2H). These cell populations were validated by Resolve MC® in n=2 additional pancreata (Supplemental Figure 3), albeit without any intact ductal glands, hence no LUM-D cluster was defined.

**Figure 2:**
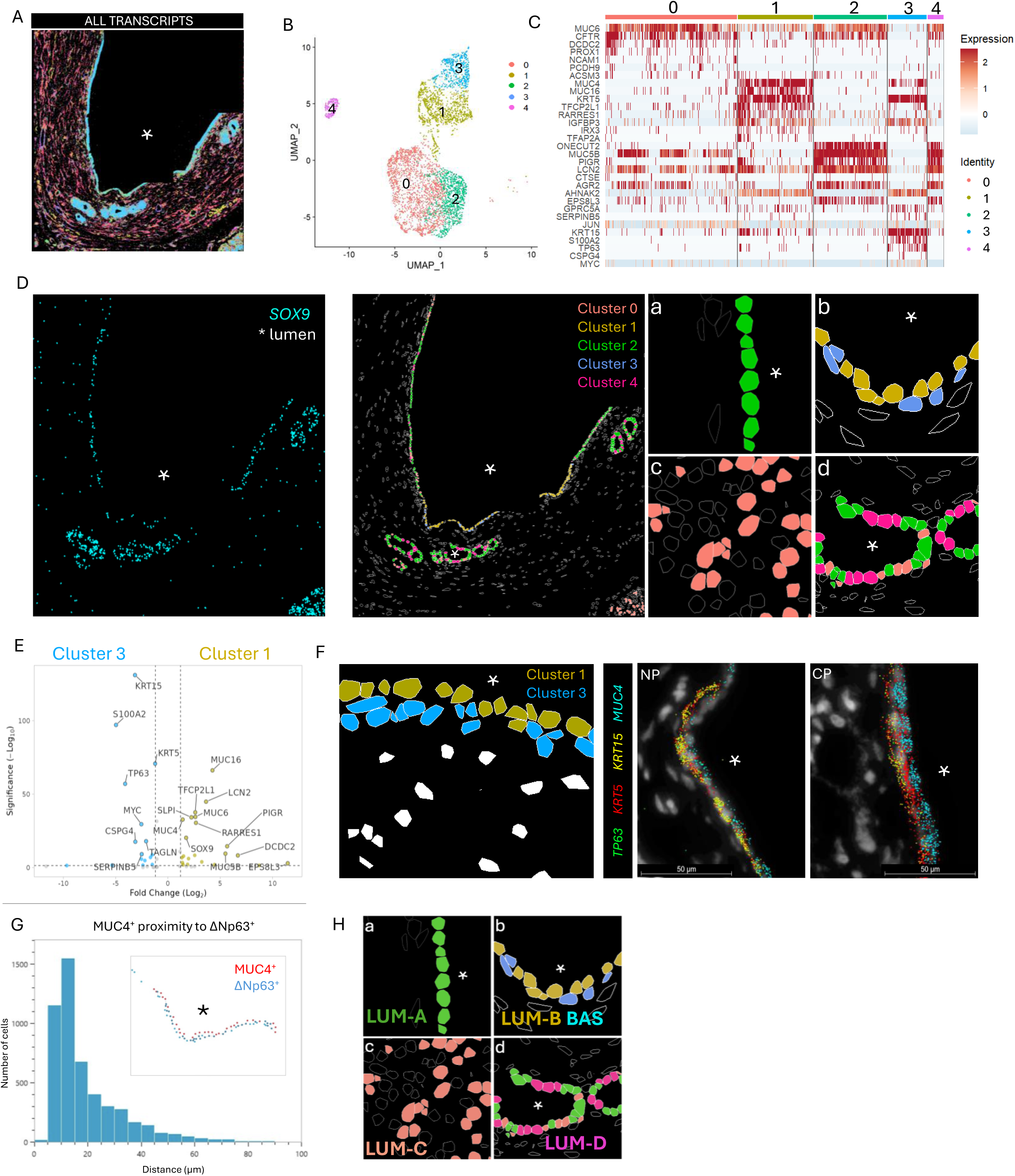
The non-neoplastic human pancreatic duct system is a stratified epithelium with a basal and four luminal populations. (A) Spatial plot of non-neoplastic pancreas region showing all one hundred transcripts of Resolve MC®. (B) Clustered UMAP of the subsetted SOX9^+^ duct populations. (C) Heatmap demonstrating top marker genes for every cluster from panel B using the FindAllMarkers function. Wilcoxon rank sum test was used for statistical analysis with min.diff.pct= 0.2. (D) Spatial plot showing the ductal system by SOX9 expression and the location of the clusters from panel B. Panels a-d are higher magnification insets of the overview plot. (E) Volcano plot showing differentially expressed genes between the two KRT5^+^ clusters, i.e. cluster 3 and cluster 1. Parameters for statistical analysis were logfc.treshold = 1, min.pct = 0, p-value < 0.05. (F) Spatial plot of cluster 1 and 3 to illustrate their location and expression of four marker genes (TP63, KRT5, KRT15, MUC4). (G) MUC4-ΔNp63 proximity analysis, i.e. distance in micrometer of MUC4^+^ cells to ΔNp63^+^ cells. (H) Spatial plot same as in panel D a-d while now using the newly proposed basal-luminal cell nomenclature as explained in main text. Asterisk labels lumen of the duct.

In summary, we identified five subsets of cells within human pancreatic ducts, where KRT5^+^ΔNp63^-^LUM-B reside uniquely superjacent to KRT5^+^ΔNp63^+^ BAS.

### Mucin expression can discriminate the basal and luminal populations

Since we discovered a population in non-neoplastic pancreas that expressed MUC4 and MUC16, we decided to further focus on mucin RNA expression patterns in the clusters (Supplemental Figure 4A-C); MUC5B discriminated best for LUM-A, MUC4 and MUC16 for LUM-B. BAS showed minimal MUC4 expression.

This RNA expression was validated by immunostainings (Figure 3A-F, Supplemental Figure 5) in a larger cohort of non-neoplastic tissue (n=50 NP, n=22 CP with each n=9 sections) of which n=15 NP and n=4 CP comprised LUM-D, n=32 NP and n=12 CP showed BAS with superjacent LUM-B. LUM-A expressed MUC5B in intra/interlobular ducts and the main duct (Figure 3A, Supplemental Figure 6A-B). LUM-B expressed MUC4 (Figure 3B) and MUC16 (Figure 3C). LUM-C, marked by CFTR (Figure 3D), expressed MUC6 (Figure 3E). Notably, LUM-B expressed membrane-bound mucins (MUC4, MUC16), whereas LUM-A and LUM-C expressed gel-forming mucins (MUC5B, MUC6), suggesting functional differences. LUM-D co-expressed MUC6 and MUC5B (Figure 3F). BAS lacked protein expression of all assessed mucins (Figure 3B-C), including MUC4. This mucin expression pattern was quantified in an additional cohort of n=9 NP where large ducts and intercalated ducts could be assessed in parallel in head, body and tail of the pancreas. The large ducts are mainly composed of cells expressing MUC5B^+^ LUM-A (75.5± 10.5% with 6.5± 6% that co-express MUC5B and MUC6) and contain 10.6± 8.9% MUC4^+^ LUM-B and <1% MUC6 only positive cells (Figure 3G, Supplemental Figure 6C). In contrast, the intercalated ducts contain 90.6±4.4% MUC6^+^ LUM-C (Figure 3G, Supplemental Figure 6D). Pancreatic ductal glands contain 72.6±12.2% MUC5B^+^MUC6^+^ LUM-D cells (Figure 3G, Supplemental Figure 6E). Additionally, ΔNp63^+^ cells made up 5.8±5.9% of the large ducts (Figure 3G). Overlap between the mucins was negligible, except for 8.7±6.0% of MUC5B^+^ cells in large ducts expressing MUC6 (Figure 3H-J). ΔNp63^+^ cells rarely expressed mucins (2.3±1.3%). The mucin expression patterns did not show regional differences between head, body and tail (Supplemental Figure 6F-H).

**Figure 3:**
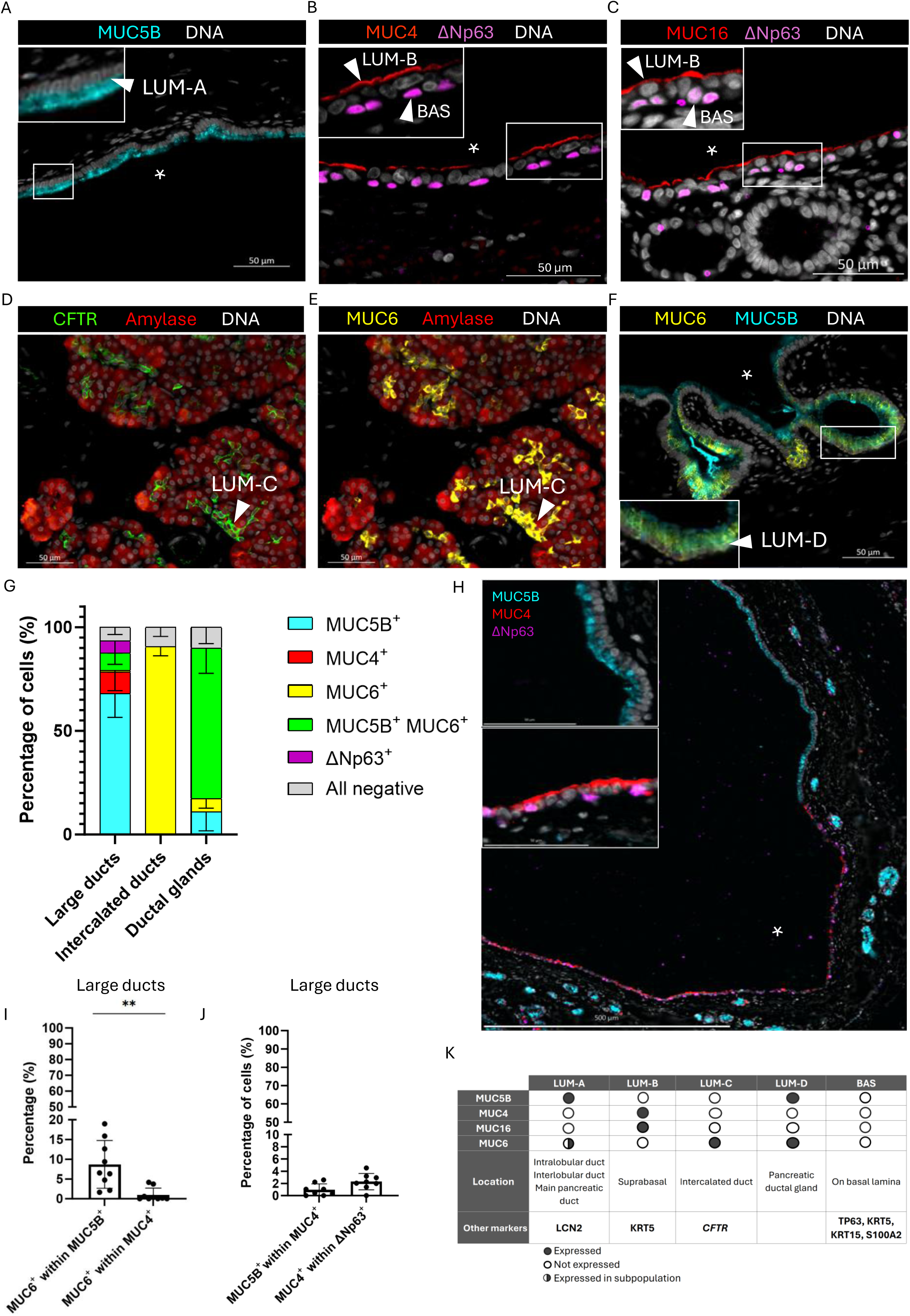
Mucin expression can discriminate the basal and luminal populations. (A-F) Immunofluorescent stainings of the different markers used to distinguish luminal and basal populations, namely (A) MUC5B, (B) MUC4, KRT17, ΔNp63, (C) MUC16 and ΔNp63, (D) CFTR and Amylase, (E) MUC6 and Amylase, (F) MUC6 and MUC5B. White arrowheads indicate the different populations. (G) Quantification of population markers in large ducts, intercalated ducts and ductal glands, n=9. (H) Representation of non-neoplastic pancreatic duct with regions containing LUM-A and with BAS + LUM-B analyzed with Comet™. (I) Quantification of overlap between MUC6 - MUC5B and MUC6 – MUC4 in large ducts. (J) Quantification of overlap between MUC5B – MUC4 and MUC4 – ΔNp63 in Comet™ analysis (K) Classification of luminal and basal populations with their specific mucin expression patterns, corresponding location and additional markers. Genes in bold have also been confirmed at the protein level in Figure 3A-F and Supplemental Figure 7D. Lumen is indicated by asterisk.

These findings, along with other antibody-validated markers (from Figure 2C-H and Supplemental Figure 6I), are summarized in Figure 3K as a tool for classification of ductal cell types. No unique markers for LUM-D were identified beyond MUC5B and MUC6 co-expression in the current gene panel.

Mucin profiles thus distinctly mark ductal cell populations, with membrane-bound mucins in LUM-B, gel-forming mucins in LUM-A/C/D, and absence in BAS, reflecting functional divergence.

### BAS and LUM-B signatures associate with basal-like subtypes and poor survival

We assessed basal/luminal gene expression in a cohort of n=276 treatment-naïve PDAC patients(4), excluding LUM-D due to lack of specific markers. High expression of LUM-B and BAS markers was associated with significantly worse disease-free survival, while LUM-C markers correlated with better prognosis (Figure 4A).

**Figure 4:**
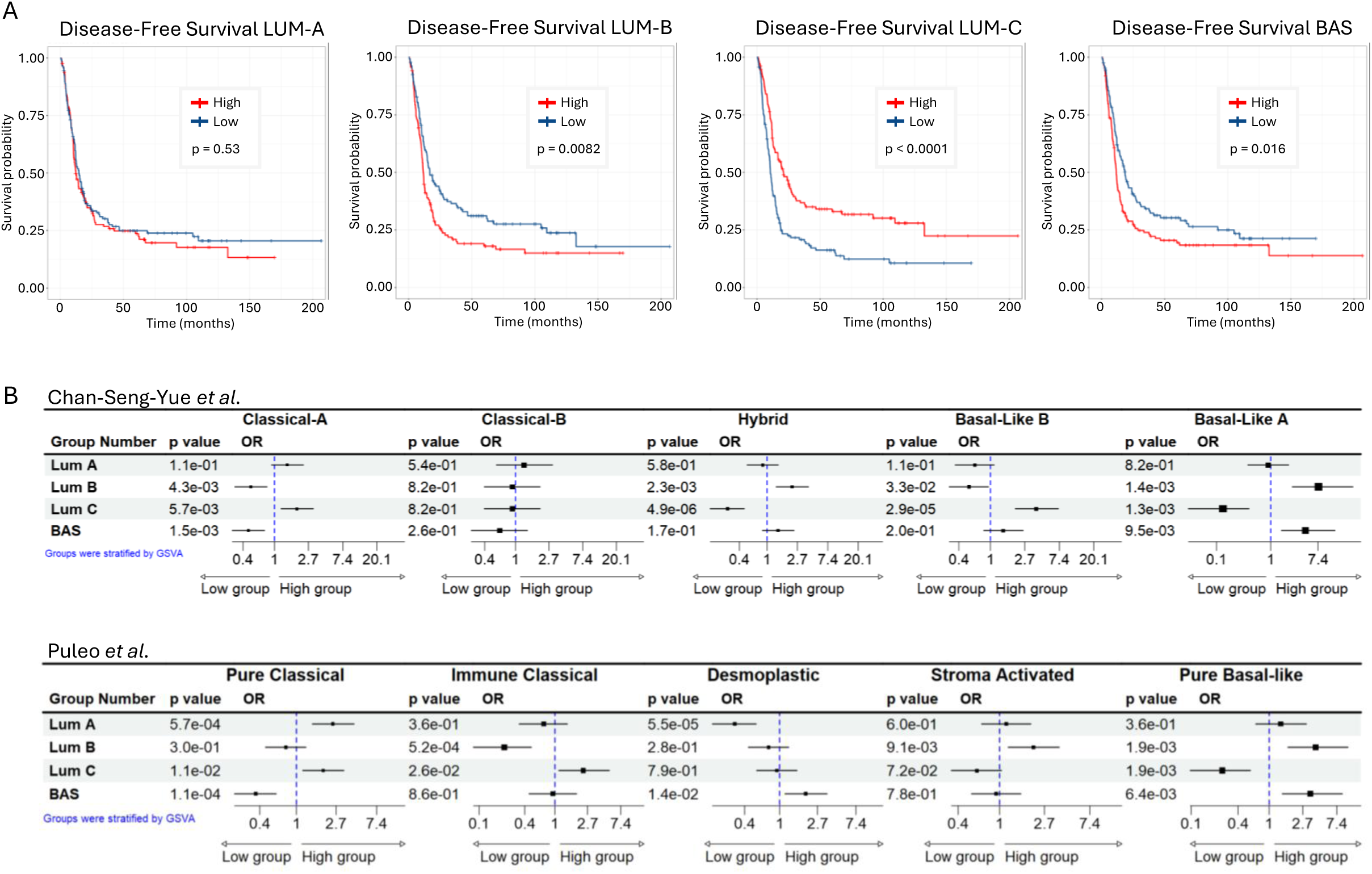
BAS and LUM-B signatures associate with basal-like subtypes and poor survival. (A) Kaplan-Meier plots depicting the disease-free survival of a cohort of n=276 treatment-naive PDAC patients. Red lines represent patients with high expression of the signature, blue lines represent low expressing patients. Groups were based on median value. Log rank test was used, and statistically significant changes were accepted at p-value < 0.05. (B) Forest plots showing the odds ratio (OR) for luminal and basal signatures in patient groups with specific subtypes of Chan-Seng-Yue et al.(7) and Puleo et al.(4). Low and high groups were based on median value.

Comparison with previously established bulk transcriptomic subtypes of PDAC(4,7), revealed that LUM-A and LUM-C profiles associated with CL-A and Pure CL subtypes (associated with better outcomes)(7), whereas LUM-B and BAS profiles associated with BL subtypes (BL-A and Pure BL) (Figure 4B) that have worse survival(4,7,49).

### Basal-like PDAC show fragmented LUM-B and BAS signatures and a mixed spatial pattern

To assess how native ductal pancreas populations were represented in cancer, the 100-plex Resolve MC® was used (n=3 PDAC of which one is CL and two are BL, n=1 ASCP). ASCP cells formed two distinct clusters (0 and 1), while PDAC cells distributed across clusters 2–7 (Figure 5A). When assessing two markers (from Figure 3K) per cell population, at most one of two LUM-A/B markers was expressed in PDAC clusters, while they were co-expressed in cluster 1 of ASCP (Figure 5B). LUM-C markers were expressed in few clusters in PDAC and absent in ASCP. Co-expression of BAS markers TP63 and KRT5 was found in the ASCP cluster 0 (Figure 5B), PDAC cells were rarely positive for TP63 and did not co-express KRT5. Together this indicated that PDAC seems to have a fragmented representation of these native cell population markers while ASCP reproduced the BAS markers.

**Figure 5:**
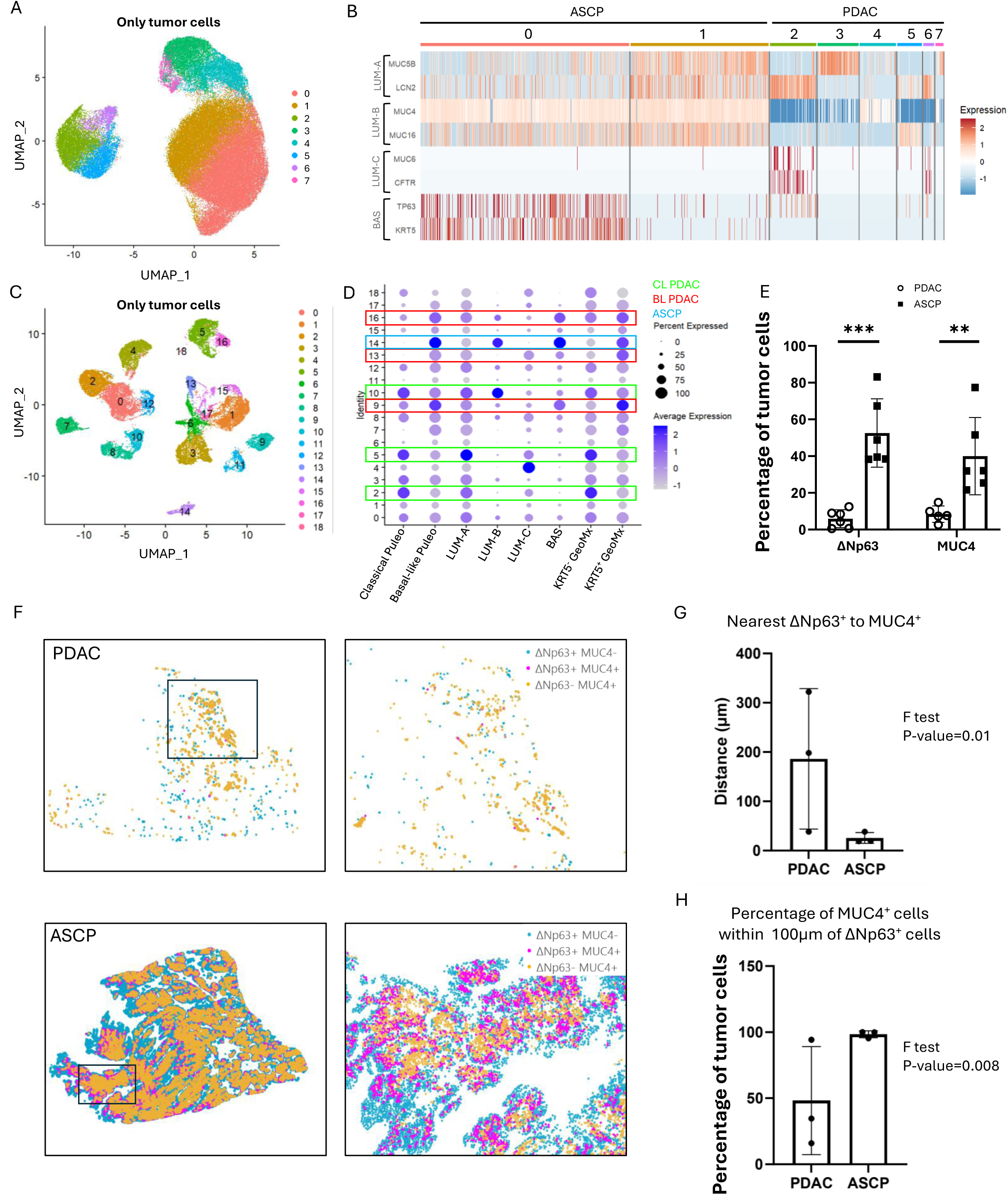
Basal-like PDAC shows fragmented BAS and LUM-B signatures and a mixed spatial pattern. (A) Clustered UMAP of only tumor cells of Pancreatic Ductal Adenocarcinoma (PDAC) and Adenosquamous Carcinoma of the Pancreas (ASCP) samples analyzed by Resolve MC® (B) Heatmap showing the different ductal classification markers (LUM-A = MUC5B and LCN2, LUM-B = MUC4 and MUC16, LUM-C = MUC6 and CFTR, BAS = TP63 and KRT5). Wilcoxon rank sum test was used for statistical analysis with min.diff.pct= 0.2. (C) Clustered UMAP of the merged scRNA-seq object of Chijimatsu et al.(50) and Zhao et al.(51) containing only tumor cells. (D) Dotplot of merged scRNA-seq object of Chijimatsu et al.(50) and Zhao et al.(51) showing expression levels of transcriptomic profiles of PDAC subtypes as defined by Puleo et al.(4) and the healthy luminal and basal ductal profiles. UCell was used to implement the profiles into the Seurat object. (E) Quantification of immunofluorescently stained ΔNp63^+^ BAS cells and MUC4^+^ LUM-B cells in PDAC and ASCP in the squamous nests using HALO® image analysis software (Indica Labs, New-Mexico, USA). Data is represented as means with standard deviation. Unpaired t-test was used to determine statistical significance. *p < 0.05, **p < 0.01, ***p < 0.001. n=6 PDAC and n=6 ASCP. (F) Spatial plot showing BAS (blue), intermediate (pink) and LUM-B (yellow) states in ASCP and PDAC. (G) Nearest neighbour analysis showing the average shortest distance from a BAS to a LUM-B cell in PDAC and ASCP. F test was used to compare variances. n=3 PDAC and n=3 ASCP. (H) Percentage of LUM-B cells located within 100µm of a BAS cell in PDAC and ASCP. F test was used to compare variances. n=3 PDAC and n=3 ASCP.

A fragmented gene signature was indeed observed in a broader gene set from 6 independent scRNA-seq studies47 (n=58 PDAC) with ASCP sample48 integrated; Eighteen tumor clusters (Figure 5C) were stratified(4) into CL and BL subtype (Figure 5D). In BL PDAC clusters 9, 13 and 16, the BAS, LUM-B and KRT5^+^ signatures were indeed fragmented, while the ASCP cluster 14 showed coherent expression (Supplemental Figure 7A). The observations were further confirmed in a single nucleus RNAseq dataset containing PDAC and ASCP samples (Hwang et al.(8), Supplemental Figure 7B cluster 5= ASCP). At the protein level, ΔNp63⁺ cells were significantly less frequent in PDAC than in ASCP (5.9 ± 4.9% vs 52.6 ± 18.6%; p = 0.0001) and rarely co-expressed KRT5, indicating fragmented BAS marker expression in PDAC compared with ASCP (Figure 5E; Supplemental Fig. 8A-B. Similarly, the LUM-B marker MUC4 was markedly reduced in PDAC (8.4 ± 4.4%) relative to ASCP (40 ± 21%; p = 0.0098).

Next to fragmentation, in the spatial plots, PDAC showed a very mixed spatial expression pattern of mucins with low TP63, while ASCP showed strong expression of MUC4 and minor MUC5B and TP63 (Supplemental Figure 9). Indeed, only in ASCP the ΔNp63⁺ and the MUC4⁺ populations are arranged in a spatial pattern, with ΔNp63⁺ cells in the periphery, MUC4⁺ cells located in the centre and a double-positive population in between (Figure 5F). A pronounced spatial coupling is observed with the nearest distance of ΔNp63⁺ cell to a MUC4^+^ cell being 22.10µm in ASCP compared to 186µm in PDAC (Figure 5G). Nearly all MUC4⁺ cells in ASCP are positioned within 100µm of a ΔNp63⁺ cell, whereas this is only half in PDAC (Figure 5H). PDAC has a significantly more heterogeneous pattern compared to the consistent ASCP pattern (Figure 5G-H, with F-test to compare the variances).

In summary, the spatial heterogeneity of the native ductal cell population is best reflected in ASCP while fragmented and mixed in BL PDAC.

### ASCP display functionally different, spatially unmixed LUM-B and BAS cells in the tumor nests

The above data warranted a closer look into the tumor cell populations of ASCP. Subclustering the ASCP cells from Figure 5A resolved two populations (Figure 6A), with respectively BAS genes (including TP63)(cluster 0) and LUM-B genes (cluster 1)(Supplemental Figure 10A). Pathway analysis using the MSigDB “C2: curated gene sets” indicated that the BAS cluster was enriched for proliferation, cell cycle, metastasis and epithelial-to-mesenchymal transition (EMT), whereas the LUM cluster displayed luminal, keratinization, and epithelial differentiation signatures (Figure 6B).

**Figure 6:**
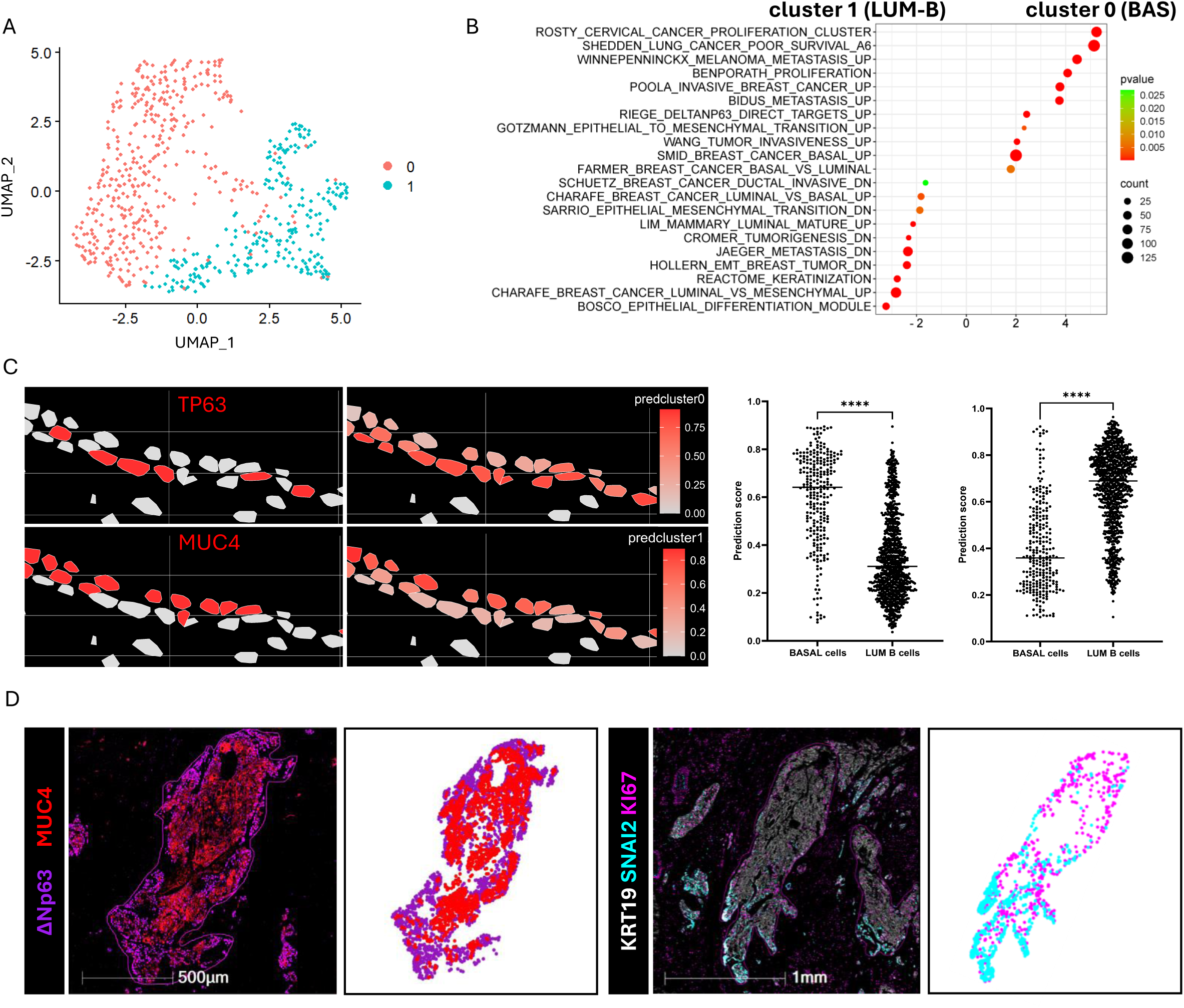
ASCP displays a spatially unmixed pattern of LUM-B and BAS cells consistent with the native tissue. (A) Clustered UMAP of subsetted ASCP cells (cluster 14) from Figure 5A. (B) GSEA analysis applied to data from panel A using curated genesets (c2.all.v2024.1.Hs.symbols.gmt). (C) Projection of reference cell signatures obtained from the clusters of panel A, i.e. cluster 0/BAS and cluster 1/LUM-B, on sections of NP analyzed by Resolve MC®, as described in Stuart et al.(52). Statistical significance was determined using an unpaired t-test. *p < 0.05, **p < 0.01, ***p < 0.001, ****p<0.0001. (D) Immunofluorescent stainings for BAS (ΔNp63), LUM-B (MUC4), proliferation (MKI67) and epithelial-to-mesenchymal transition (SNAI2) markers in ASCP.

We then performed cell type classification using an integrated reference(52), where we projected the two ASCP profiles from Figure 6A back onto a region of NP analyzed by Resolve MC® which confirmed that BAS cluster 0 had a higher prediction score in the TP63^+^ BAS of NP (mean= 0.65) compared to the LUM-B cluster 1 (mean= 0.35), while this tumoral LUM-B cluster scored higher in the NP LUM-B cells (mean= 0.65 versus 0.35)(Figure 6C), underscoring their resemblance to the native tissue.

The same spatial pattern was seen with Resolve MC® as in Figure 5F (Supplemental Figure 10B) where BAS were at the periphery of squamous nests while LUM-B genes projected to the center. Immunostainings in a larger cohort (n=14 ASCP) validated this spatial unmixing, with ΔNp63^high^ BAS in the outer rim of tumor nests, and MUC4^+^MUC16^+^ LUM-B occupying the center (Figure 6D, Supplemental Figure 10C) and confirmed that the proliferation marker KI67^+^ and EMT markers SNAI2 and S100A2 were higher in outer ΔNp63^high^ regions (Figure 6D, Supplemental Figure 10C, n=14). Indeed, among the KI67^+^ cells, 33.48 ± 11% expressed only ΔNp63, 14.8 ± 10.4% only MUC4 and 25.8 ± 17% both, indicating 59% of proliferative cells express ΔNp63 in ASCP (Supplemental Figure 10D, n=6). We noted a difference with LUM-B MUC4 and MUC16 shed into lumens in ASCP (Supplemental Figure 10C, and E as an H&E representation of ASCP) versus the membrane localization in non-neoplastic tissue (Figure 3B,C).

We also assessed a set of markers from a recent report on squamoid, basaloid, and mesenchymal cell states in pancreatic cancer(8). Basaloid markers overlapped with BAS regions, while squamoid markers aligned with LUM-B regions (Supplemental Figure 11A), and mesenchymal vimentin was confined to the outer rim.

ASCP thus recapitulates spatially unmixed BAS and LUM-B with the former population proliferating and undergoing EMT at the outer rim of the tumor nests.

### ΔNp63 drives cell plasticity to BAS, conserved from the native tissue to cancer

With TP63 identified as a key regulator in the KRT5^+^ cells (Figure 1G), and the new knowledge that this group consists of BAS and LUM-B (Figure 2), we (re-)evaluated the role of ΔNp63–a known lineage-specifier of basal cells(10,11,18)–from this new perspective. ΔNp63-knockdown (KD) in the main duct-derived KRT5^+^ HPDE cell line(18) reduced BAS markers (TP63, S100A2) and induced LUM-B markers (MUC4, MUC16)(Figure 7A). Additionally, overexpression of ΔNp63 in organoid cultures from the native tissue (derived from a NP exocrine cell fraction, Figure 7B, Supplemental File 3) induced the BAS gene signature including TP63, S100A2, KRT17, TFAP2A with minor induction of LUM-B markers (MUC4, MUC16). Cell type geneset (MSigDB) analysis showed downregulation of the BAS genesets and upregulation of luminal genesets in HPDE with ΔNp63 KD (Figure 7C) while in the organoid overexpression experiment more basal and squamous genesets were enriched at the expense of (fetal) duct cells (Figure 7D).

**Figure 7:**
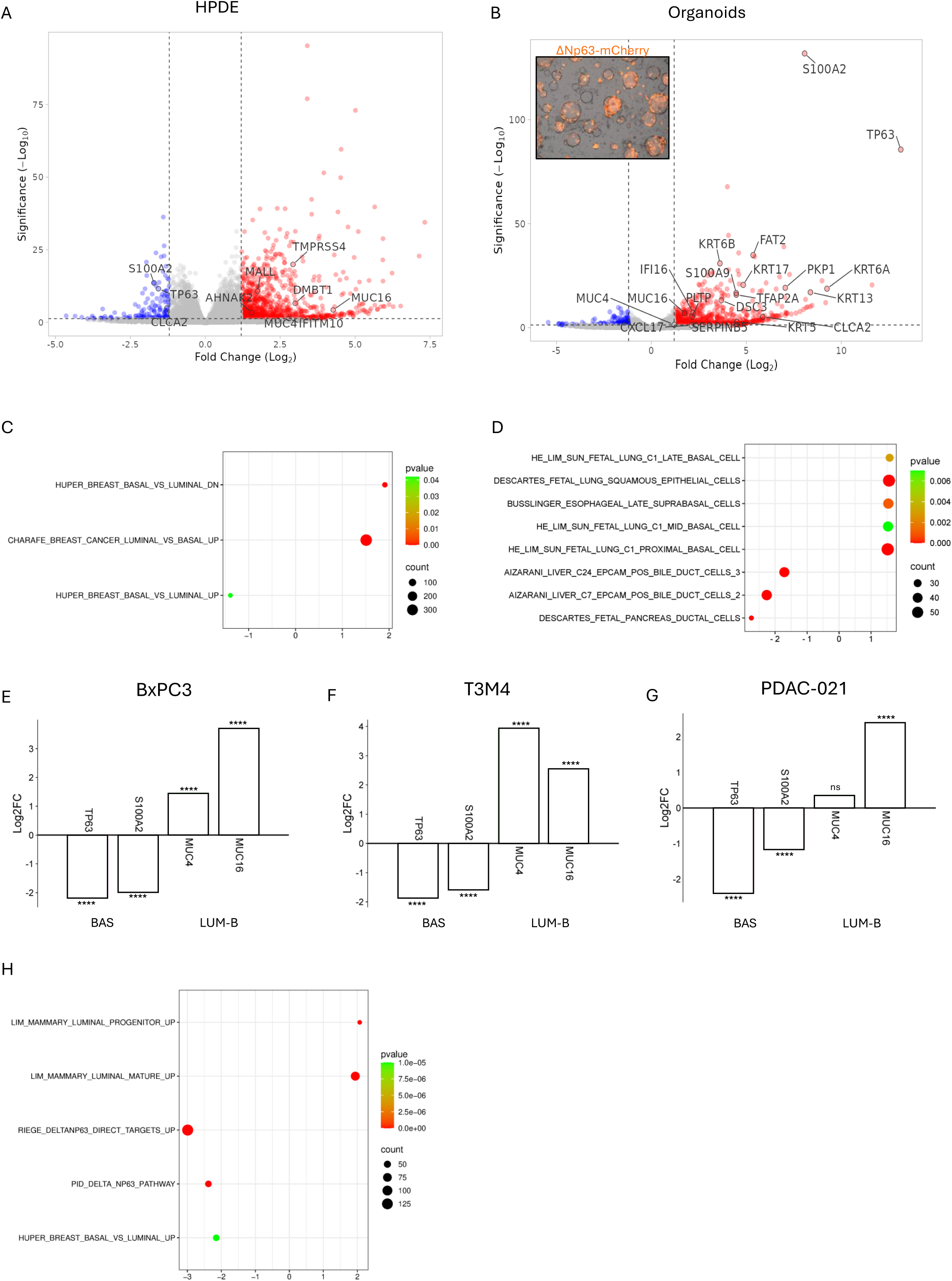
ΔNp63 drives cell plasticity to BAS, conserved from the native tissue to cancer. (A) Volcano plot showing Human Pancreatic Ductal Epithelial (HPDE) cells with and without knockdown of ΔNp63. Genes shown overlap with the GeoMx® data of Figure 1C. (B) Lentiviral transduction of exocrine-derived organoids in passage 2 for introduction of ΔNp63-mCherry. Volcano plot showing transduced versus control organoids. Genes shown overlap with the GeoMx® data of Figure 1C. Logfc.treshold = 1, min.pct = 0, p-value < 0.05. (C) GSEA analysis applied to data from panel A with C2 of MSigDB. (D) GSEA analysis applied to data from panel B with C2 of MSigDB. (E-G) Log2FC of BAS and LUM-B genes after knockdown of ΔNp63 in (E) BxPC3, (F) T3M4 and (G) PDAC-021. ns= not significant, *p < 0.05, **p < 0.01, ***p < 0.001, ****p<0.0001. (H) GSEA analysis on squamous cell lines after ΔNp63 knockdown using C2 of MSigDB.

Subsequent analysis in basal-like/squamous pancreatic cancer cell lines (BxPC3 cells(10), T3M4 and PDAC-021(53)) confirmed that ΔNp63 KD resulted in an enrichment for luminal signatures, in particular LUM-B genes like MUC4 and MUC16, while genes aligned with BAS (TP63 and S100A2) were downregulated (Figure 7E-G). Pathway analysis (MsigDB, C2) confirmed the shift from a basal to a luminal state (Figure 7H). This cell plasticity agreed with the transition from ΔNp63^+^ cells to MUC4^+^ cells and an intermediate zone in Figure 5F.

These findings establish ΔNp63 as a conserved lineage regulator that drives reversible luminal–basal plasticity, promoting a BAS identity from native pancreatic epithelium through to basal-like/squamous pancreatic cancer.

## Discussion

Investigating normal human pancreatic duct cells and their malignant counterparts offers reciprocal insights—a rationale that underpinned our study. While traditionally regarded as a uniform simple epithelium, the pancreatic duct is more heterogeneous than previously appreciated. Building on our earlier work, where we identified a rare (1.6% of duct cells) KRT5⁺ΔNp63⁺ basal cells(18), we now defined five spatially and transcriptionally distinct ductal populations—four luminal (LUM-A-D) and one basal (BAS)— through advanced spatial transcriptomic technologies.

Our study provided new biological insights: GeoMx® profiling of groups of KRT5⁺ cells revealed strong transcriptomic similarities with (supra)basal cells from lung and prostate(32,54) but lacked single-cell resolution. Spatial mapping with Resolve MC® further discriminated the KRT5^+^ group into BAS cells and suprabasal LUM-B cells, distinct from neighboring LUM-A cells, yet, the signatures remained incomplete due to the limited custom gene panel. Given the absence of specific purification markers and the limitations of single-cell and single-nucleus RNA-seq technologies—including shallow sequencing depth and poor capture of rare cell types—new spatial transcriptomic platforms offering full-transcriptome, single-cell resolution hold promise for resolving complete molecular signatures and lineage trajectories. Nevertheless, we identified mucins as distinguishing markers providing new insights. LUM-B cells express MUC4 and MUC16, previously reported not to be present in the healthy pancreas and restricted to aggressive pancreatic cancer(35–38,55–57). We thus uncover a cell type in the native tissue that warrants further investigation, next to the enigmatic BAS with their presumed stem cell potential —as suggested by the gene expression analysis and role in other organs. The unique positioning of LUM-B cells superjacent to BAS cells suggests a potential lineage relationship, reminiscent of basal-to-suprabasal transitions observed in other organs during epithelial repair(28,58). Studies on the cellular dynamics and the regulation of ΔNp63 in the adult (human) pancreas highlight the necessity for good experimental models. While we reported that ΔNp63 is transiently expressed in murine pancreatic development(59), it is absent in the adult mouse pancreas(18), hampering more mechanistic work. While our study did not directly address this, it is noteworthy that LY6D—a marker for rare, Wnt-responsive progenitor-like cells in the murine pancreas(26,27) —is a LUM-B marker. In human, we occasionally observed BAS in ductal invaginations or ductal gland structures hypothesized to function as progenitor niches, from which cells migrate to larger ducts upon injury(60). The development of dedicated human experimental models is again essential to further investigate these dynamics.

Our findings are also relevant for pancreatic cancer. PDAC is currently stratified using transcriptomic subtypes (CL vs. BL). ASCP is mostly considered separate from PDAC —albeit not in all studies—, yet it lacks specific clinical guidelines and is treated as PDAC(20,21). While the term “BL” in PDAC draws from similarities with basal tumors in other organs, our findings suggest that PDAC is only loosely “BL”. In contrast, ASCP more faithfully recapitulates BAS and LUM-B identities, preserving both gene signatures and spatial organization. These results align with the recent works of Hilmi et al.(61) and Chan-Seng-Yue et al.(7), where they show that BL PDAC can exist in different types, to which our data now adds that, in comparison, ASCP seems to be in a more robust basal state. We demonstrated that ΔNp63 is a conserved driver of BAS identity and its loss facilitates a basal to luminal transition in both the non-neoplastic and cancer contexts. Therapeutic targeting of ΔNp63(62) has been proposed, with drugs that could potentially convert the aggressive (proliferating, EMT-undergoing) BAS to the more indolent LUM-B state(63,64). One could envision a combination approach where BAS are driven to LUM-B that on their turn would become the target of a second therapy. Potential targets are listed in Supplemental Figure 12; notably, MUC4 expression suggests potential utility of bosutinib (MUC4 inhibitor)(65) and CAR-T cells designed against specific mucins(56). It has already been demonstrated that mucin regulation can influence the subtype of tumor cells(66). Further preclinical studies are timely and can benefit from a recent murine model recapitulated the development of ASCP(67). The fact that LUM-A and LUM-C cells – the majority of duct cells— have functionally distinct mucins suggests that those cells would remain unaffected during treatment.

Finally, our refined classification of pancreatic ductal cell populations raises new questions regarding the cell of origin in pancreatic cancer. Both acinar and ductal cells have been proposed as PDAC progenitors(2), but our data suggest this framework should be revisited to account for ductal heterogeneity and to distinguish PDAC from ASCP.

In summary, our spatial transcriptomic analysis of the human pancreas improves biological discrimination between pancreatic tumor types—particularly BL PDAC and ASCP—and provides a foundation for future studies of pancreatic biology in health and disease.

## Abbreviations

PDAC: Pancreatic Ductal Adenocarcinoma
ASCP: Adenosquamous Cancer of the Pancreas
NP: Normal human Pancreas
CP: Chronic Pancreatitis
BAS: Basal cell
LUM: Luminal cell

## Disclosures

All authors declare no competing interests.

## Author contributions

<u>Conceptualization:</u> JL VDB, IR. <u>Data curation:</u> JL VDB, MVDV. <u>Formal analysis:</u> JL VDB, MVDV. <u>Funding acquisition:</u> IR. <u>Investigation:</u> JL VDB, MVDV, EM, OAS, ZM, KC, JBr, SVL, PL. <u>Methodology:</u> JL VDB, MVDV. <u>Project administration:</u> JL VDB, IH, IR. <u>Software:</u> MVDV. <u>Resources:</u> MN, NM, Emo, CB, LV, JN, JF, NS, IR. <u>Supervision:</u> TA, MR, JL VL, JB, IH, IR. <u>Writing original draft:</u> JL VDB, IR.

## Data Transparency Statement

Spatial data will be made available on VUB-IR and R script on GitHub.

## Acknowledgements

The authors thank the Visual and Spatial Tissue Analysis (VSTA) core facility and specifically express their appreciation for the high-quality efforts of VSTA technician Emmy De Blay and lab technician Veerle Laurysens. The authors also thank the team at Resolve Biosciences, whose Molecular Cartography® technology was utilized in this study as part of a collaboration.

---

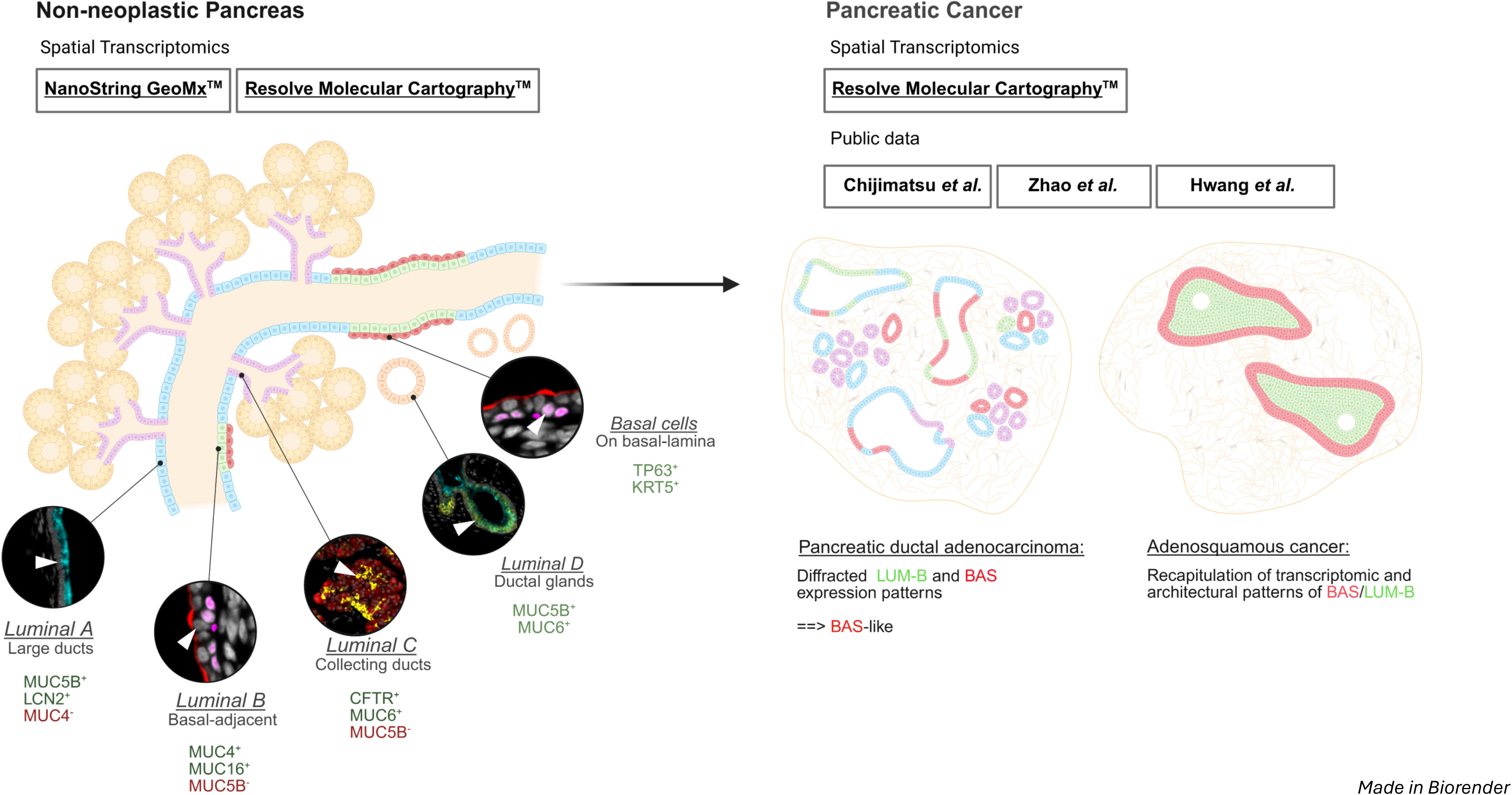

## Notes

**Grant Support:** IR was awarded research grants from Research Foundation Flanders (FWO, G0A7322N) and Wetenschappelijk Fonds Willy Gepts (WFWG). J-L VDB is a recipient of a PhD Fellowship of the FWO (Grant ID 11L7724N) and a finishing grant of Kom Op Tegen Kanker (KOTK)(Grant ID KotK_VUB/2025/14284). EMo acknowledges the support of grant PI22/00334 from Instituto de Salud Carlos III, co-financed by the European Regional Development Fund (ERCF). IH was financially supported by the bequest of Ms. Esther Desmedt and Ms. Irma Noë, and FWO, G0A7322N. SVL was financially supported by the Award Cancer Research - Oncology Center VUB (bequests of Ms. Esther Desmedt and Ms. Irma Noë) and Kom Op Tegen Kanker grant (KOTK_VUB/2024/13916). NM is a senior clinical investigator of the FWO (Grant ID 1801425N). JB is supported by a Foundation Against Cancer Fundamental mandate (2023-038). KC is a recipient of a PhD Fellowship of the FWO (Grant ID 1157221N).

### Competing Interest Statement

The authors have declared no competing interest.

### Summary of Updates

A quantification of mucin expression was performed on a cohort of non-neoplastic pancreata (Figure 3G-J). The conservation of spatial patterns in pancreatic cancers was quantified in Figure 5F-H. Two additional cell lines were used for functional experiments (Figure 7E-H).

