## Supplemental Figures for "A Luminal–Basal Stratification of the Native Human Pancreatic Duct is Differentially Represented in Pancreatic Cancers"

### Slide 1
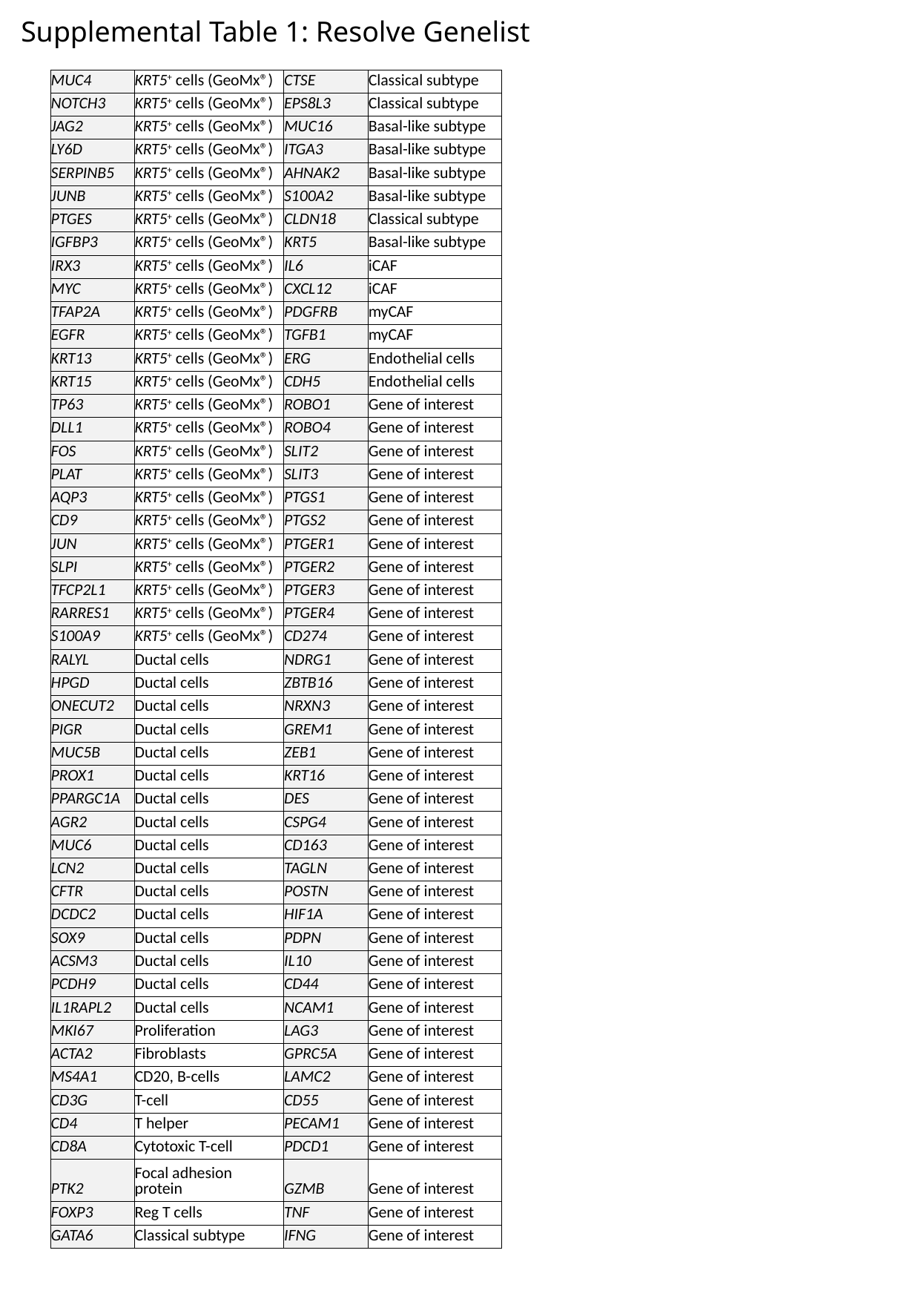

Supplemental Table 1: Resolve Genelist
| MUC4 | KRT5^+^ cells (GeoMx®) | CTSE | Classical subtype |
| --- | --- | --- | --- |
| NOTCH3 | KRT5^+^ cells (GeoMx®) | EPS8L3 | Classical subtype |
| JAG2 | KRT5^+^ cells (GeoMx®) | MUC16 | Basal-like subtype |
| LY6D | KRT5^+^ cells (GeoMx®) | ITGA3 | Basal-like subtype |
| SERPINB5 | KRT5^+^ cells (GeoMx®) | AHNAK2 | Basal-like subtype |
| JUNB | KRT5^+^ cells (GeoMx®) | S100A2 | Basal-like subtype |
| PTGES | KRT5^+^ cells (GeoMx®) | CLDN18 | Classical subtype |
| IGFBP3 | KRT5^+^ cells (GeoMx®) | KRT5 | Basal-like subtype |
| IRX3 | KRT5^+^ cells (GeoMx®) | IL6 | iCAF |
| MYC | KRT5^+^ cells (GeoMx®) | CXCL12 | iCAF |
| TFAP2A | KRT5^+^ cells (GeoMx®) | PDGFRB | myCAF |
| EGFR | KRT5^+^ cells (GeoMx®) | TGFB1 | myCAF |
| KRT13 | KRT5^+^ cells (GeoMx®) | ERG | Endothelial cells |
| KRT15 | KRT5^+^ cells (GeoMx®) | CDH5 | Endothelial cells |
| TP63 | KRT5^+^ cells (GeoMx®) | ROBO1 | Gene of interest |
| DLL1 | KRT5^+^ cells (GeoMx®) | ROBO4 | Gene of interest |
| FOS | KRT5^+^ cells (GeoMx®) | SLIT2 | Gene of interest |
| PLAT | KRT5^+^ cells (GeoMx®) | SLIT3 | Gene of interest |
| AQP3 | KRT5^+^ cells (GeoMx®) | PTGS1 | Gene of interest |
| CD9 | KRT5^+^ cells (GeoMx®) | PTGS2 | Gene of interest |
| JUN | KRT5^+^ cells (GeoMx®) | PTGER1 | Gene of interest |
| SLPI | KRT5^+^ cells (GeoMx®) | PTGER2 | Gene of interest |
| TFCP2L1 | KRT5^+^ cells (GeoMx®) | PTGER3 | Gene of interest |
| RARRES1 | KRT5^+^ cells (GeoMx®) | PTGER4 | Gene of interest |
| S100A9 | KRT5^+^ cells (GeoMx®) | CD274 | Gene of interest |
| RALYL | Ductal cells | NDRG1 | Gene of interest |
| HPGD | Ductal cells | ZBTB16 | Gene of interest |
| ONECUT2 | Ductal cells | NRXN3 | Gene of interest |
| PIGR | Ductal cells | GREM1 | Gene of interest |
| MUC5B | Ductal cells | ZEB1 | Gene of interest |
| PROX1 | Ductal cells | KRT16 | Gene of interest |
| PPARGC1A | Ductal cells | DES | Gene of interest |
| AGR2 | Ductal cells | CSPG4 | Gene of interest |
| MUC6 | Ductal cells | CD163 | Gene of interest |
| LCN2 | Ductal cells | TAGLN | Gene of interest |
| CFTR | Ductal cells | POSTN | Gene of interest |
| DCDC2 | Ductal cells | HIF1A | Gene of interest |
| SOX9 | Ductal cells | PDPN | Gene of interest |
| ACSM3 | Ductal cells | IL10 | Gene of interest |
| PCDH9 | Ductal cells | CD44 | Gene of interest |
| IL1RAPL2 | Ductal cells | NCAM1 | Gene of interest |
| MKI67 | Proliferation | LAG3 | Gene of interest |
| ACTA2 | Fibroblasts | GPRC5A | Gene of interest |
| MS4A1 | CD20, B-cells | LAMC2 | Gene of interest |
| CD3G | T-cell | CD55 | Gene of interest |
| CD4 | T helper | PECAM1 | Gene of interest |
| CD8A | Cytotoxic T-cell | PDCD1 | Gene of interest |
| PTK2 | Focal adhesion protein | GZMB | Gene of interest |
| FOXP3 | Reg T cells | TNF | Gene of interest |
| GATA6 | Classical subtype | IFNG | Gene of interest |

### Slide 2
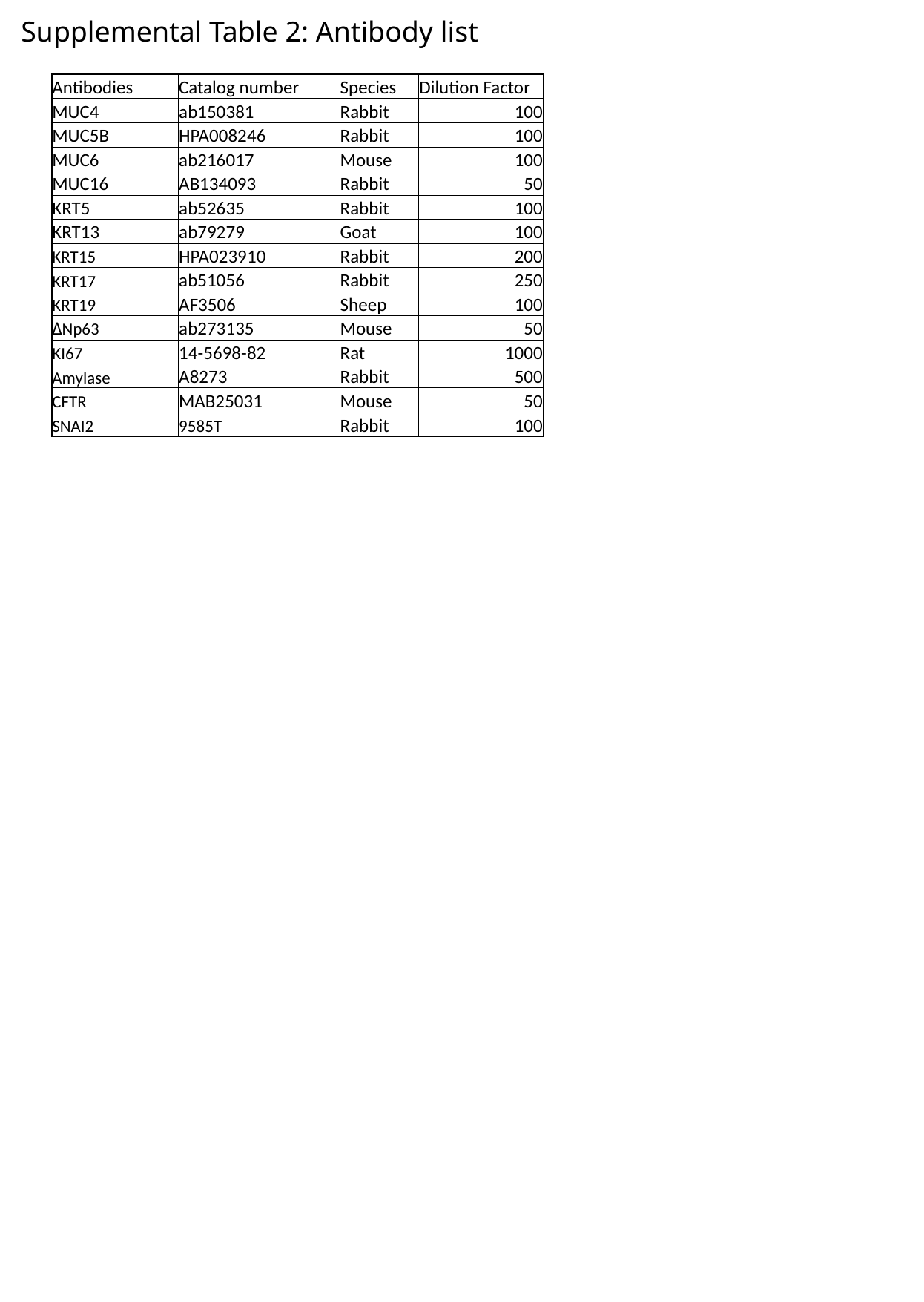

Supplemental Table 2: Antibody list
| Antibodies | Catalog number | Species | Dilution Factor |
| --- | --- | --- | --- |
| MUC4 | ab150381 | Rabbit | 100 |
| MUC5B | HPA008246 | Rabbit | 100 |
| MUC6 | ab216017 | Mouse | 100 |
| MUC16 | AB134093 | Rabbit | 50 |
| KRT5 | ab52635 | Rabbit | 100 |
| KRT13 | ab79279 | Goat | 100 |
| KRT15 | HPA023910 | Rabbit | 200 |
| KRT17 | ab51056 | Rabbit | 250 |
| KRT19 | AF3506 | Sheep | 100 |
| ΔNp63 | ab273135 | Mouse | 50 |
| KI67 | 14-5698-82 | Rat | 1000 |
| Amylase | A8273 | Rabbit | 500 |
| CFTR | MAB25031 | Mouse | 50 |
| SNAI2 | 9585T | Rabbit | 100 |

### Slide 3
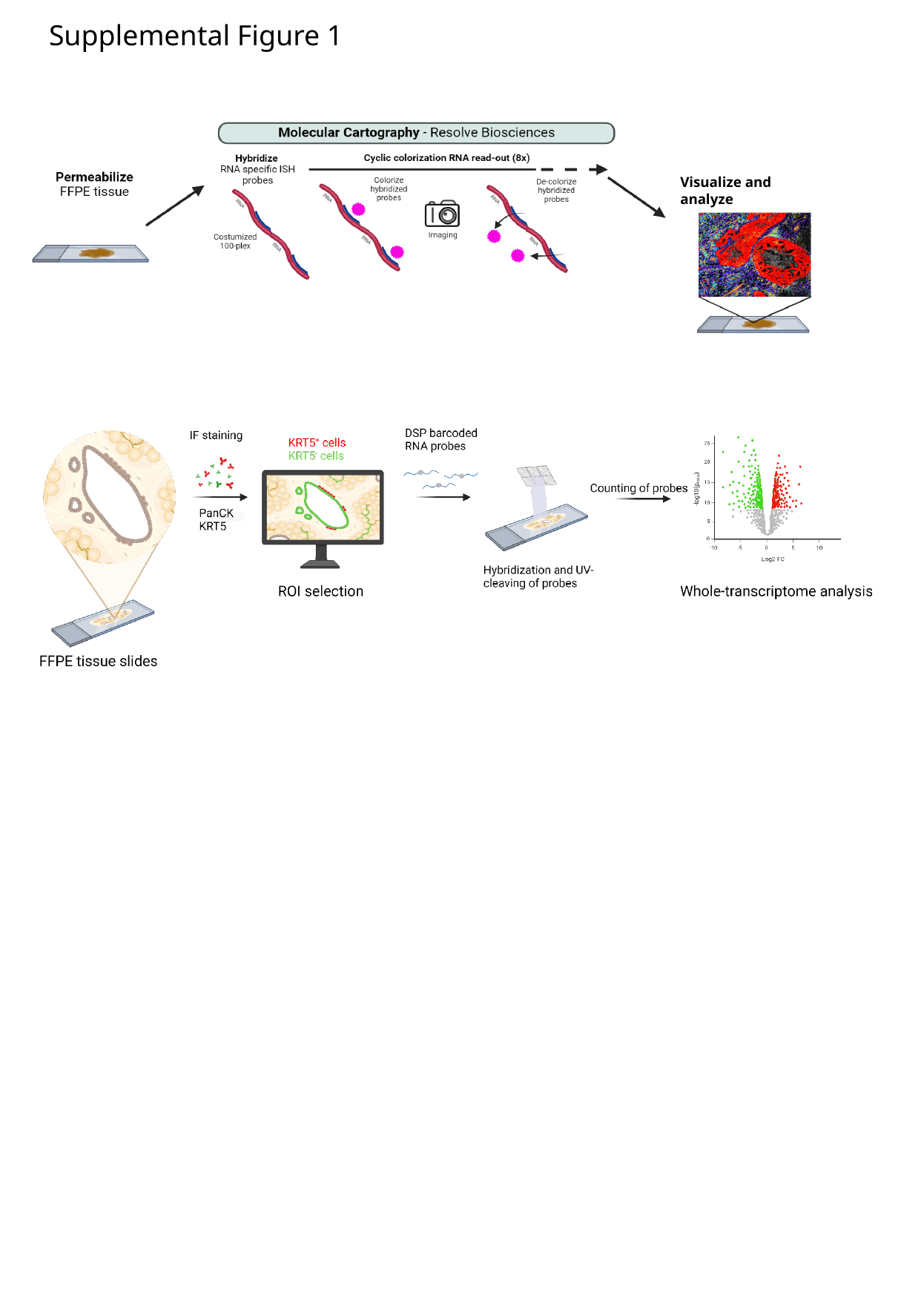

Supplemental Figure 1
Visualize and analyze

### Slide 4
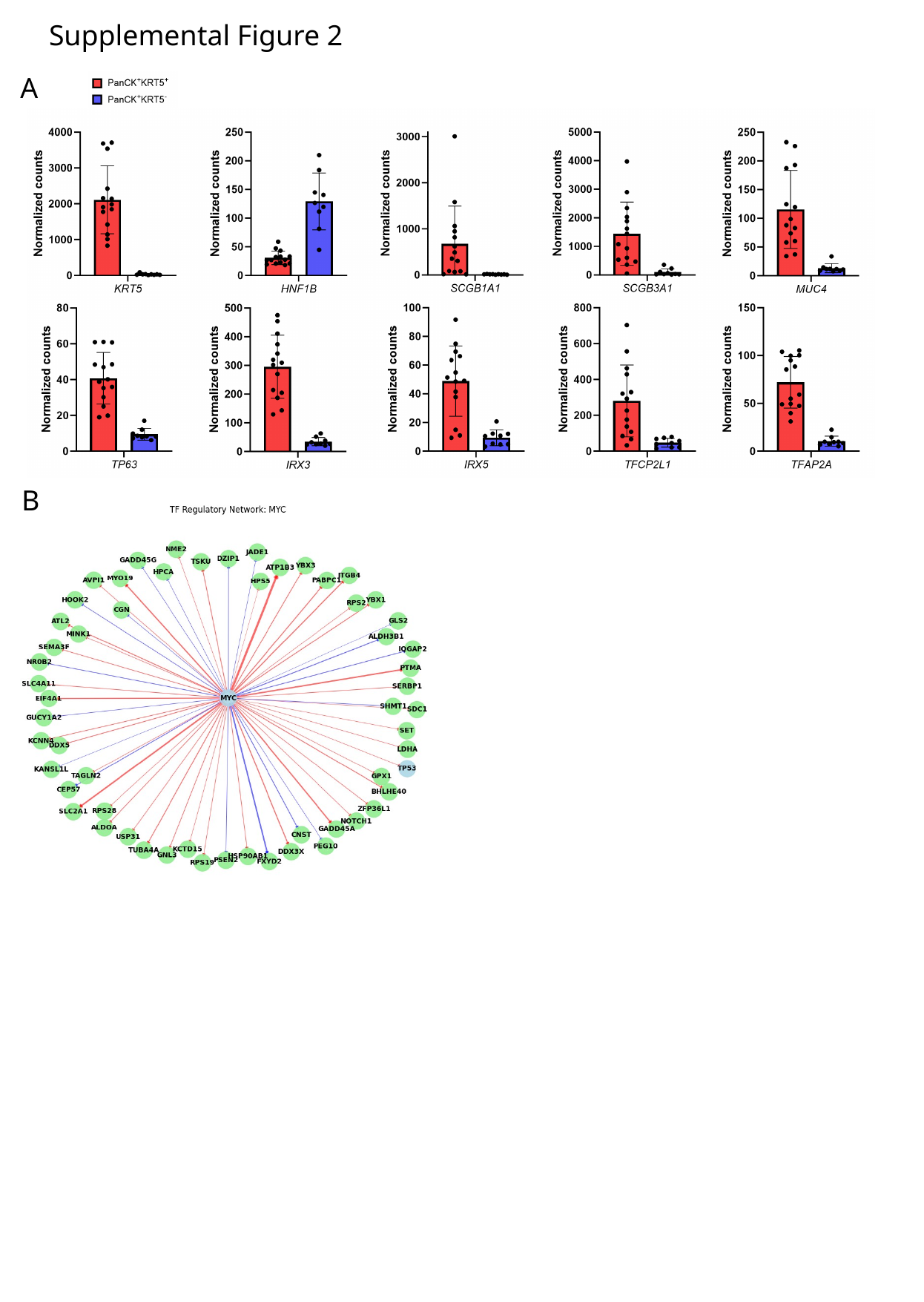

Supplemental Figure 2
A
B

### Slide 5
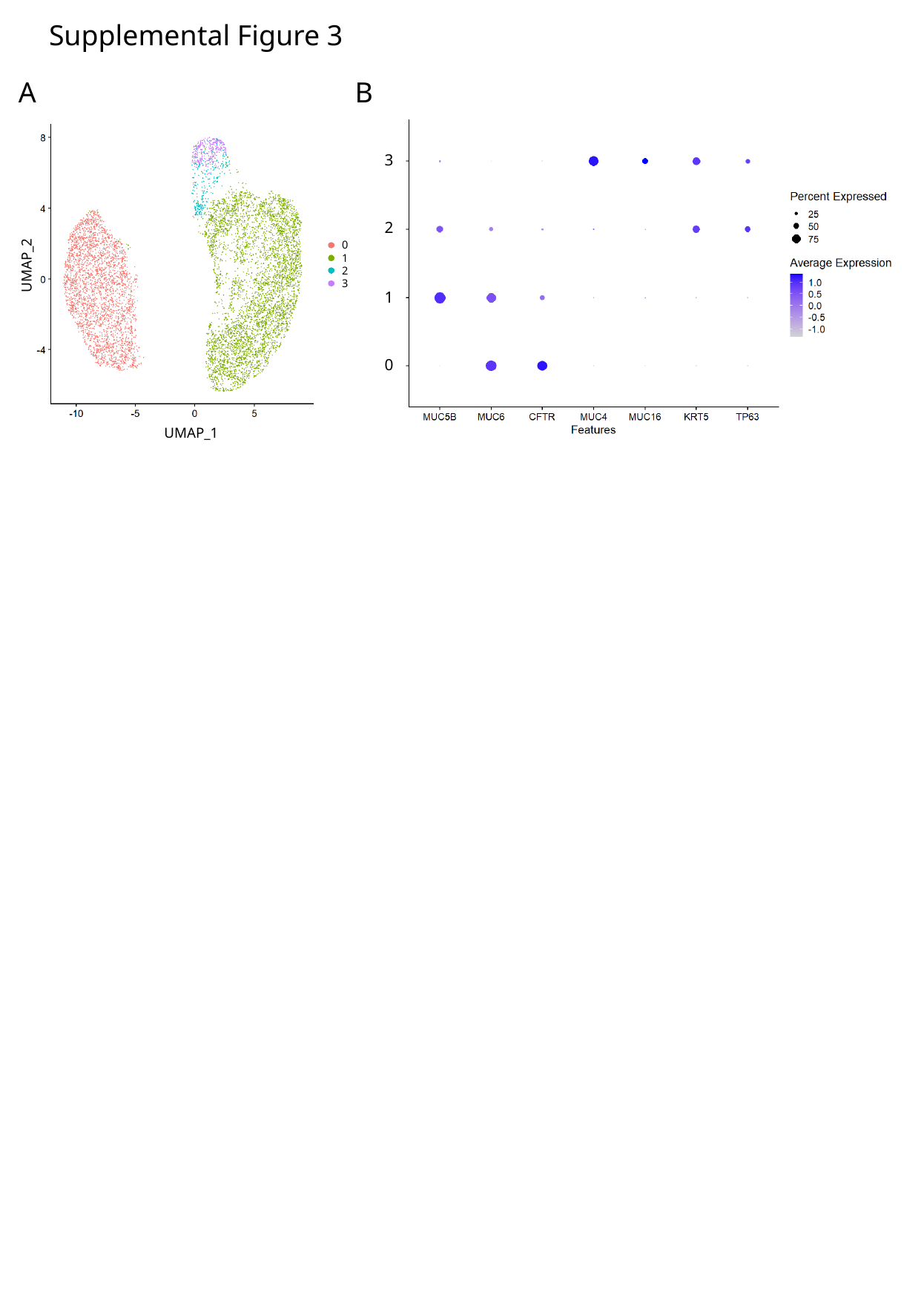

Supplemental Figure 3
A
B
3
2
0
1
2
3
UMAP_2
1
0
UMAP_1

### Slide 6
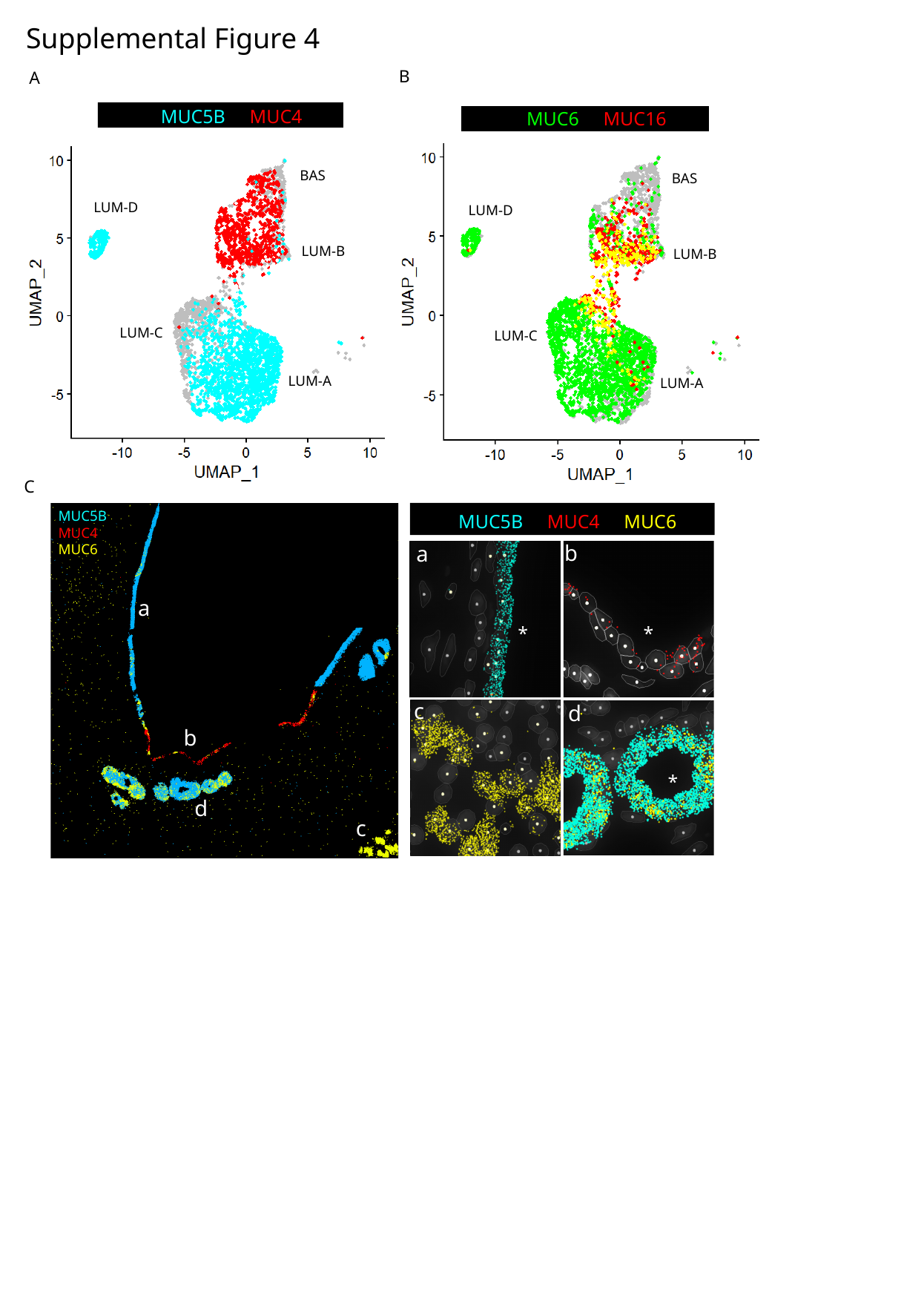

Supplemental Figure 4
B
A
 MUC5B MUC4
 MUC6 MUC16
BAS
LUM-D
LUM-B
LUM-C
LUM-A
BAS
LUM-D
LUM-B
LUM-C
LUM-A
*
*
*
C
MUC5B
MUC4
MUC6
a
b
d
c
 MUC5B MUC4 MUC6
b
a
c
d
*
*
*

### Slide 7
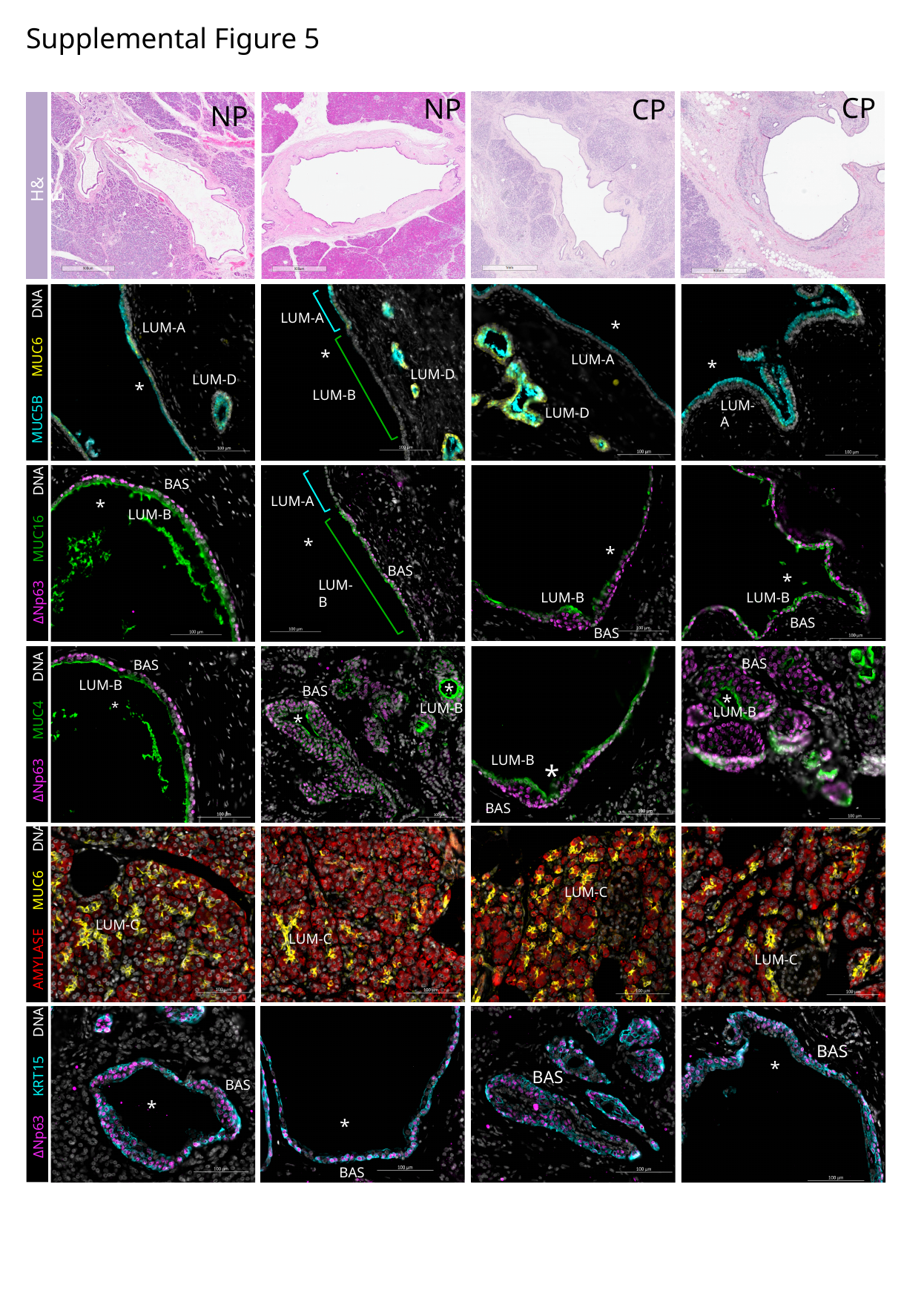

Supplemental Figure 5
CP
NP
CP
NP
H&E
LUM-A
*
MUC5B MUC6 DNA
LUM-A
*
LUM-A
*
LUM-D
LUM-D
*
LUM-B
LUM-A
LUM-D
BAS
LUM-A
*
LUM-B
ΔNp63 MUC16 DNA
*
*
BAS
*
LUM-B
LUM-B
LUM-B
BAS
BAS
*
*
BAS
BAS
LUM-B
*
BAS
*
*
LUM-B
LUM-B
ΔNp63 MUC4 DNA
LUM-B
BAS
LUM-C
AMYLASE MUC6 DNA
LUM-C
LUM-C
LUM-C
BAS
BAS
*
ΔNp63 KRT15 DNA
BAS
*
*
BAS

### Slide 8
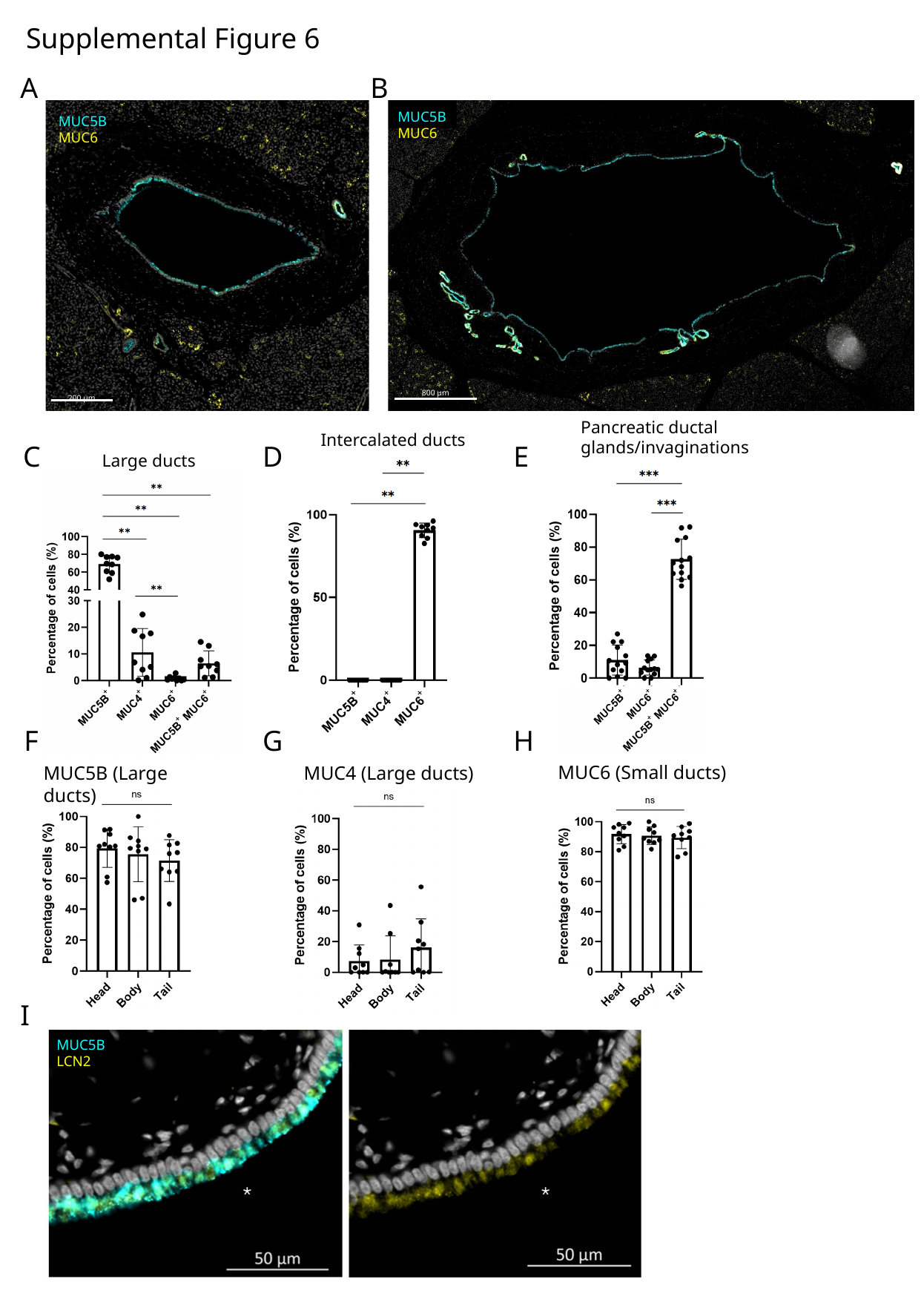

Supplemental Figure 6
A
B
MUC5B
MUC6
200 µm
MUC5B
MUC6
800 µm
Pancreatic ductal glands/invaginations
Intercalated ducts
C
D
E
Large ducts
F
G
H
MUC6 (Small ducts)
MUC5B (Large ducts)
MUC4 (Large ducts)
I
MUC5B
LCN2

### Slide 9
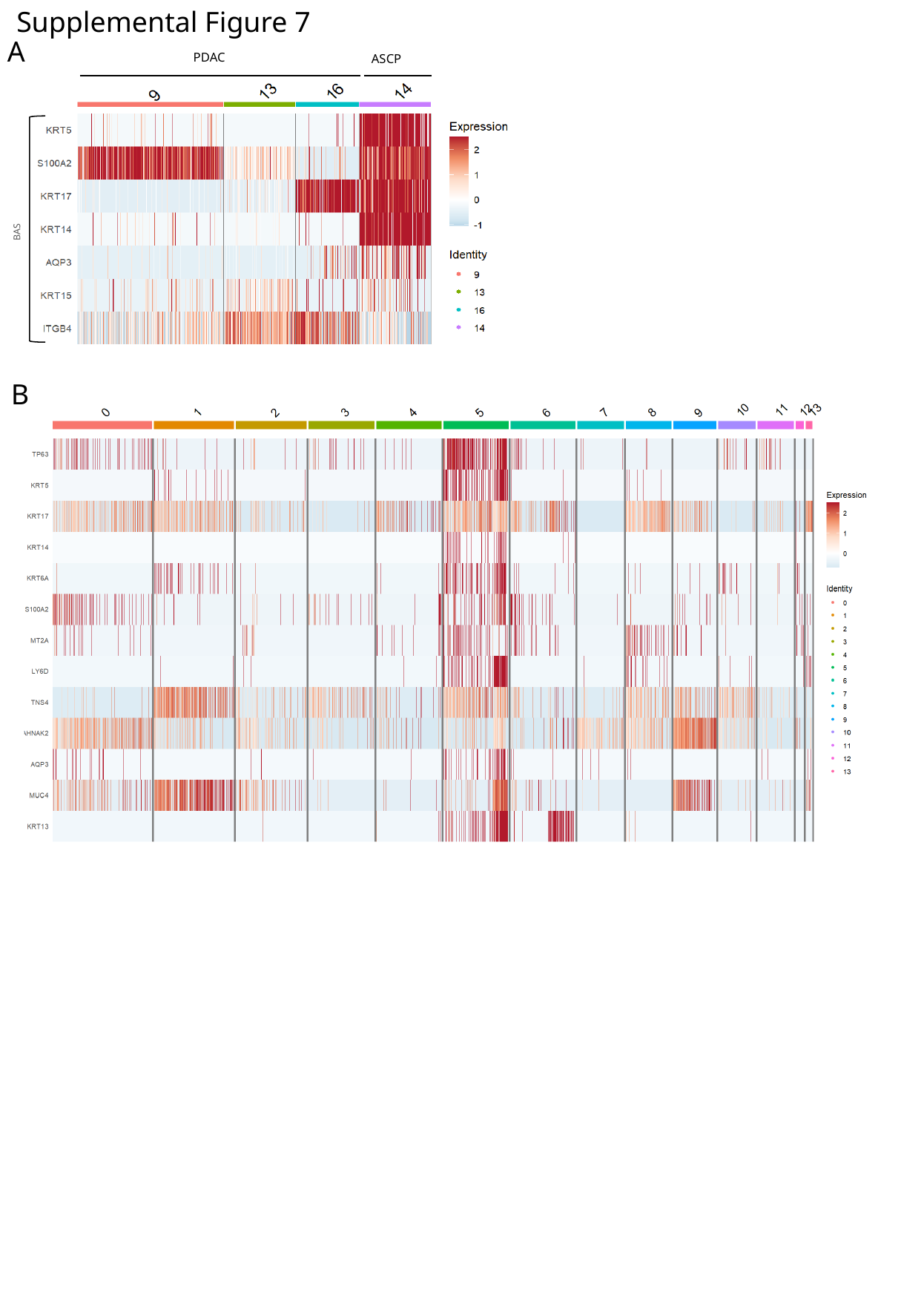

Supplemental Figure 7
A
PDAC
ASCP
BAS
B

### Slide 10
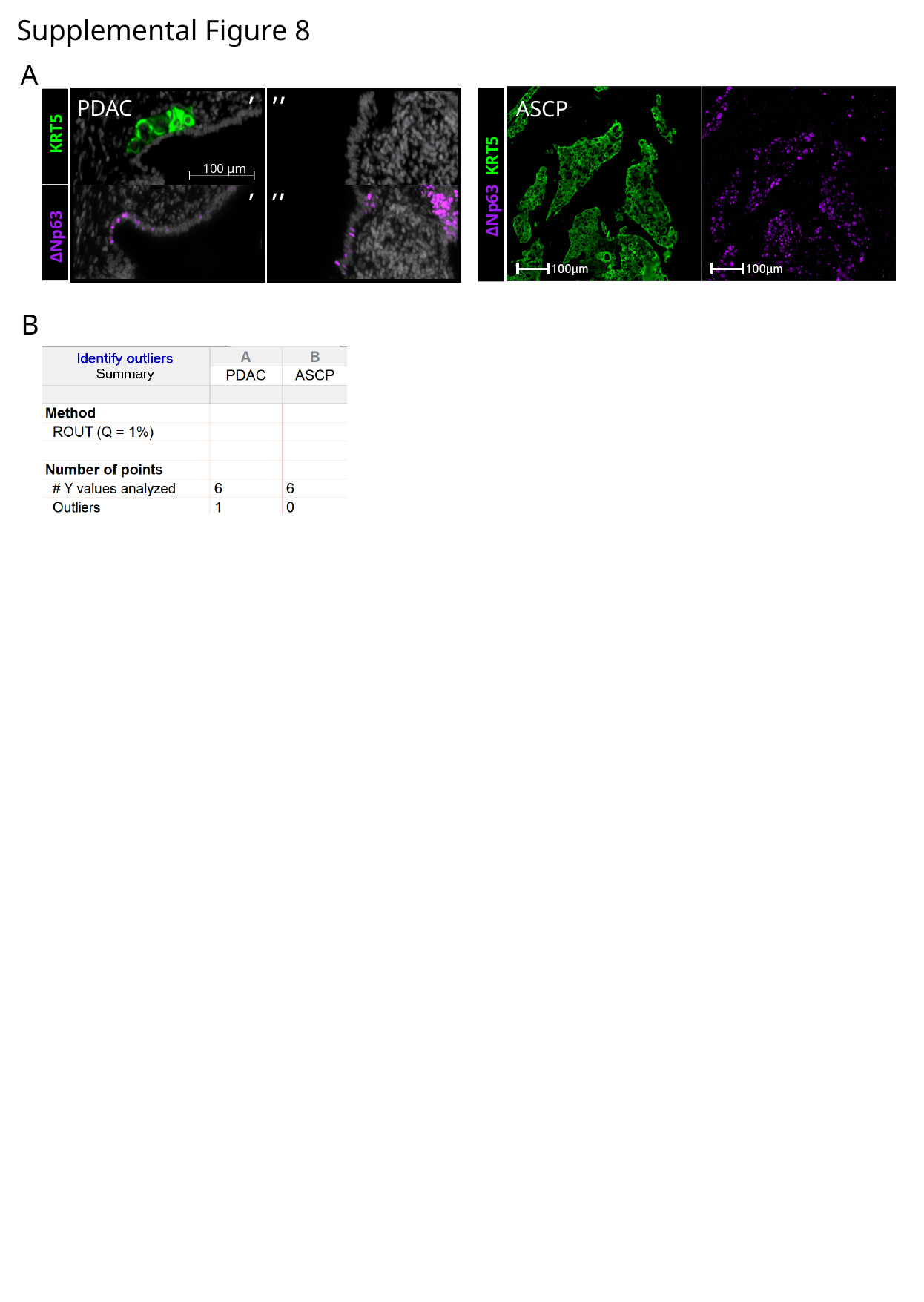

Supplemental Figure 8
A
,
,,
PDAC
  KRT5
100 µm
,
,,
ΔNp63
ASCP
ΔNp63  KRT5
B

### Slide 11
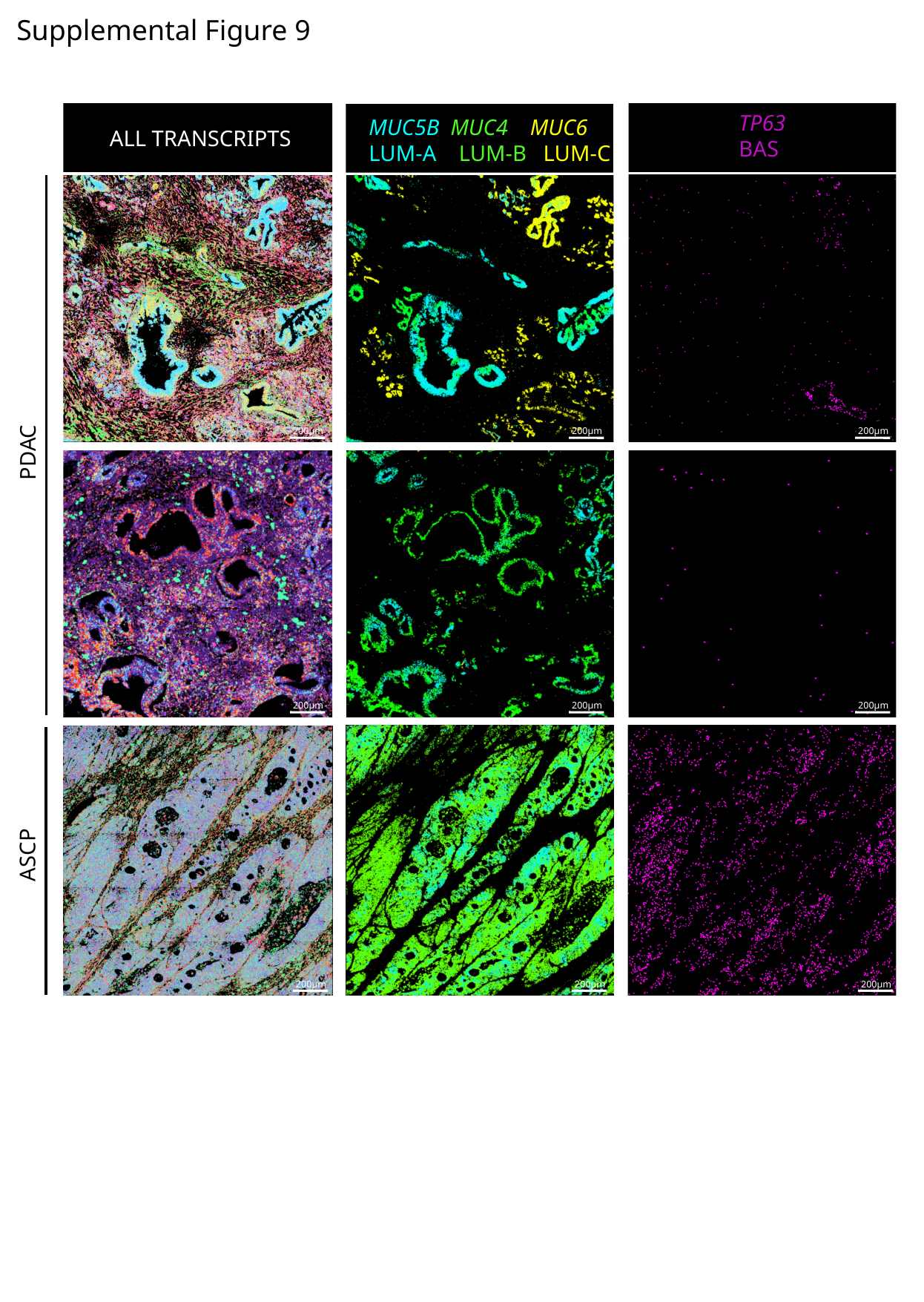

Supplemental Figure 9
TP63
BAS
MUC5B MUC4 MUC6
LUM-A LUM-B LUM-C
ALL TRANSCRIPTS
200µm
200µm
200µm
PDAC
200µm
200µm
200µm
ASCP
200µm
200µm
200µm
200µm
200µm
200µm

### Slide 12
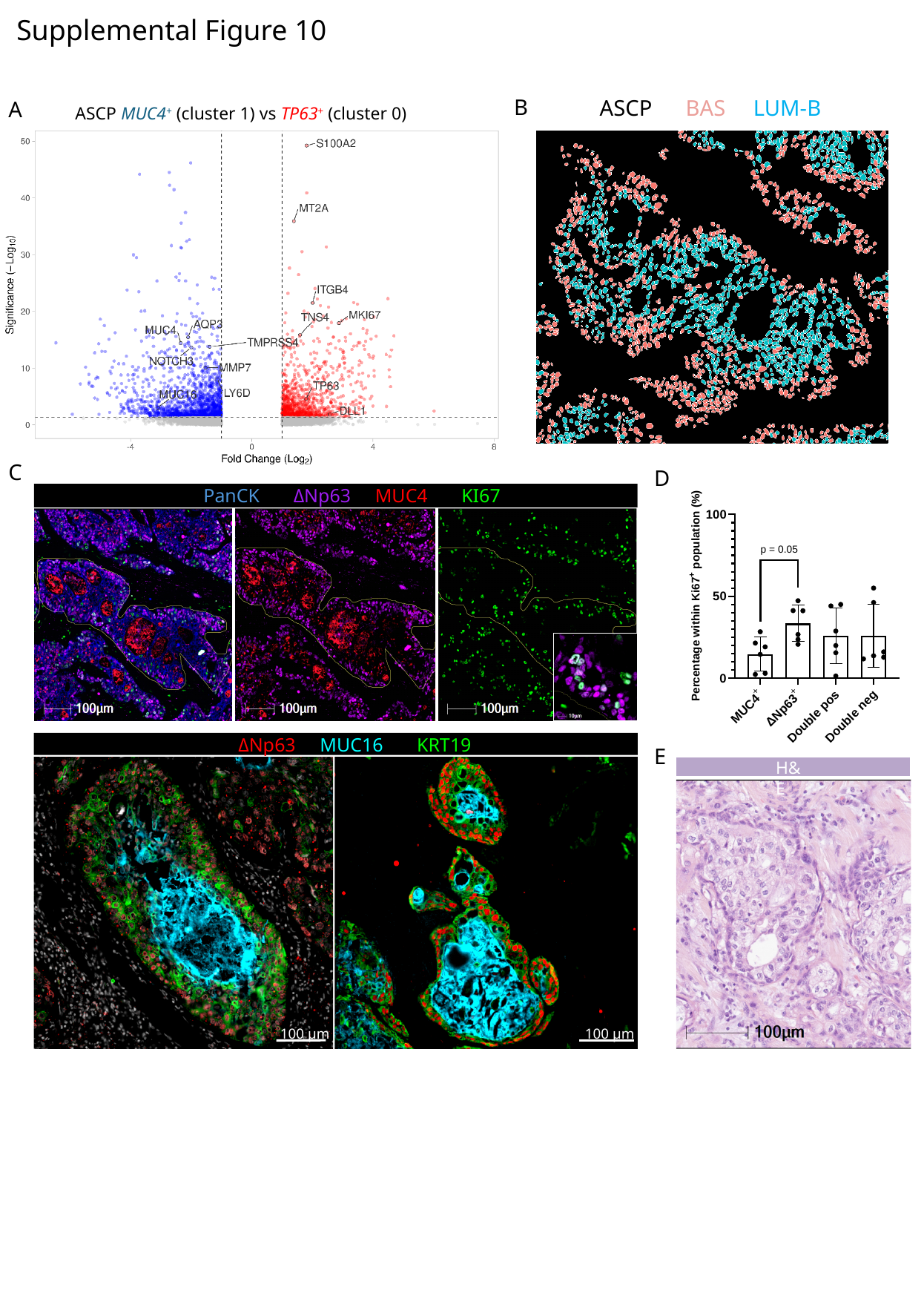

Supplemental Figure 10
B
ASCP BAS LUM-B
A
ASCP MUC4^+^ (cluster 1) vs TP63^+^ (cluster 0)
C
D
PanCK ΔNp63  MUC4  KI67
 ΔNp63  MUC16  KRT19
100 µm
100 µm
E
H&E

### Slide 13
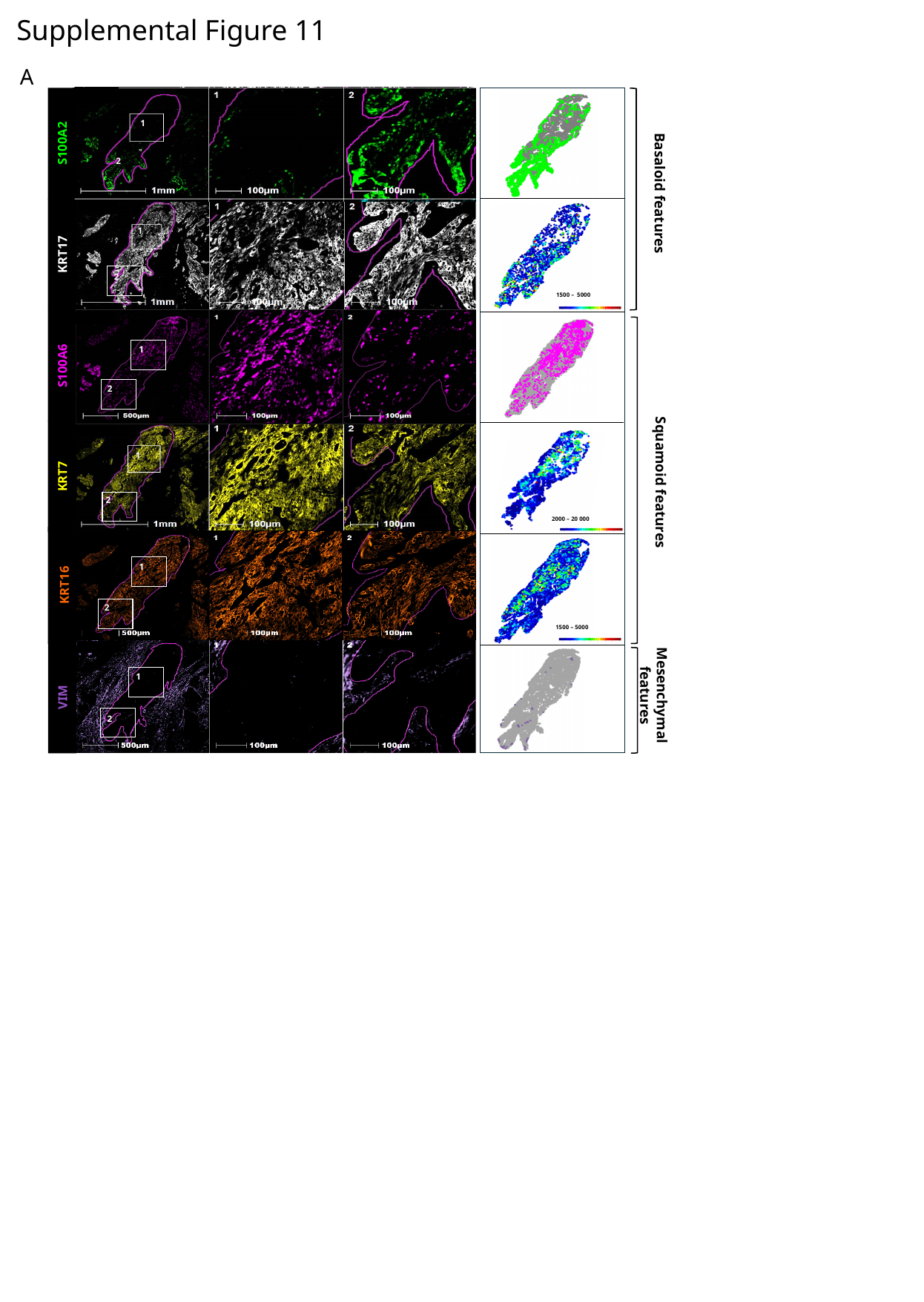

Supplemental Figure 11
A
1
S100A2
2
2
1
KRT17
2
S100A6
1
2
1
2
1
2
1500 – 5000
2000 – 20 000
1
2
Basaloid features
1
1
2
2
Squamoid features
KRT7
1
KRT16
2
1500 – 5000
1
Mesenchymal
features
VIM
2

### Slide 14
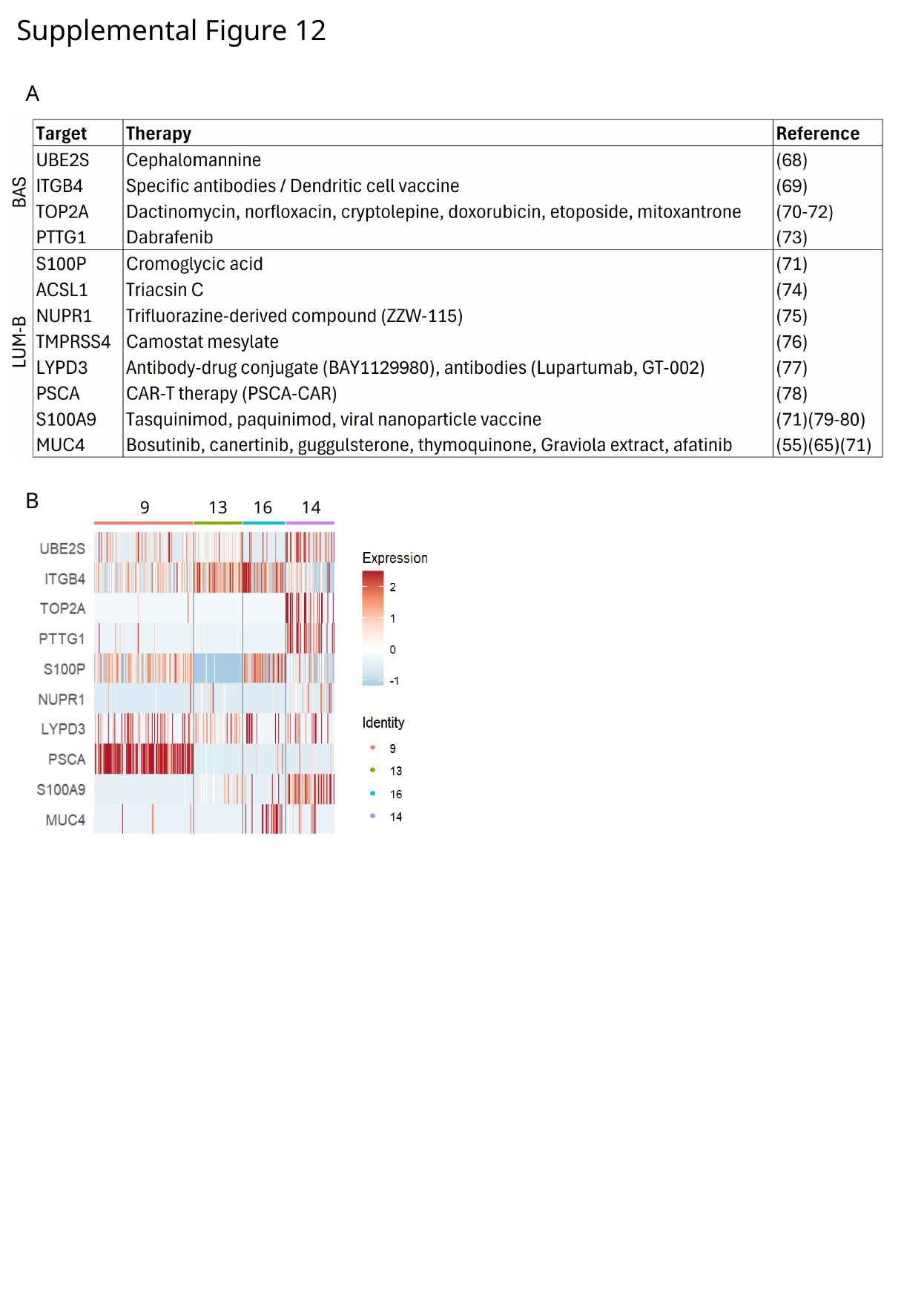

Supplemental Figure 12
A
B
9
13
16
14
