## Supplemental Figure Legends for "A Luminal–Basal Stratification of the Native Human Pancreatic Duct is Differentially Represented in Pancreatic Cancers"

**Supplemental Table 1:**

Overview of custom designed 100-plex gene panel for Resolve Molecular Cartography®.

**Supplemental Table 2:**

Overview of primary antibodies.

**Supplemental Figure 1:**

(A) Stepwise protocol of GeoMx® set-up for FFPE slides of healthy pancreas and chronic pancreatitis. (B) Stepwise protocol of single-cell spatial transcriptomics tools Resolve Molecular Cartography® (Resolve Biosciences) and CosMx Spatial Molecular Imager® (NanoString).

**Supplemental Figure 2:**

(A) Normalized counts of genes of interest from GeoMx® data comparing KRT5^+^ and KRT5^-^ cells (*n=* 23 ROI). (B) Regulatory network of MYC, based on GeoMx® expression data.

**Supplemental Figure 3:**

(A) Ductal cells were subset based on SOX9 expression prior to UMAP visualization of unsupervised clustering, using n=2 non-neoplastic pancreata from Resolve MC® data. (B) Dotplot of luminal/basal classification genes in different clusters of duct cells.

**Supplemental Figure 4:**

(A-B) UMAPs illustrating the expression of (A) MUC5B, MUC4 and (B) MUC6 and MUC16 in different duct populations. Cells highlighted in yellow co-express both markers. (C) Spatial plot of mucin expression (*MUC5B, MUC4, MUC6*) in NP ducts, analysed by Resolve Molecular Cartography®. Letters a-d indicate insets from Figure 2D.

**Supplemental Figure 5:**

First row shows representative Hematoxylin-Eosin stainings of two NP and two CP samples. Following rows show immunofluorescent stainings of the luminal-basal classification markers on samples of non-neoplastic pancreata (*n=*32 NP, *n=*12 CP). Four distinct samples are shown per row. Asterisk labels the lumen.

**Supplemental Figure 6:**

(A-B) MUC5B and MUC6 immunofluorescent staining of interlobular (A) and main pancreatic duct (B) in NP. (C-E) Quantification of mucins in different compartments of the non-neoplastic pancreas. Wilcoxon matched-pairs signed rank test was used to test significance. *p < 0.05, **p < 0.01, ***p < 0.001, *n=*9. (F-H) Quantification of mucins in the head, body and tail of NP. Wilcoxon matched-pairs signed rank test was used to test significance. ns= not significant, *p < 0.05, **p < 0.01, ***p < 0.001, *n=*9. (I) Immunofluorescent staining of MUC5B and LCN2 in LUM-A cells. Asterisk labels the lumen.

**Supplemental Figure 7:**

(A) Heatmap of selected basal-like PDAC clusters and the ASCP cluster for specific BAS markers (*KRT5, S100A2, KRT17, KRT14, AQP3, KRT15, ITGB4*). Wilcoxon rank sum test was used for statistical analysis with min.diff.pct= 0.2. (B) Heatmap of tumor clusters in the snRNA-seq dataset of Hwang *et al.*(8), showing expression levels for top upregulated genes of KRT5^+^ cells as defined by GeoMx in Figure 1. ASCP cells cluster together in cluster 5.

**Supplemental Figure 8:**

(A) Serial immunofluorescent staining for BAS markers ΔNp63 and KRT5 in PDAC. (B) Outlier analysis performed in GraphPad Prism 10® using the Robust regression Outlier removal (ROUT) method.

**Supplemental Figure 9:**

Spatial plots of transcripts in the Recognize® platform of Resolve MC® in PDAC and ASCP samples. First column shows all 100 transcripts, second column shows the mucins from the luminal-basal classification and final column shows BAS marker *TP63*.

**Supplemental Figure 10:**

(A) Volcano plot of differential gene expression between cluster 0 (BAS) and cluster 1 (LUM-B) cells in the ASCP scRNA-seq data of Zhao *et al.*(51). Genes were considered differentially expressed when Log_2_Fold Change > 1.2 and p-value < 0.05. Genes shown in the plot overlap with the KRT5^+^ GeoMx® profile. (B) Spatial plot of BAS and LUM-B clusters in ASCP analysed by Resolve MC®. (C) Immunofluorescent staining for epithelial marker PanCK, BAS marker ΔNp63, LUM-B marker MUC4 and proliferation marker Ki67 (top row) and BAS marker ΔNp63, LUM-B marker MUC4 and epithelial marker KRT19 (bottom row) on ASCP tumor nests. (D) Quantification of ΔNp63^+^ and MUC4^+^ cells within the KI67^+^ population in ASCP. Double pos = double positive cells for ΔNp63 and MUC4, Double neg = double negative for both markers. Statistical significance was determined using a paired t-test. Statistical significance was accepted at p < 0.05. Error bars show standard deviation. (E) H&E staining of an adenosquamous tumor nest.

**Supplemental Figure 11:**

(A) Immunofluorescent stainings on ASCP nests for the basal-like signatures as defined by Hwang *et al.*(8): Basaloid markers (S100A2, KRT17), Squamoid markers (S100A6, KRT7, KRT16) and a Mesenchymal marker (VIM). The rightmost panel shows the spatial plot obtained using HALO® image analysis software, demonstrating the positively detected signal for every marker.

**Supplemental Figure 12:**

(A) List of conserved targets overlapping with the GeoMx® KRT5^+^ profile and the scRNAseq profiles of BAS and LUM-B populations in ASCP (B) Heatmap demonstrating the expression of conserved BAS (*UBE2S, ITGB4, TOP2A, PTTG1*) and LUM-B (*S100P, NUPR1, LYPD3, PSCA, S100A9, MUC4*) druggable targets in basal-like PDAC clusters (clusters 9, 13, 16) and ASCP cluster (cluster 14) in the merged object of Figure 5A.
