## Supplemental Methods for "A Luminal–Basal Stratification of the Native Human Pancreatic Duct is Differentially Represented in Pancreatic Cancers"

**Patient involvement**

The findings of this study were shared with pancreatic cancer patients in an early stage to encourage active dialogue and gather feedback. Additionally, the final data was presented at a patient event for pancreatic cancer patients, where they had the opportunity to interact with medical staff and researchers.

**Immunofluorescence**

Human pancreatic samples were fixed in 4% paraformaldehyde and embedded in paraffin. FFPE slides of 5 micron were deparaffinized in toluene and subsequently rehydrated in a serially diluted propanol sequence. 3% Hydrogen peroxidase was utilized to block endogenous peroxidase activity for 30 minutes at room temperature. Afterwards, slides were placed in a pressure cooker for 40 minutes in citrate buffer (Sigma-Aldrich, St-Louis, MO, USA, C9999) for antigen retrieval. Slides were then incubated with protein block for 30 minutes using 25% diluted casein (Thermo Fisher Scientific, Waltham, MA, USA, 37528). The primary antibodies that were used are listed in Supplementary Table 2. Primary antibodies were incubated overnight at 4°C. The secondary antibodies were added together with Hoechst dye (Thermo Fisher Scientific, H1399, diluted 1/250) for nucleus detection. Slides were mounted using ProLong Gold Antifade mounting medium (Thermo Fisher Scientific, P36930).

Basescope detection of KRASG12D was performed as described in Michiels *et al.*(17).

**Lunaphore Comet™**

Formalin-fixed paraffin-embedded (FFPE) slides were preprocessed using the PT Module (Epredia) with Dewax and HIER Buffer H (TA999-DHBH, Epredia) for 60 min at 99 °C. After rinsing, slides were stored in Multistaining Buffer (BU06, Lunaphore) until use. A 13-plex protocol template was generated using COMET Control Software, and reagents were loaded onto the instrument to run the fully automated sequential immunofluorescence (seqIF) protocol. Nuclear staining was performed with DAPI (Thermo Scientific, cat. no. 62248, 1/1000 dilution) either by dynamic incubation for 2 min or by supplementing secondary antibody cocktails with DAPI. For all staining cycles, the dynamic incubation time for primary antibody mixes was set to 8 min, while the dynamic incubation time for secondary antibody and DAPI cocktails was set to 2 min. All primary antibody cocktails were prepared in Multistaining Buffer (BU06, Lunaphore). For each imaging cycle, exposure times were 25 ms for DAPI, 250 ms for TRITC, and 400 ms for Cy5. The elution step was performed for 2 min in each cycle using Elution Buffer (BU07-L, Lunaphore), followed by a 30 s quenching step with Quenching Buffer (BU08-L, Lunaphore). Imaging was carried out in Imaging Buffer (BU09, Lunaphore). Secondary antibody mixes consisted of Alexa Fluor Plus 647 donkey anti-mouse (Invitrogen, cat. no. A32787, 1/200 dilution) and Alexa Fluor Plus 555 donkey anti-rabbit (Invitrogen, cat. no. A32794, 1/200 dilution), or Alexa Fluor Plus 647 donkey anti-rabbit (Invitrogen, cat. no. A48270, 1/200 dilution) and Alexa Fluor Plus 555 donkey anti-rat (Invitrogen, cat. no. A32727, 1/100 dilution), or Alexa Fluor Plus 647 donkey anti-mouse (Invitrogen, cat. no. A32787, 1/200 dilution) and Alexa Fluor Plus 555 donkey anti-goat (Invitrogen, cat. no. A32816, 1/200 dilution). The imaging area was 12.5mm ×12.5 mm. Upon completion of the experiment, a raw OME-TIFF file was generated by the COMET Control Software for downstream analysis using HALO Indica software.

**Primary cell organoid culture**

Healthy human exocrine cells were washed twice in 1X DPBS (14190250, Thermo Fisher), after which 1 µL of the cell pellet was resuspended in 40 µL of freshly thawed Matrigel (Corning, New York, USA). Matrigel domes were created in a Nunclon Sphera 24-well plate (174930, Thermo Fisher) and were incubated at 37 °C for 30 minutes. Afterwards, 500 µL of complete medium was added to each well (Advanced DMEM/F-12 (12643010, Thermo Fisher, Waltham, USA) supplemented with 1/100 GlutaMAX (35050061, Thermo Fisher), 100 U/mL Penicillin-Streptomycin (15140122, Thermo Fisher), 0.01 M HEPES (11560496, Thermo Fisher), 1/50 B-27 supplement (12587010, Thermo Fisher), 10 mM Nicotinamide (N0636, Sigma, Kawasaki, Japan), 10.5 mM Y-27632 (Y0503, Sigma), 10 ng/mL Wnt3a (5036, R&D Systems, Minneapolis, USA), 100 ng/mL Noggin (120-10C, Peprotech, Cranbury, USA), 1 µg/mL R-spondin (12038, Peprotech), 1.25 mM NAC (A0737, Sigma), 500 nM A83-01 (2939, Tocris Bioscience, Bristol, UK), 10 nM Gastrin1 (G9145, Sigma), 100 ng/mL hFGF (100-26, Peprotech), 50 ng/mL hEGF (AF-100-15, Peprotech).

**Lentiviral transduction of organoids**

Human exocrine organoids were digested into a single cell suspension by incubating for 5 minutes in Accumax® (Thermo Fisher), while resuspending. The cells were then incubated overnight in a Matrigel-coated 24-well plate together with the lentivirus (Multiplicity of Infection = 20) in organoid culture medium supplemented with TransDux MAX® (System Biosciences, Palo Alto, USA) to enhance transduction efficiency. The next day, medium was aspirated and cells were washed three times with 1X DPBS, after which another Matrigel layer was placed on top of the cells. After 30 minutes of solidifying, complete medium was placed in the well for further culture. Lentivirus pLV[EXP]-CMV>(deltaNp63alpha) was obtained using VectorBuilder (Neu-Isenburg, Germany).

**siRNA-mediated knockdown of HPDE, BxPC3, T3M4 and PDAC-021**

Knockdown was performed as described in Martens *et al.*(18).

**Single-cell RNA sequencing re-analysis**

Single-cell transcriptomic data from a pancreatic adenosquamous tumor was obtained from the publicly accessible dataset GSE165399(51). Preprocessing of this data was conducted using RStudio (v2023.3.0.386) and the “Seurat” package (version 5.1.0)(48). To minimize noise, genes detected in fewer than three cells and cells with fewer than 200 detected genes were excluded. The 2000 most highly variable genes were identified using variance-stabilizing transformation (VST), after which the standard Seurat workflow was applied. A single-cell neighborhood graph was generated via principal component analysis (PCA), employing the first 17 principal components. Clustering was conducted using the Louvain algorithm with a resolution parameter of 0.1, and cell cluster identities were assigned based on specific marker genes, enabling the delineation of distinct cell populations. To extend our analysis, a publicly available, pre-processed dataset containing malignant pancreatic ductal adenocarcinoma (PDAC) cell subsets was incorporated. This dataset, formatted as a Seurat object and comprising multiple PDAC samples(50), is accessible via Zenodo (<https://zenodo.org/record/6024273#.Yg2eTJZUtaY>). The processed PDAC Seurat object was merged with a subset of tumor cells (selected based on epithelial markers such as *KRT19*) from the previously mentioned adenosquamous tumor dataset, and further analysis was conducted using the standard Seurat workflow. A single-cell neighborhood graph was constructed through principal component analysis (PCA) using the first 12 principal components, with clustering achieved via the Louvain algorithm at a resolution of 0.3. Additionally, the impact of mitochondrial gene expression was assessed, and this factor was regressed out during the data scaling process if deemed necessary.

**Transcriptome gene set analysis**

R v4.2.3 and RStudio v2023.3.0.386 environments were used for transcriptomic data analysis. GSVA v1.46.00 package was applied to the Puleo *et al.* cohort and the samples were stratified by the median according to the enrichment of the gene sets of study. The package ggpubr v0.6.0 was used to calculate the Pearson correlation to assess the co-expression of the genes.

Next, transcriptomic subtype classifications of Puleo *et al.* and Chan-Seng-Yue *et al.* were applied to the cohort. To measure the association between the GSVA obtained groups and molecular subtypes, odds ratios (OR) and their corresponding 95% confidence intervals (95% CI) were calculated using univariate logistic regression models implemented with the gtsummary v1.7.2 and foresploter v1.1.2.

**Resolve Molecular Cartography® (continued)**

After hybridization, the probes are colorized and decolorized during a cyclic imaging approach. As a result, every individual target is conjugated to a unique 16-bit fluorescent barcode. Areas of interest were selected for all samples based on protein stainings for ΔNp63, totaling 26 mm^2^ analyzed across all samples. Regions of interest were imaged on a Zeiss Celldiscoverer 7, resulting in 32 Z-stacks per region. The Recognize platform used the Cellpose algorithm, employing the pre-trained *cyto* model. Key parameters were configured as follows: the *diameter* parameter was set to 50.0 and the *flow-thresh* parameter was set to 0.5. Following segmentation, the cell outlines were computed by applying the convex hull algorithm. This was achieved using the *chull* function in R, based on the spatial distribution of transcripts assigned to individual cells by Cellpose. The output files were incorporated into a Seurat object utilizing a customized pipeline developed by adapting publicly available code from GitHub (<https://github.com/ostunilab/PDAC_Nature_2023/blob/main/Molecular_Cartography/Nature2023_Human_PDAC_MolecularCartography_analyses.r>). Genes expressed in fewer than 10 cells and cells expressing fewer than 4 genes were excluded. Seurat object was normalized and scaled using the standard Seurat pipeline. Principal component analysis (PCA) was conducted to reduce dimensionality, and clustering was performed using the Louvain algorithm with 7 principal components and a resolution of 0.2. Marker genes for each identified cluster were subsequently computed using the FindAllMarkers function in Seurat.

**Statistical analysis**

For differential gene expression analysis of GeoMx® data, a linear mixed model was used with Benjamini-Hochberg correction. For differential gene expression analysis of all other data, Wilcoxon rank sum test was used. Survival analyses were conducted using the survival v3.5-3 and survminer v0.4.9 packages. The log-rank test was used to compare Kaplan-Meier curves and p-values < 0.05 were considered statistically significant. Multivariate Cox proportional hazard regression models, stratified by high and low groups based on GSVA results, were used to analyze survival outcomes, with results presented as hazard ratios (HRs) and 95% confidence interval (CI). Disease free survival (DFS) was defined as the time from diagnosis to the first documentation of recurrent disease following surgery. GraphPad Prism® 10 was used to analyze data.  Results are shown as means ± standard deviation.
