## Supplemental File 3 for "A Luminal–Basal Stratification of the Native Human Pancreatic Duct is Differentially Represented in Pancreatic Cancers"

### Non-neoplastic pancreas

(Normal pancreas – NP)

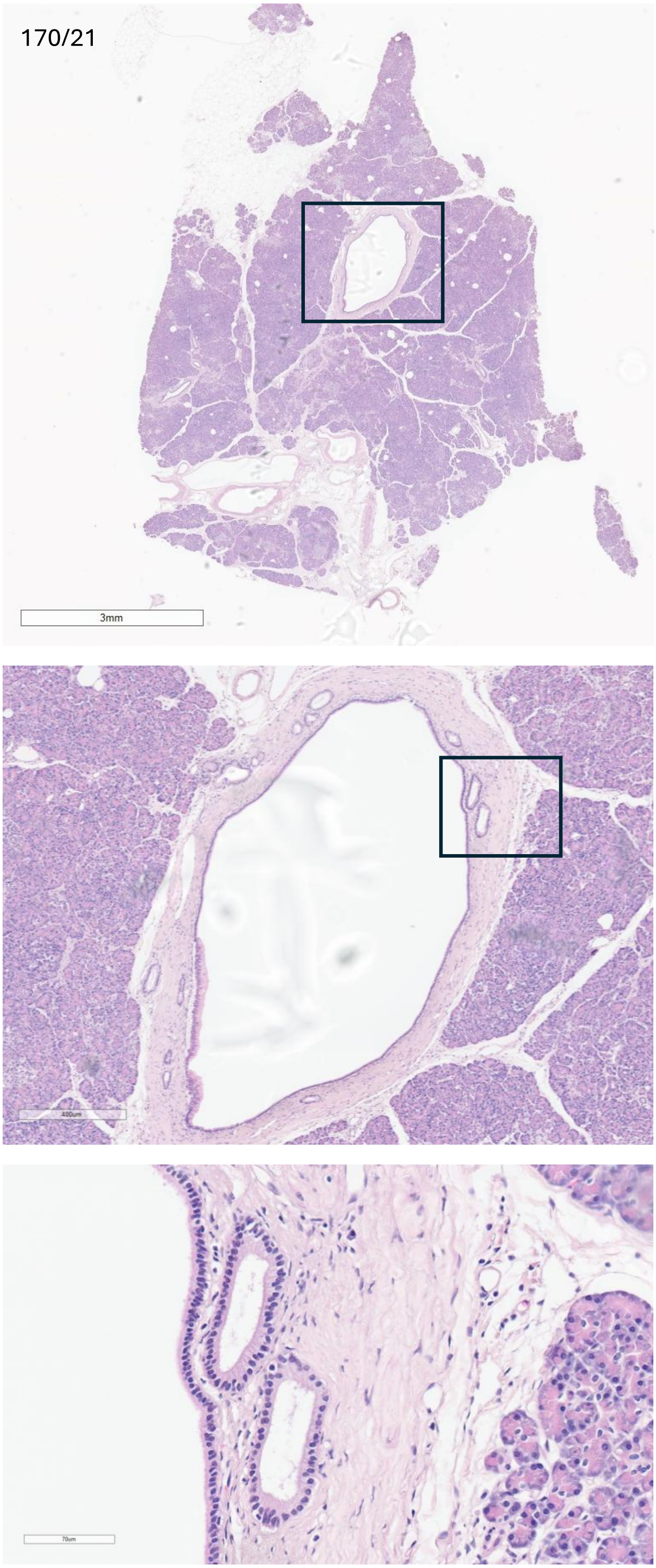
170/21
3mm
400um
70um

171/2

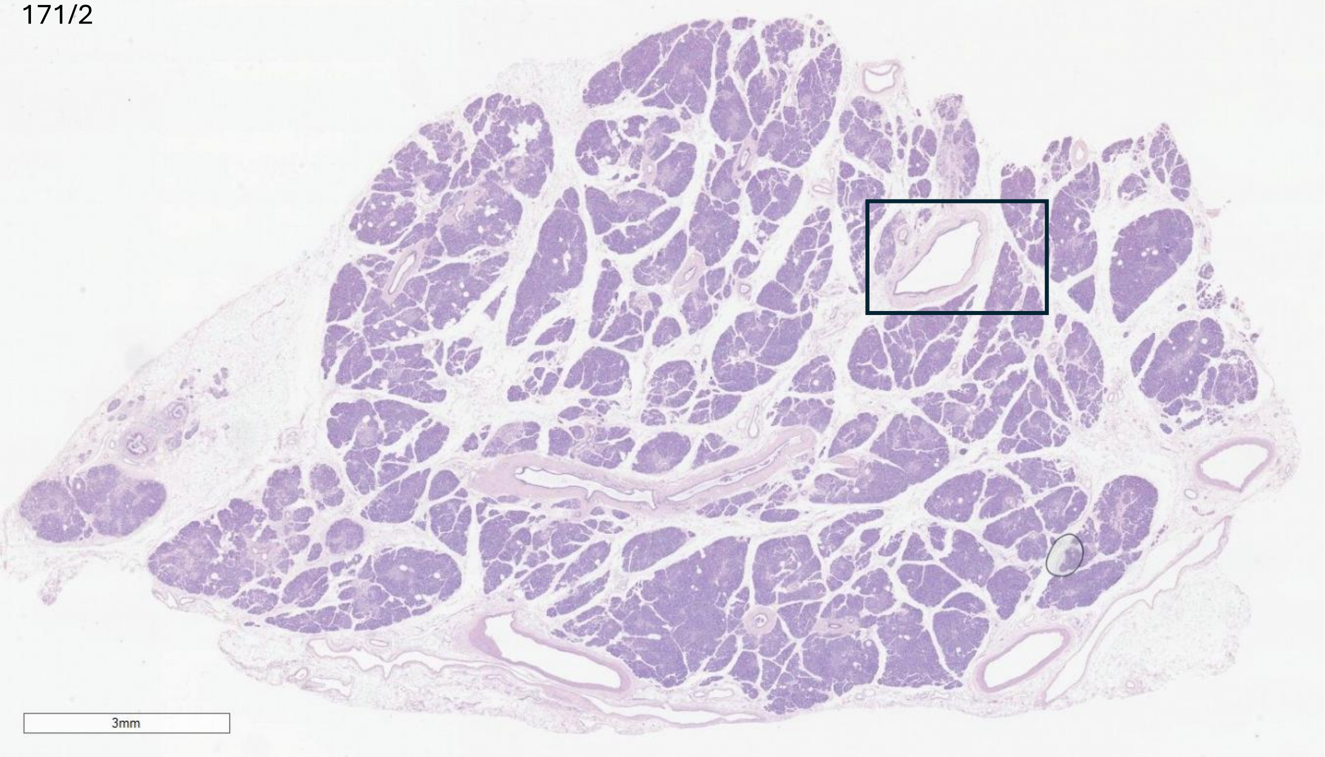


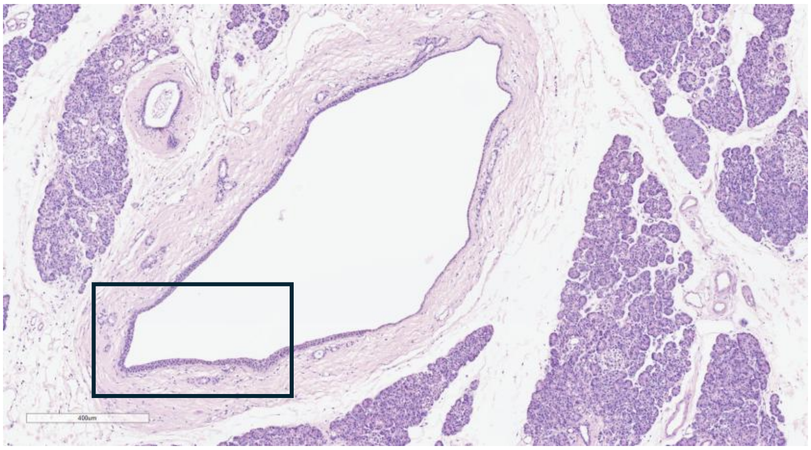


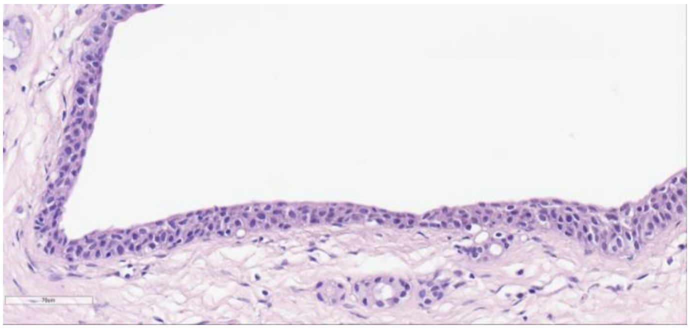

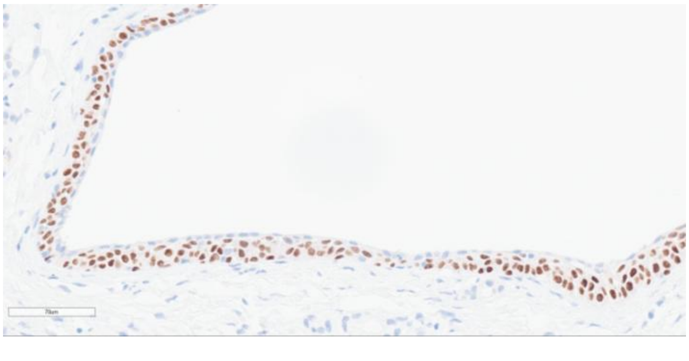

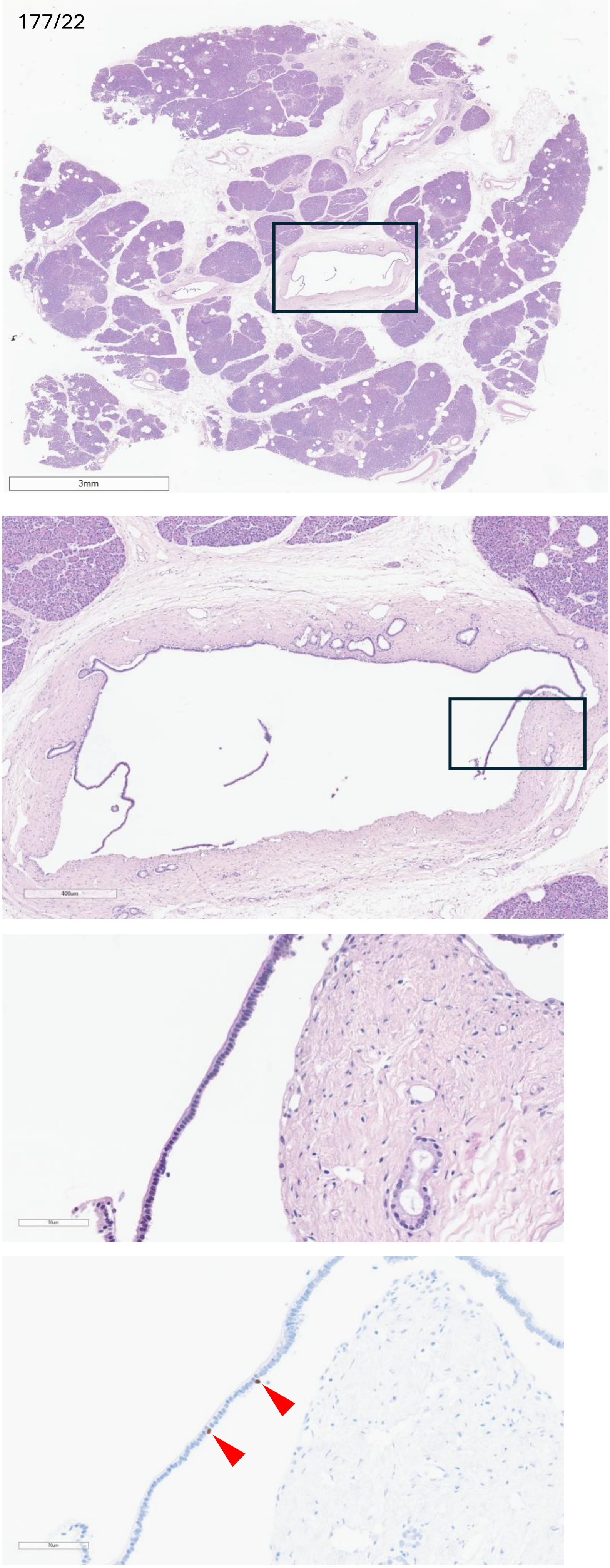
177/22
3mm
400um

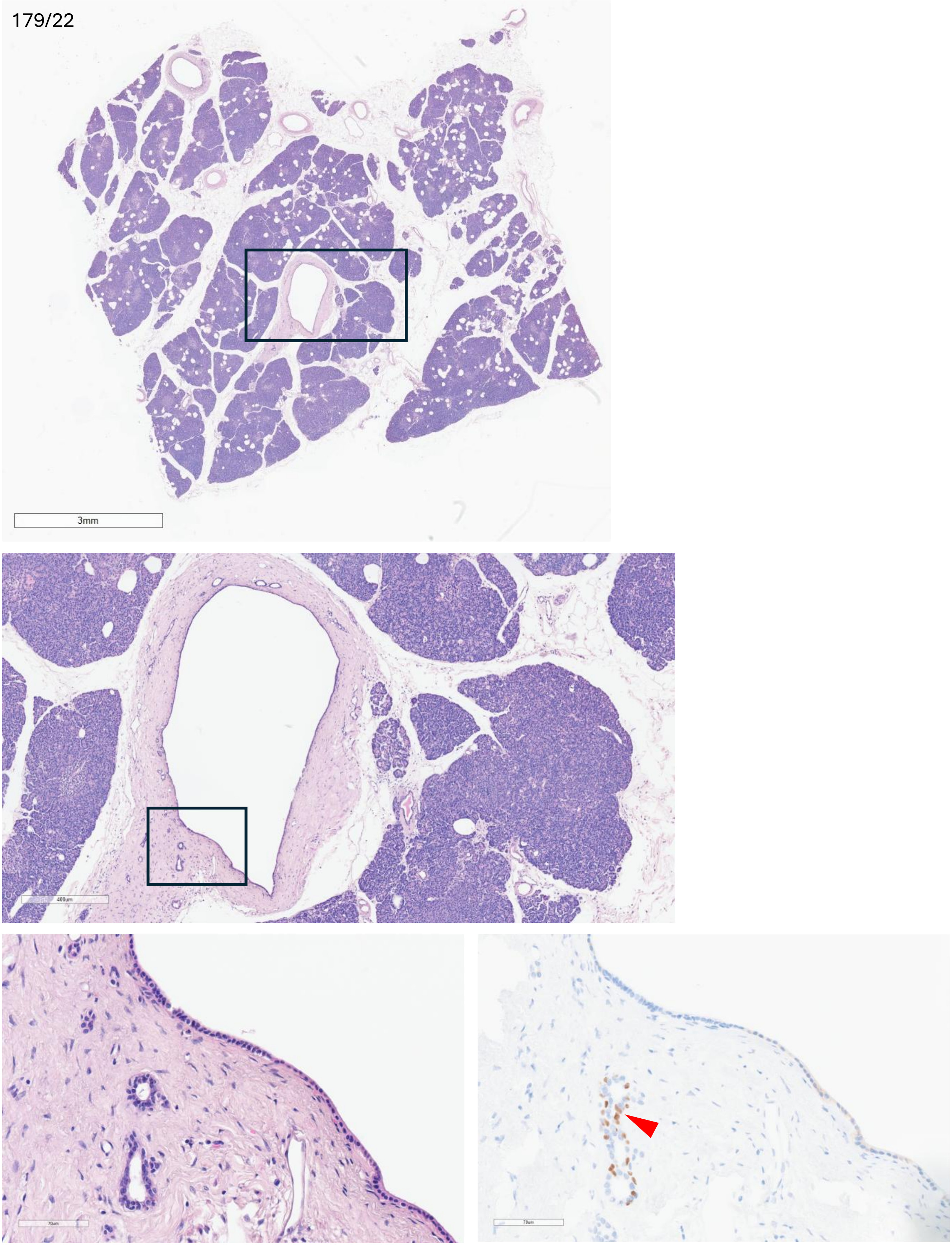
179/22
3mm
400um
70um
70um

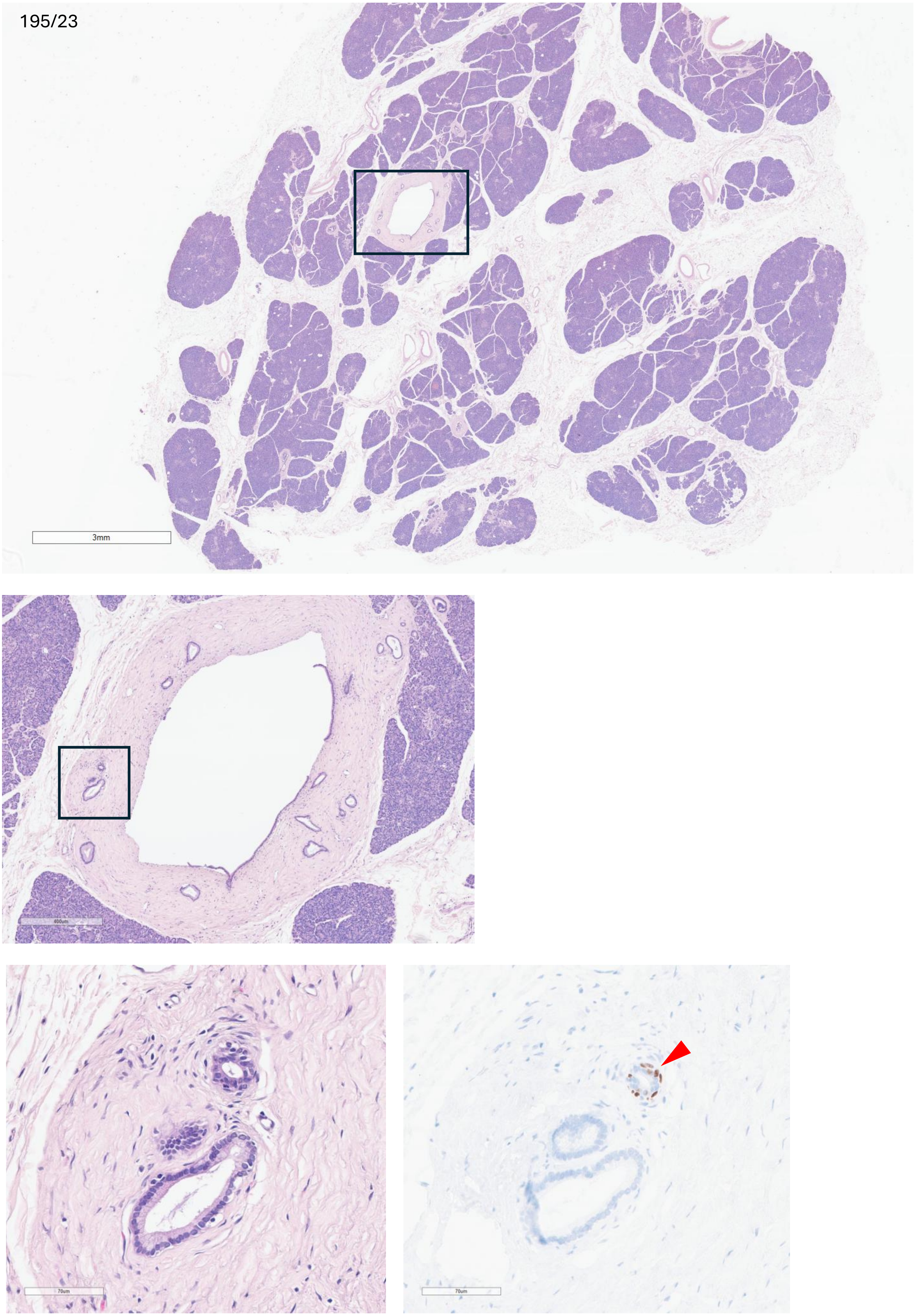
195/23
3mm
400um
70um
70um

198/23

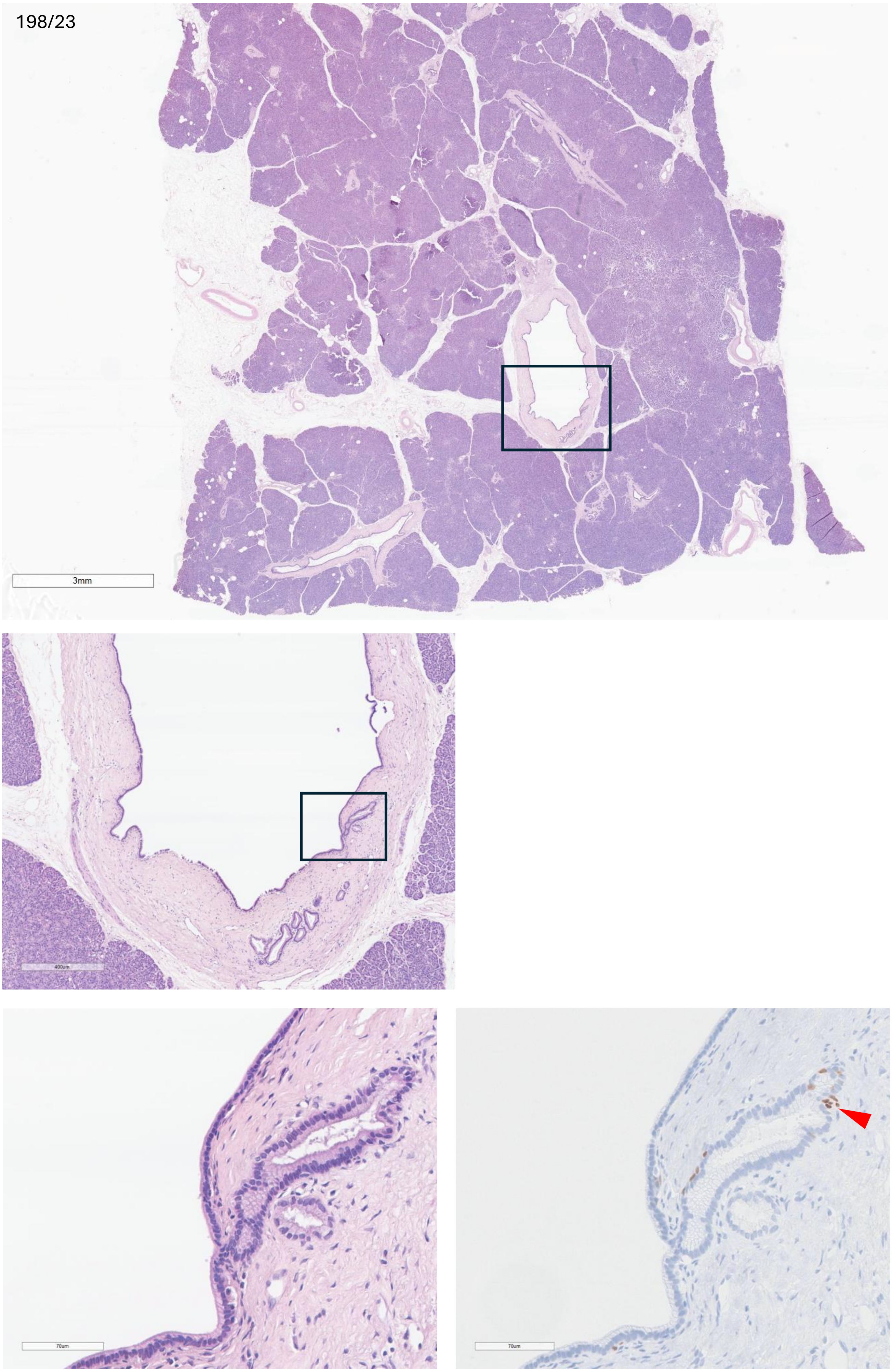

A00-169

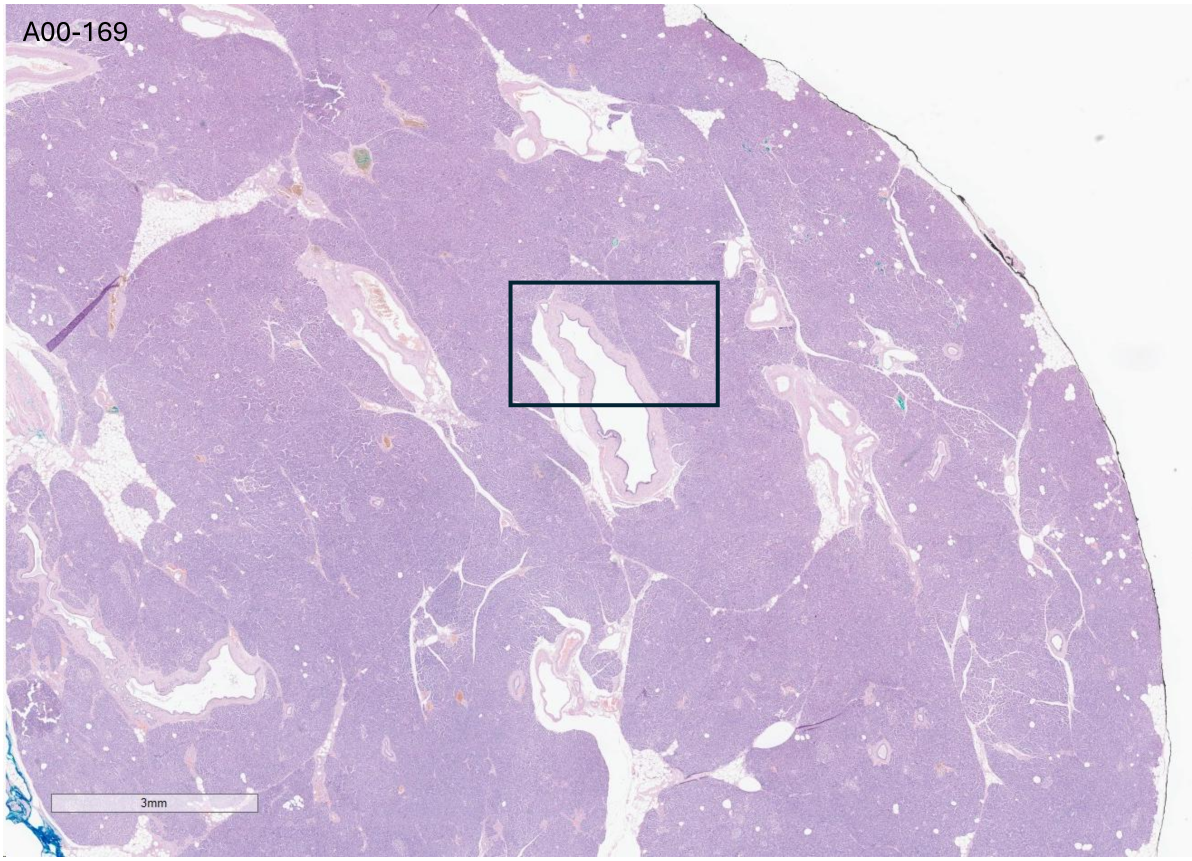


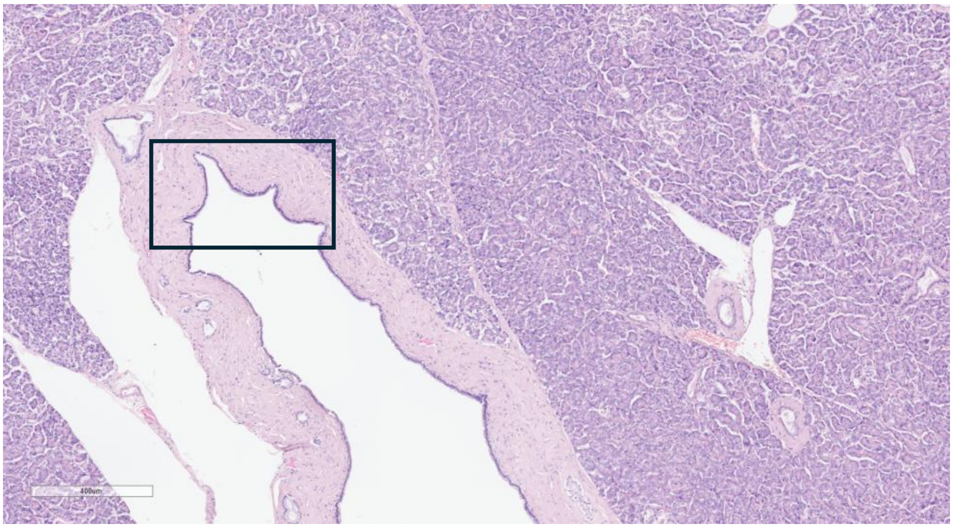


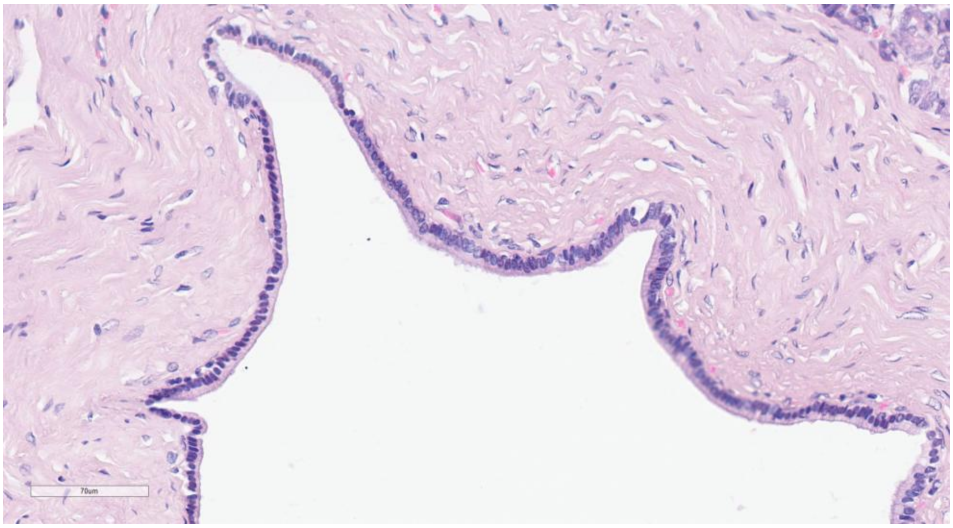

A00-188

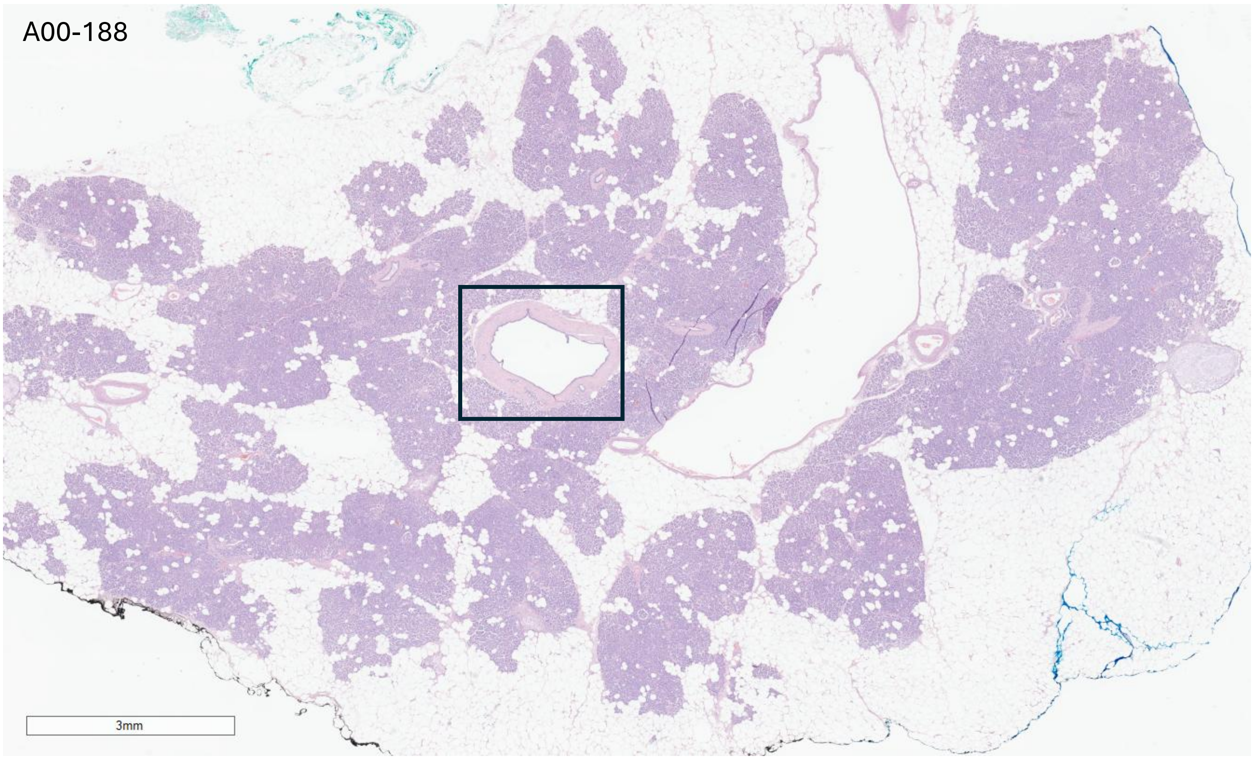


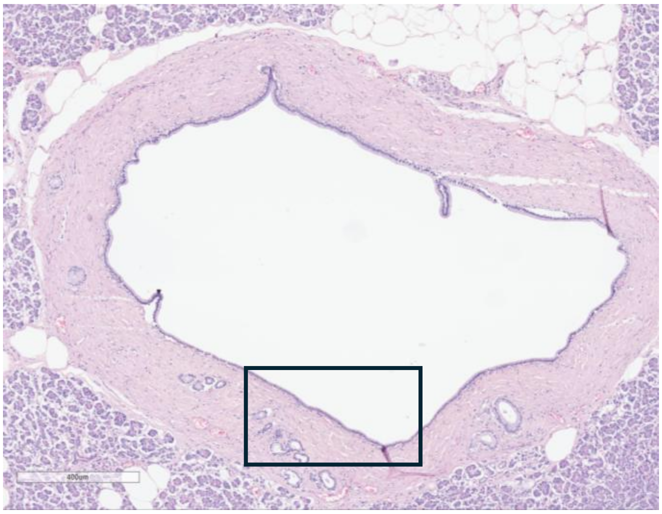


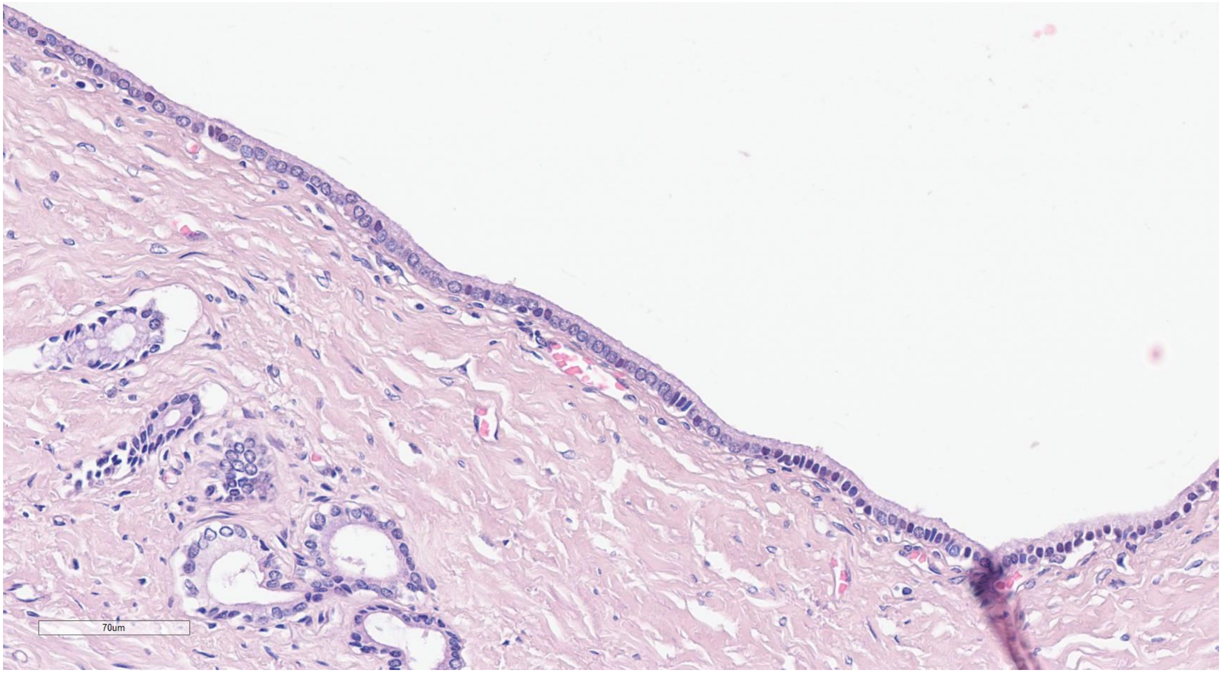

A00-407

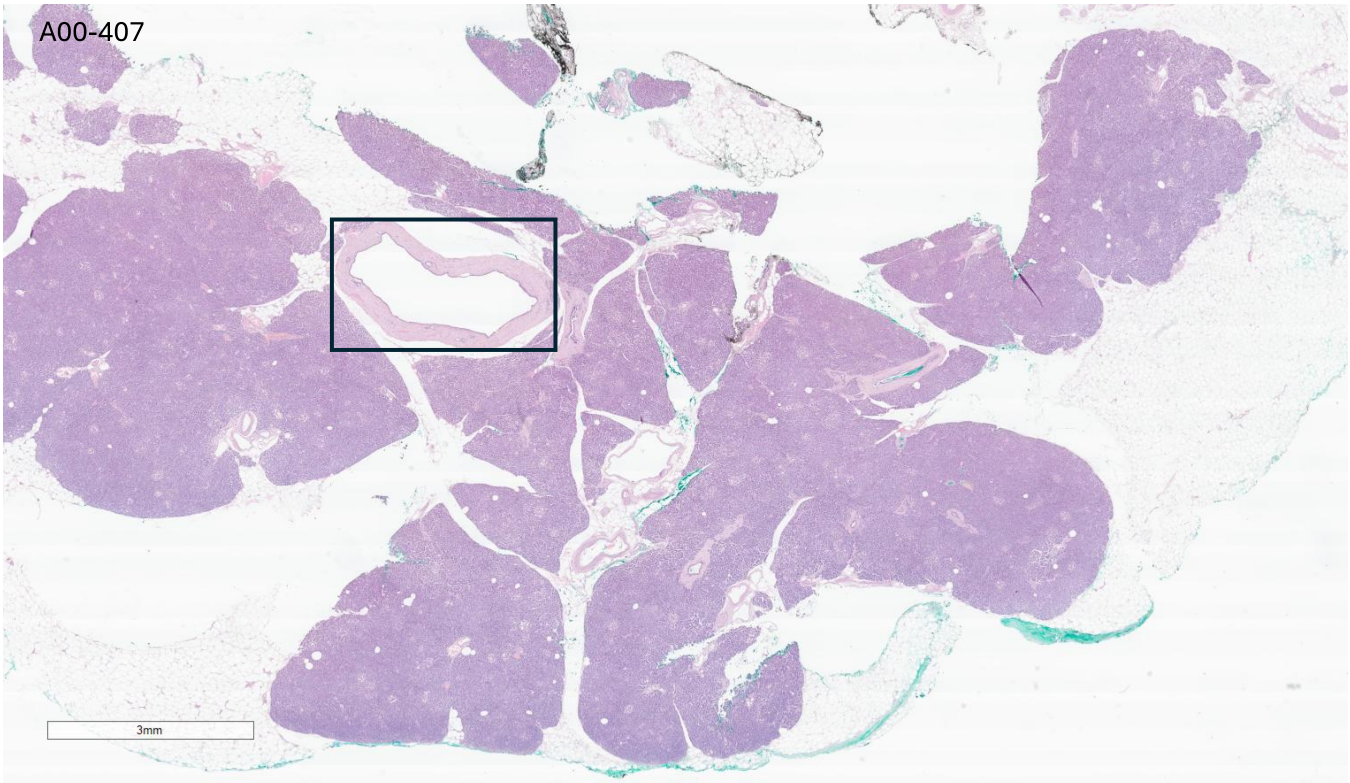


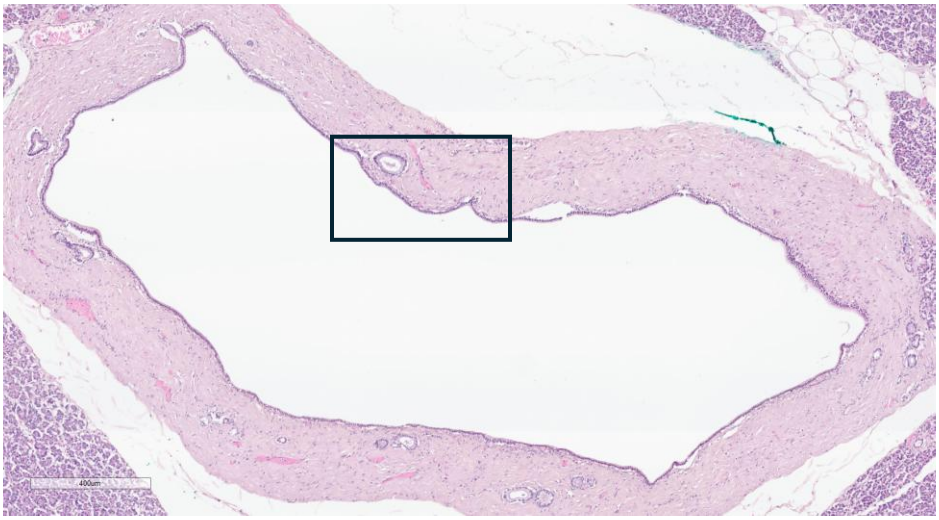


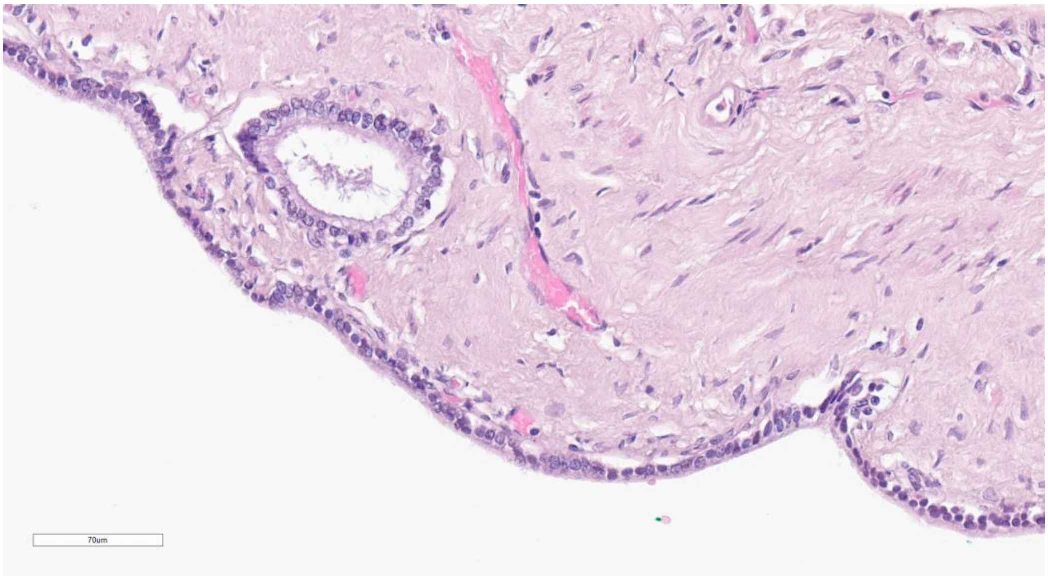

A01-339

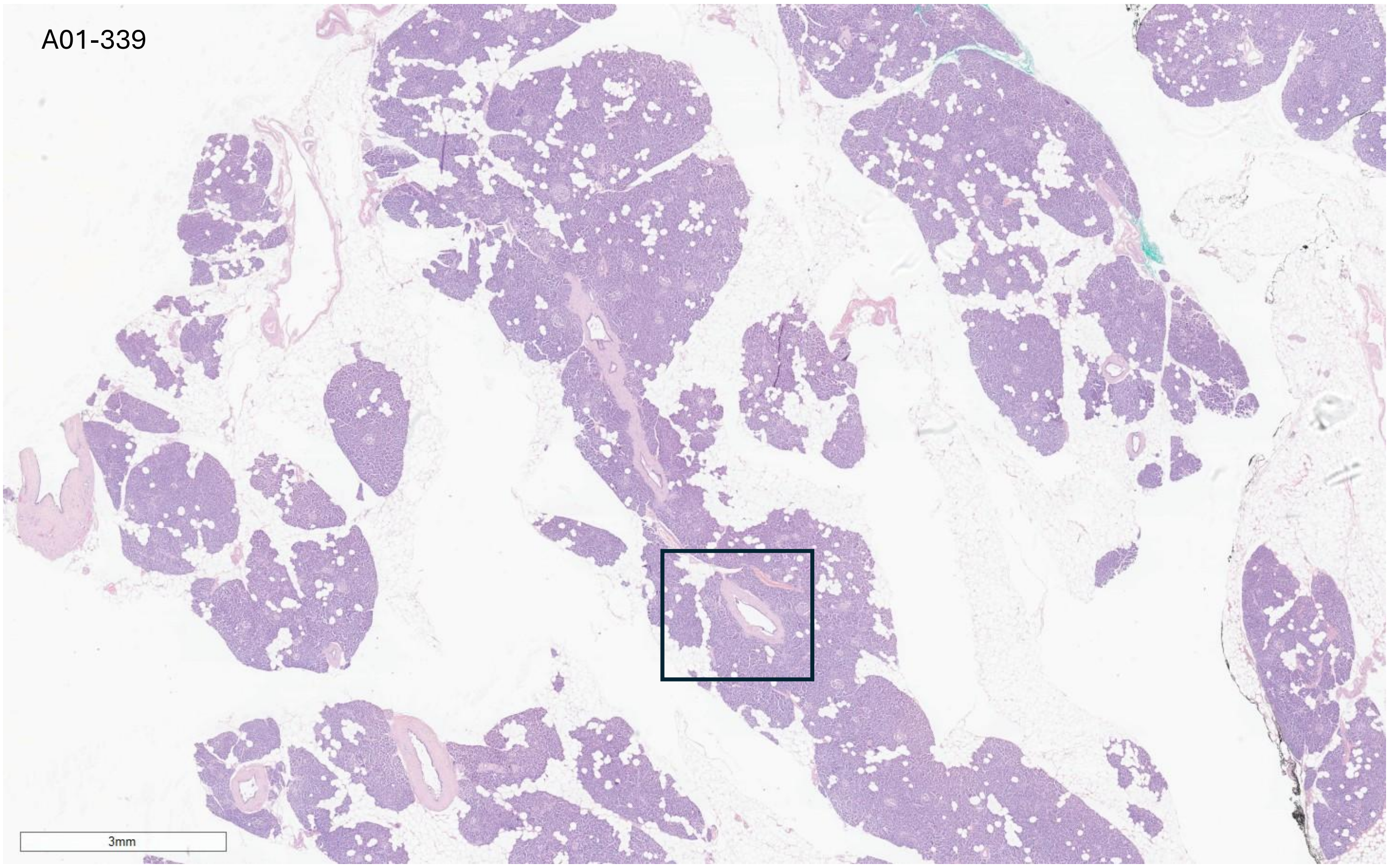


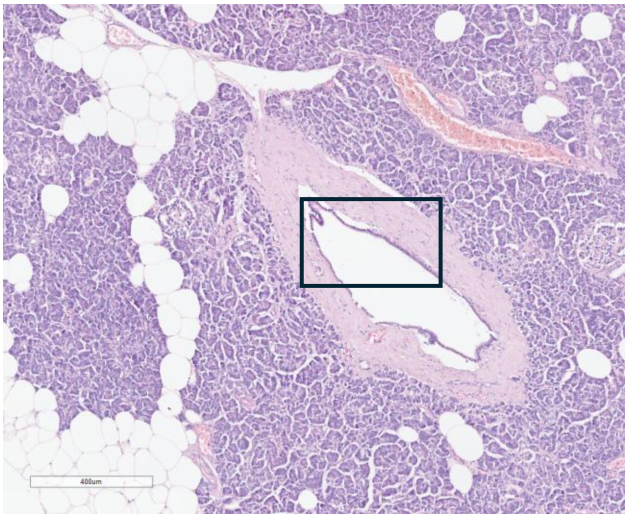


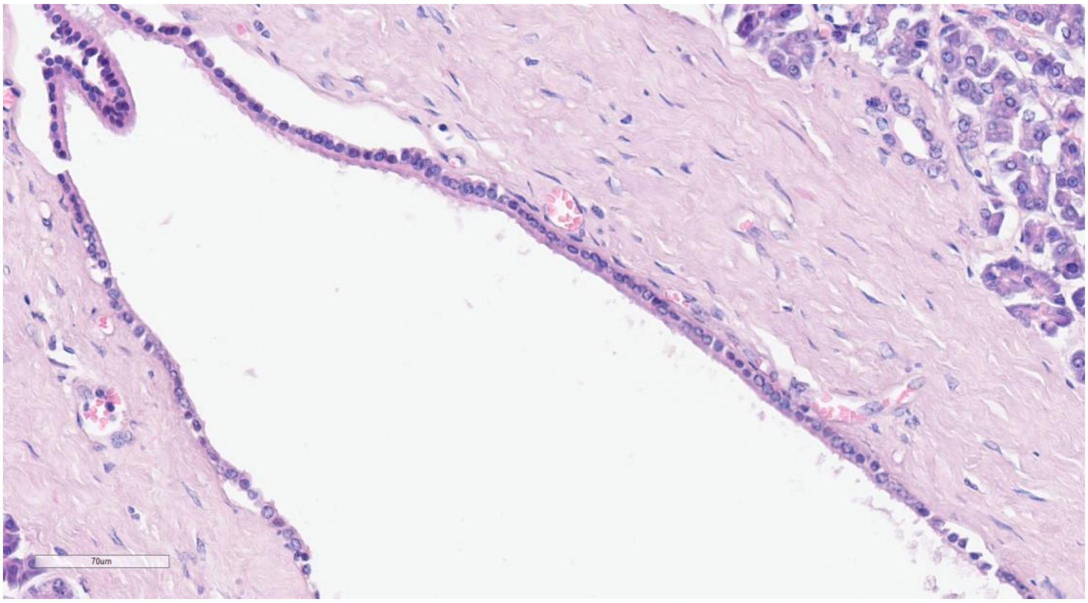

A01-340

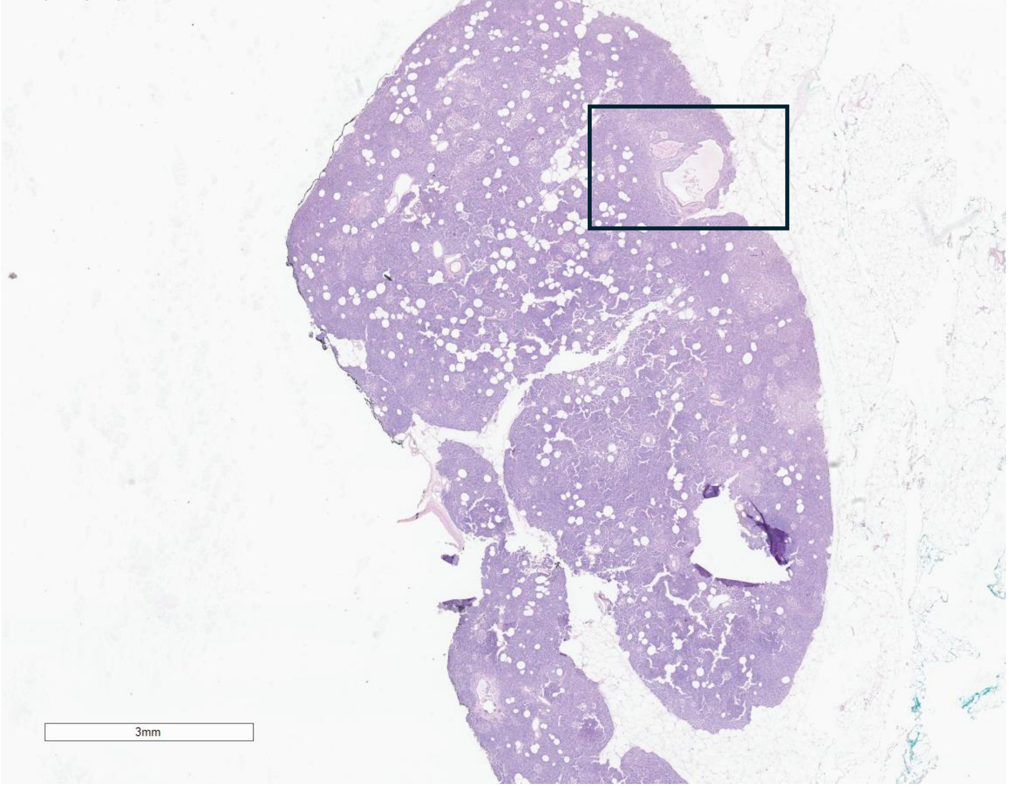


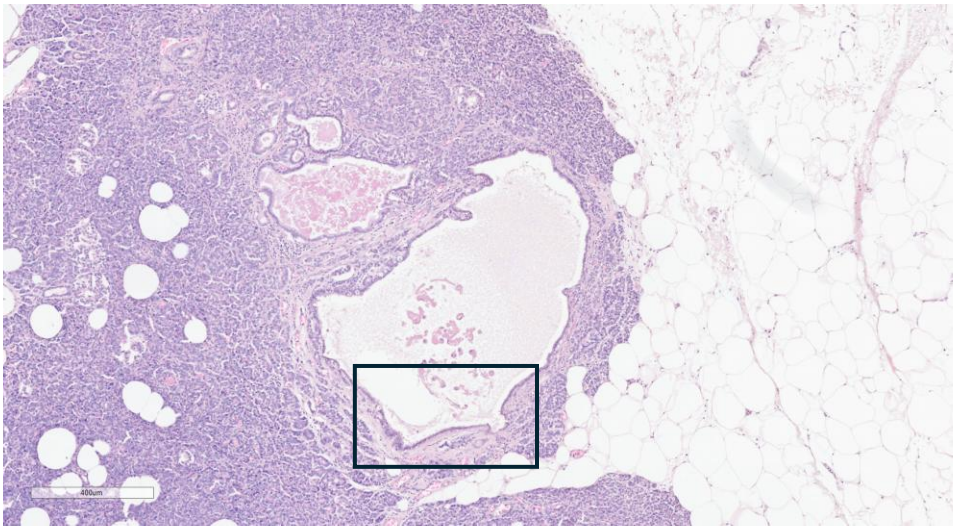


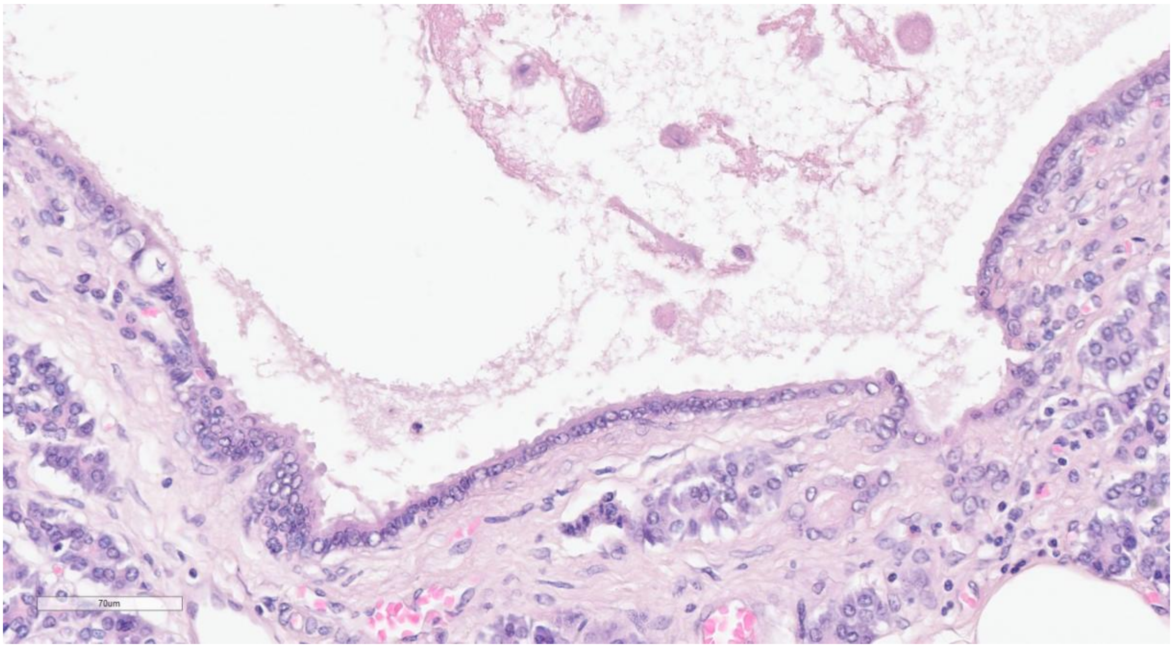

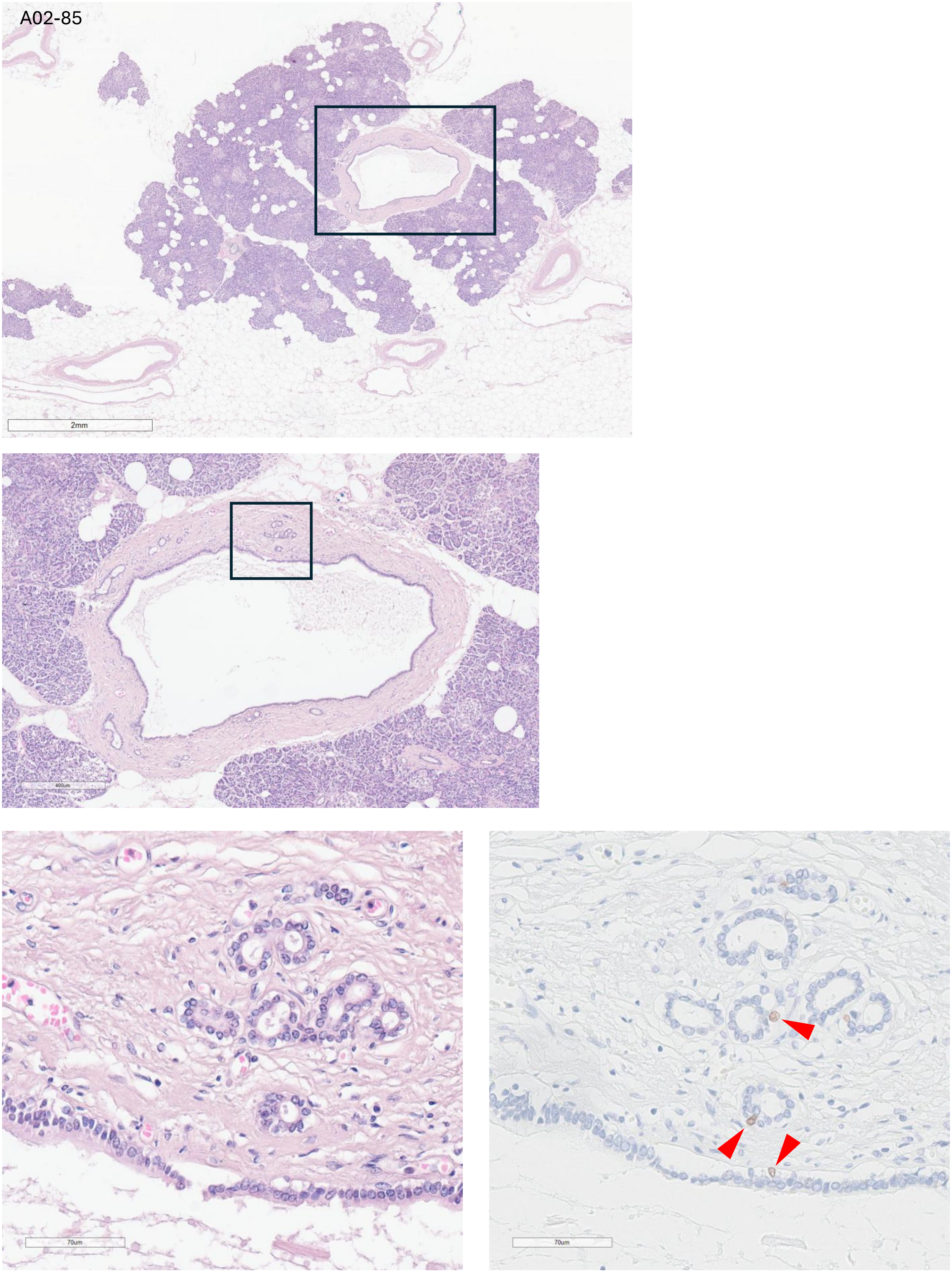
A02-85
2mm
400um
70um
70um

A02-120

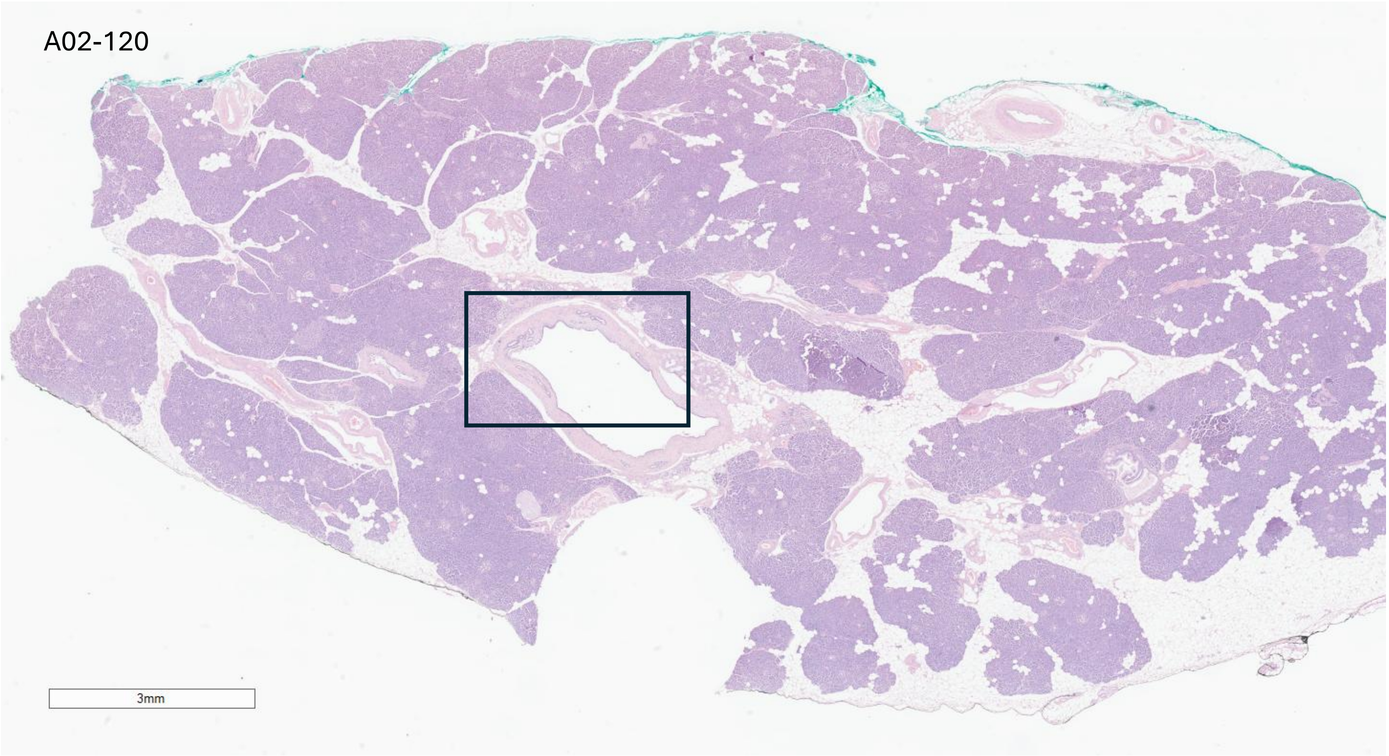


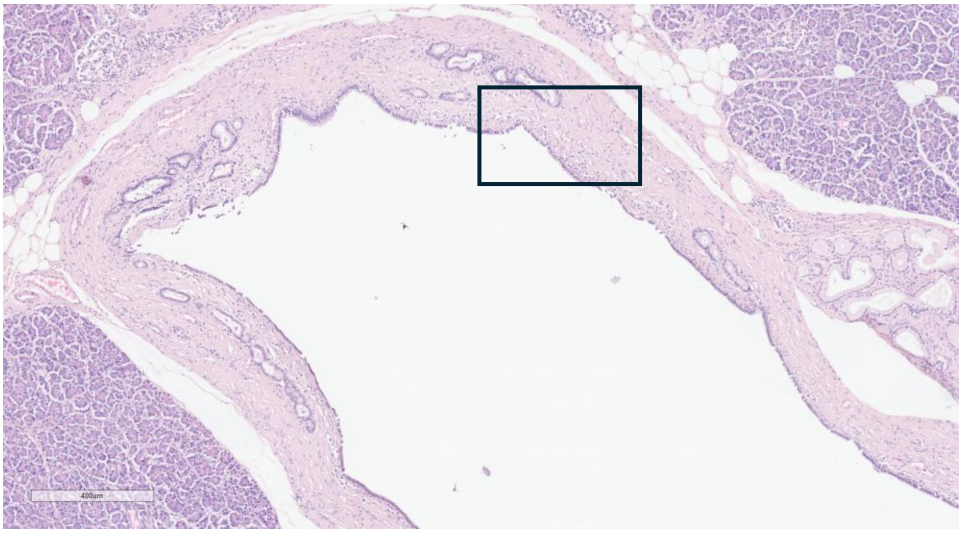


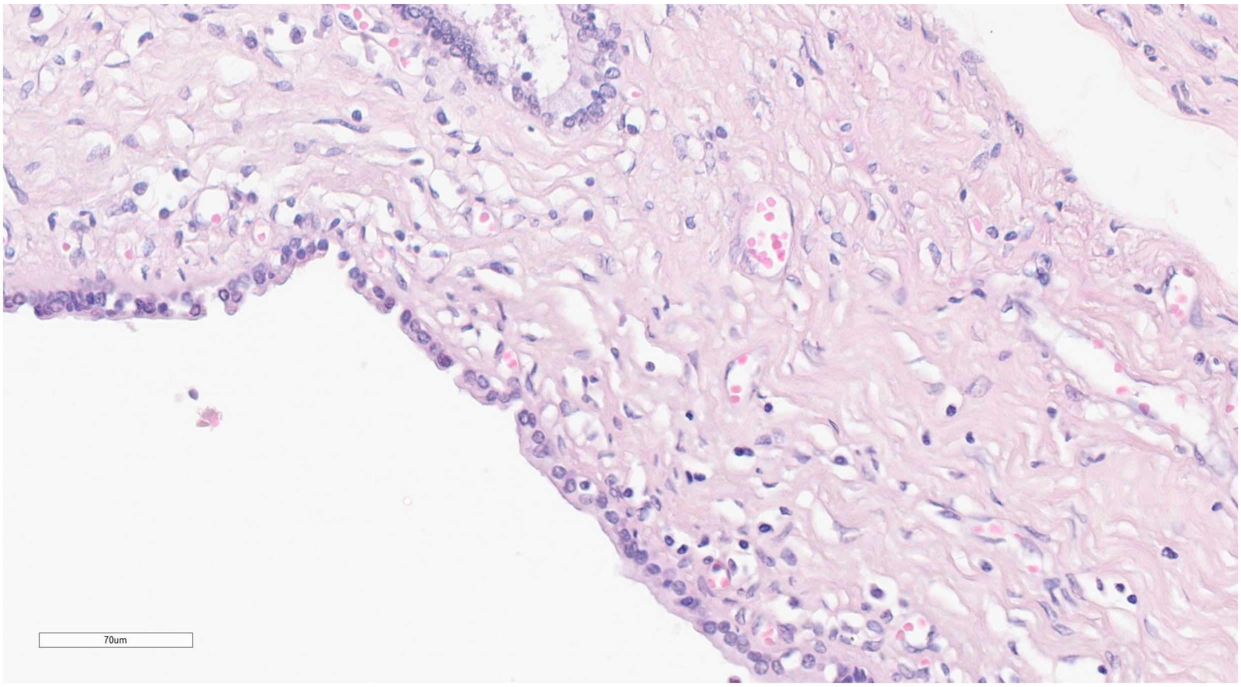

A02-134

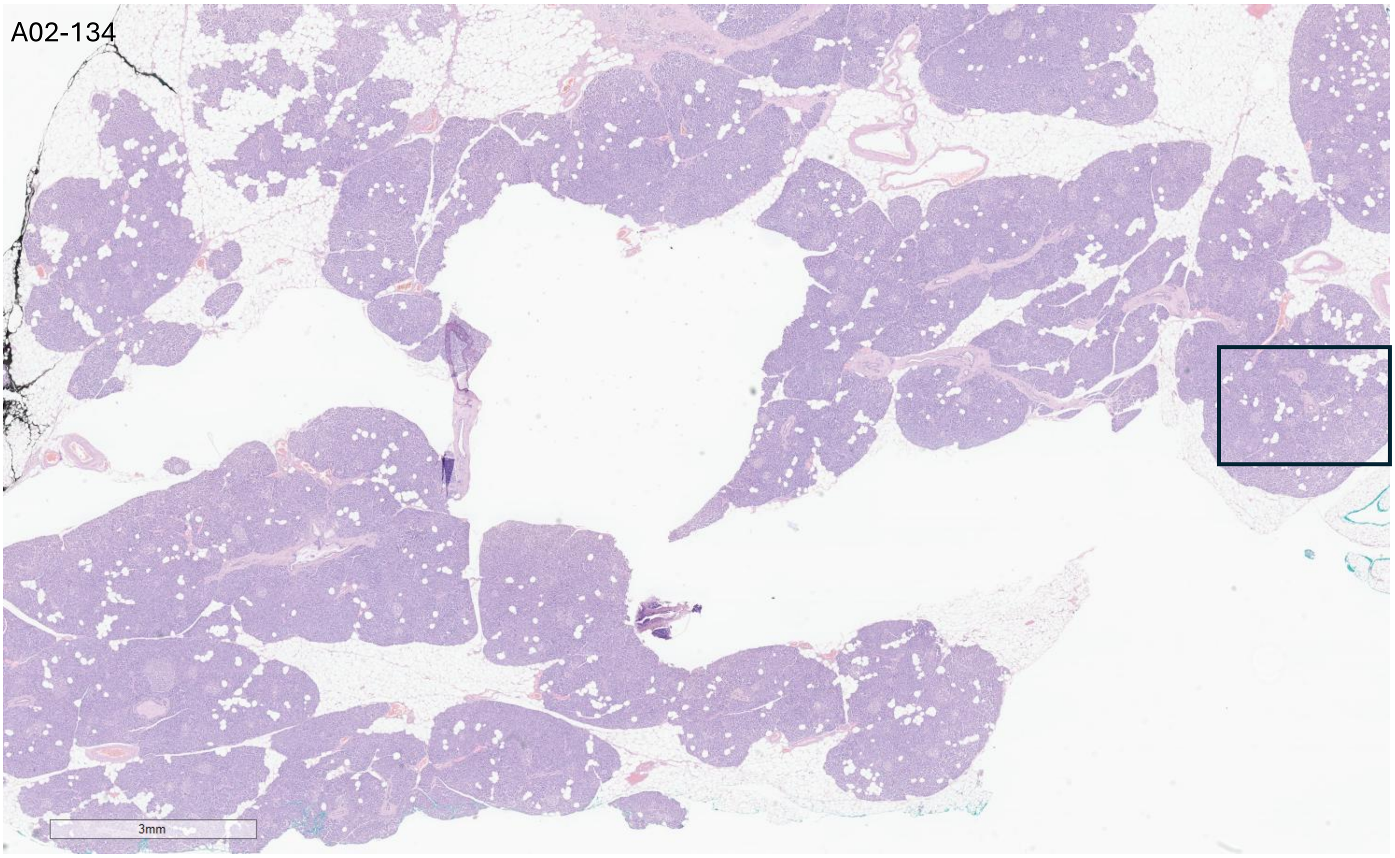


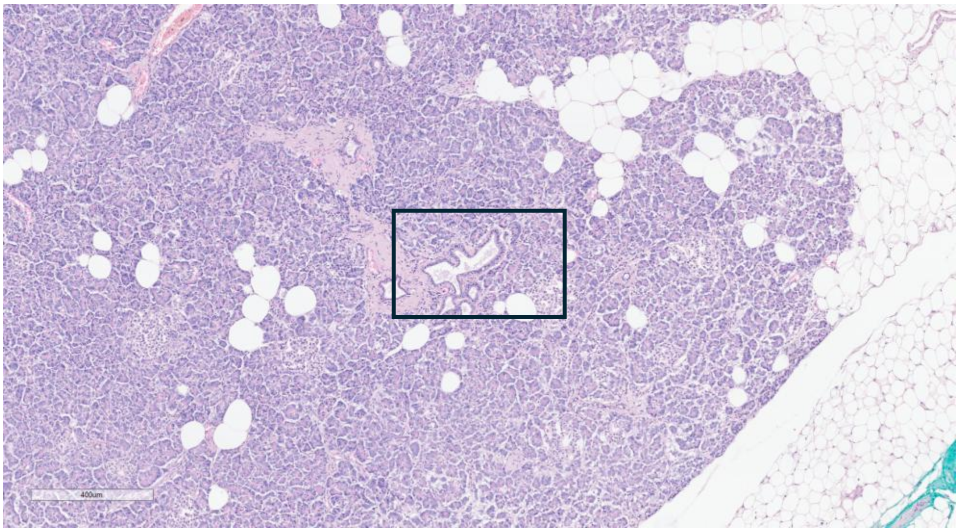


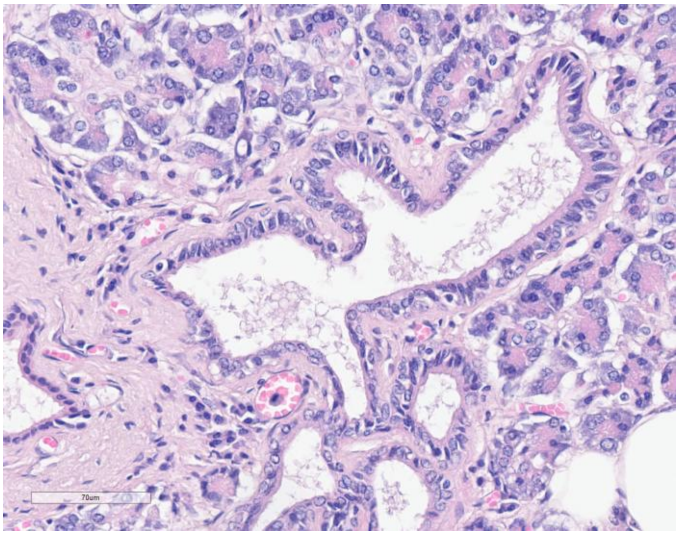


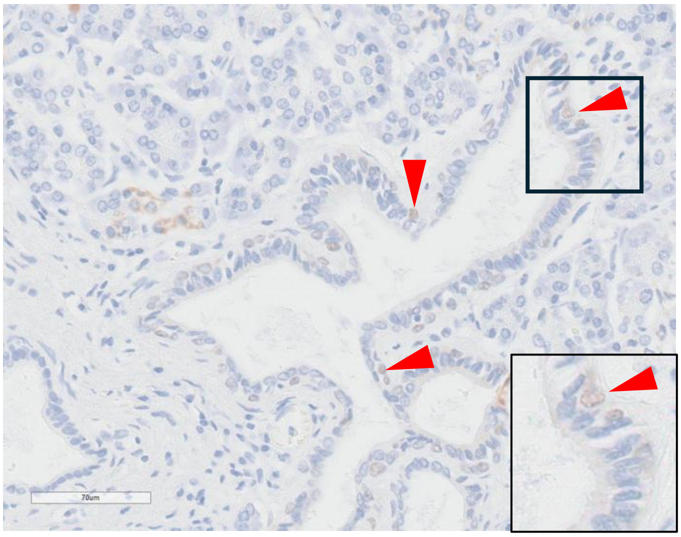

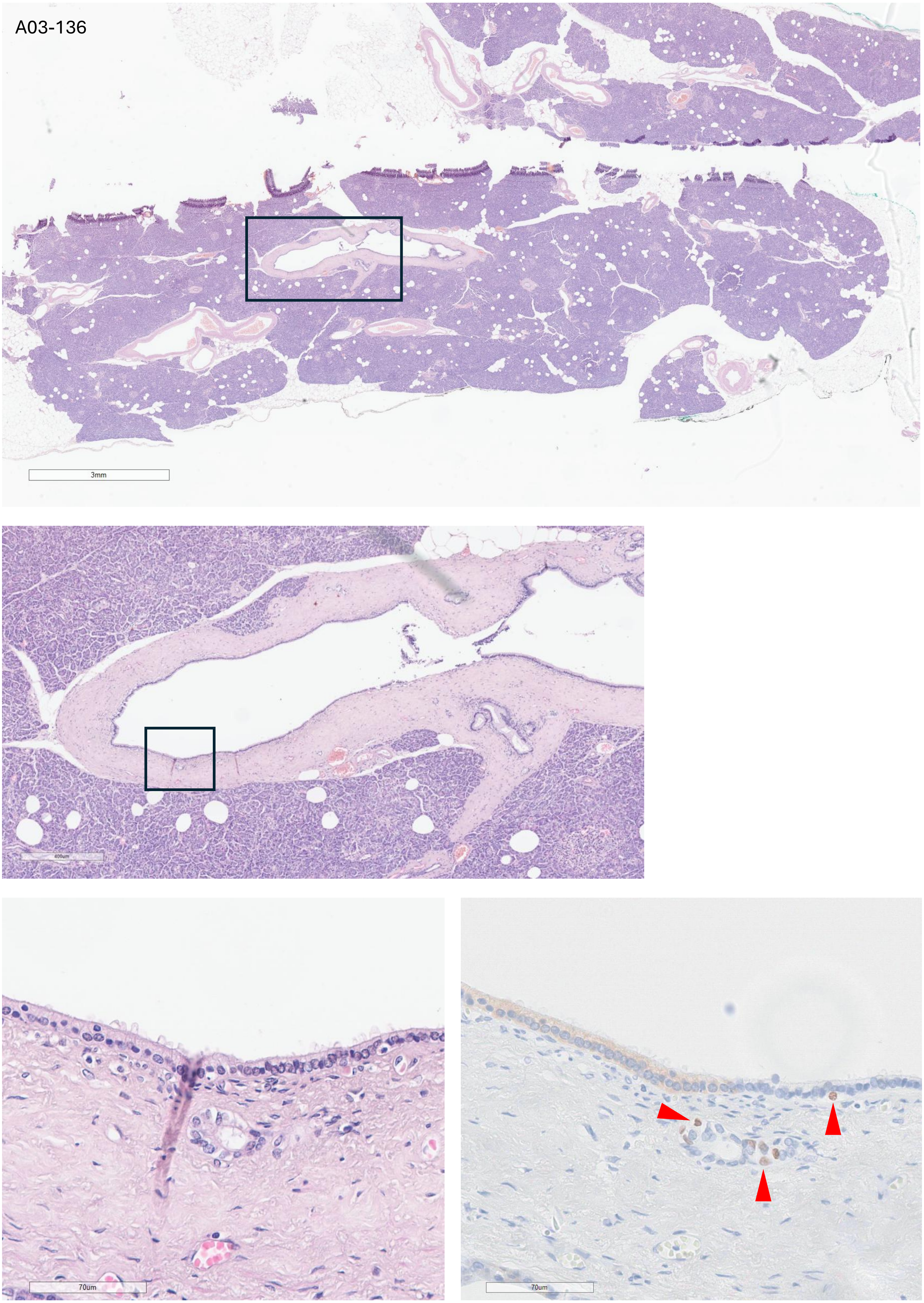
A03-136
3mm
400um
70um
70um

A04-107

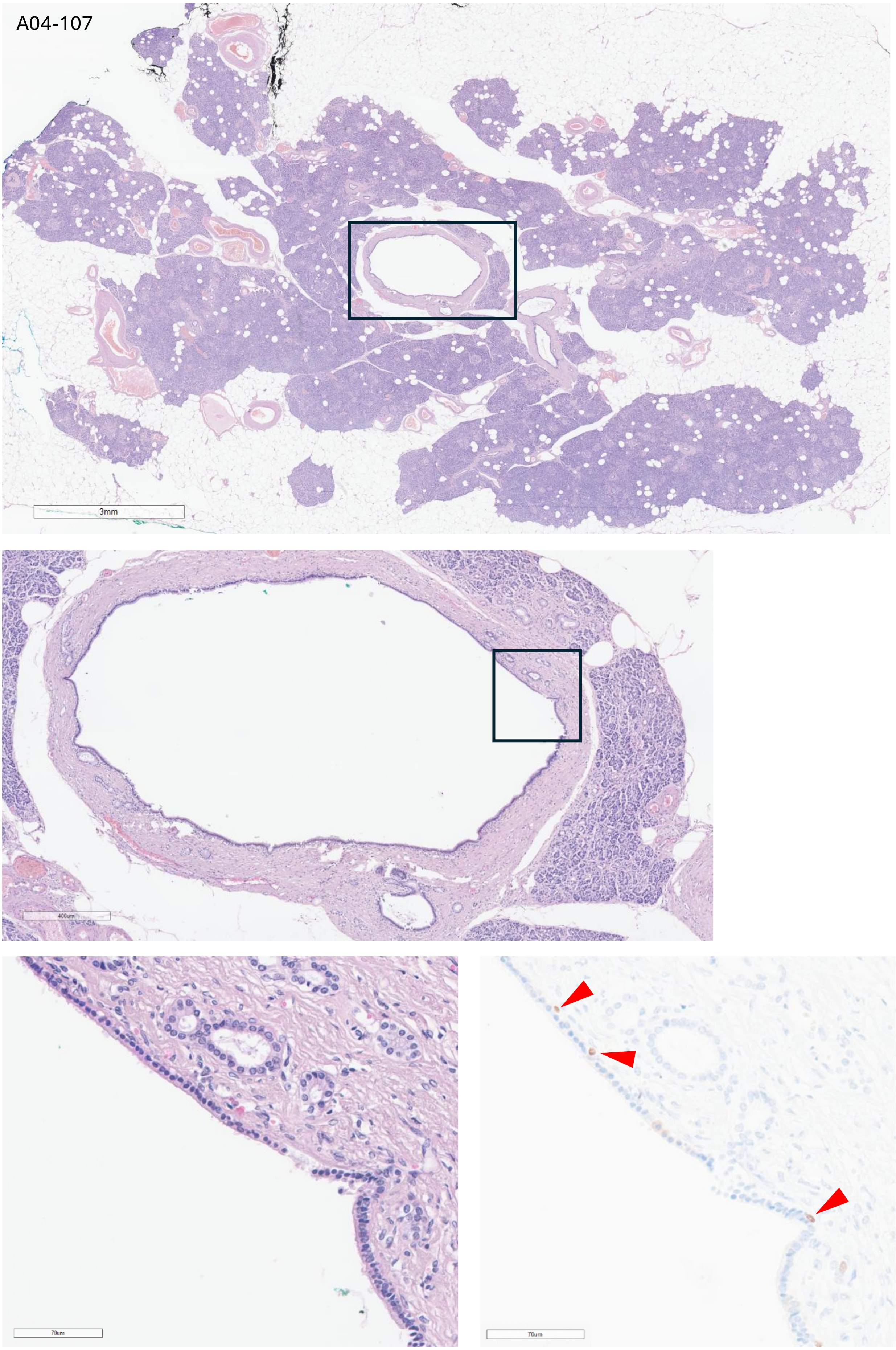

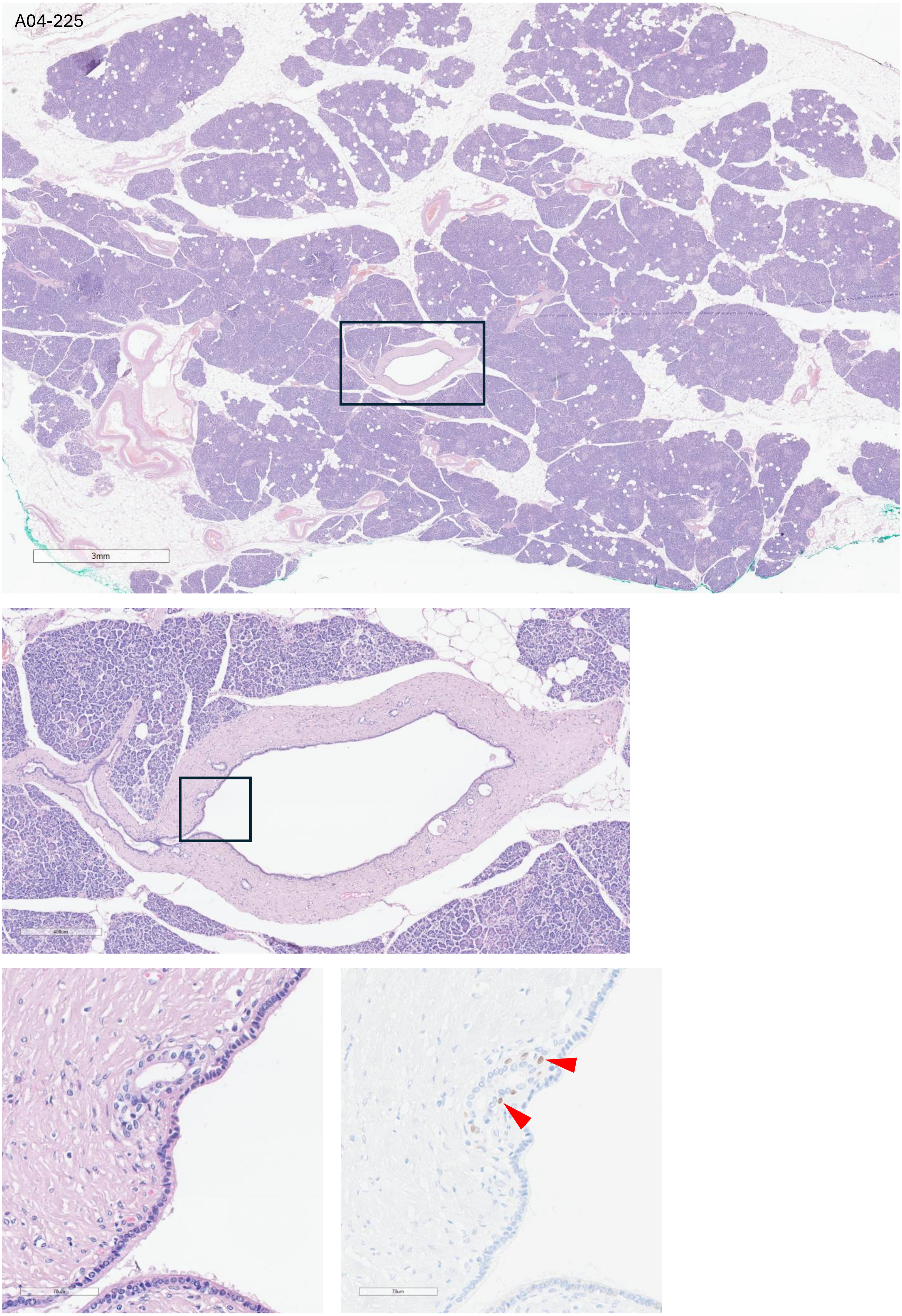
A04-225
3mm
400um
70um
70um

A04-241

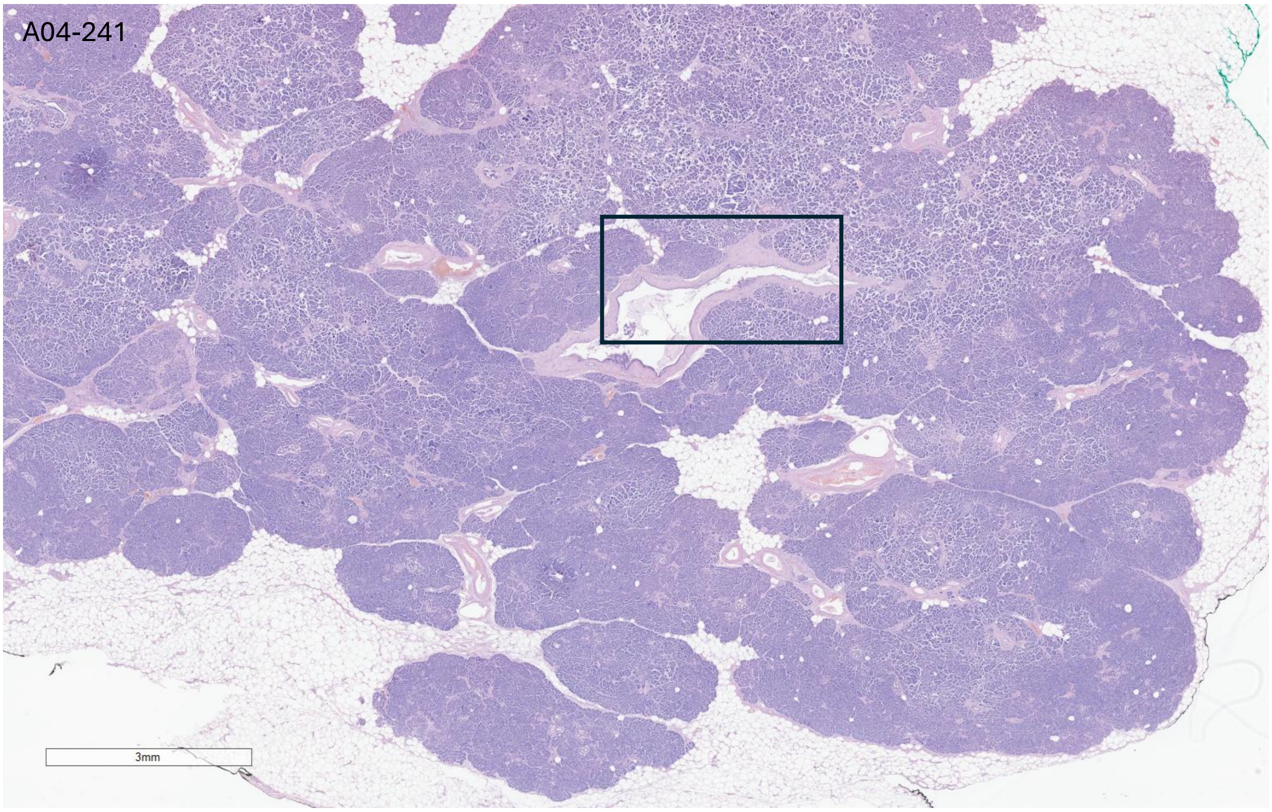


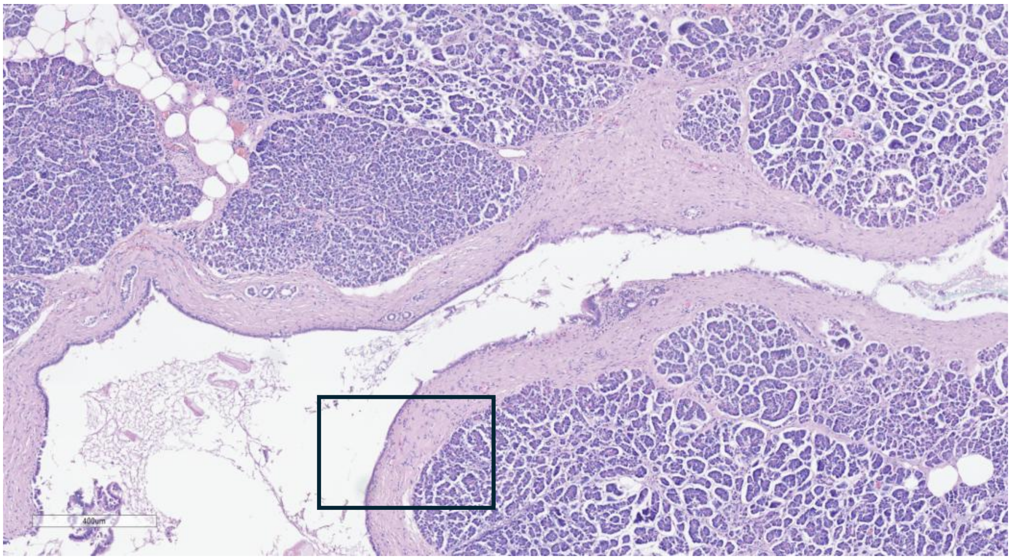


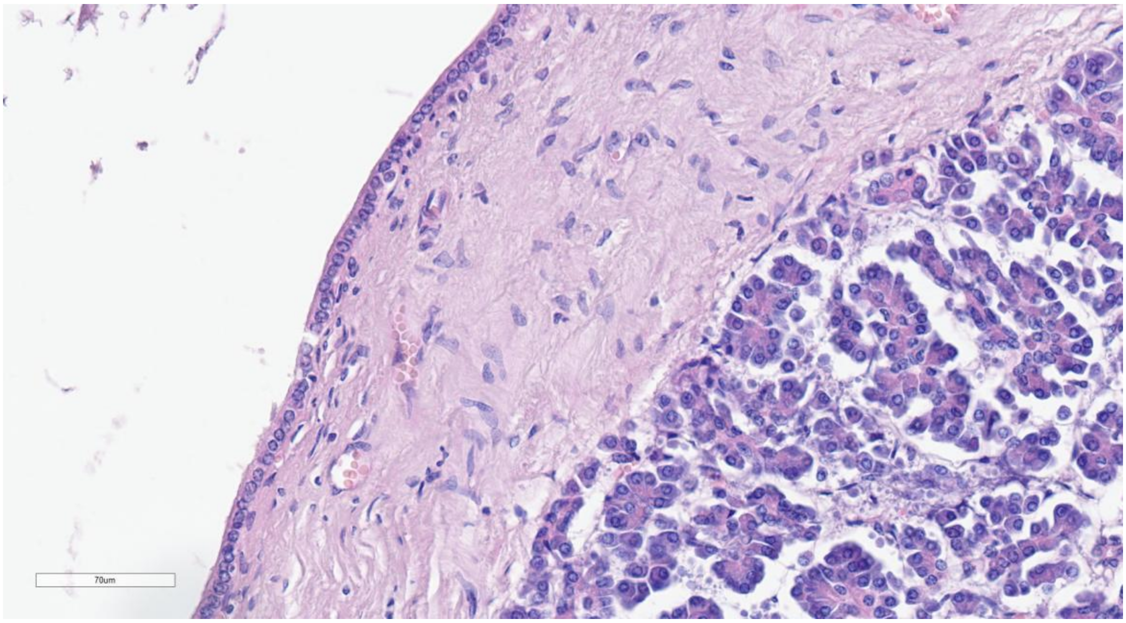

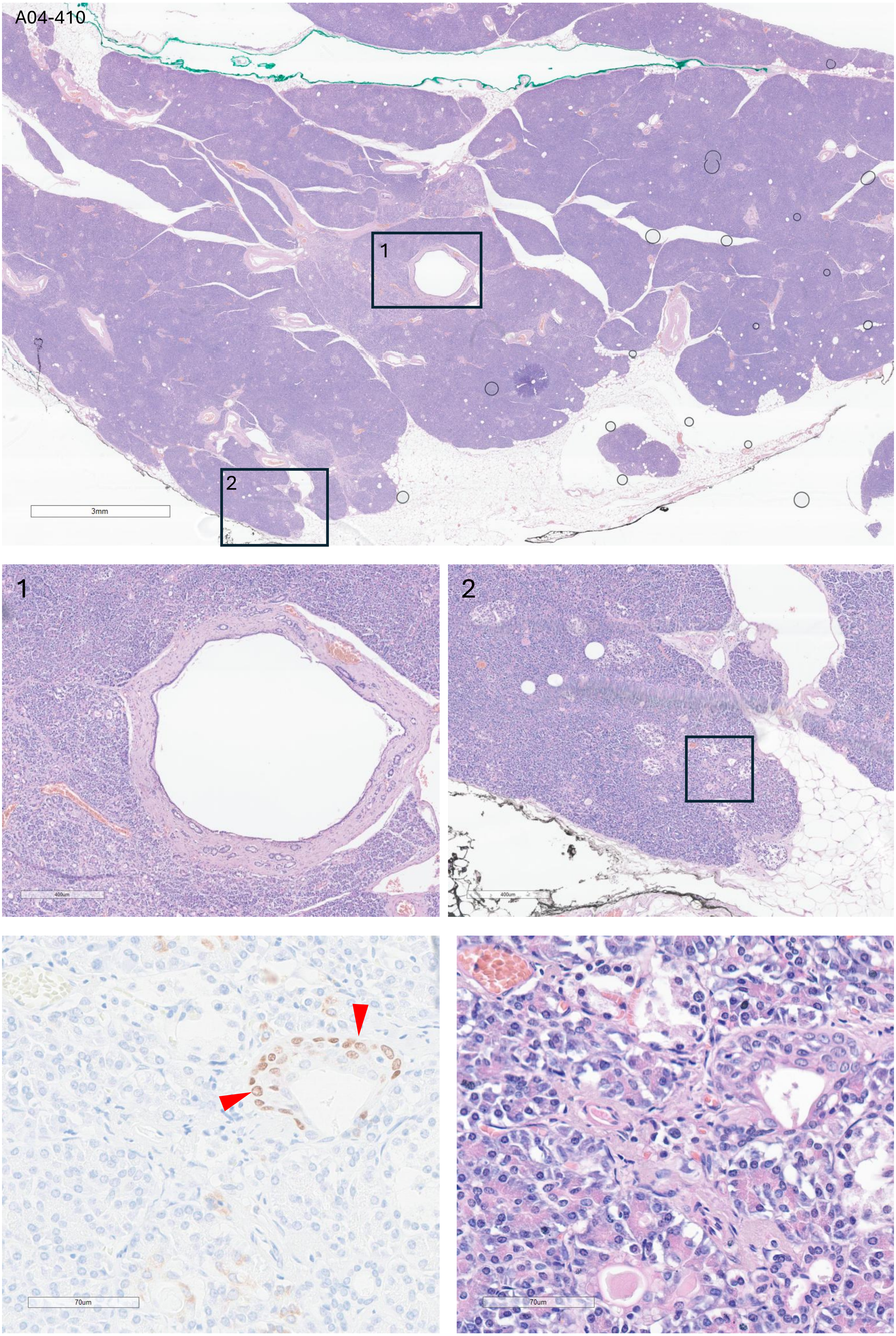
A04-410
1
2
3mm
400um
70um

A04-420

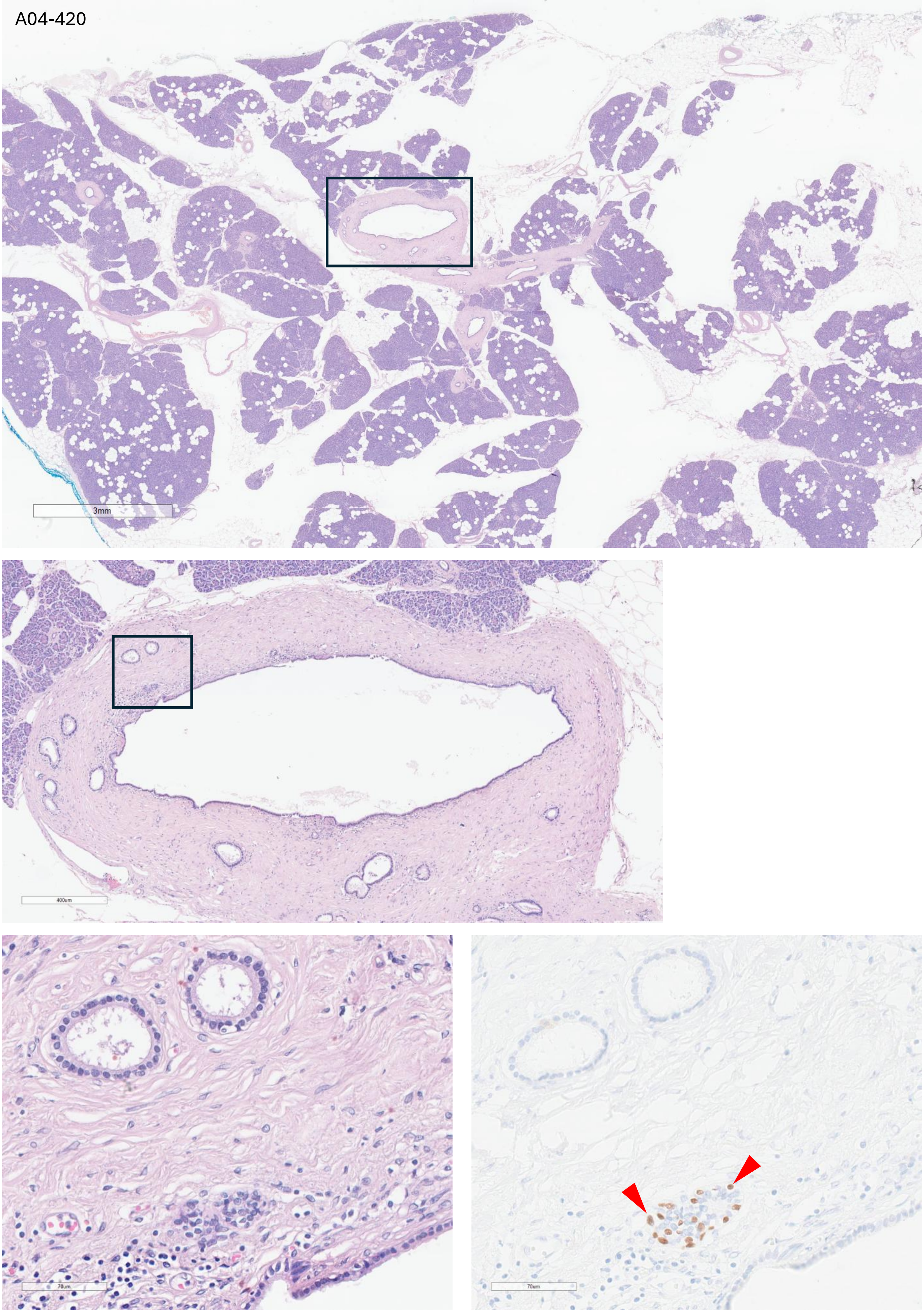

A05-64

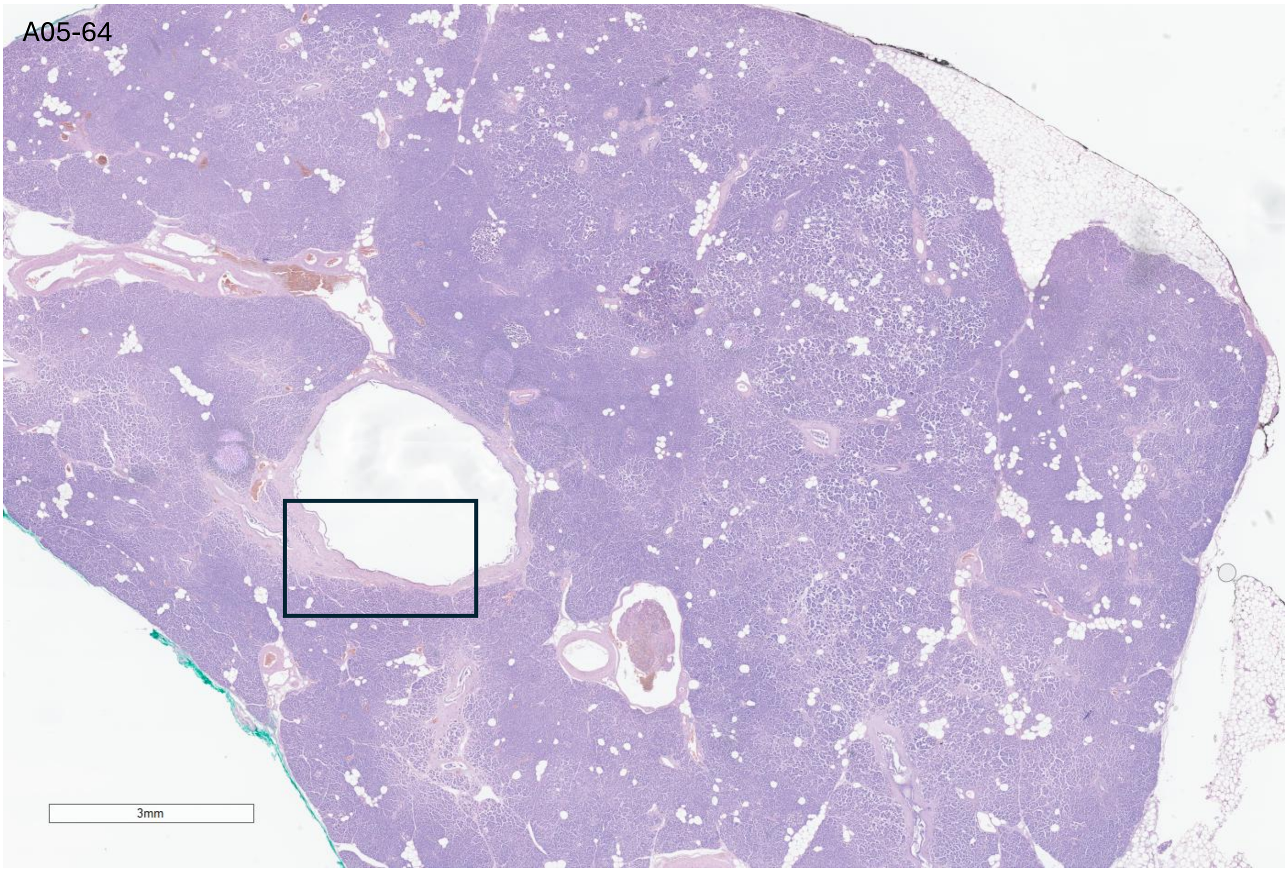


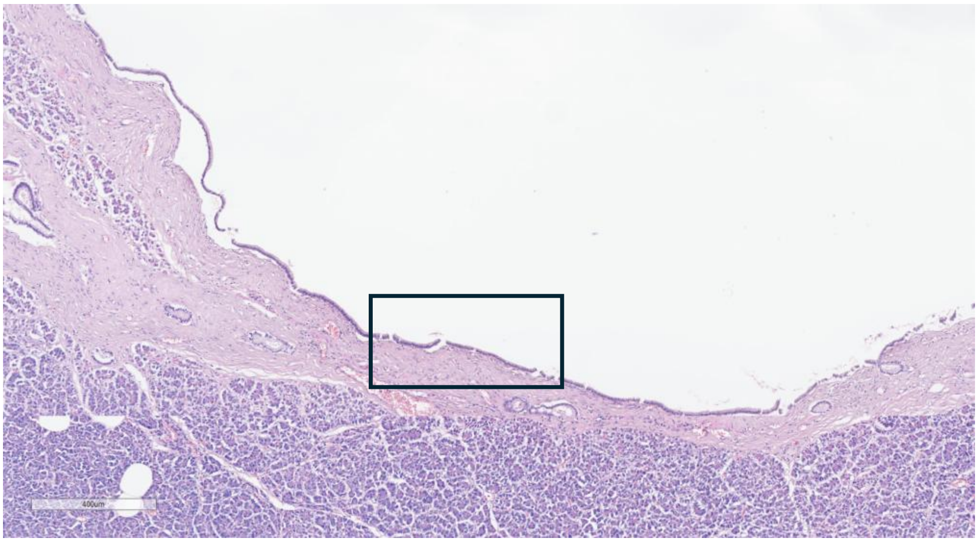


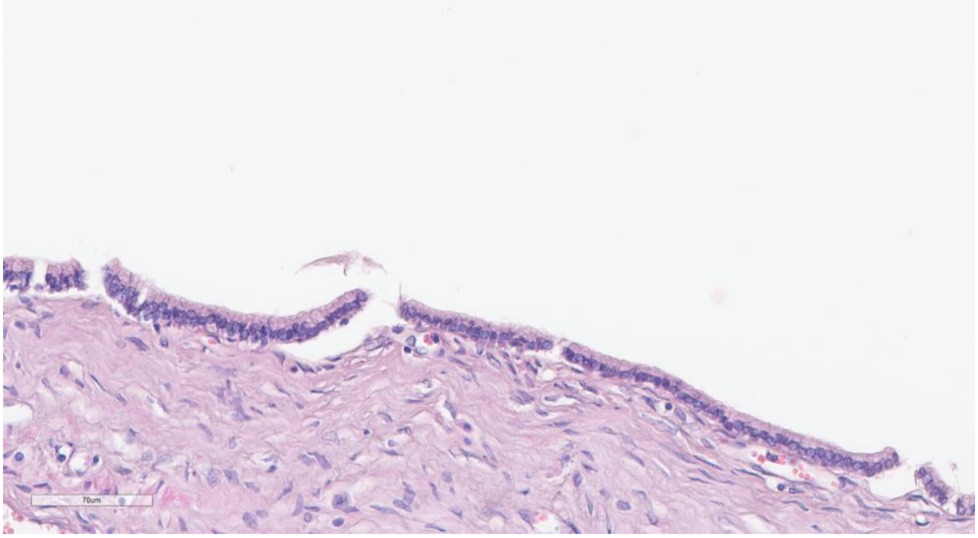

A05-405

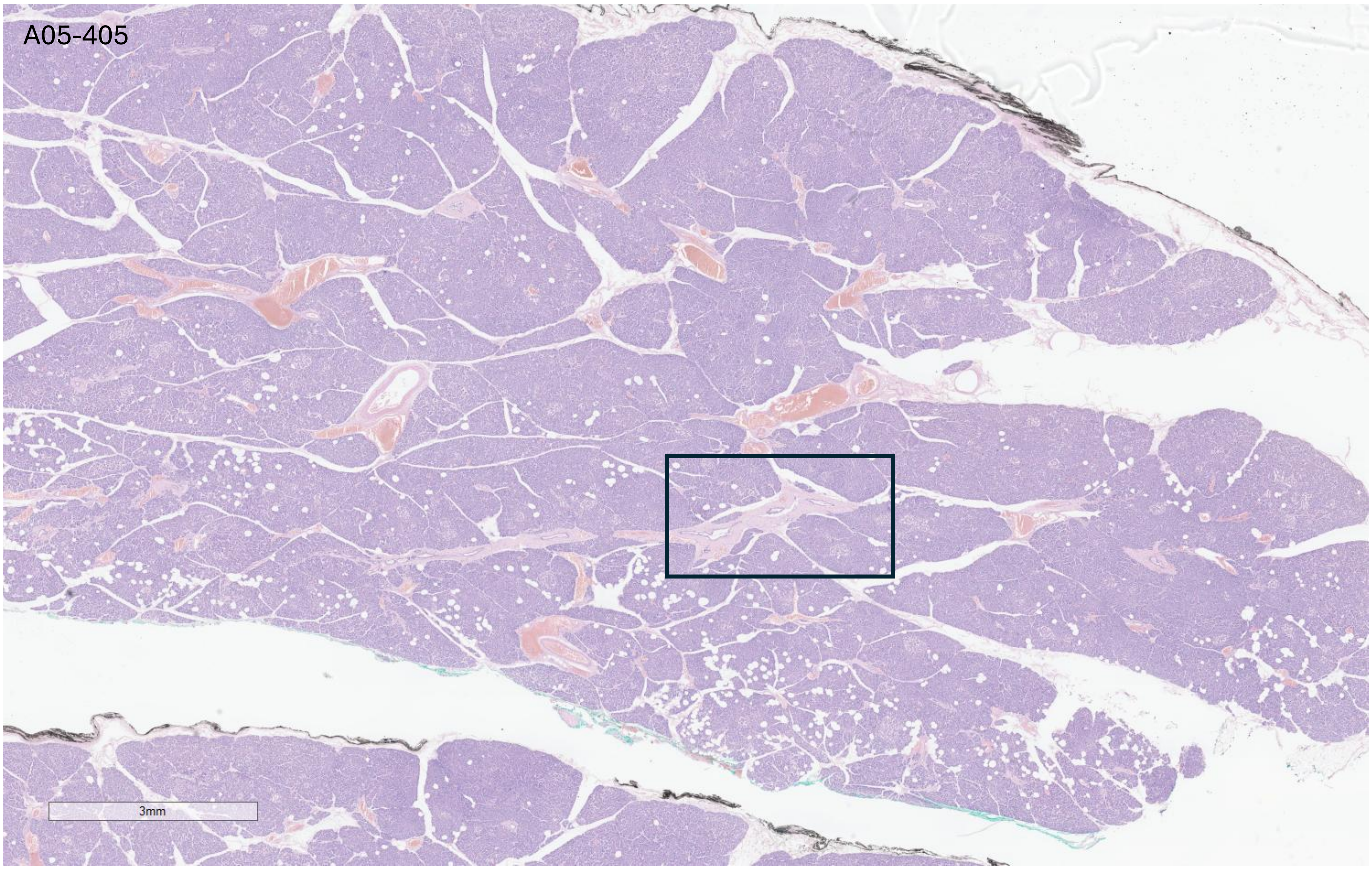


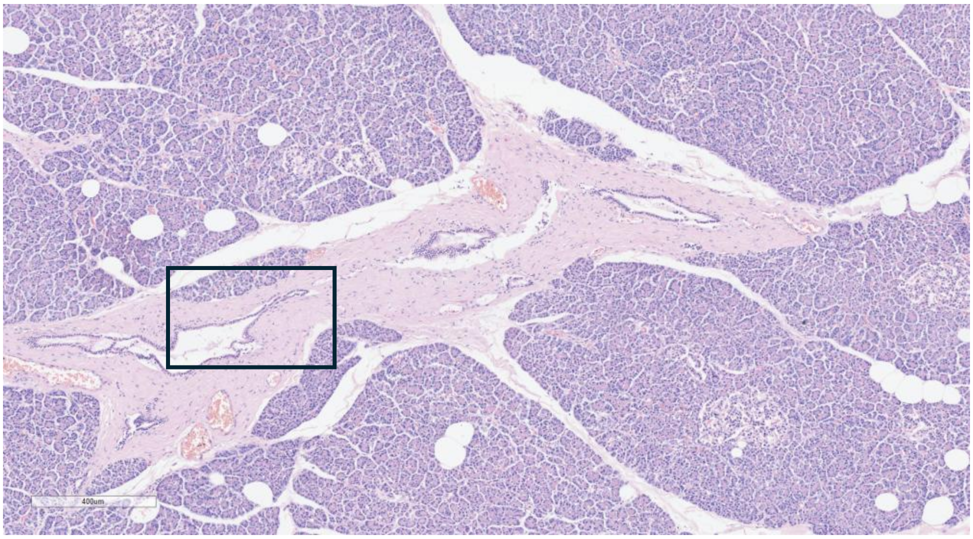


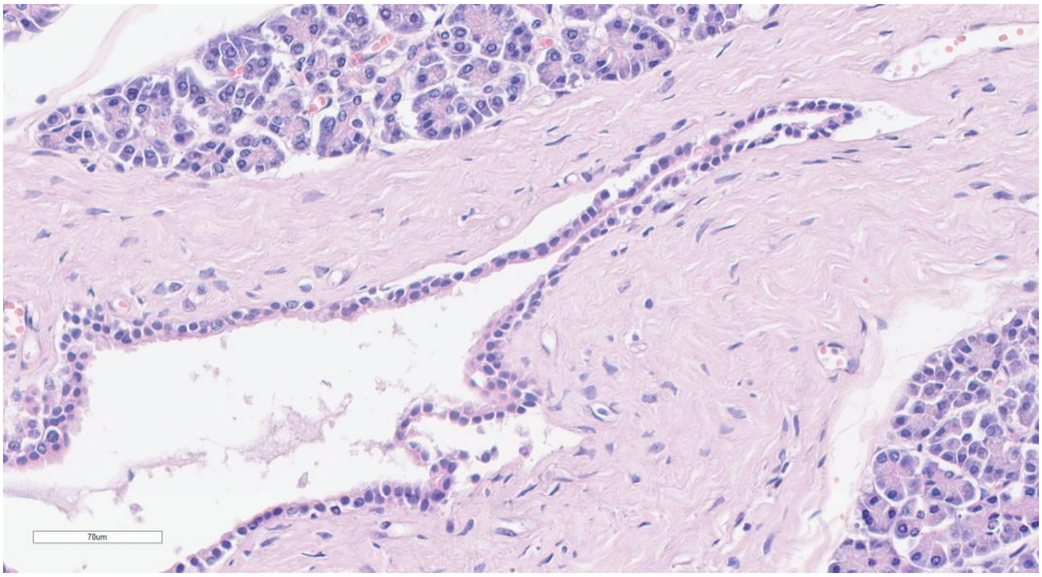

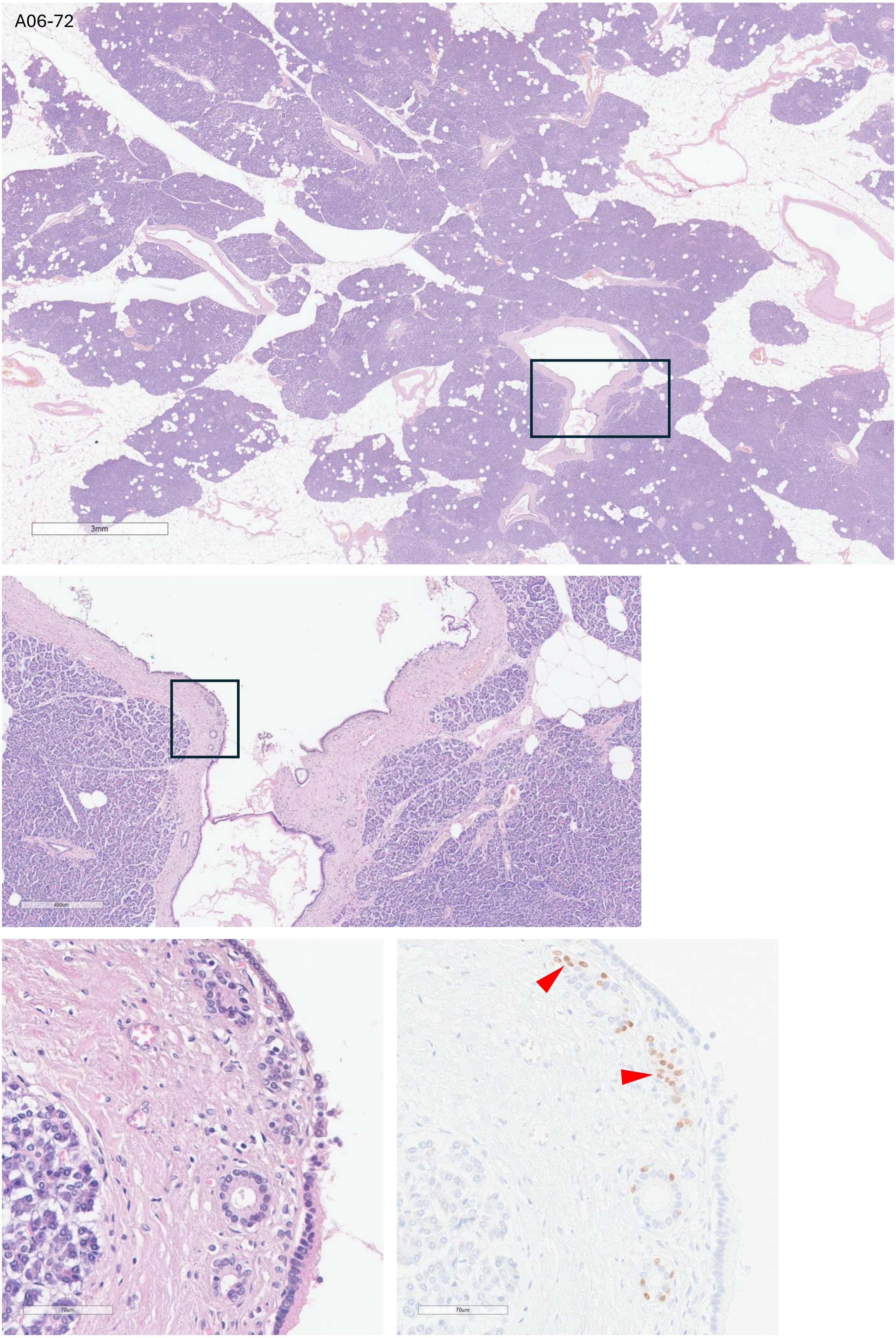
A06-72
3mm
400um
70um
70um

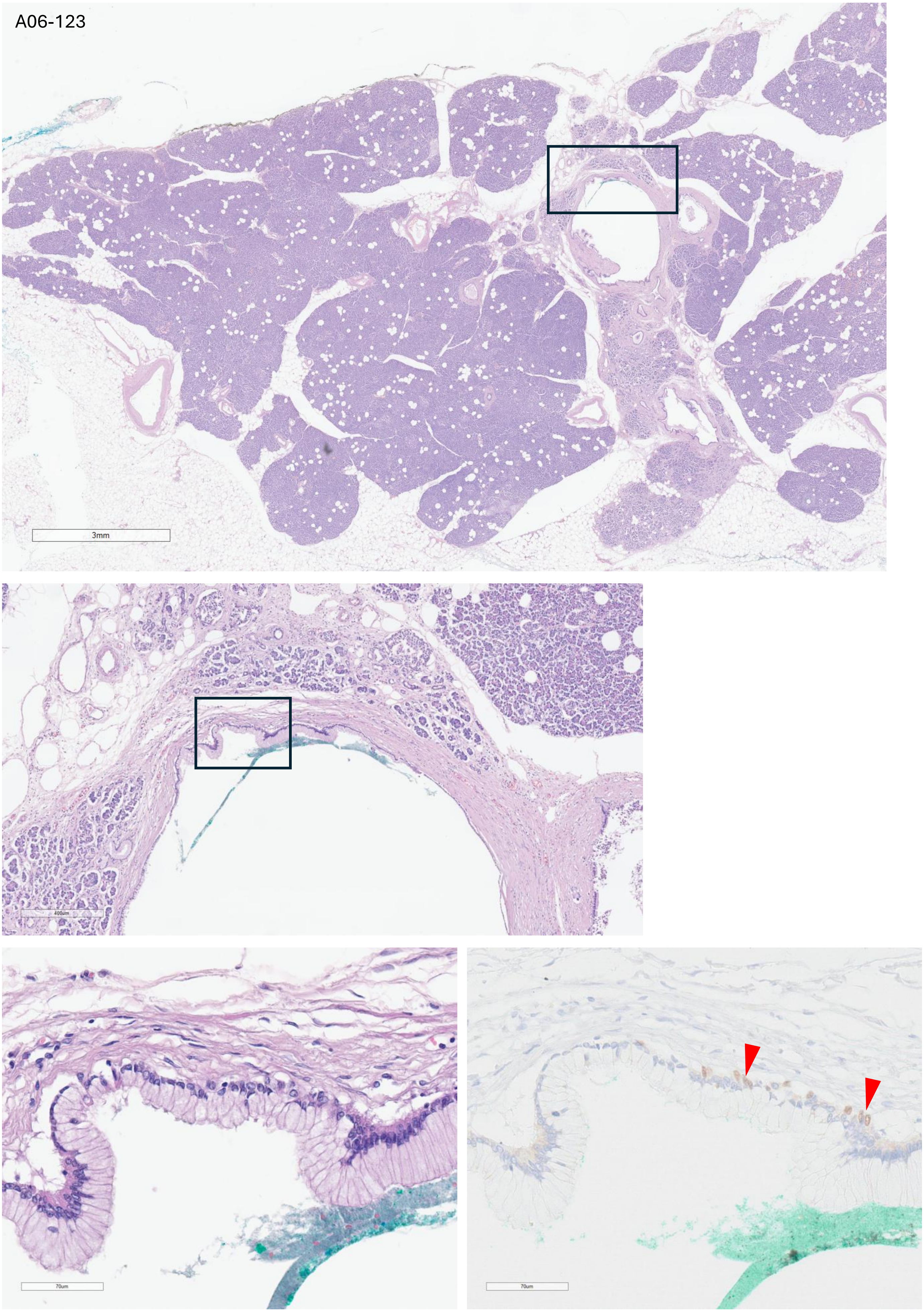
A06-123
3mm
400um
70um
70um

A08-060

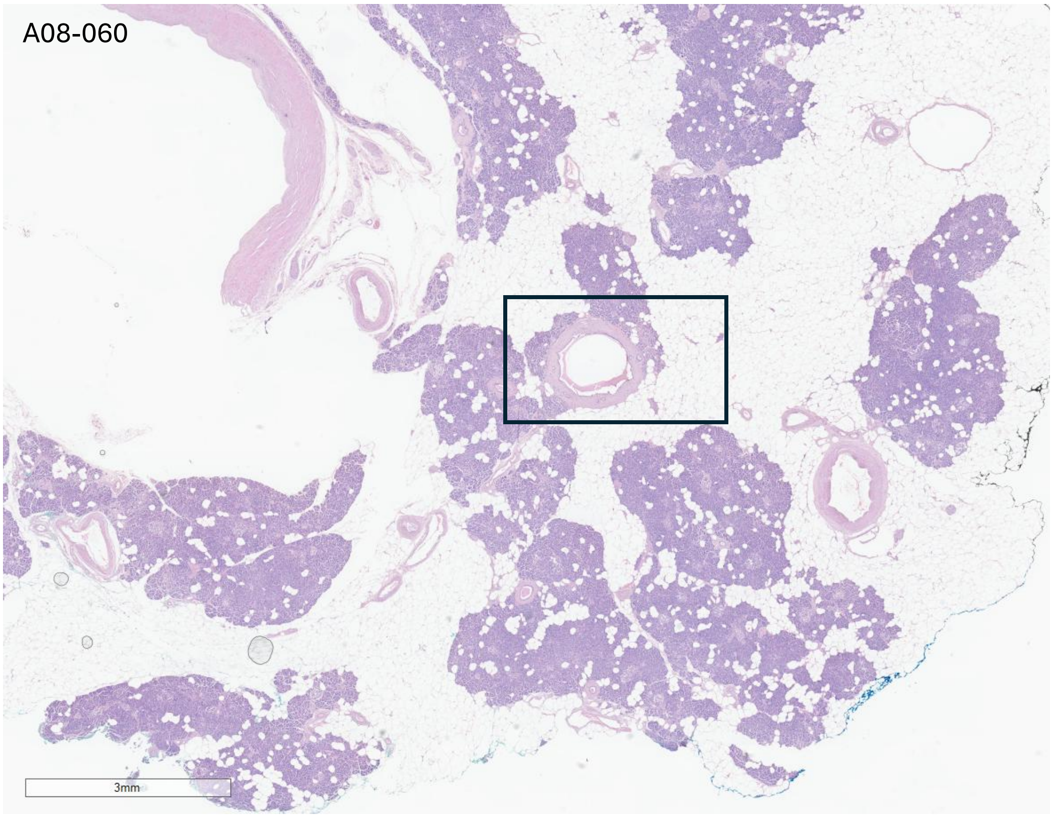


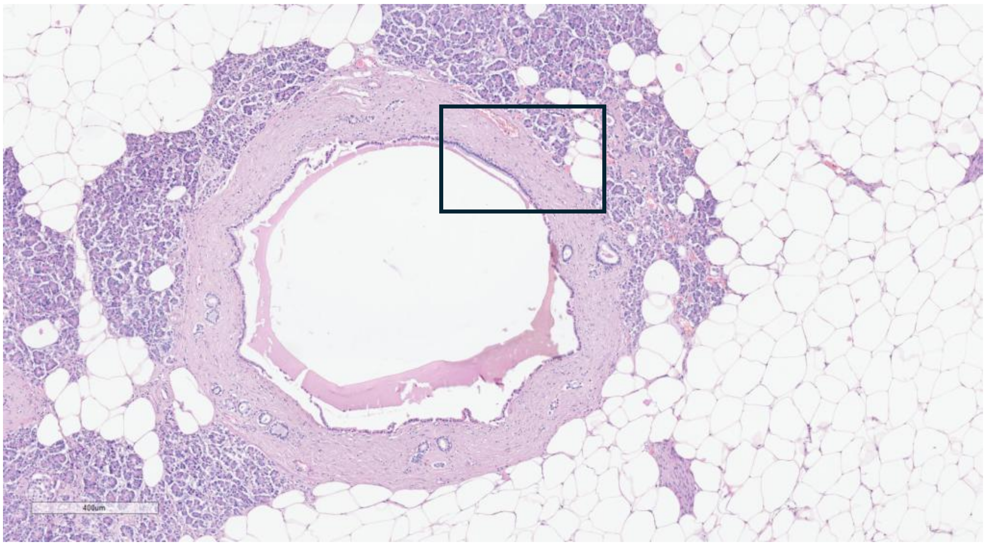


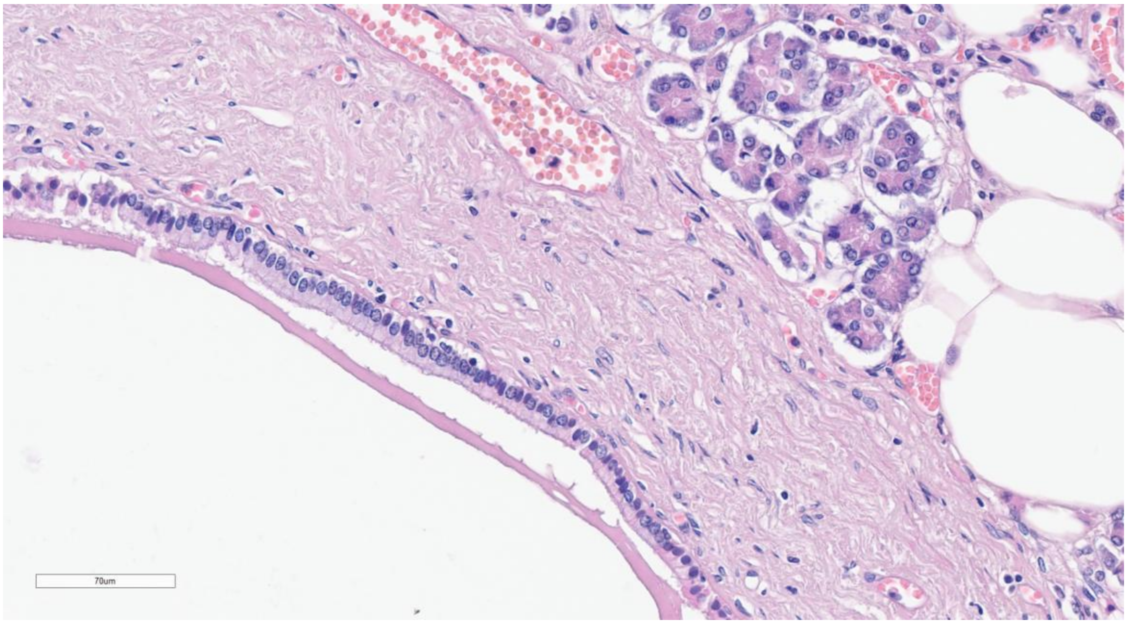

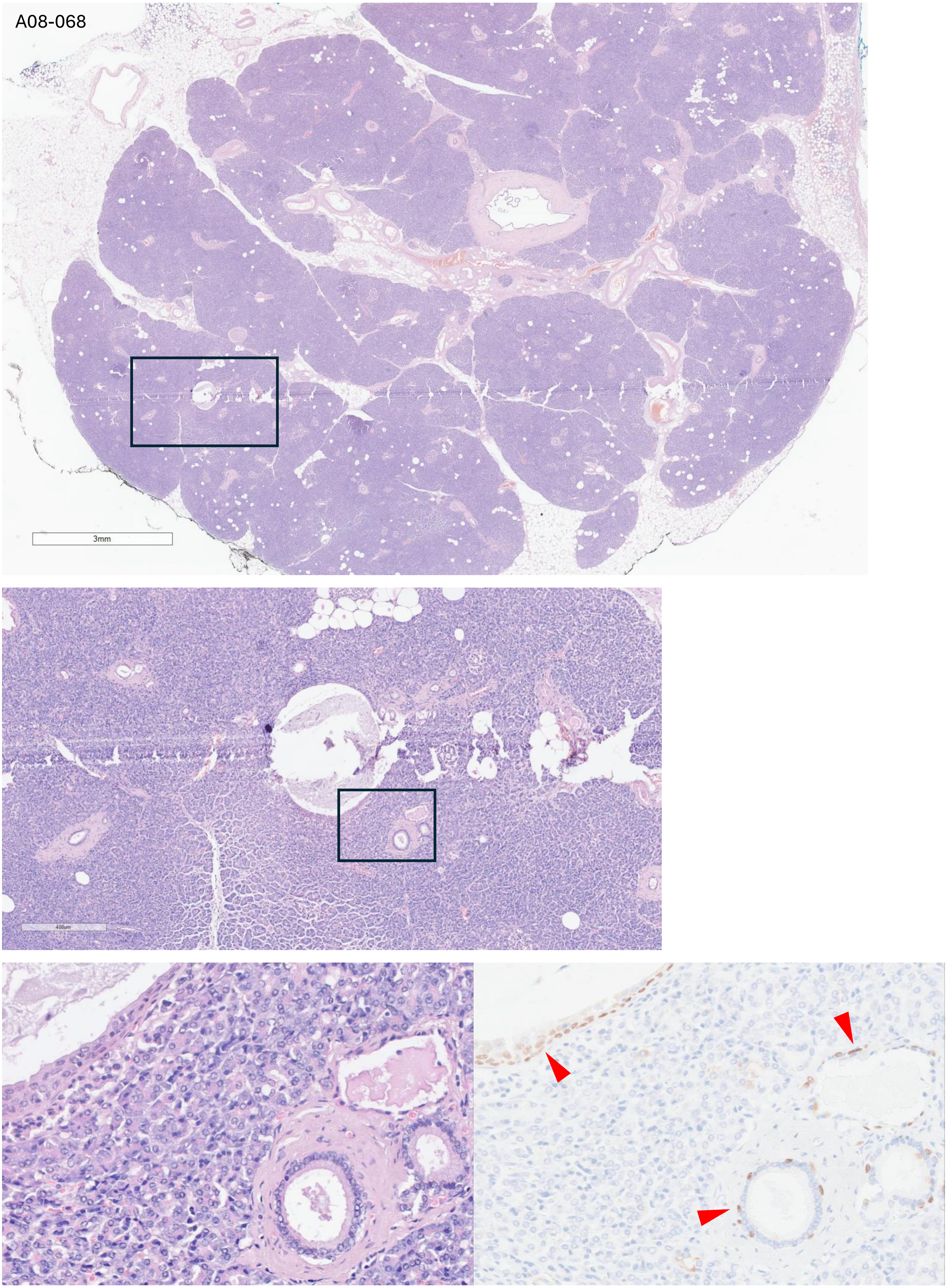
A08-068
3mm
400um

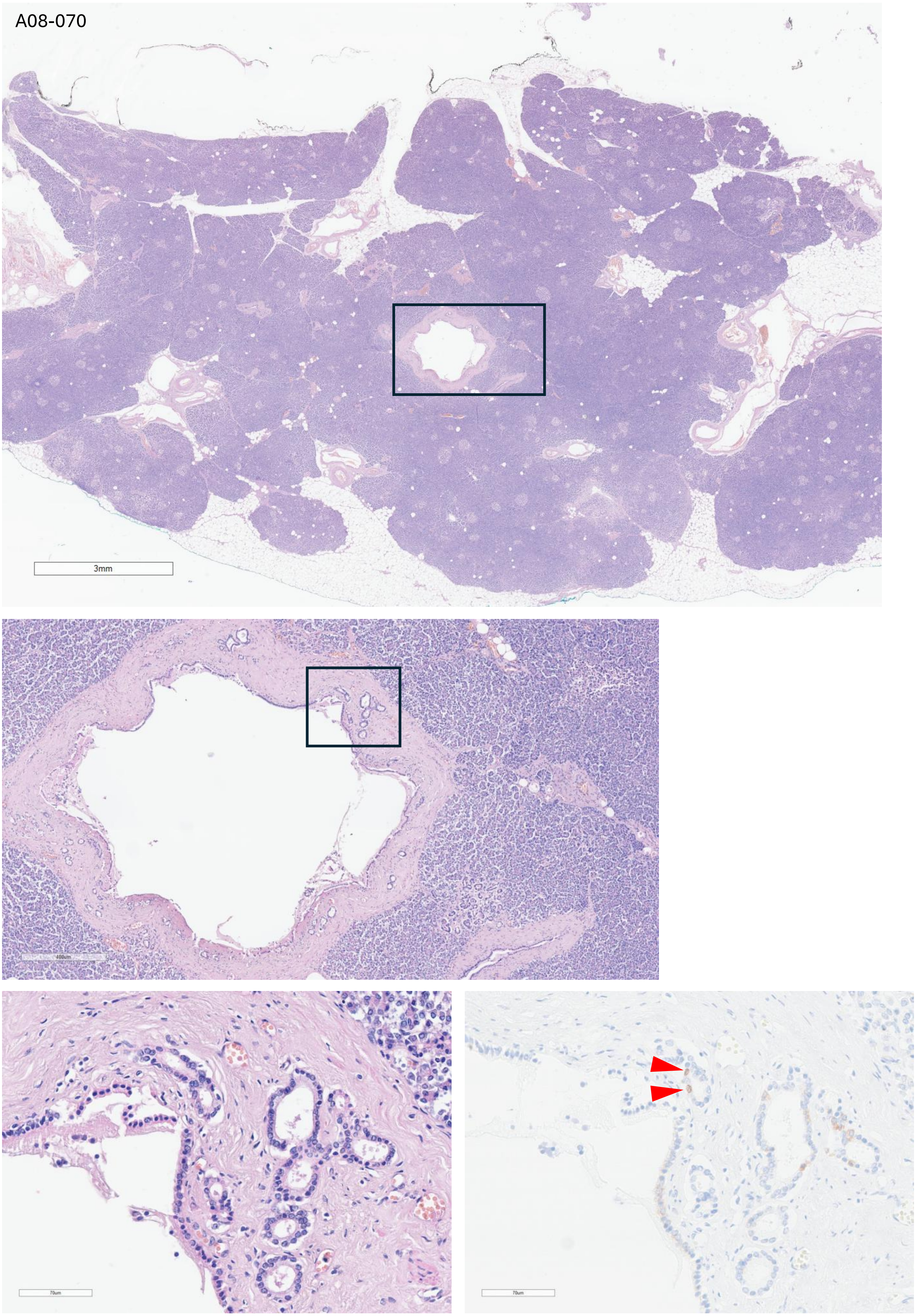
A08-070
3mm
70um
70um

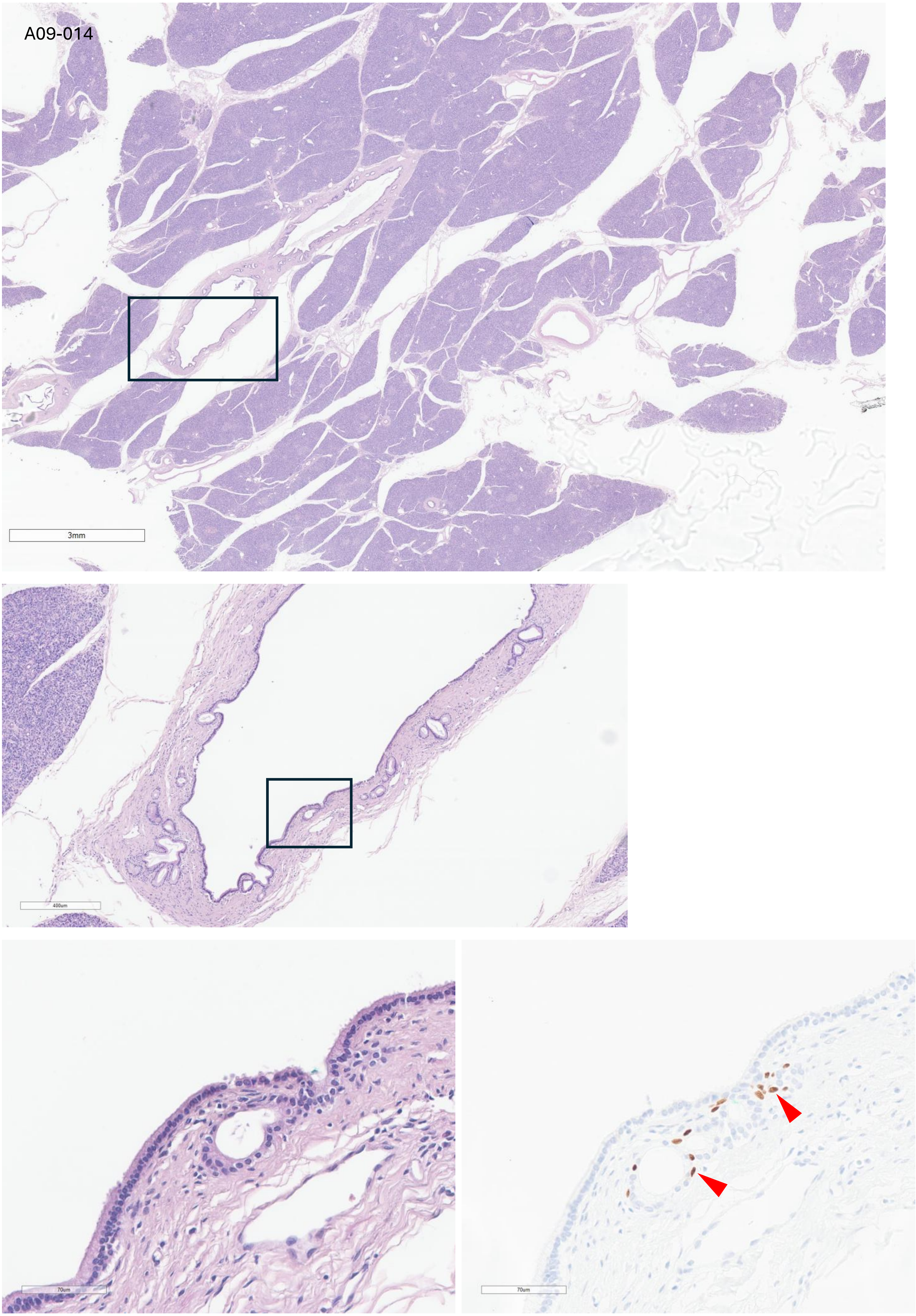
A09-014
3mm
400um
70um
70um

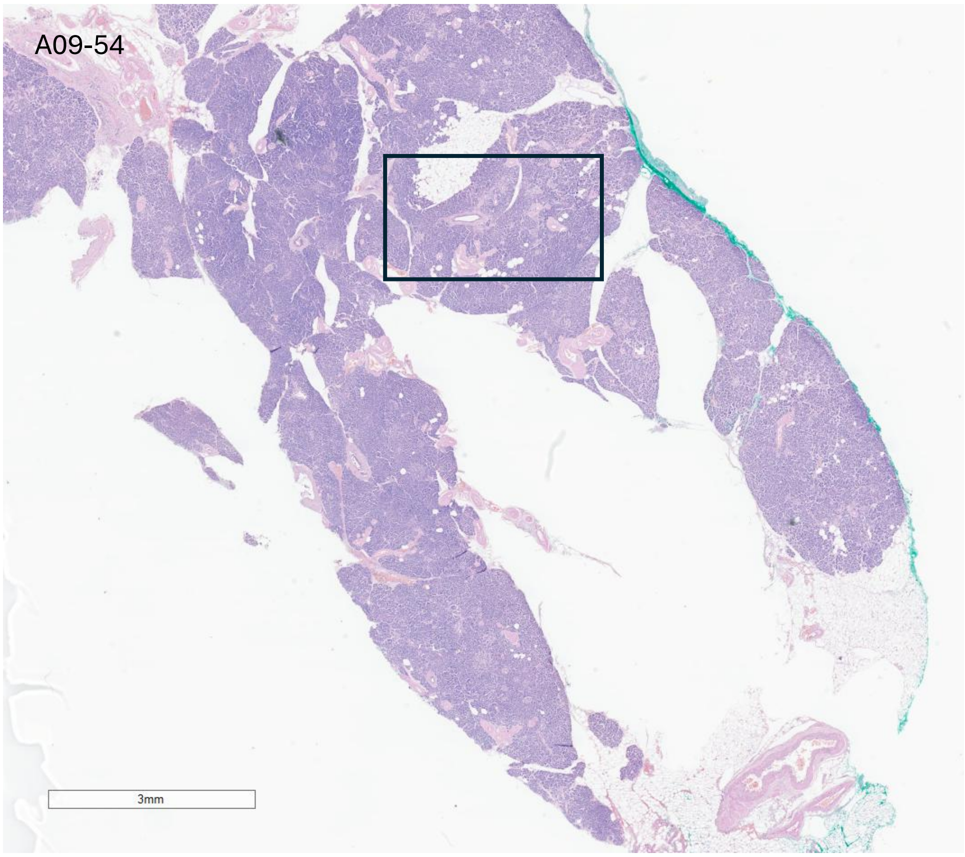
A09-54
3mm

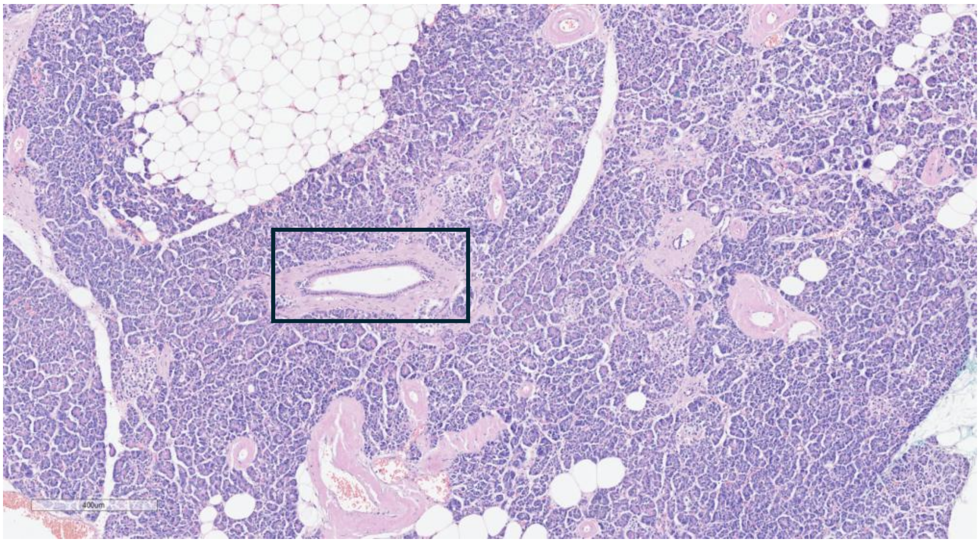
400um

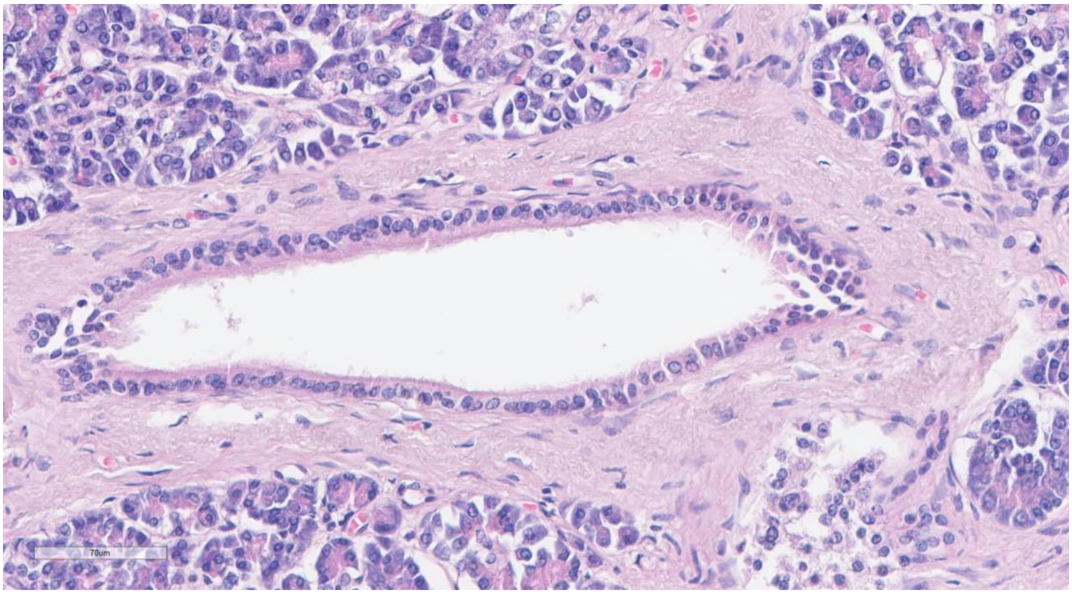
70um

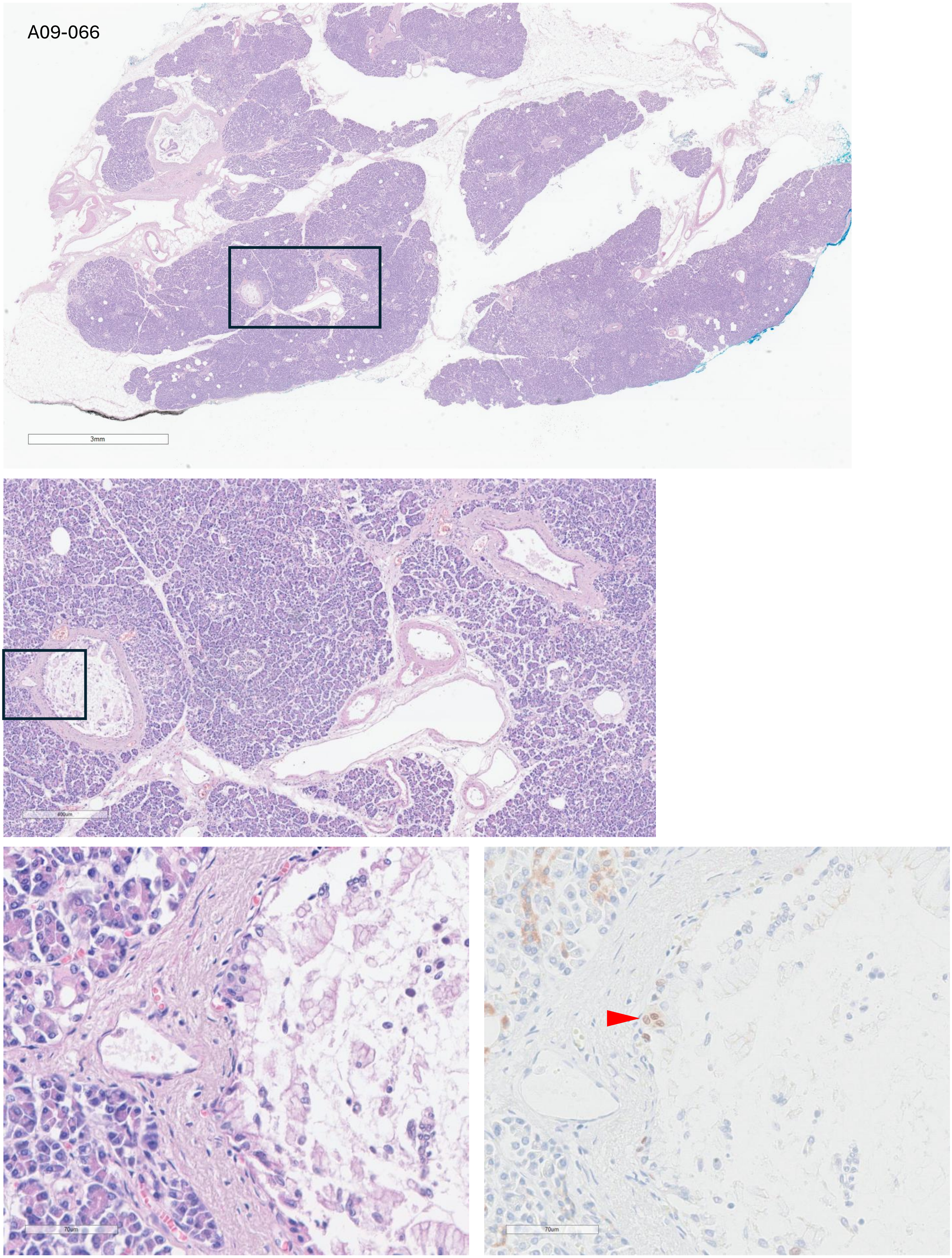

A09-066
3mm
400um
70um
70um

A404

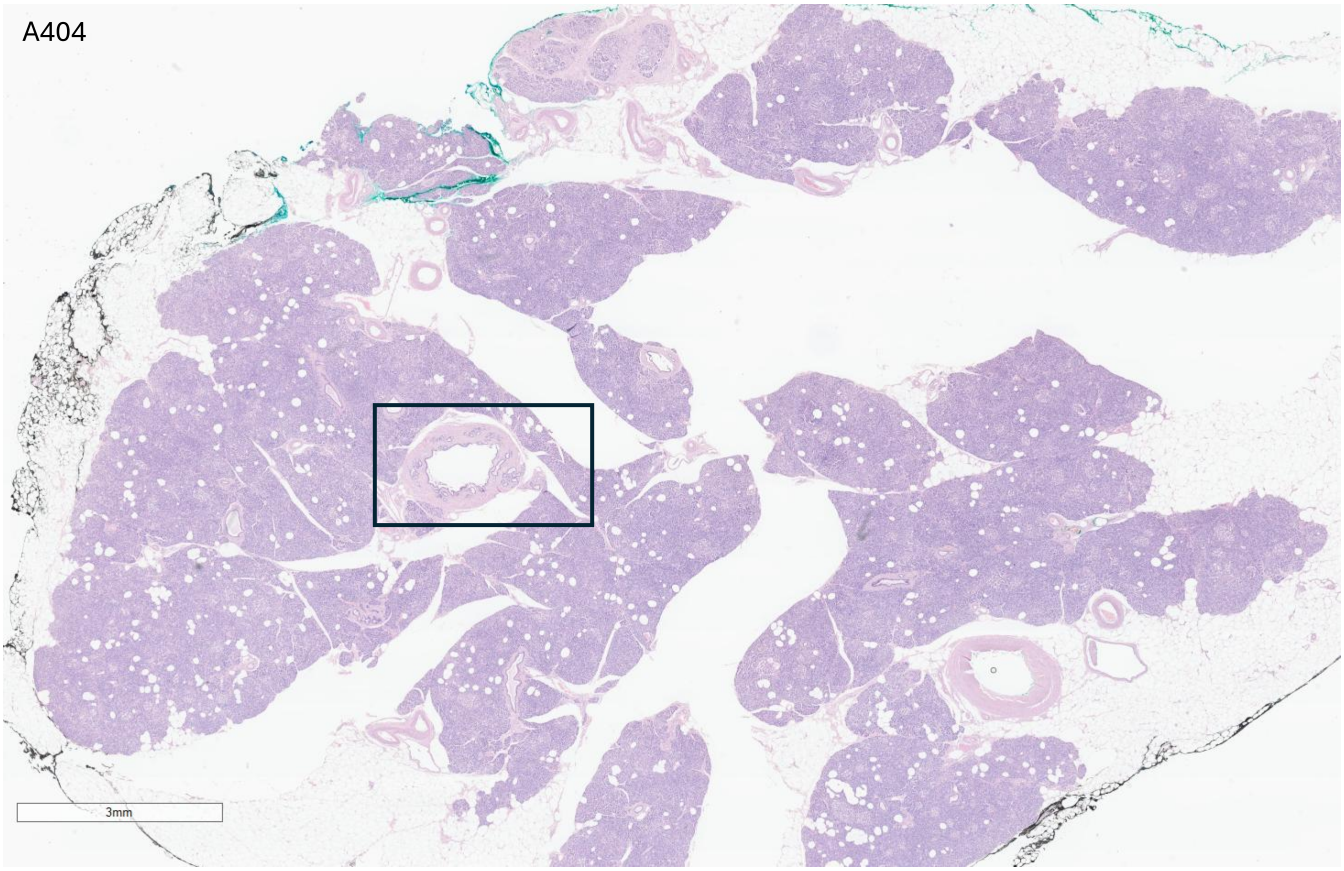


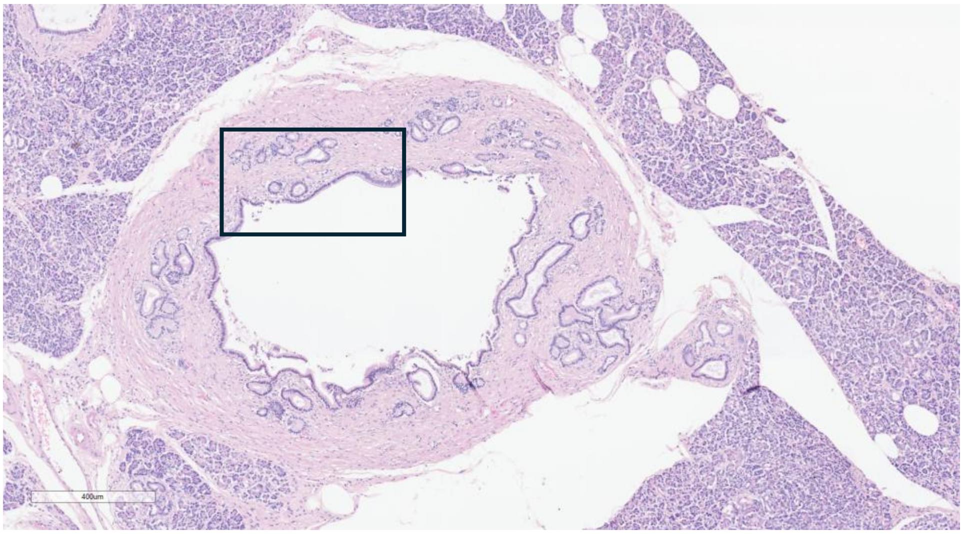


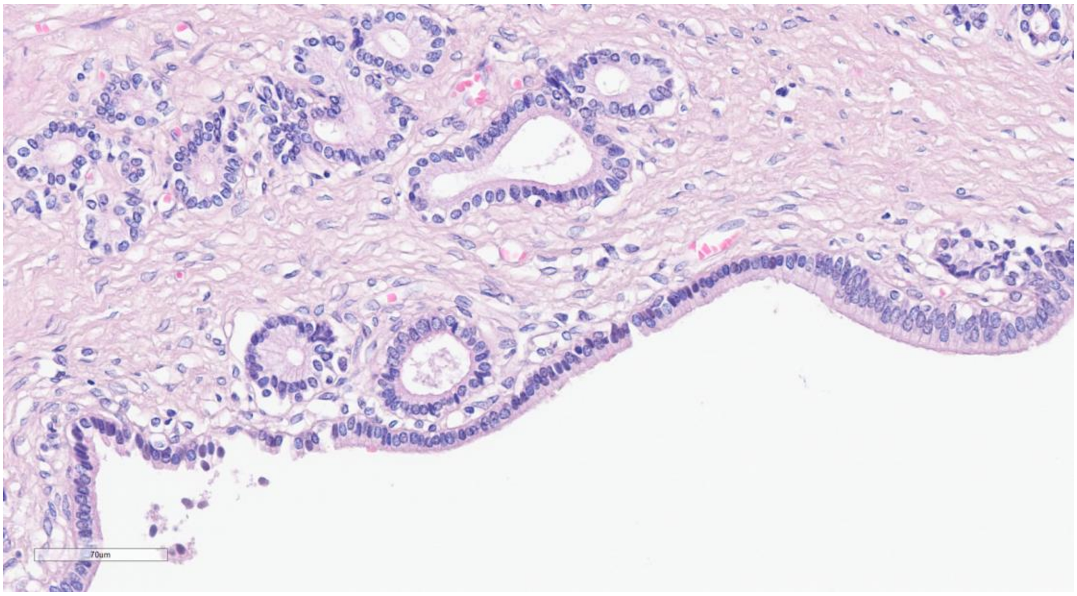

### Non-neoplastic pancreas

(Normal pancreas – NP)

Quantification cohort

NP-1

Head

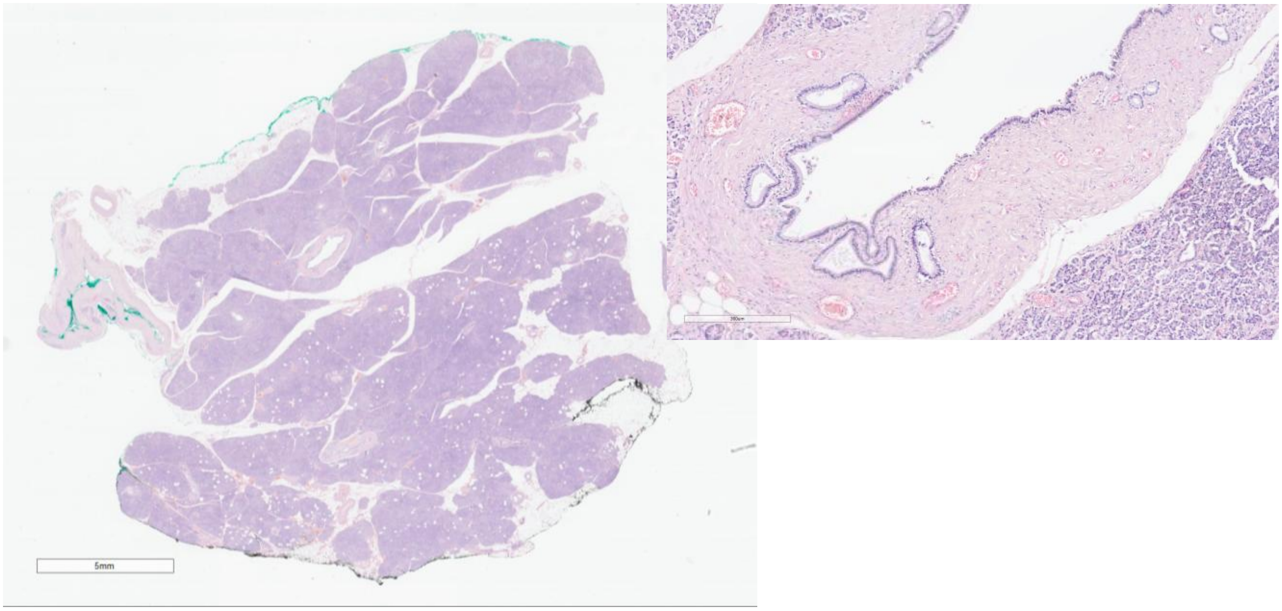


Body

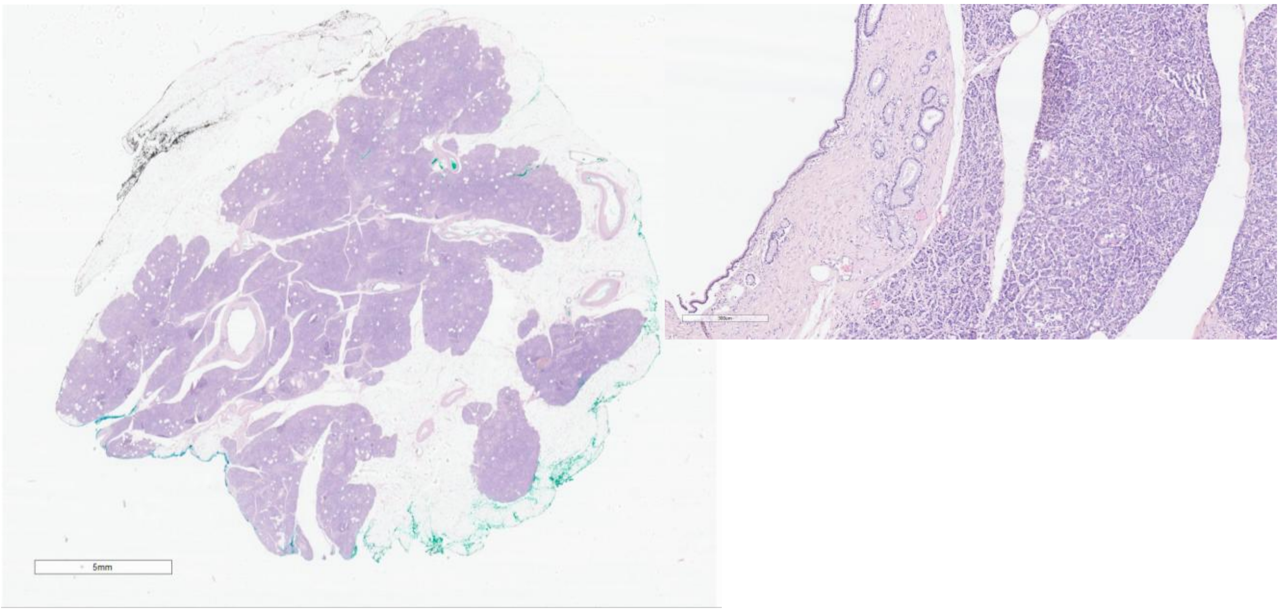


Tail

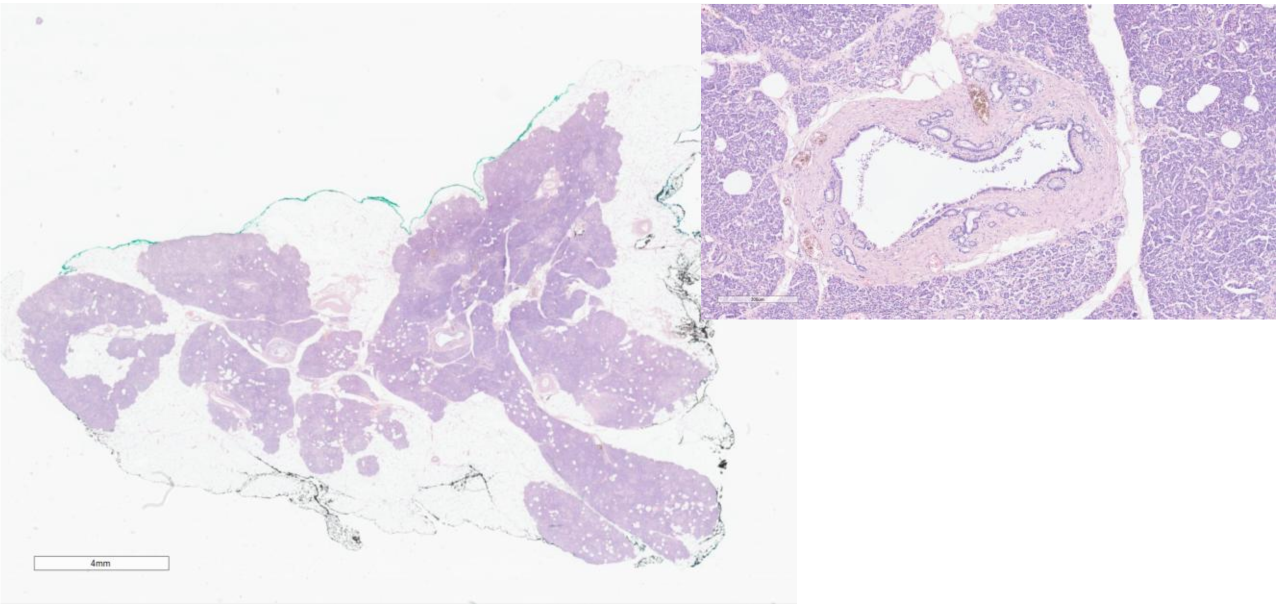

NP-2

Head

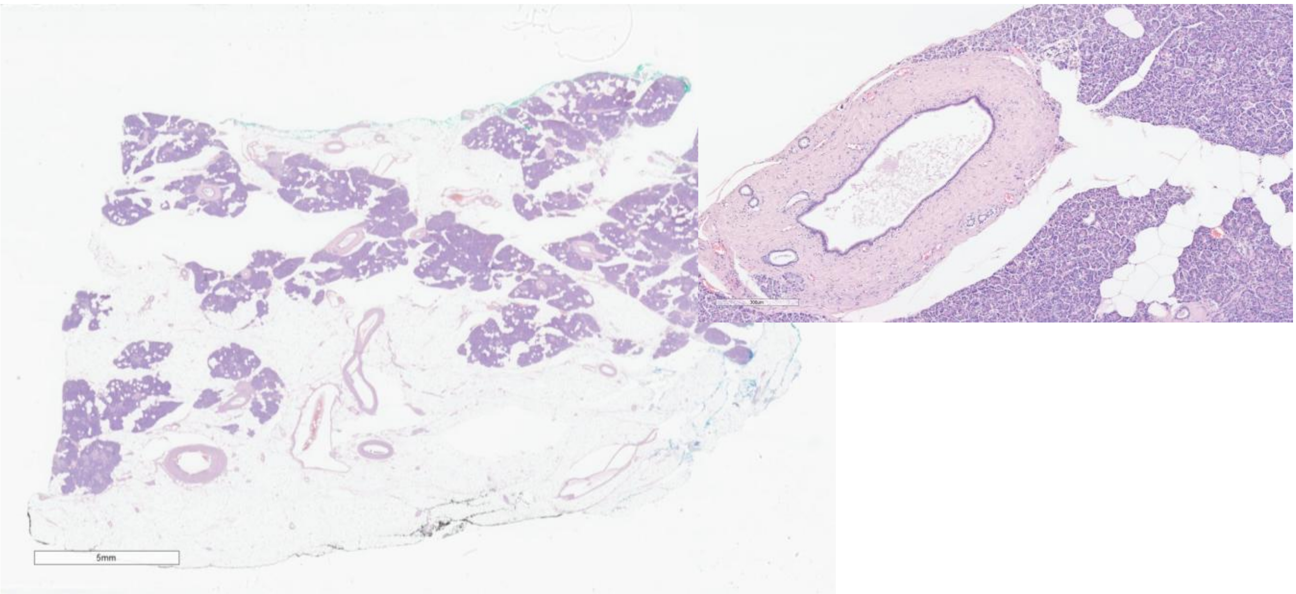


Body

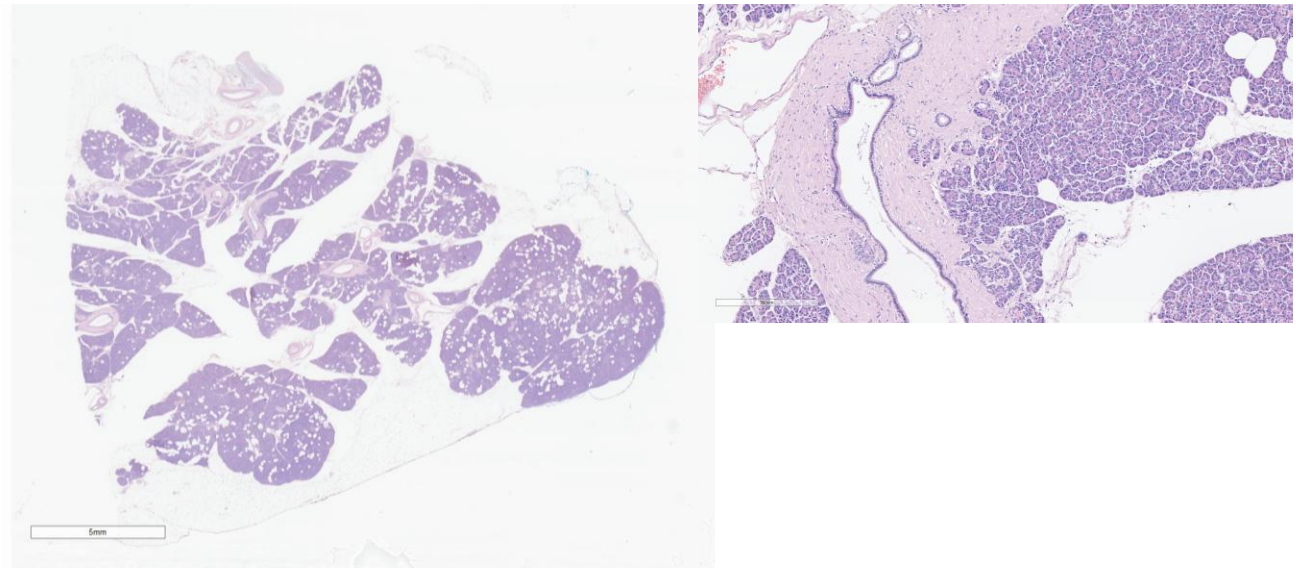


Tail

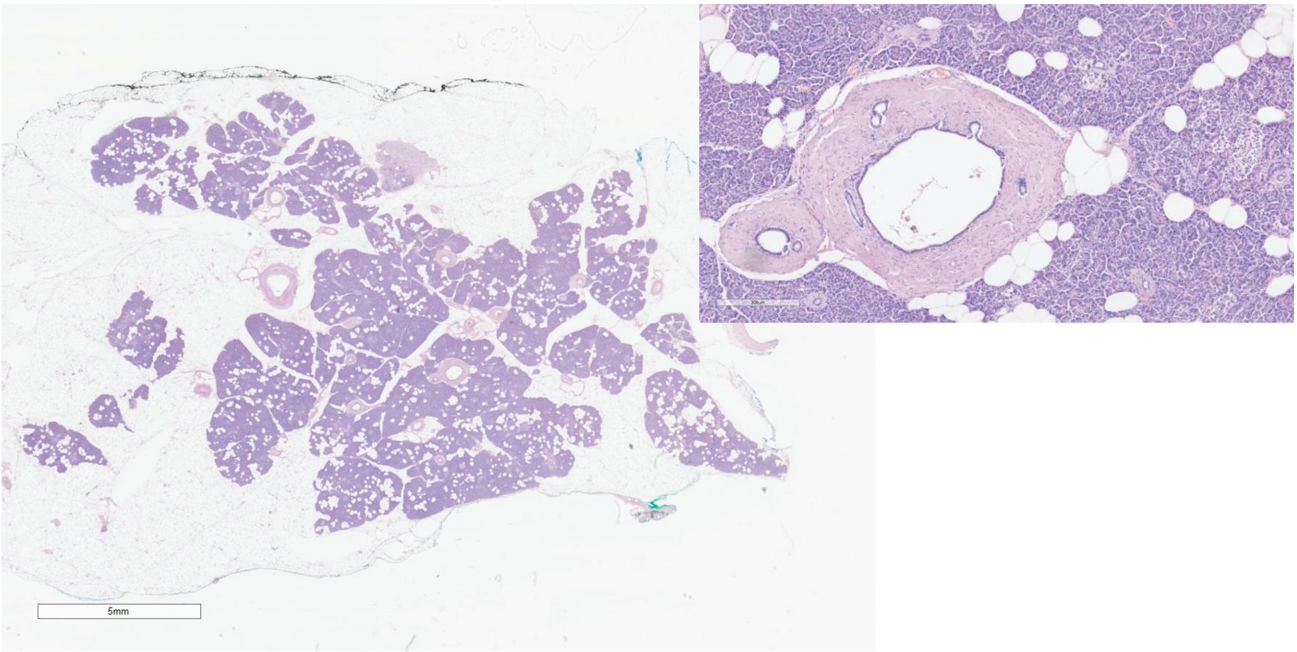

NP-3

Head

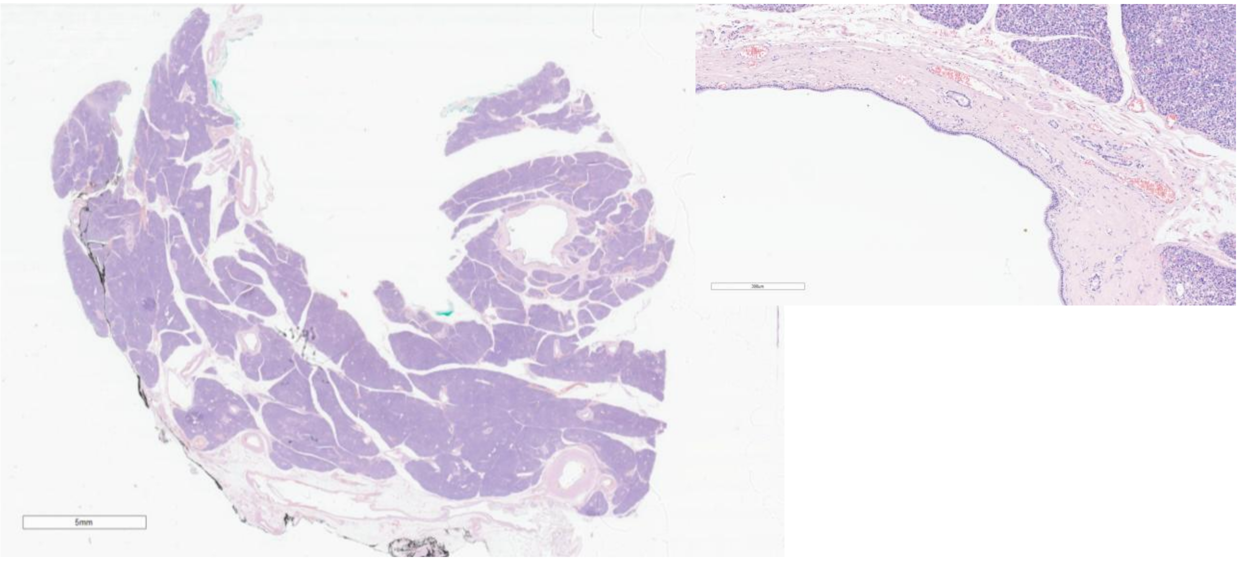


Body

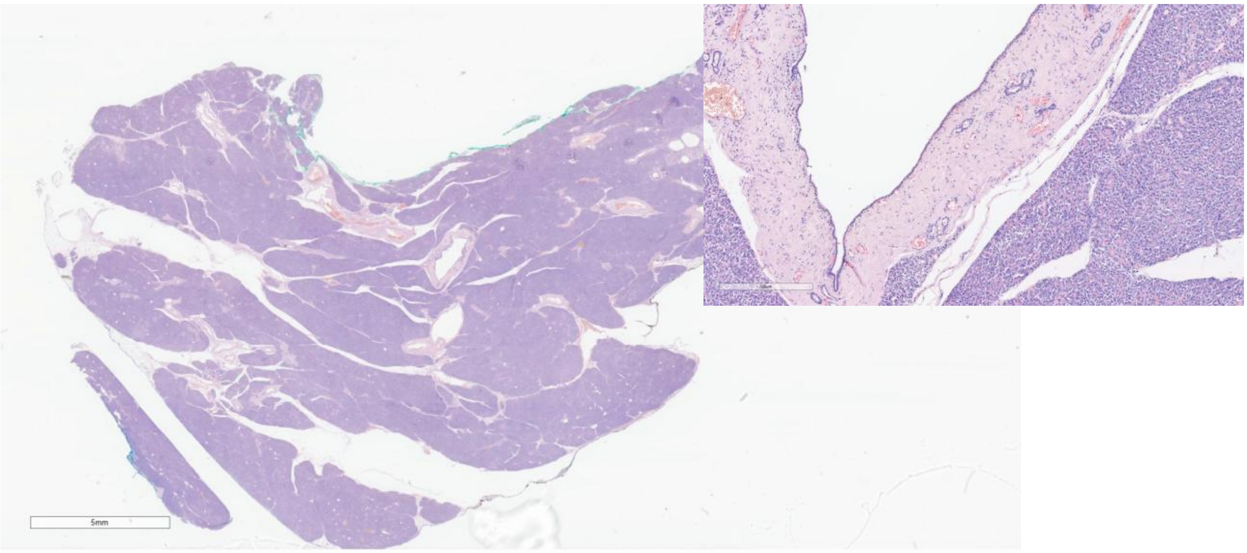


Tail

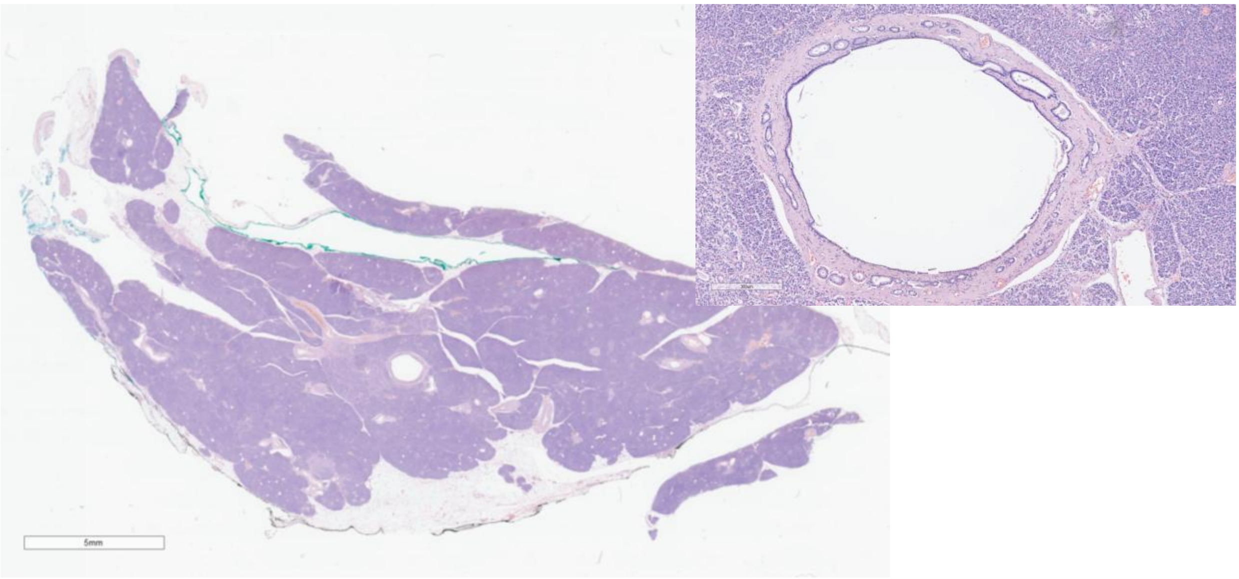

NP-4

Head

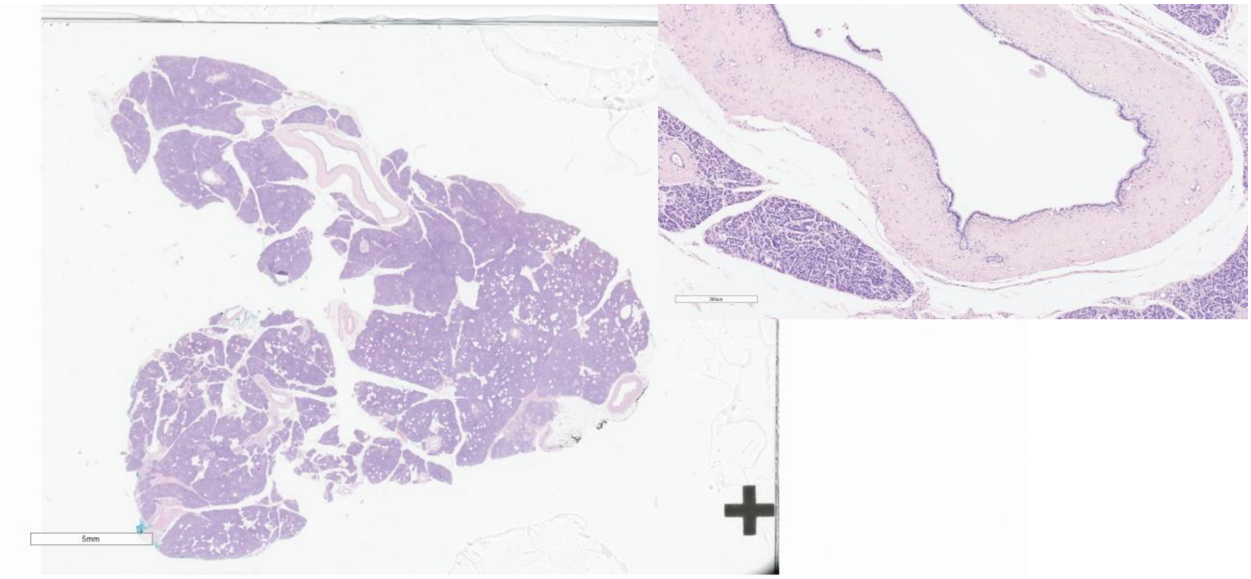


Body

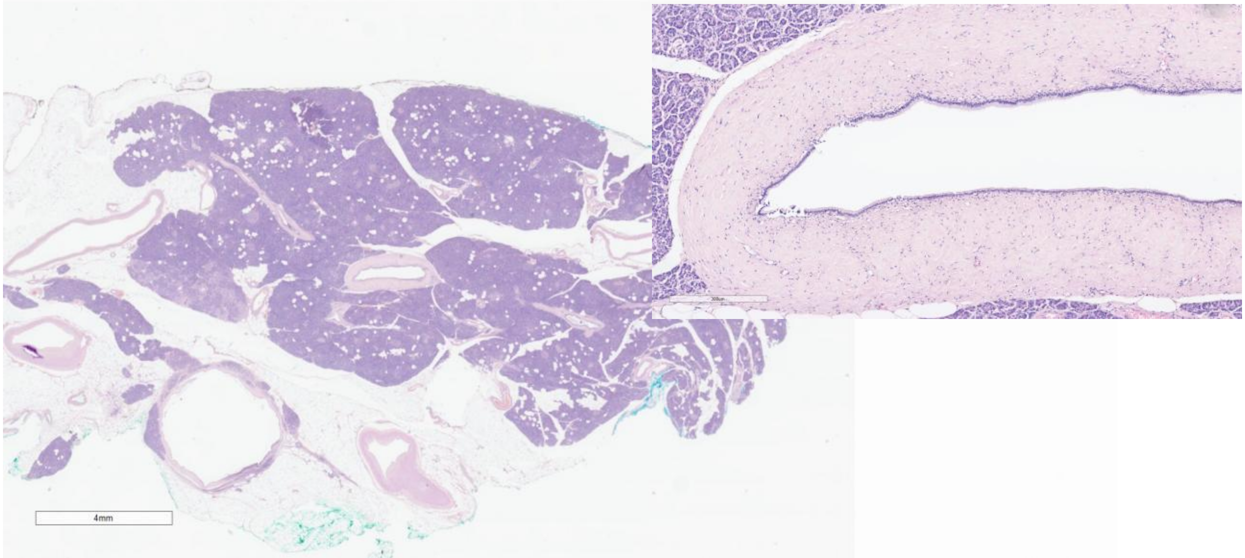


Tail

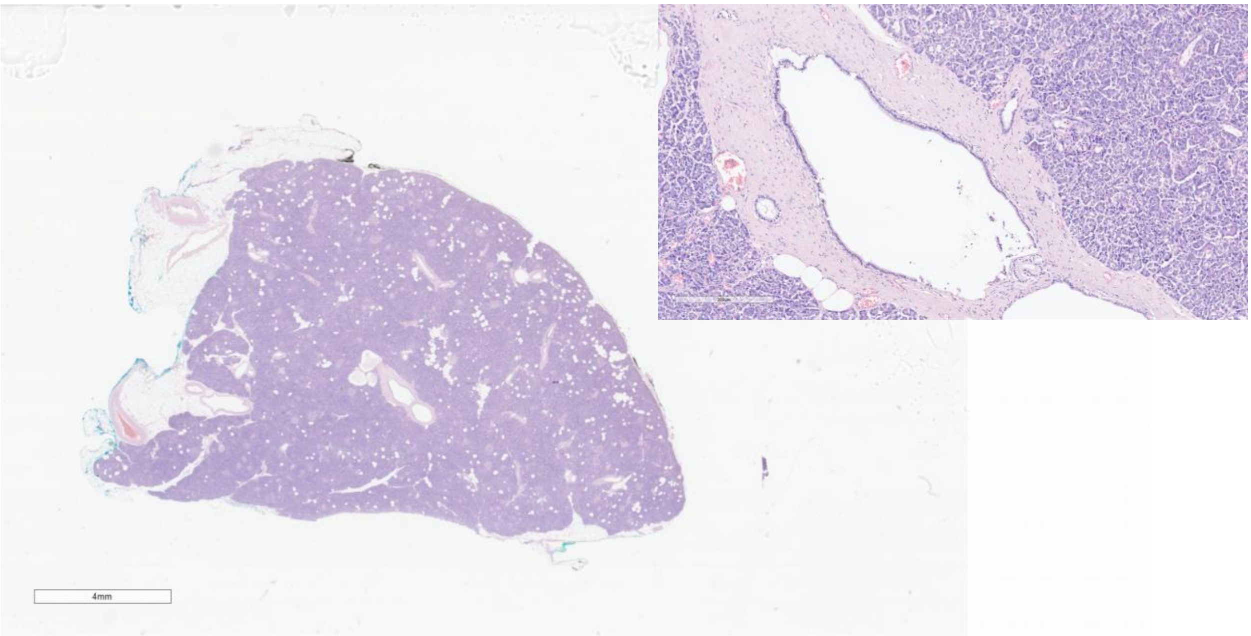

NP-5

Head

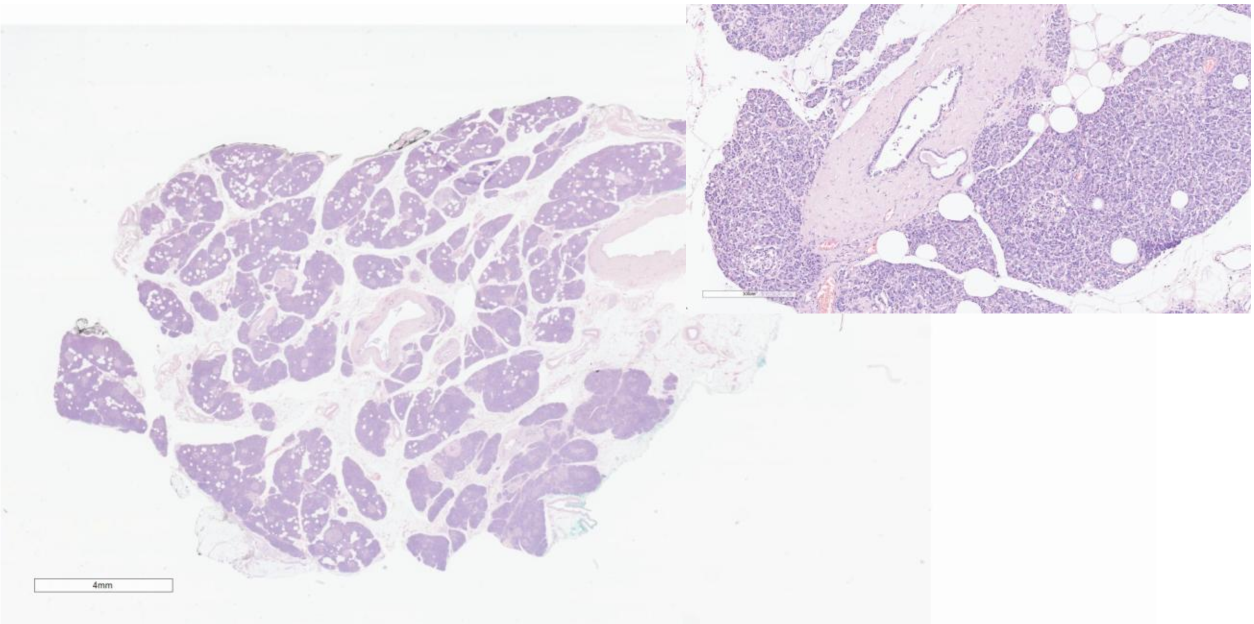


Body

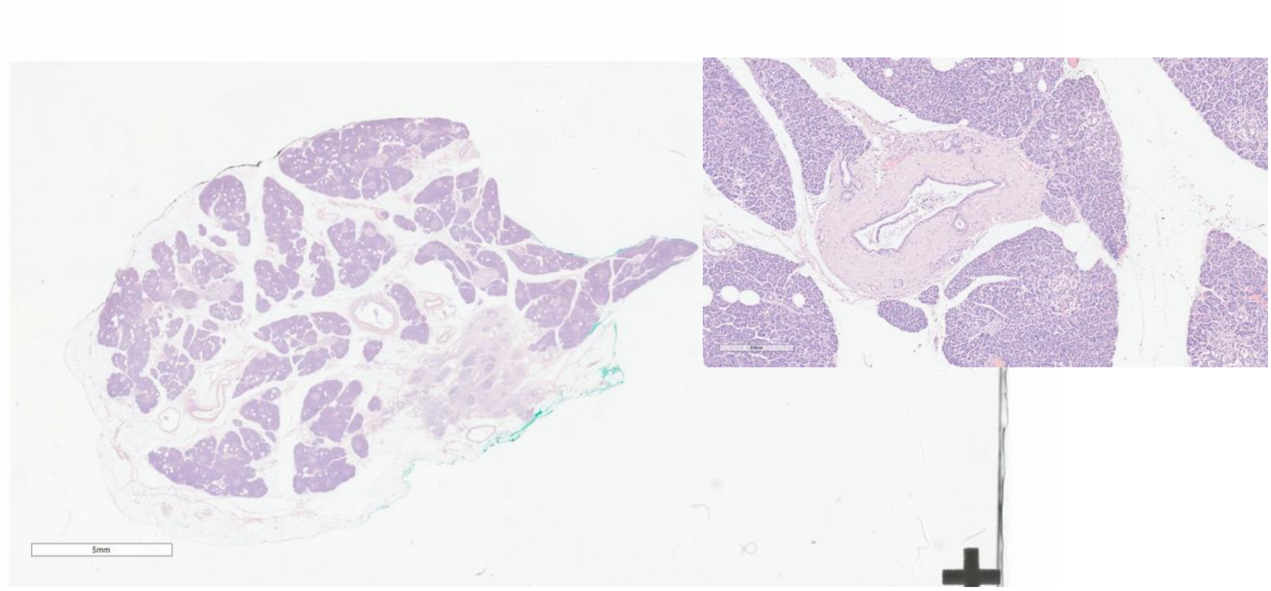


Tail

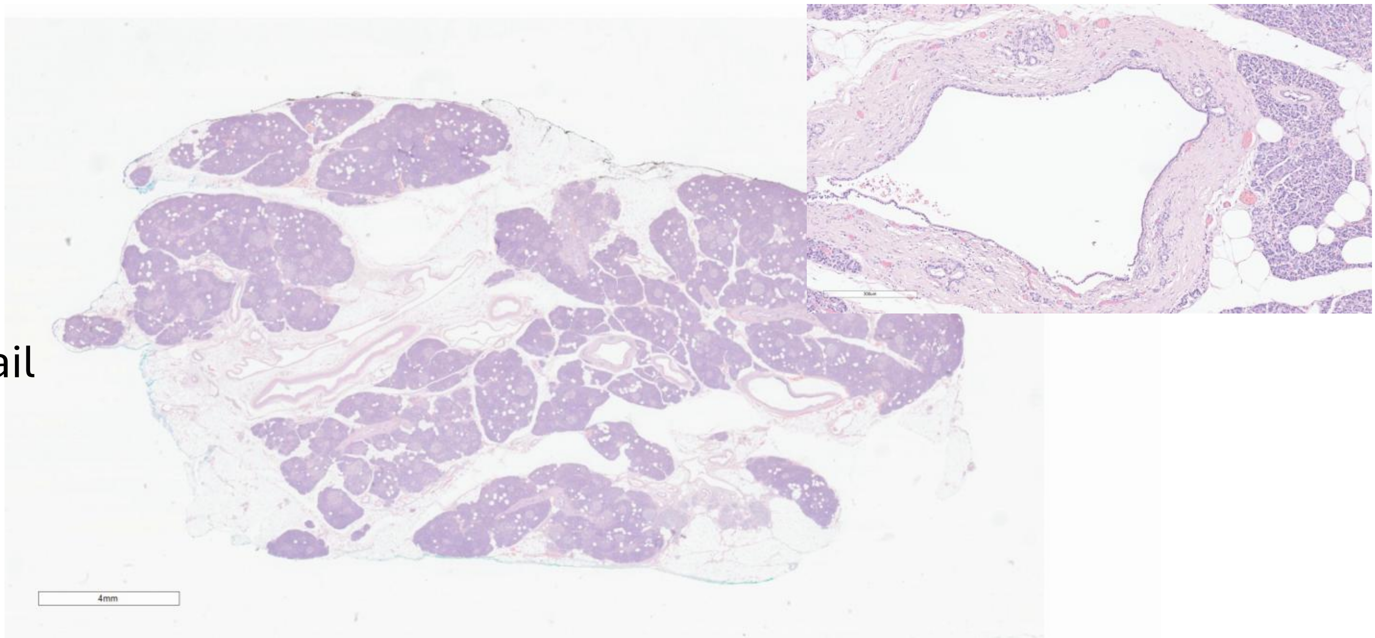

# NP-6

Head

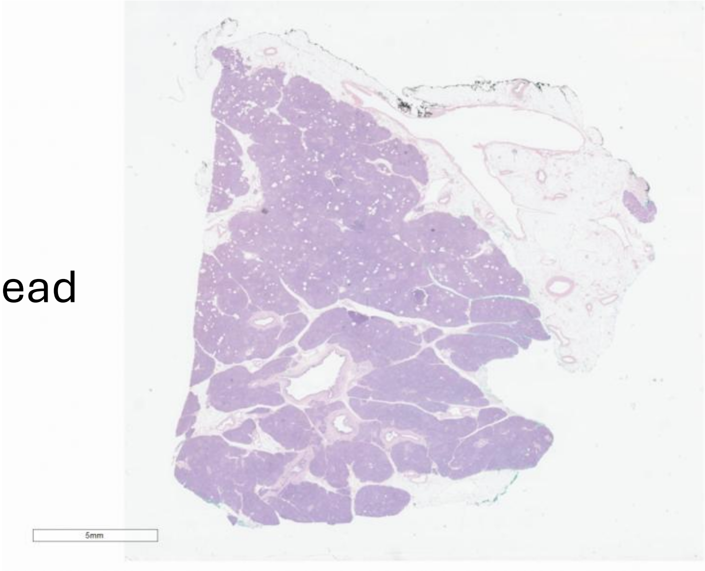


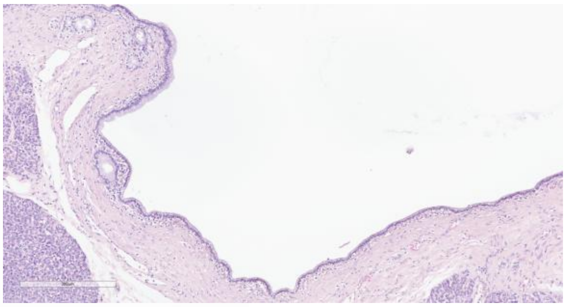

Body

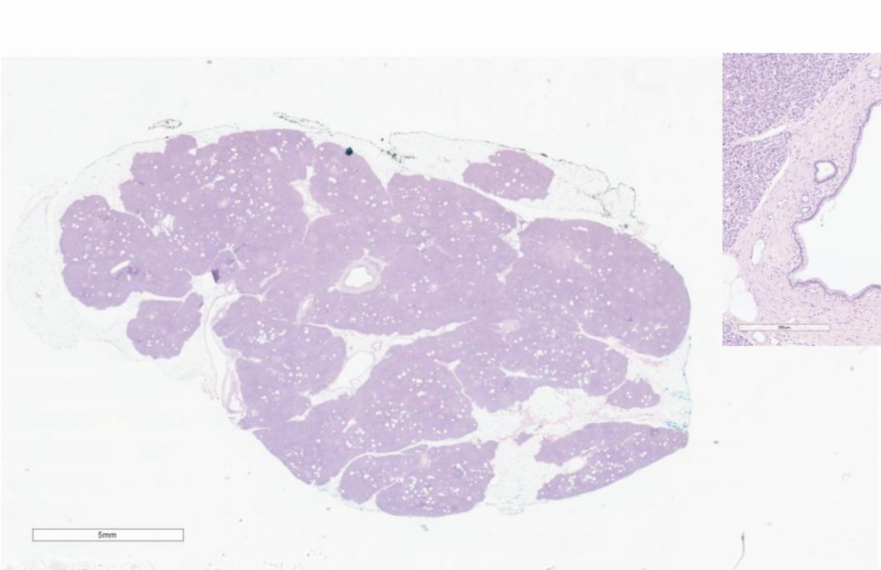


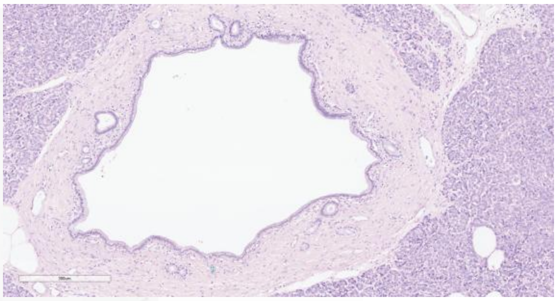

Tail

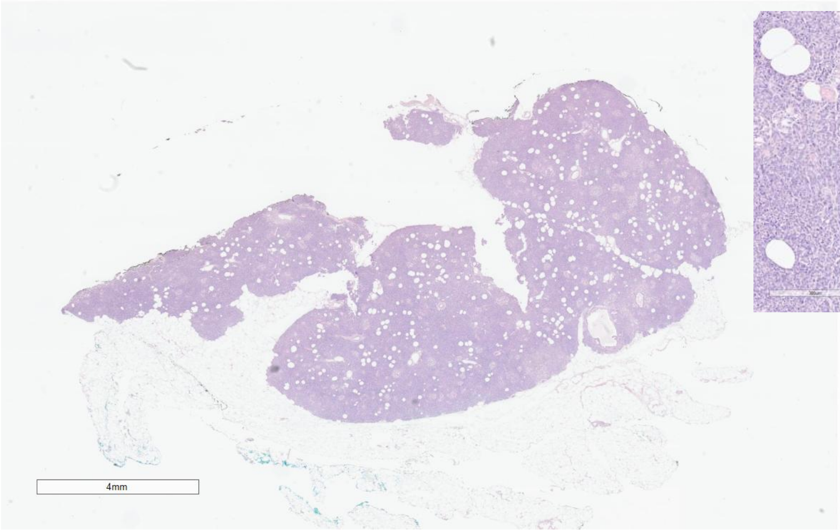


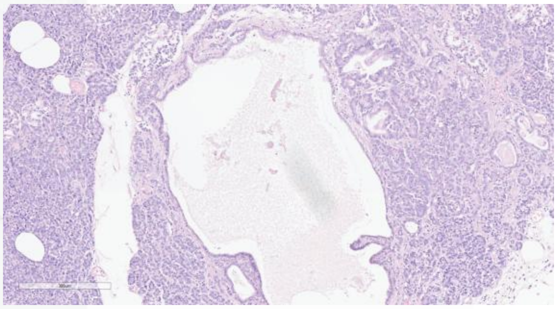

NP-7

Head

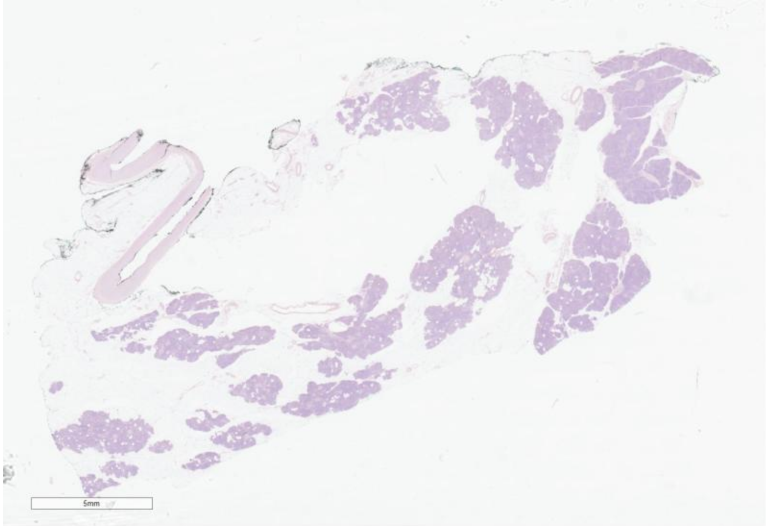


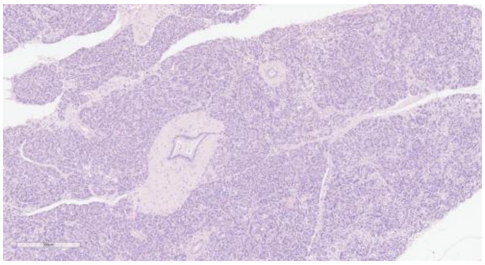

Body

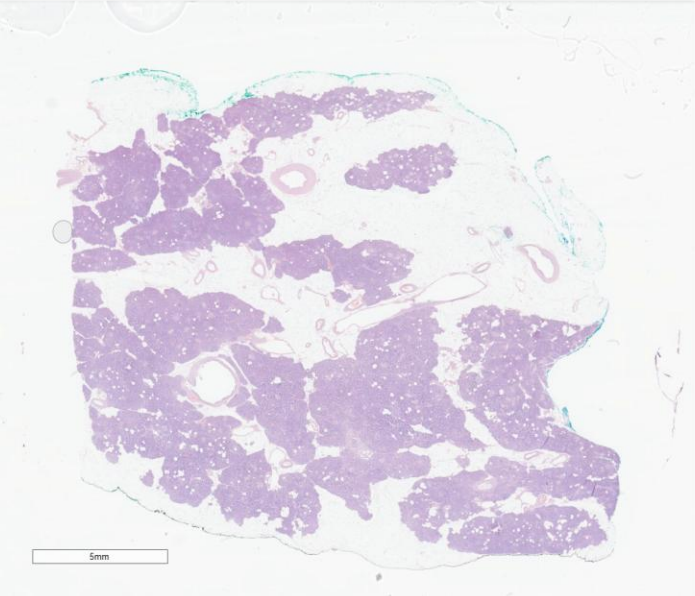


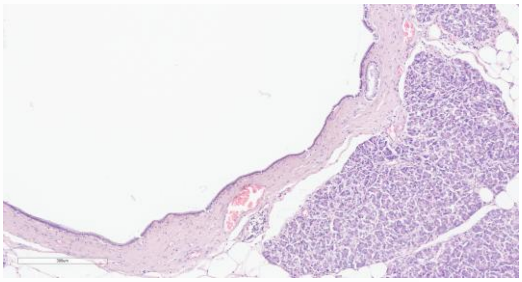

Tail

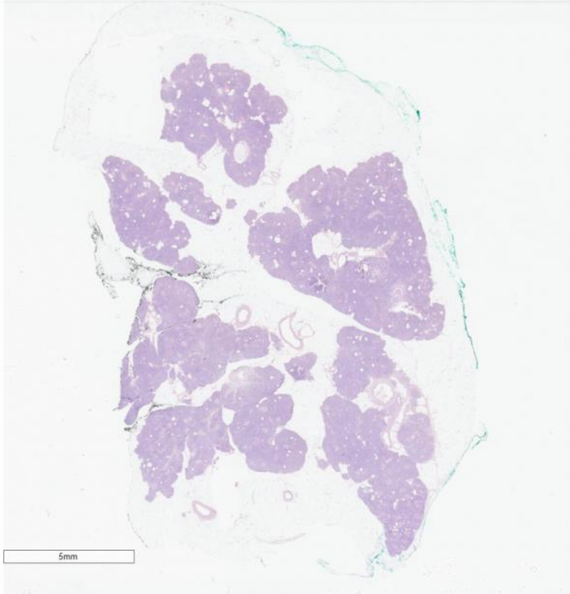


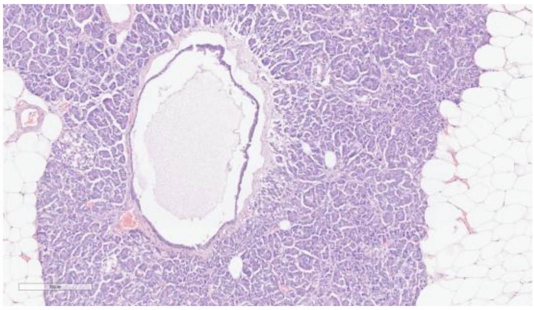

NP-8

Head

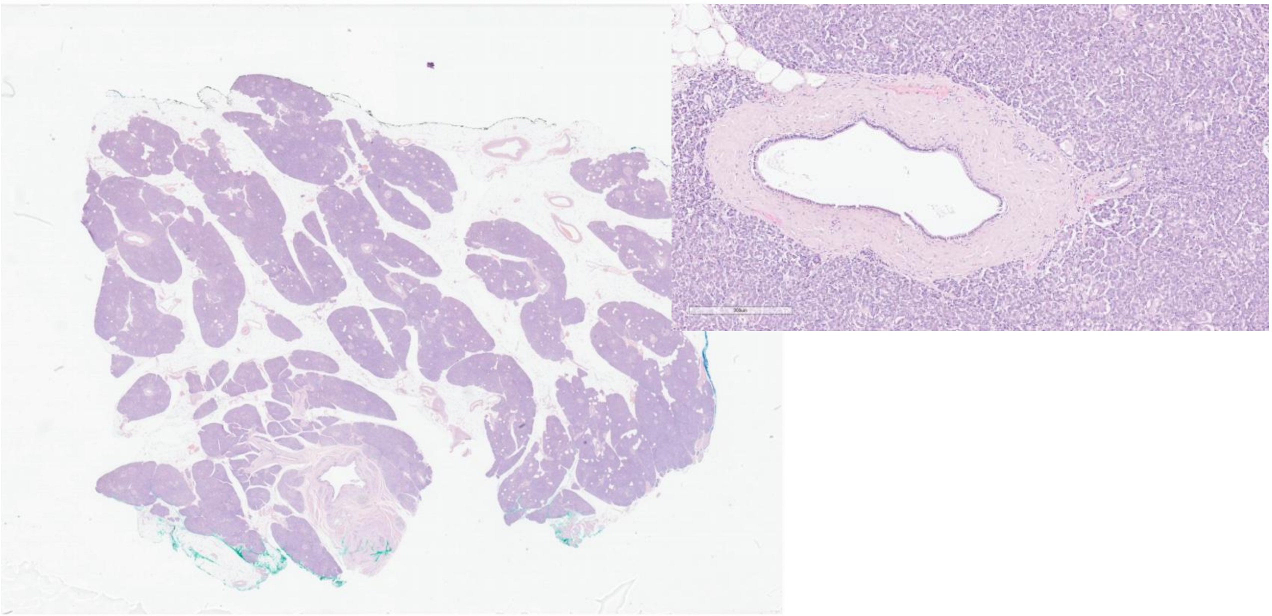

Body

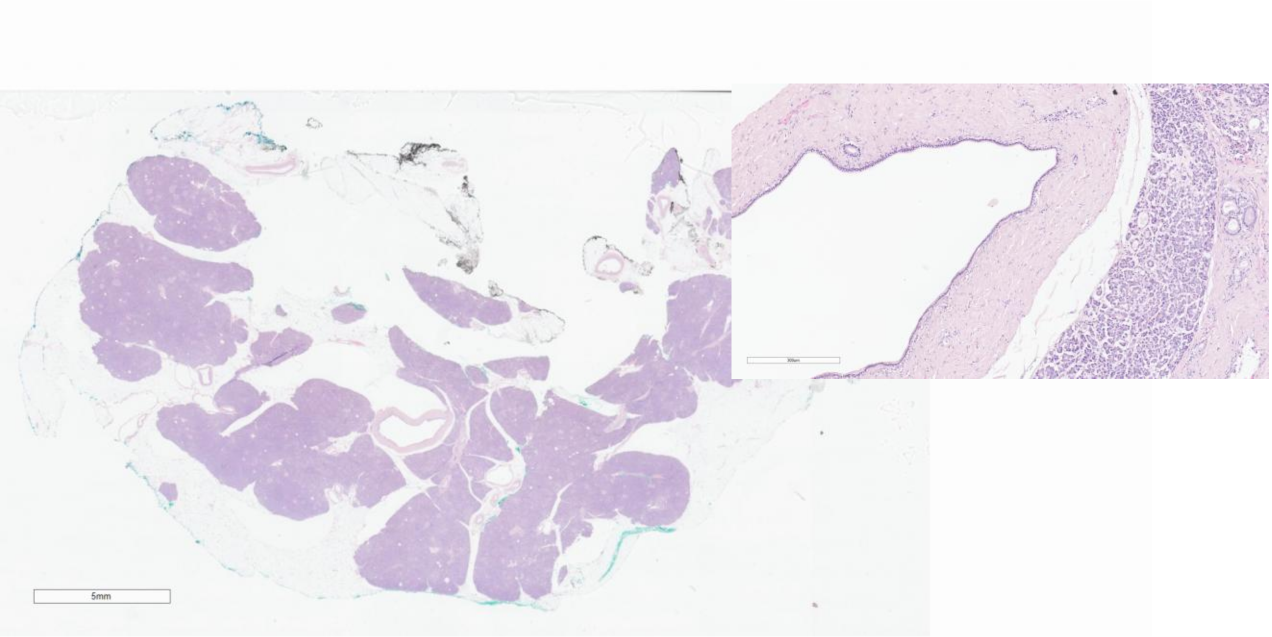


Tail

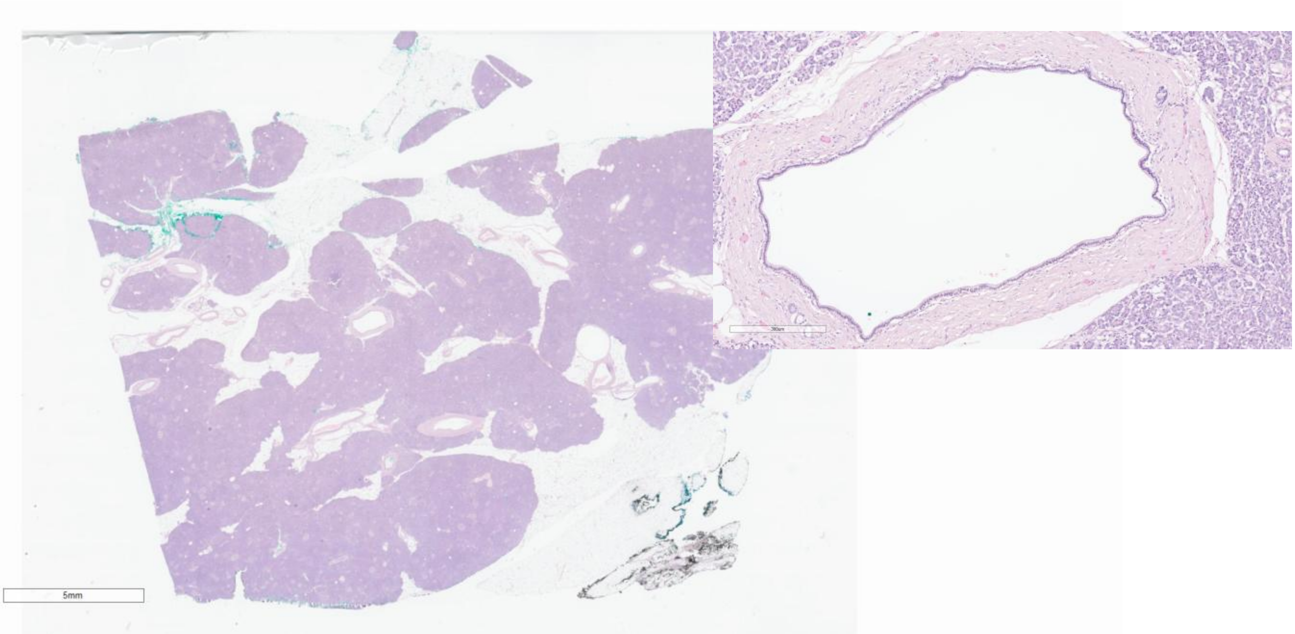

# NP-9

Head

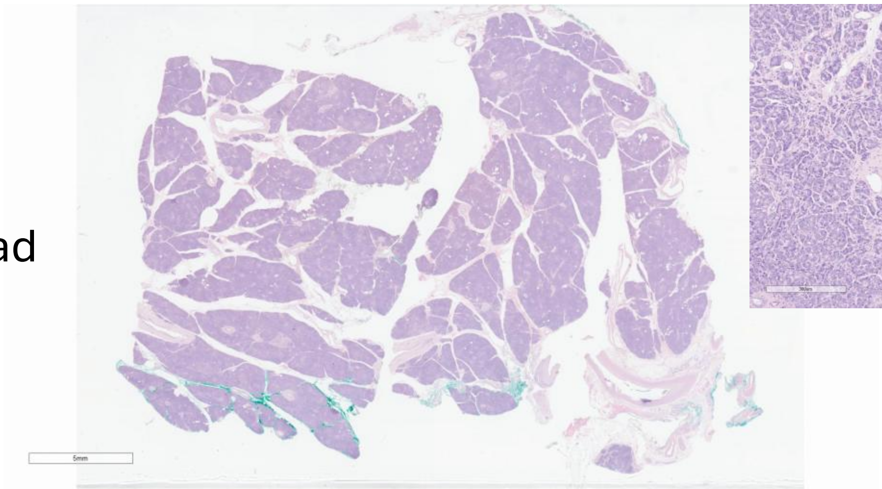


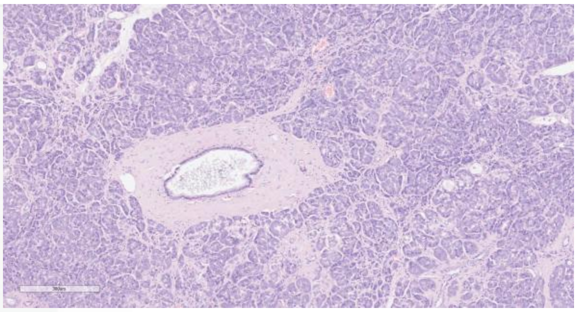

Body

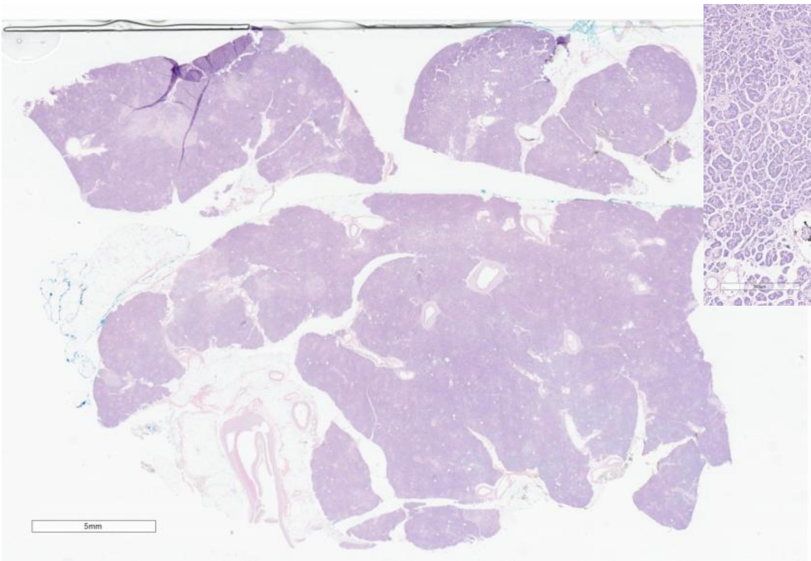


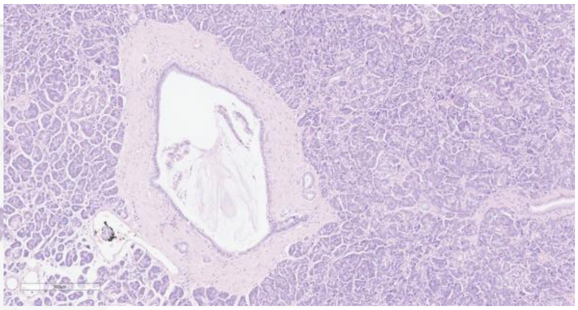

Tail

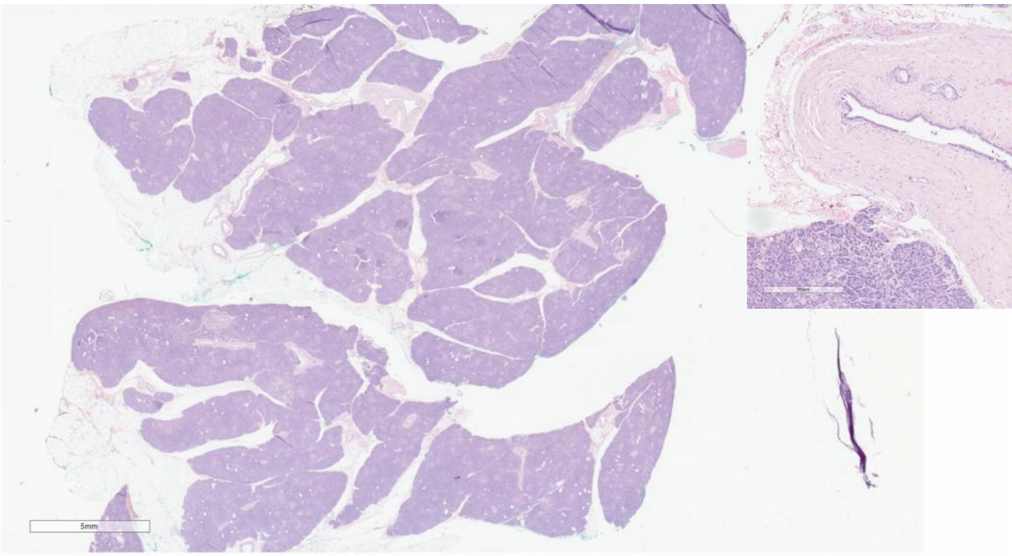


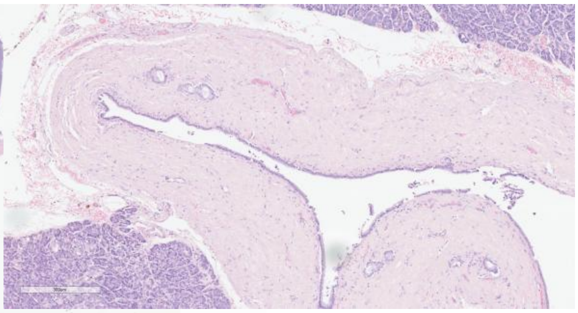

### Non-neoplastic pancreas

(Chronic pancreatitis – CP)

24EH7759E

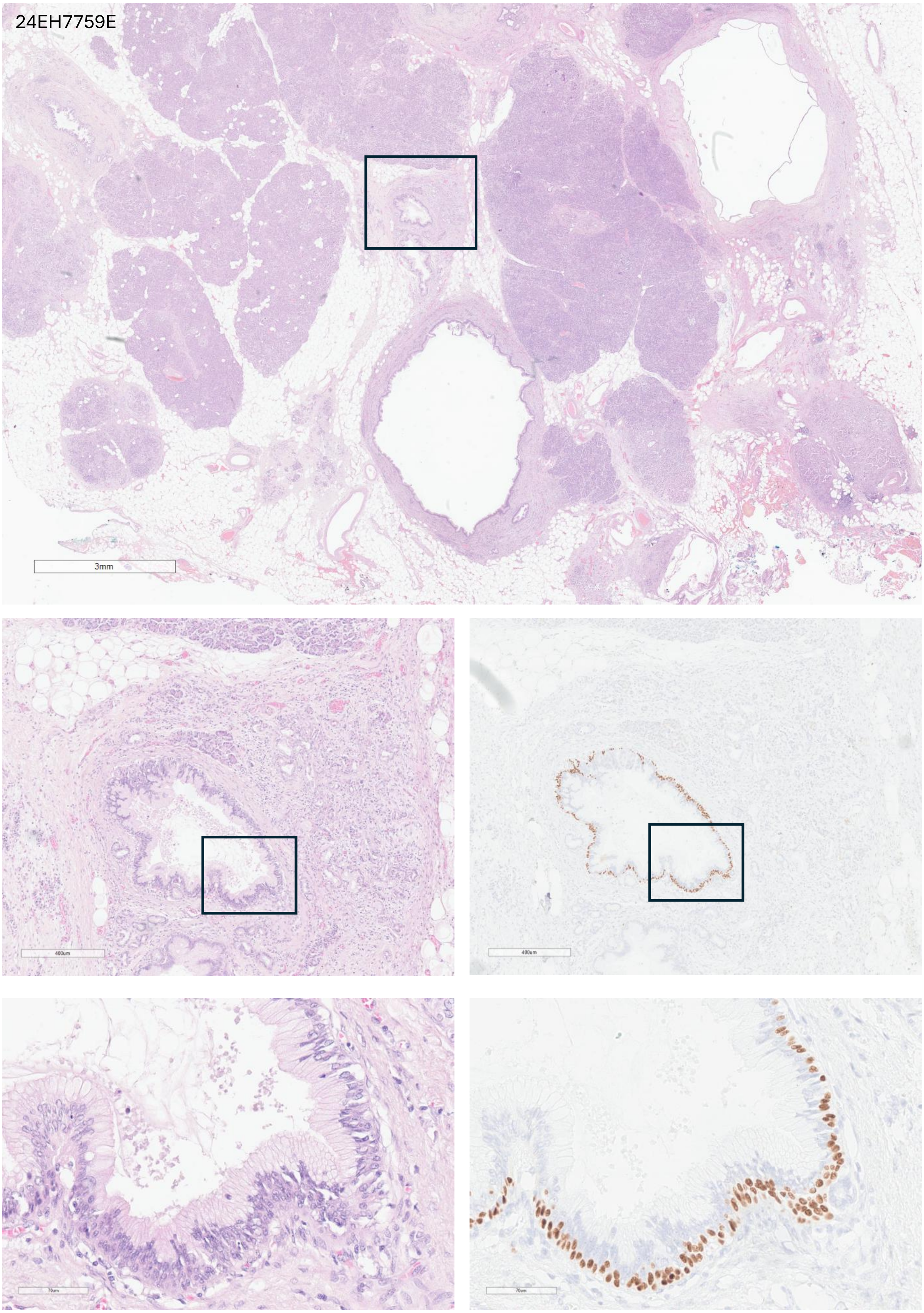

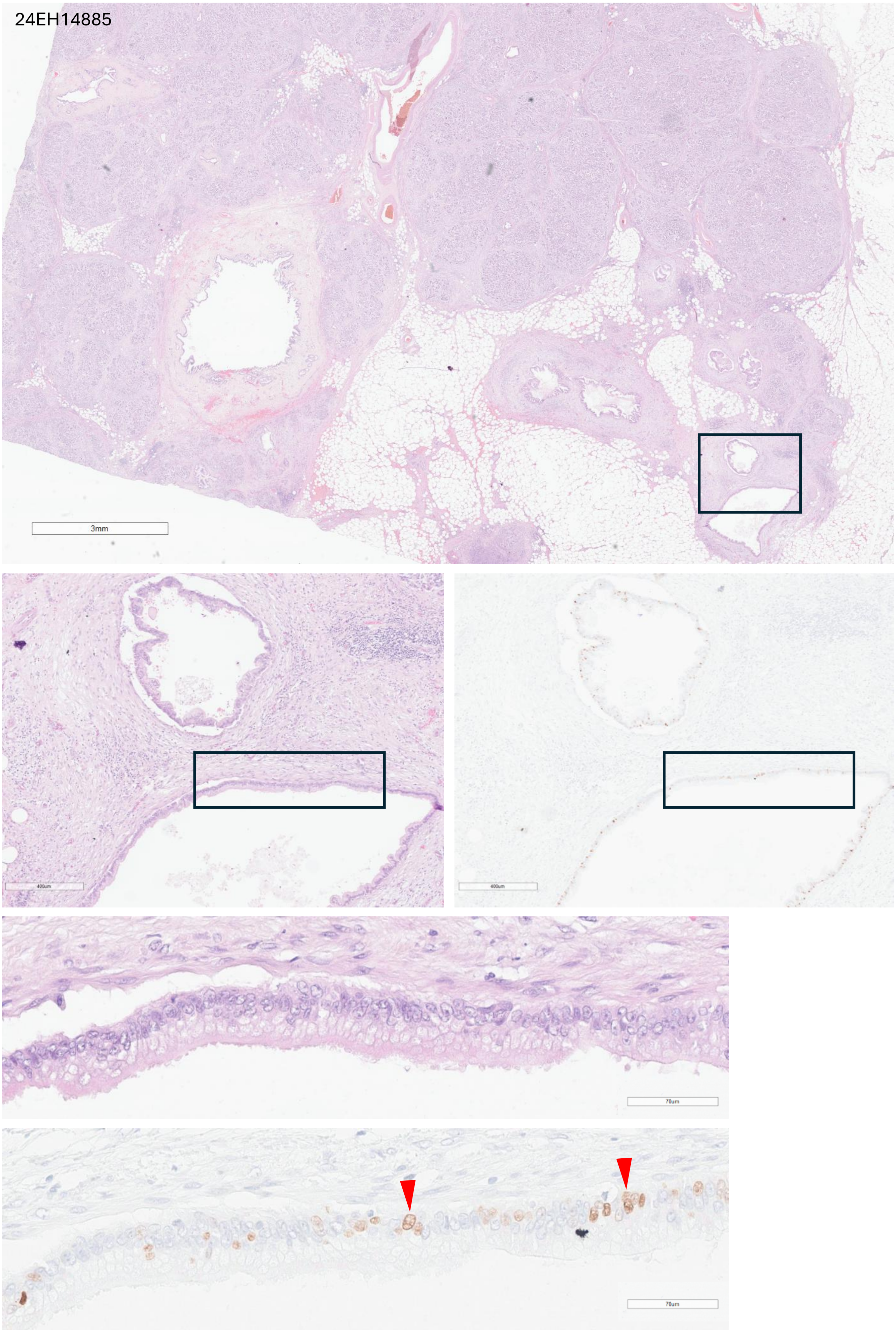
24EH14885
3mm
400um
400um
70um
70um

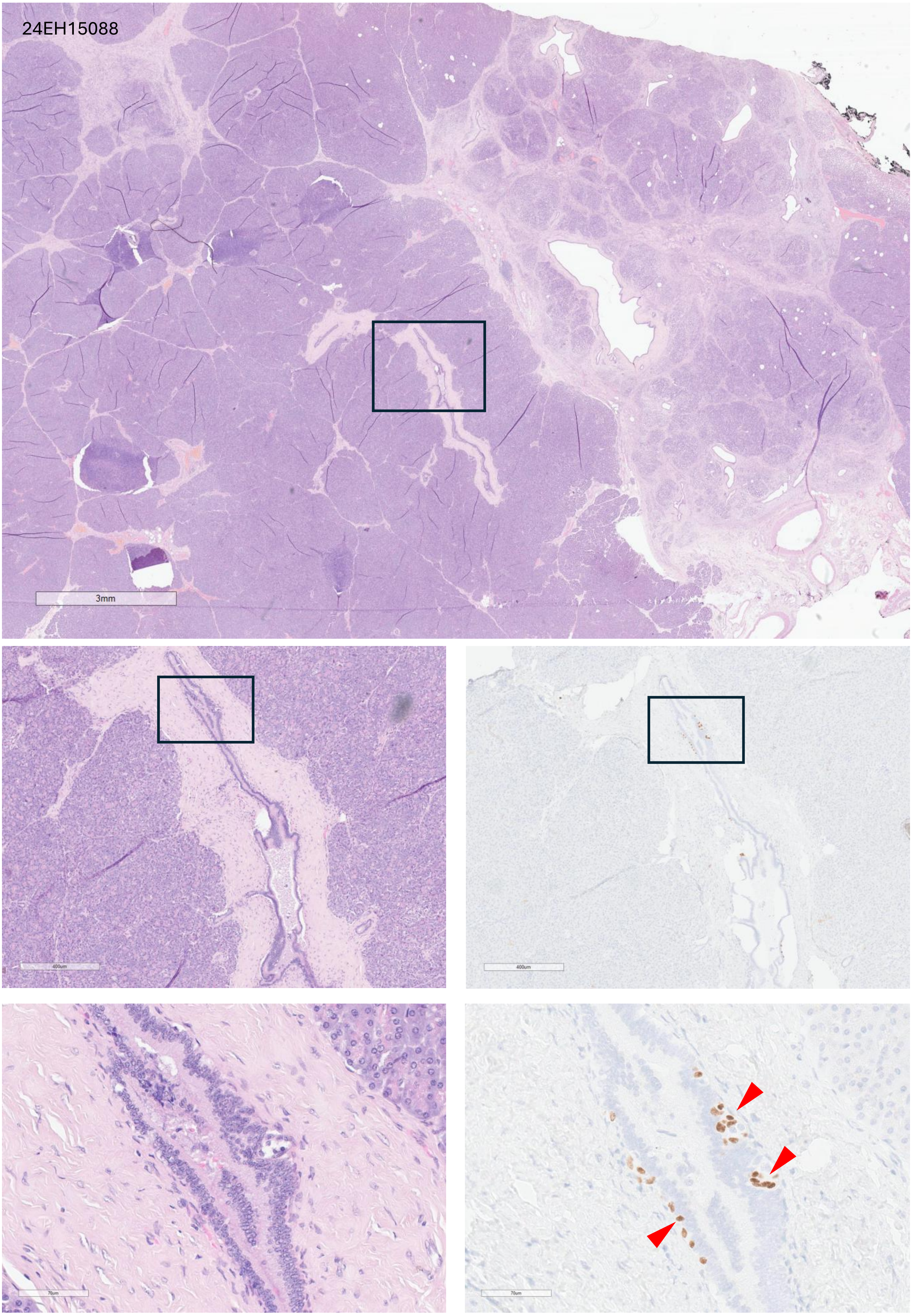
24EH15088
3mm
400um
400um
70um
70um

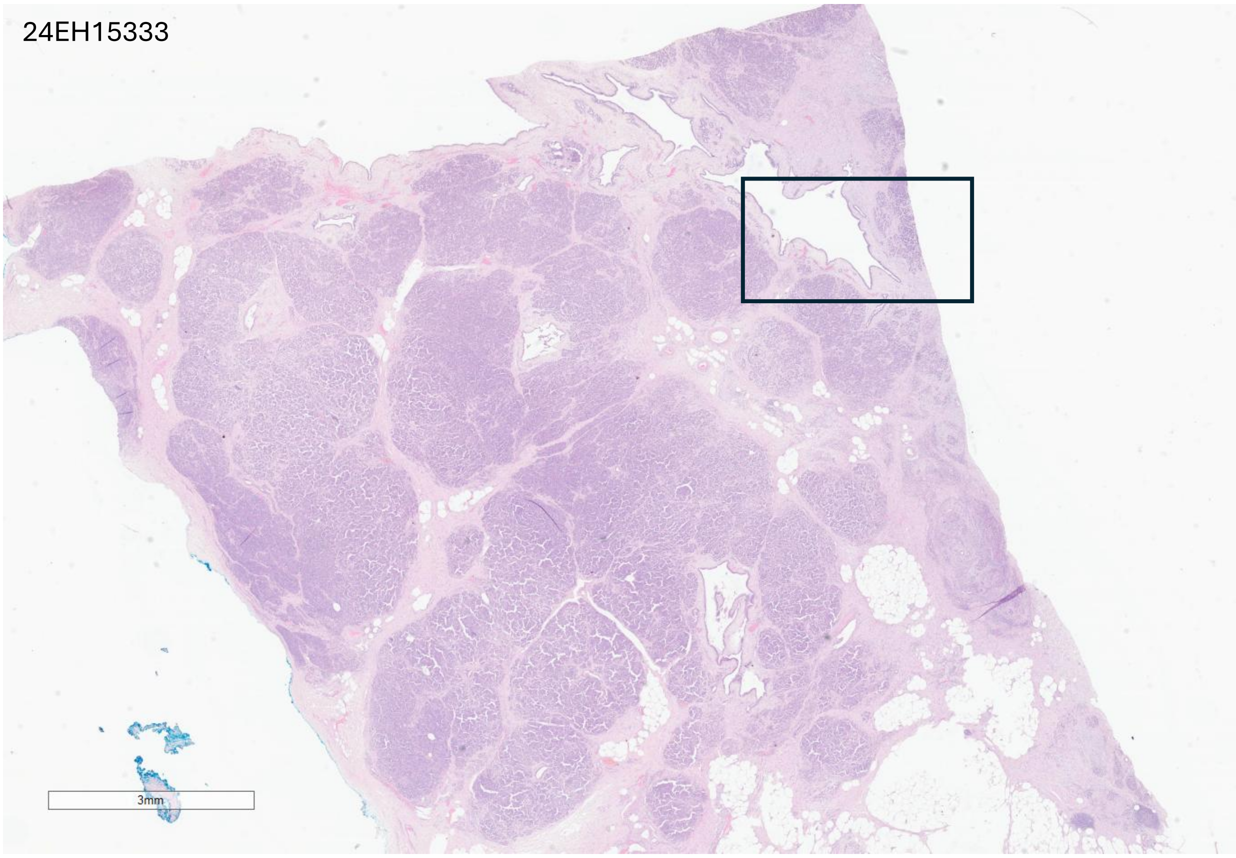
24EH15333
3mm

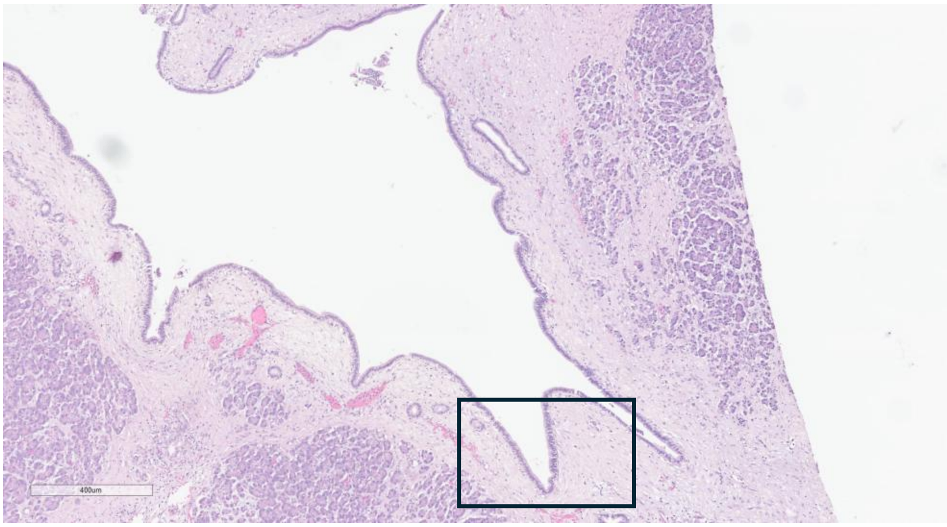
400um

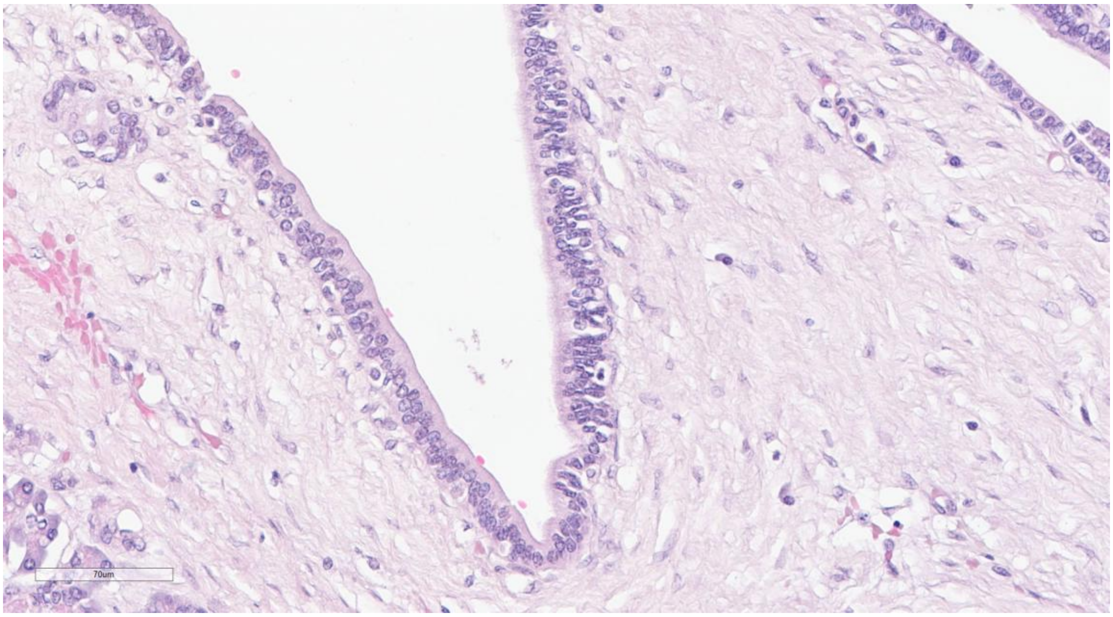
70um

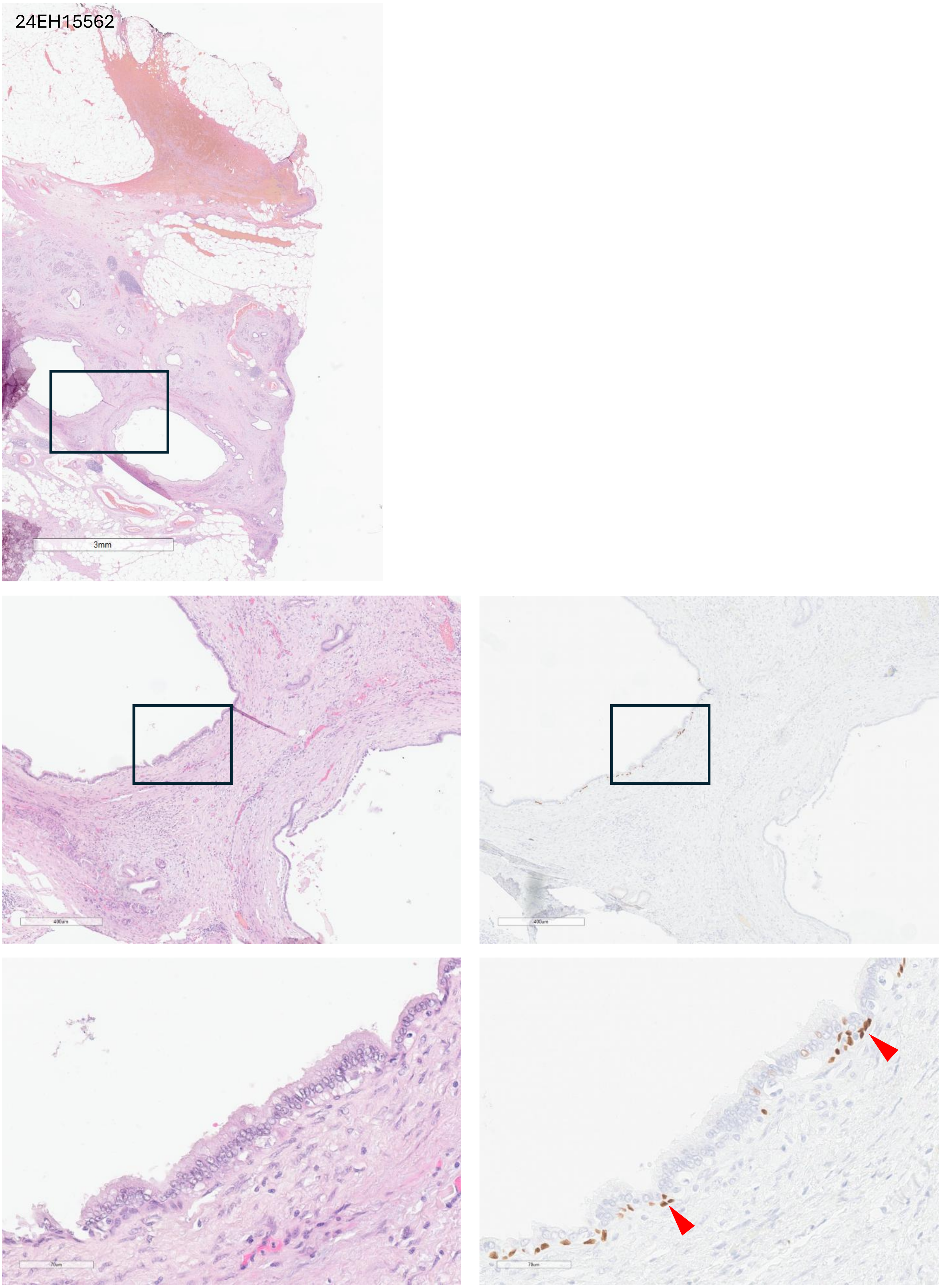
24EH15562
3mm
400um
400um
70um
70um

24EH16955

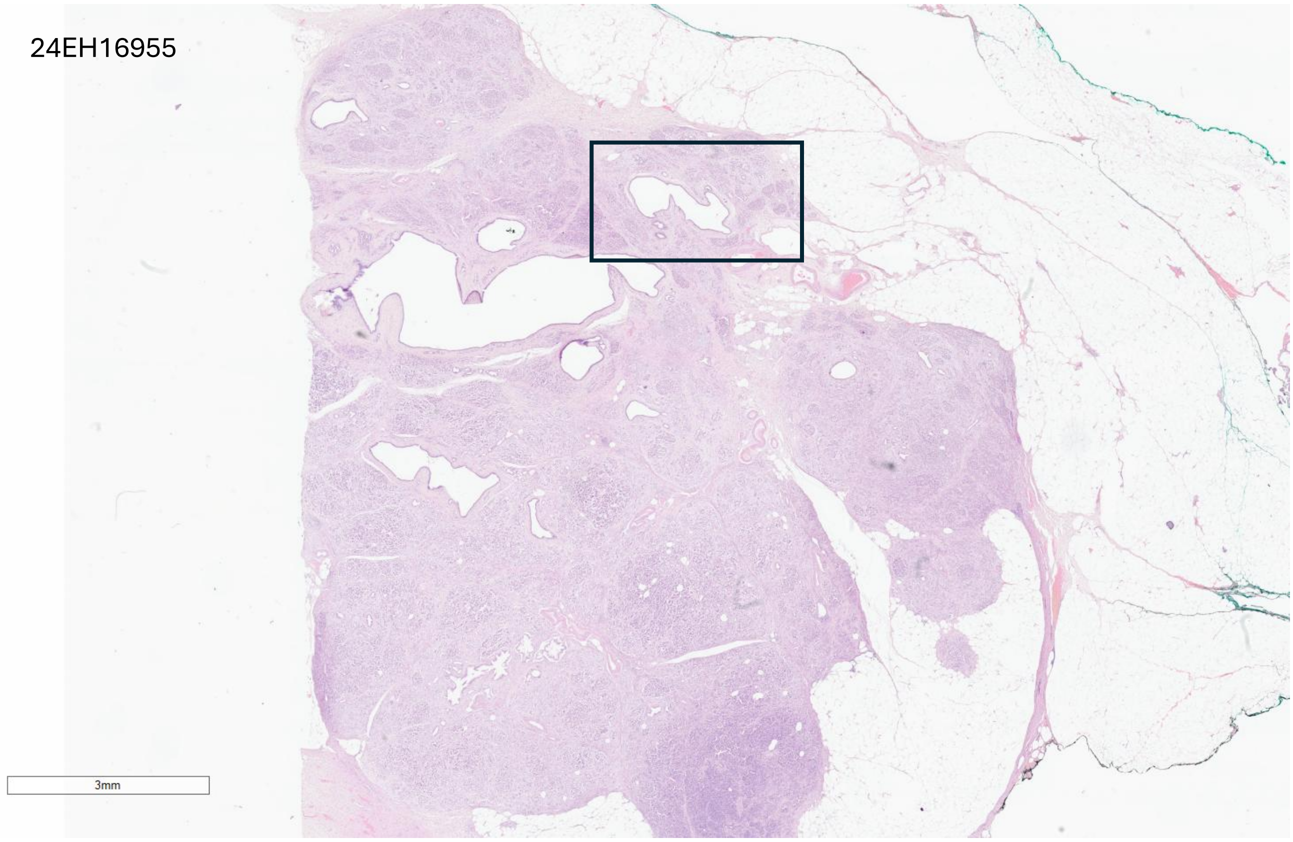


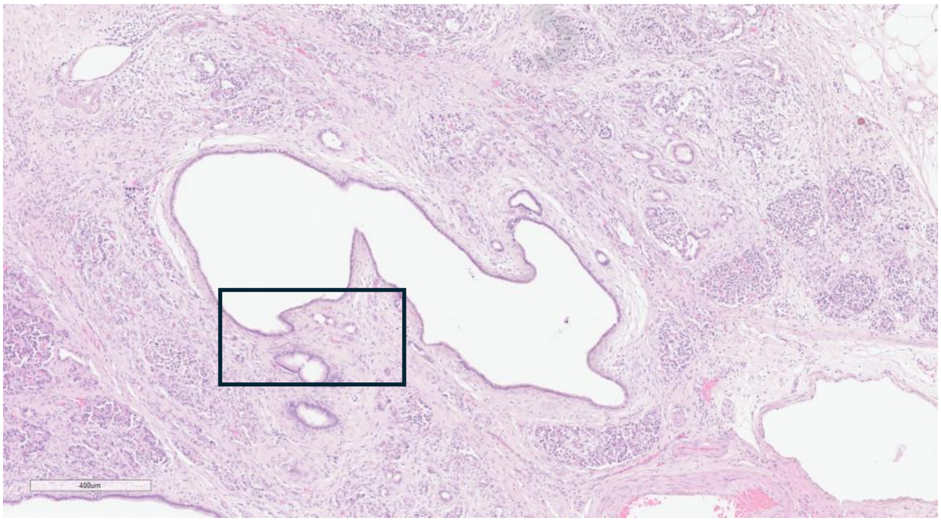


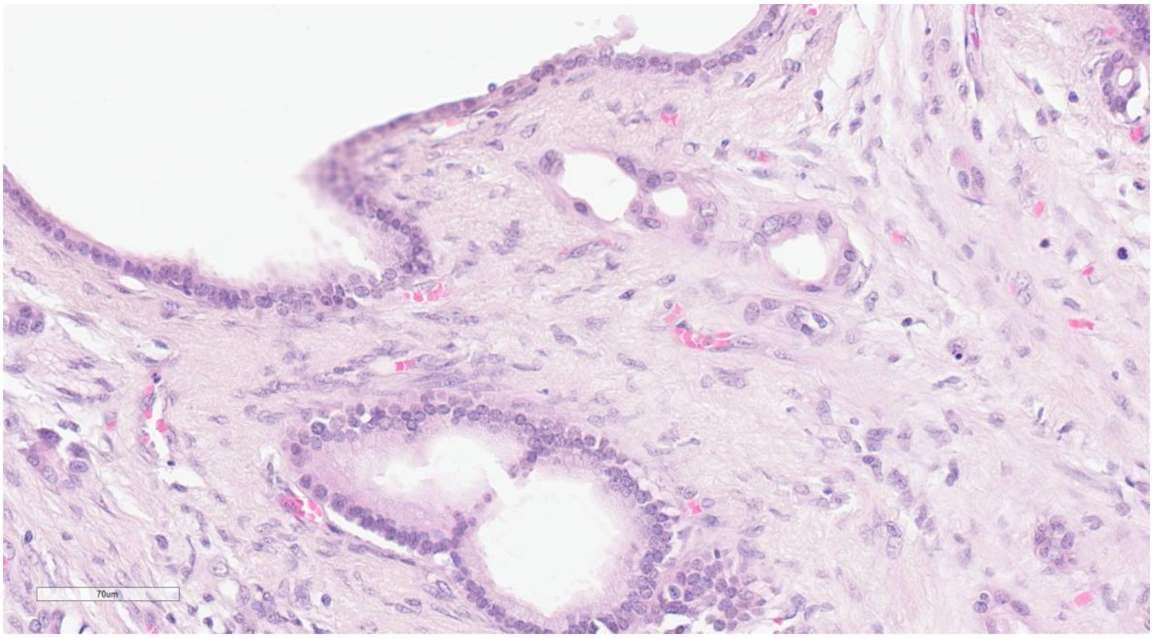

25EH018

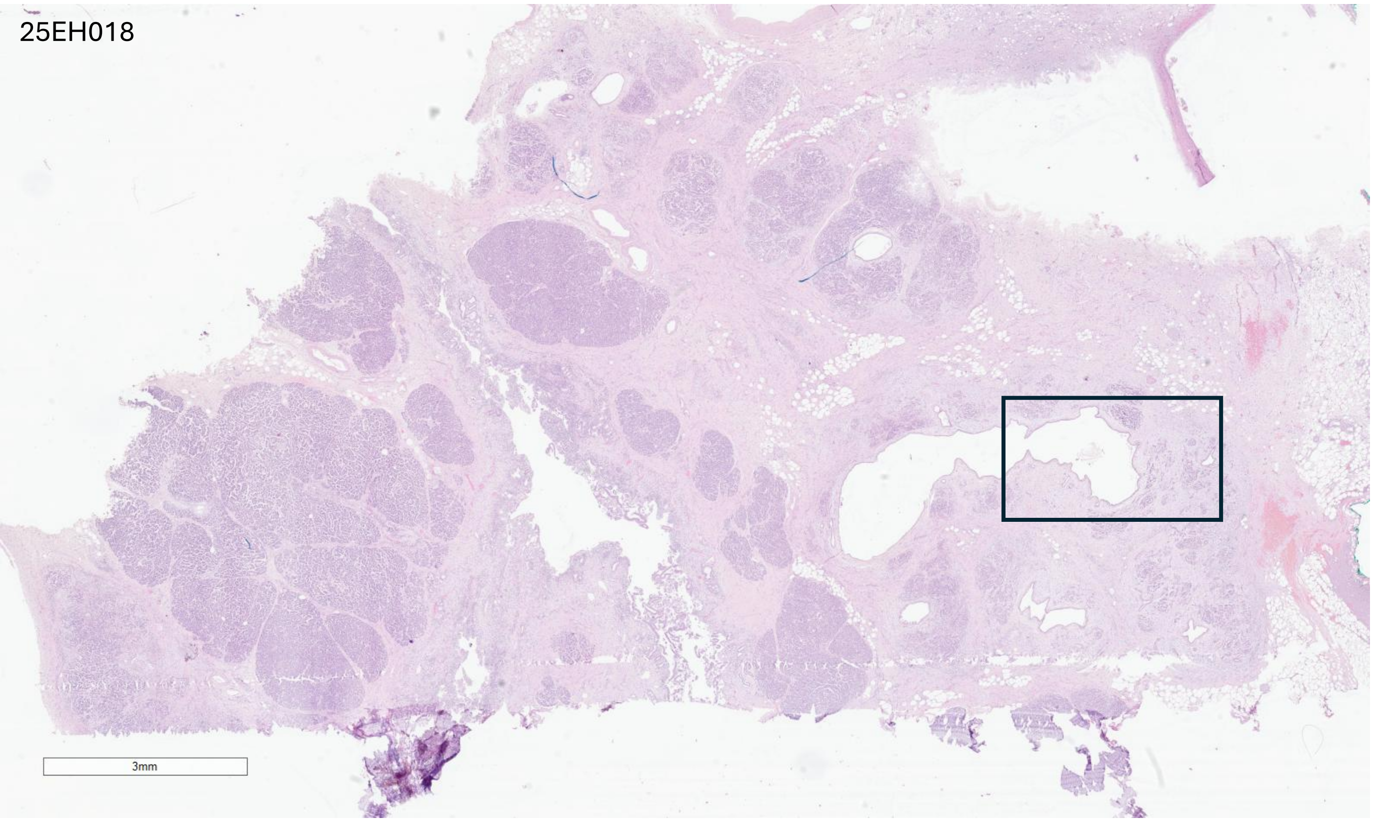


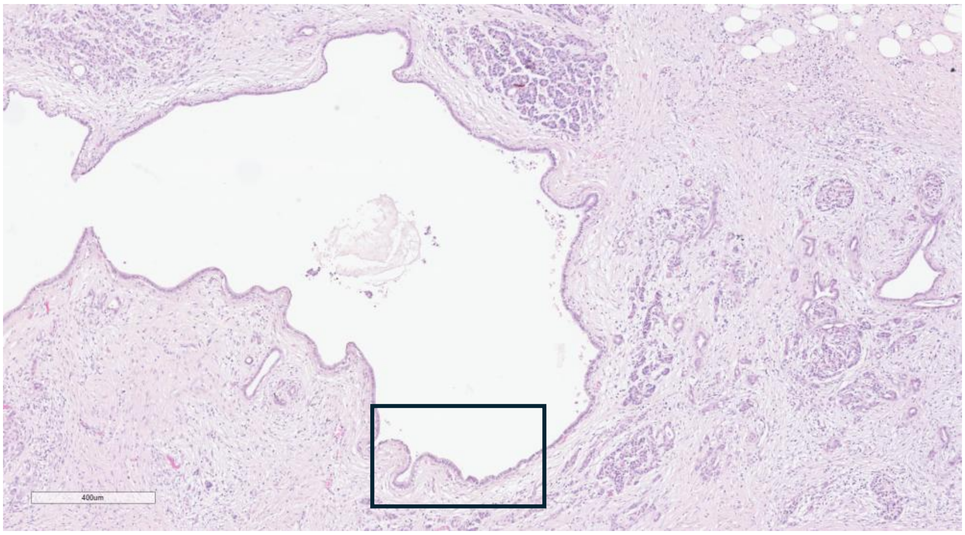


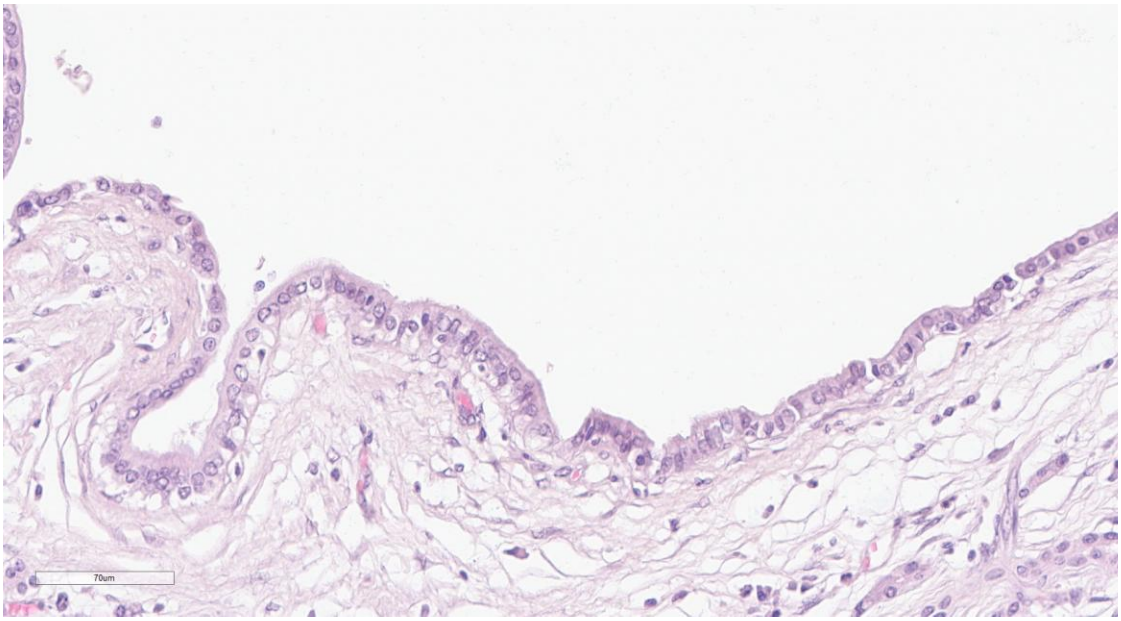

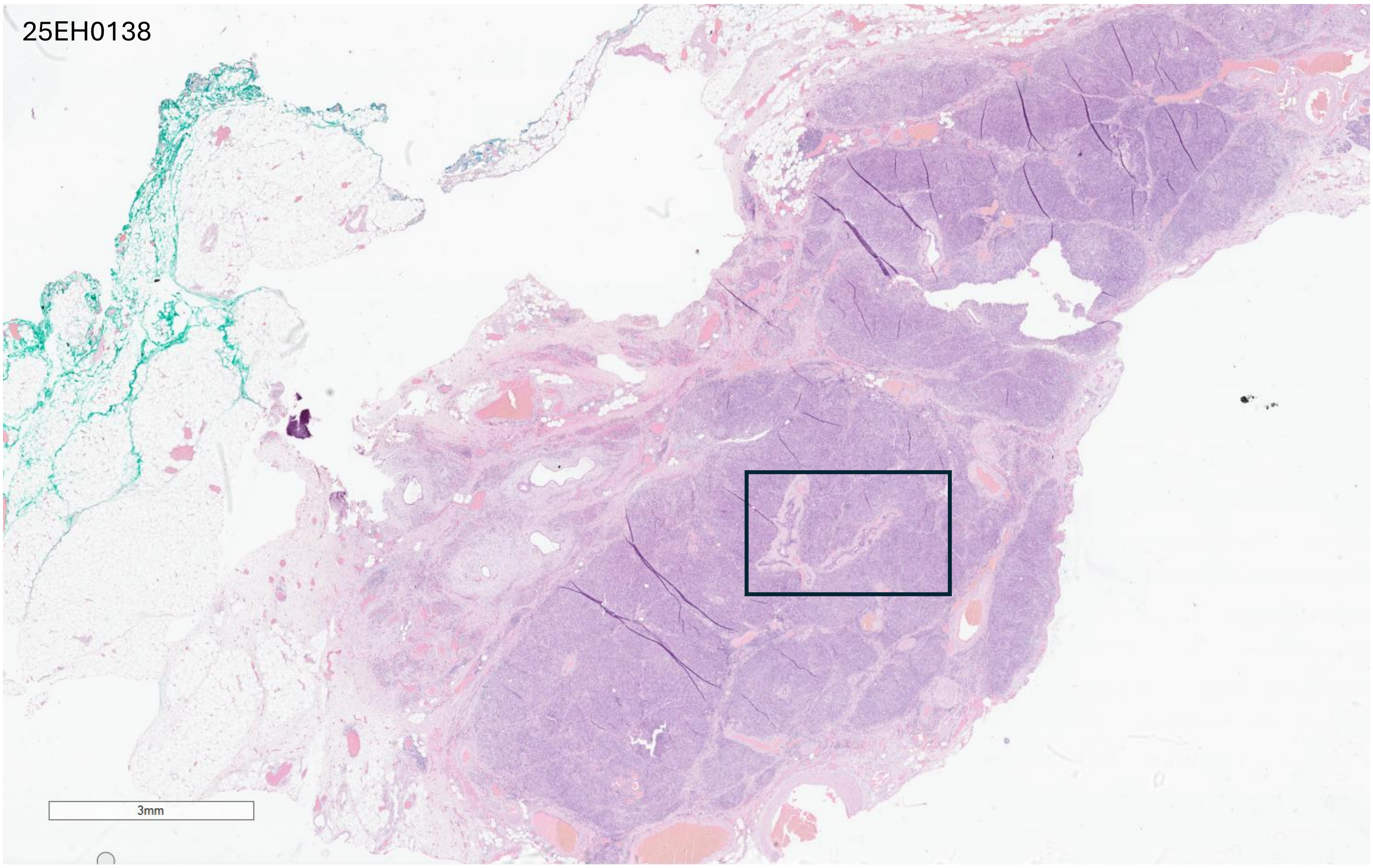
25EH0138
3mm
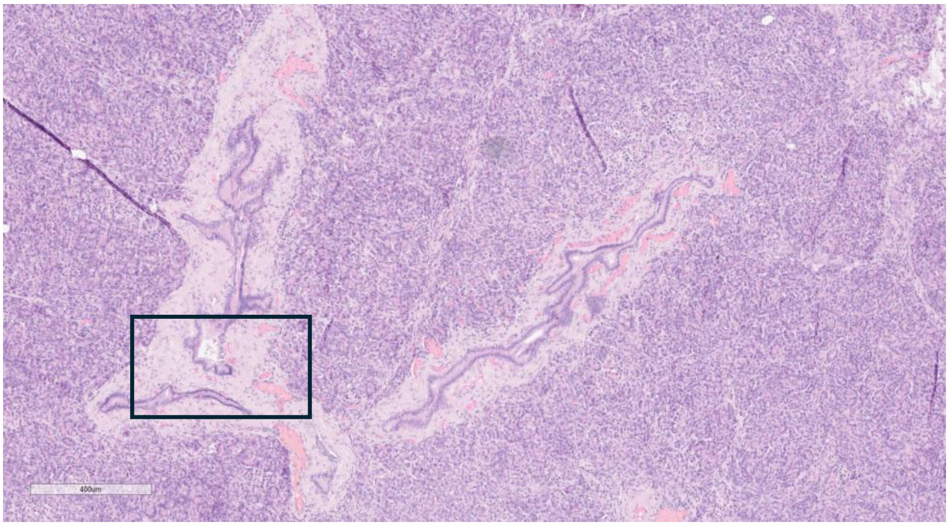
400um
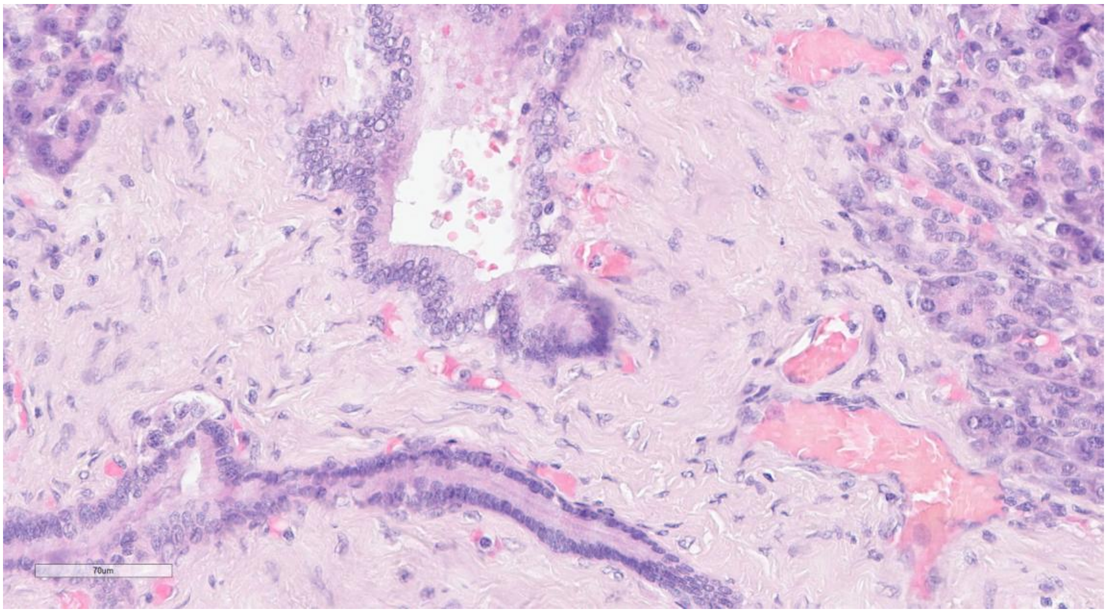
70um

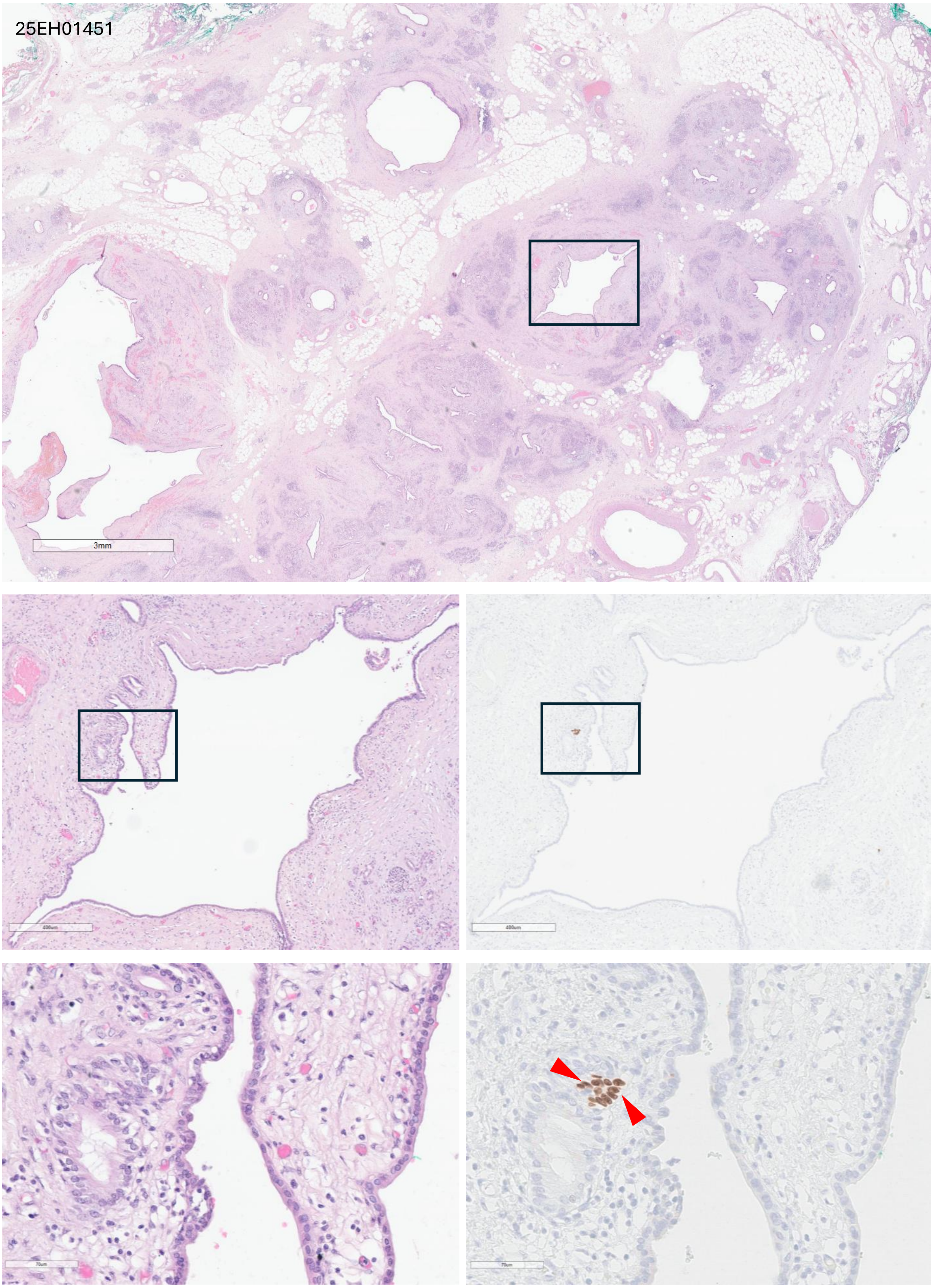

25EH01451
3mm
400um
400um
70um
70um

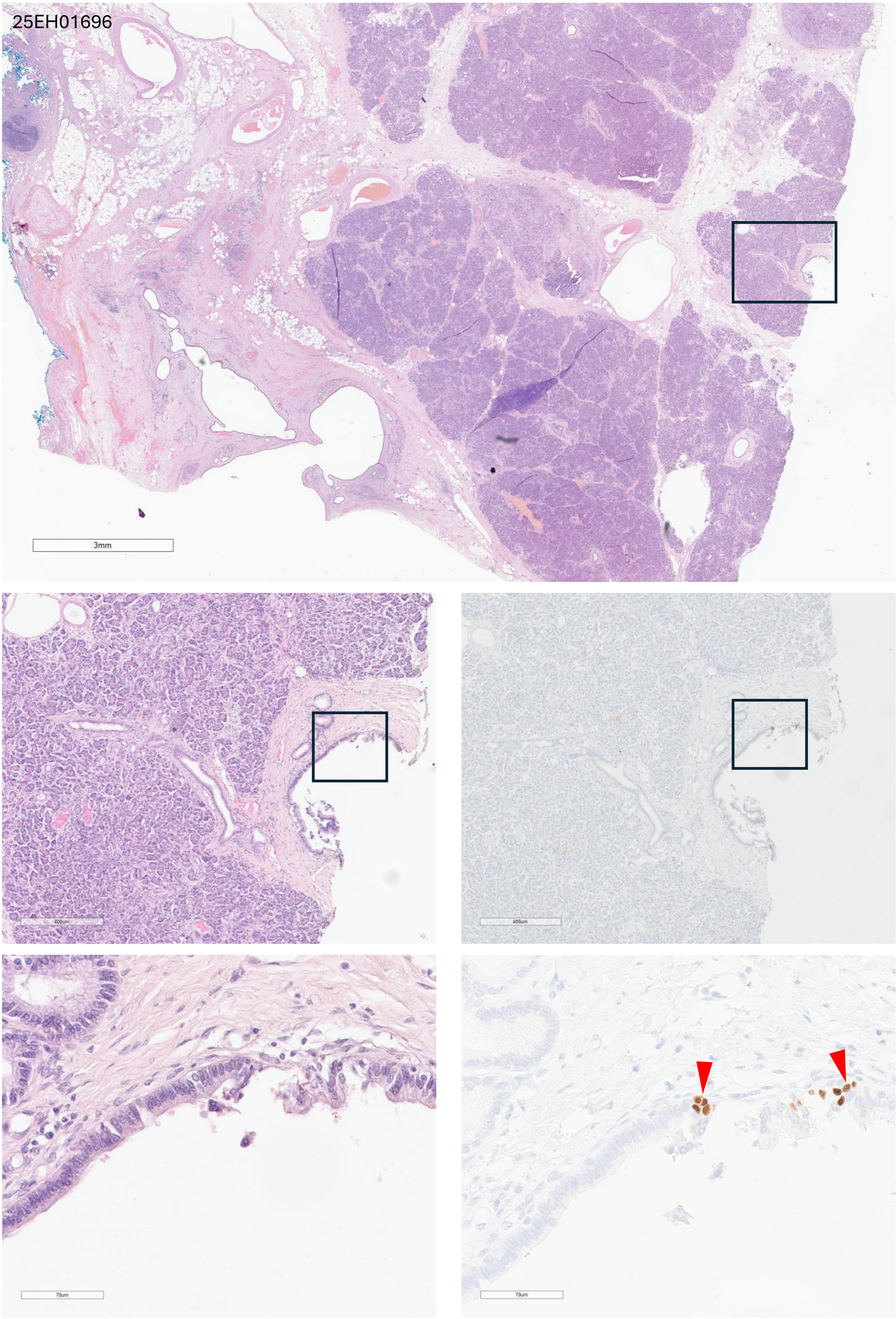
25EH01696
3mm
400um
400um
70um
70um

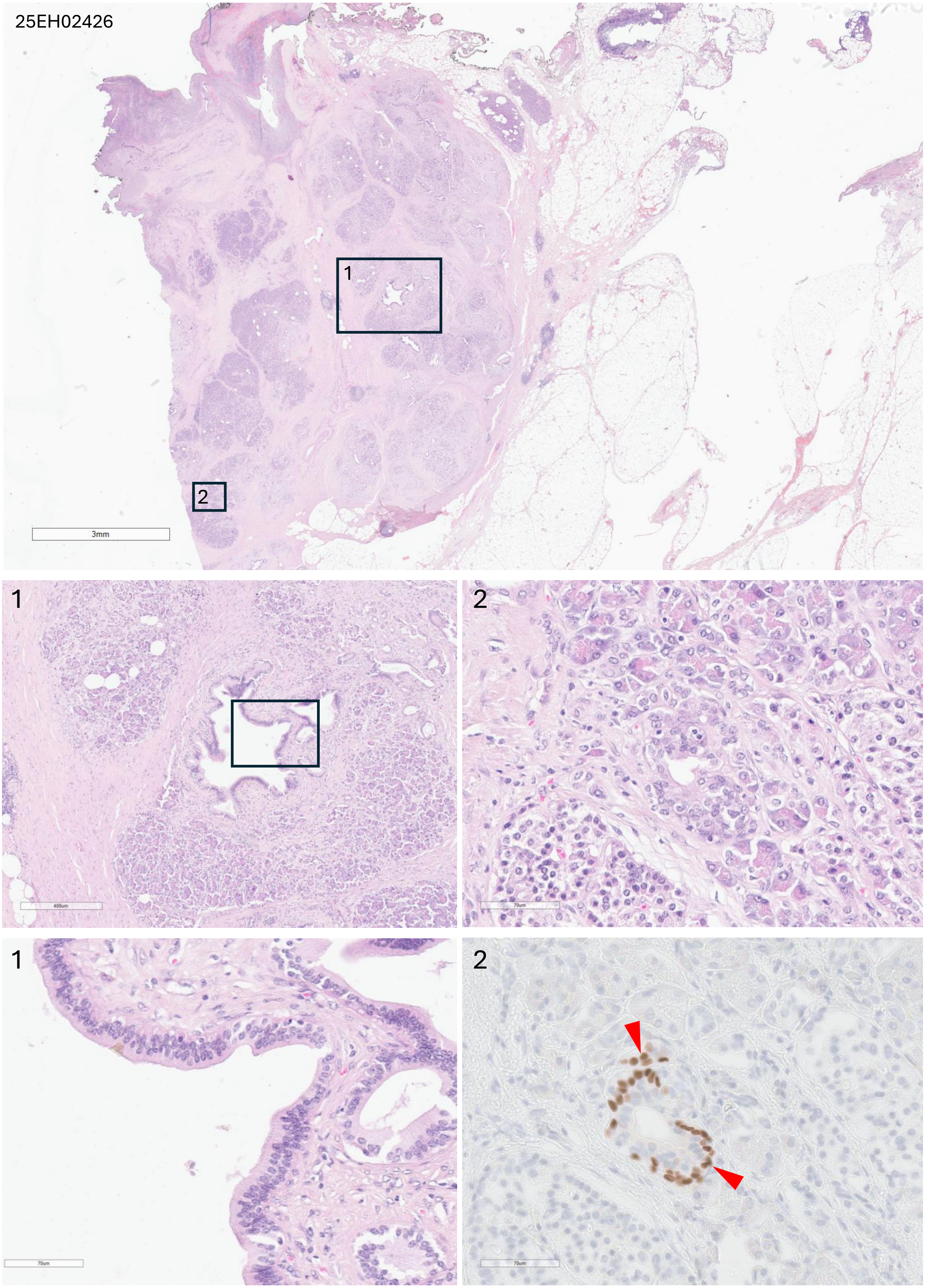
25EH02426
1
2
3mm
400um
70um

25EH03908

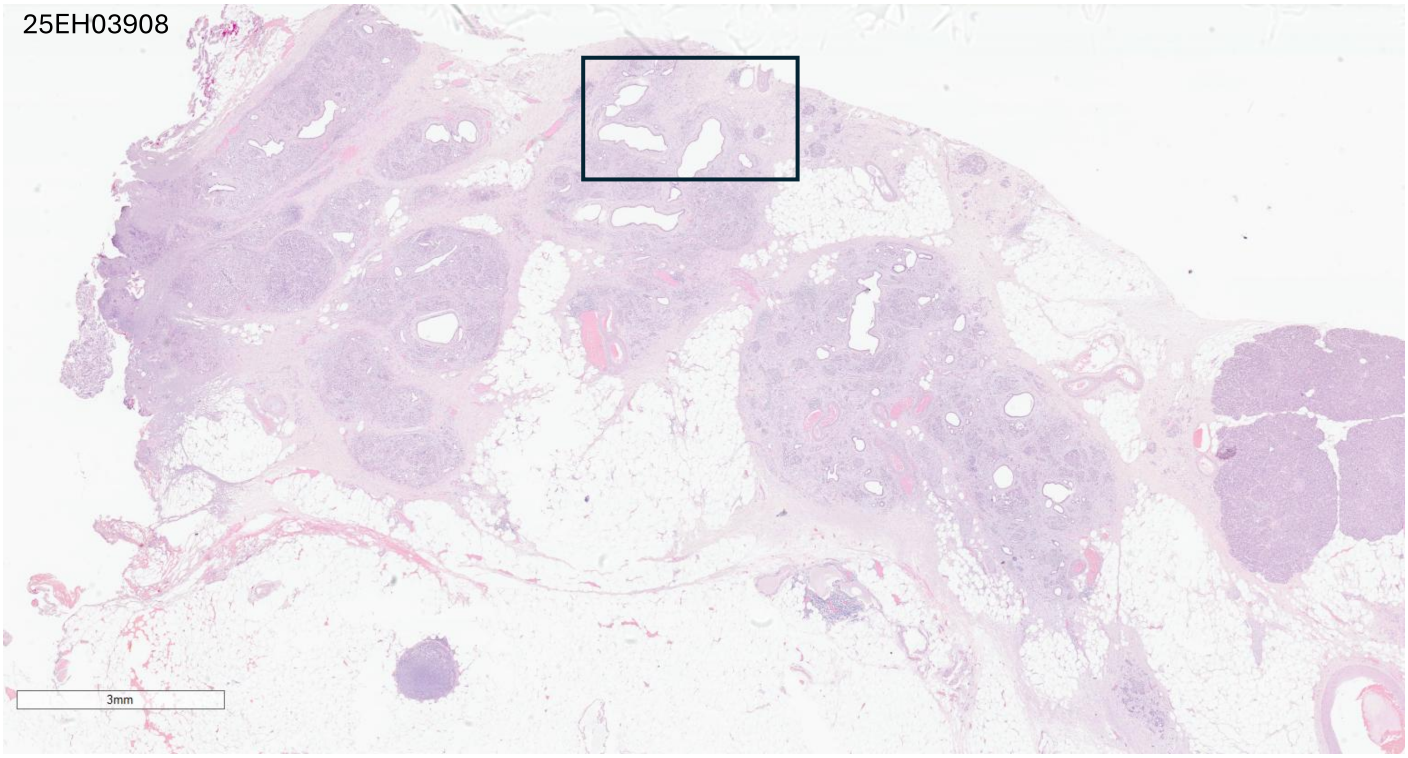


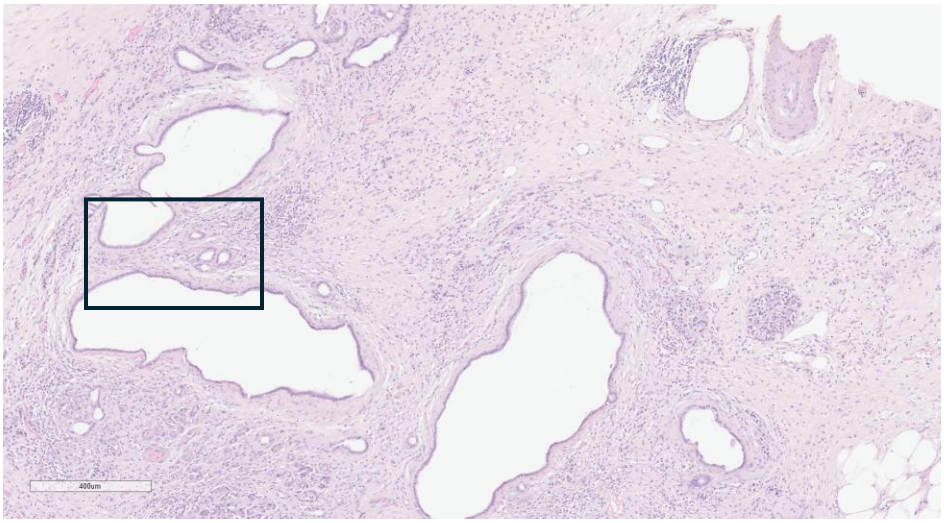


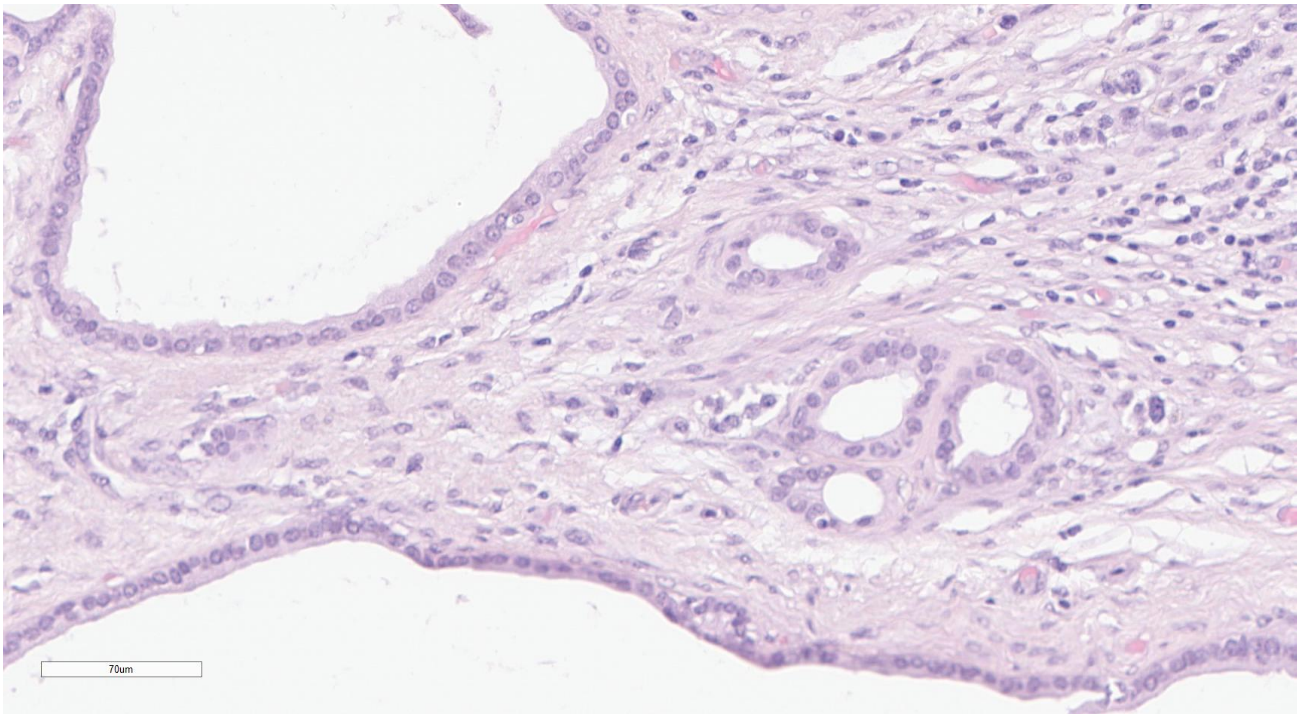

25EH04737

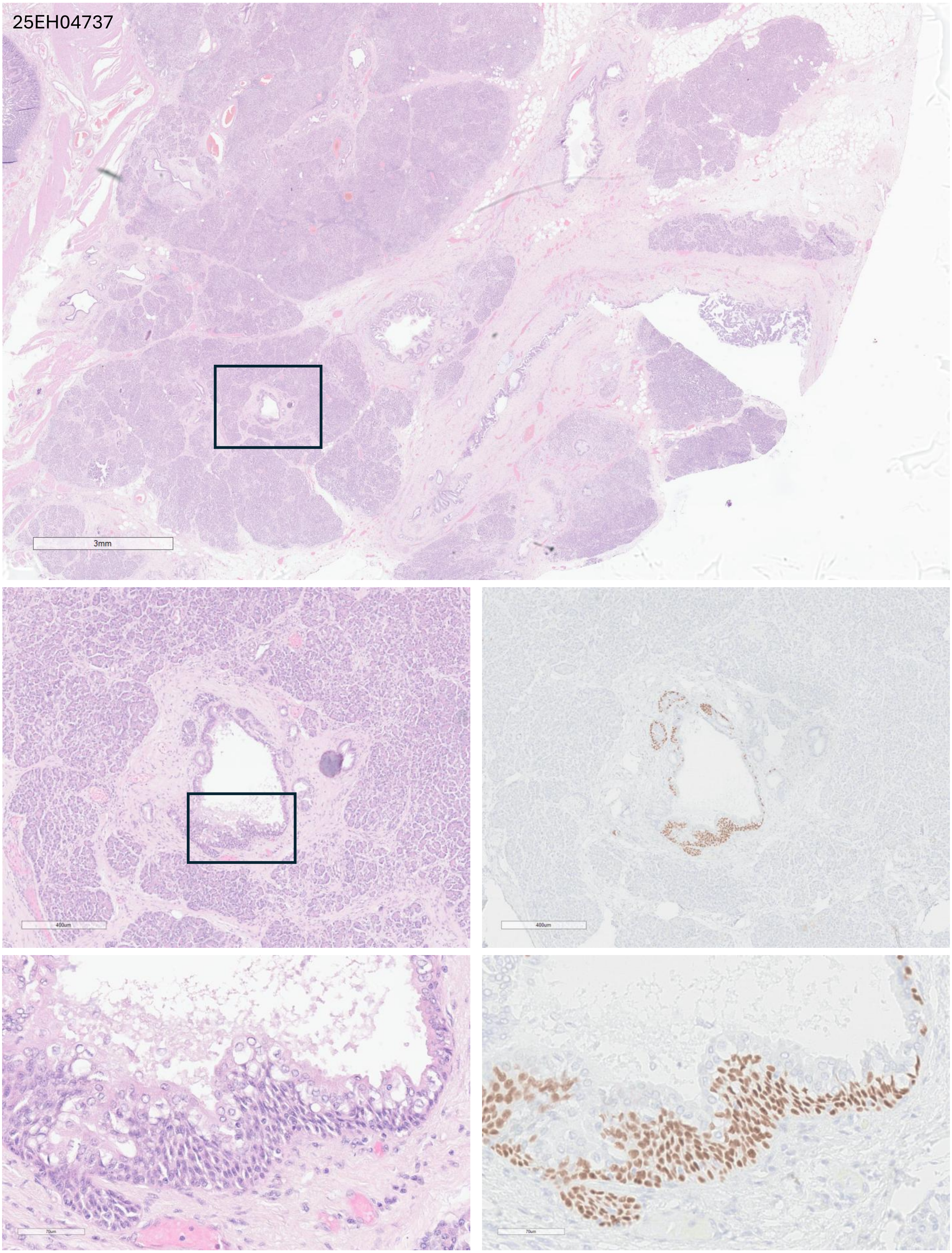

25EH04932

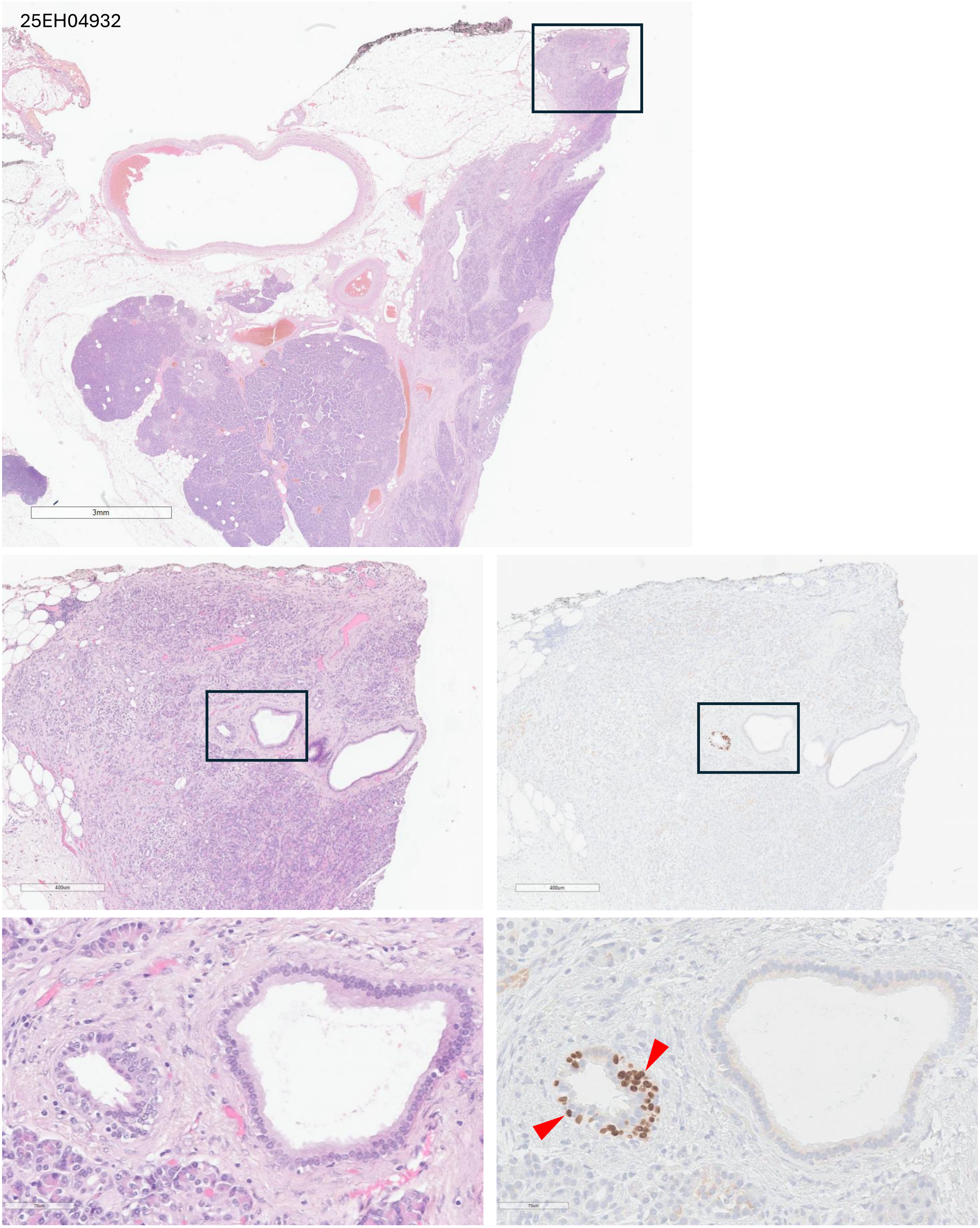

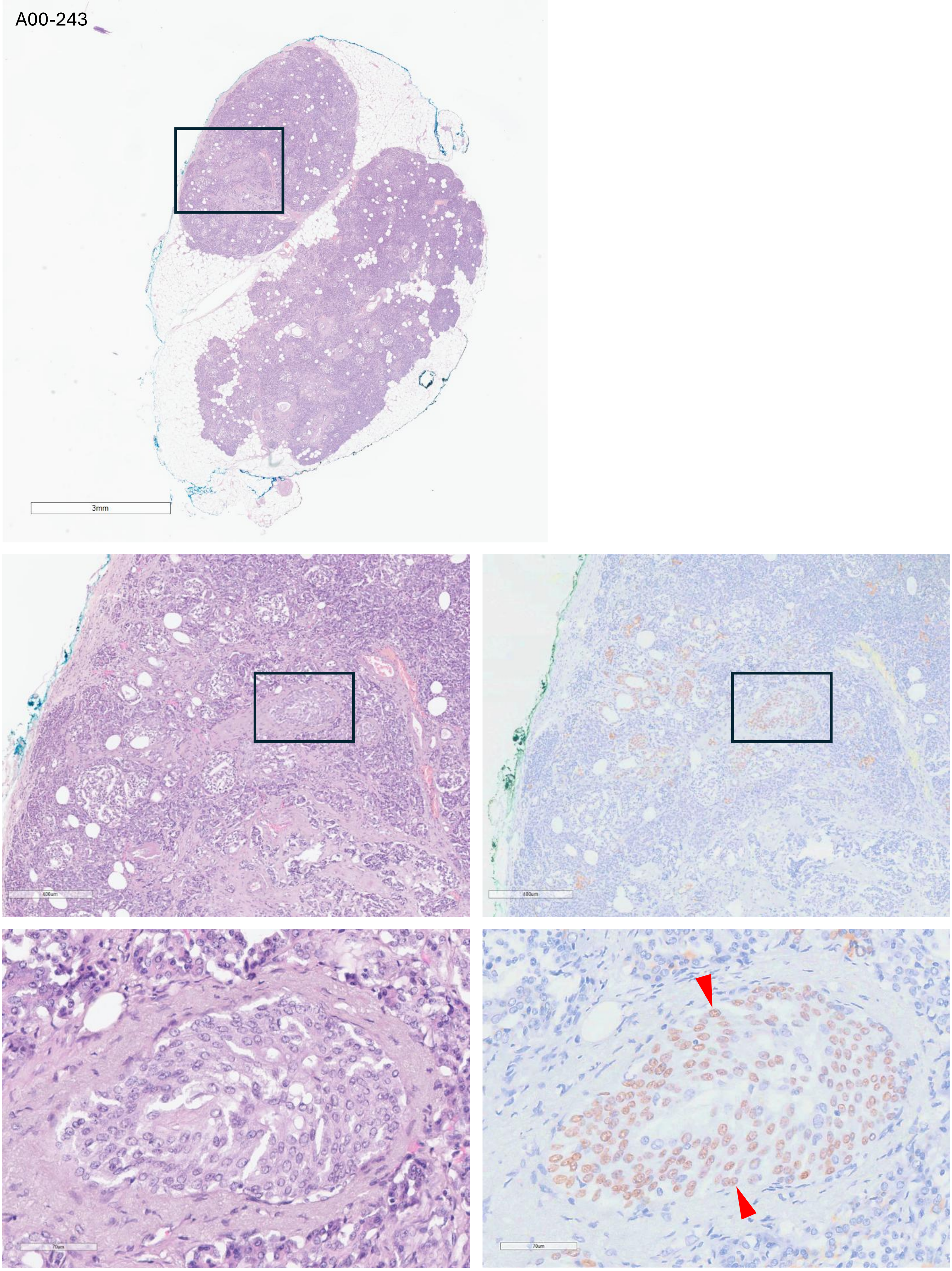
A00-243
3mm
400um
400um
70um
70um

### Adenosquamous Carcinoma

ASCP-1

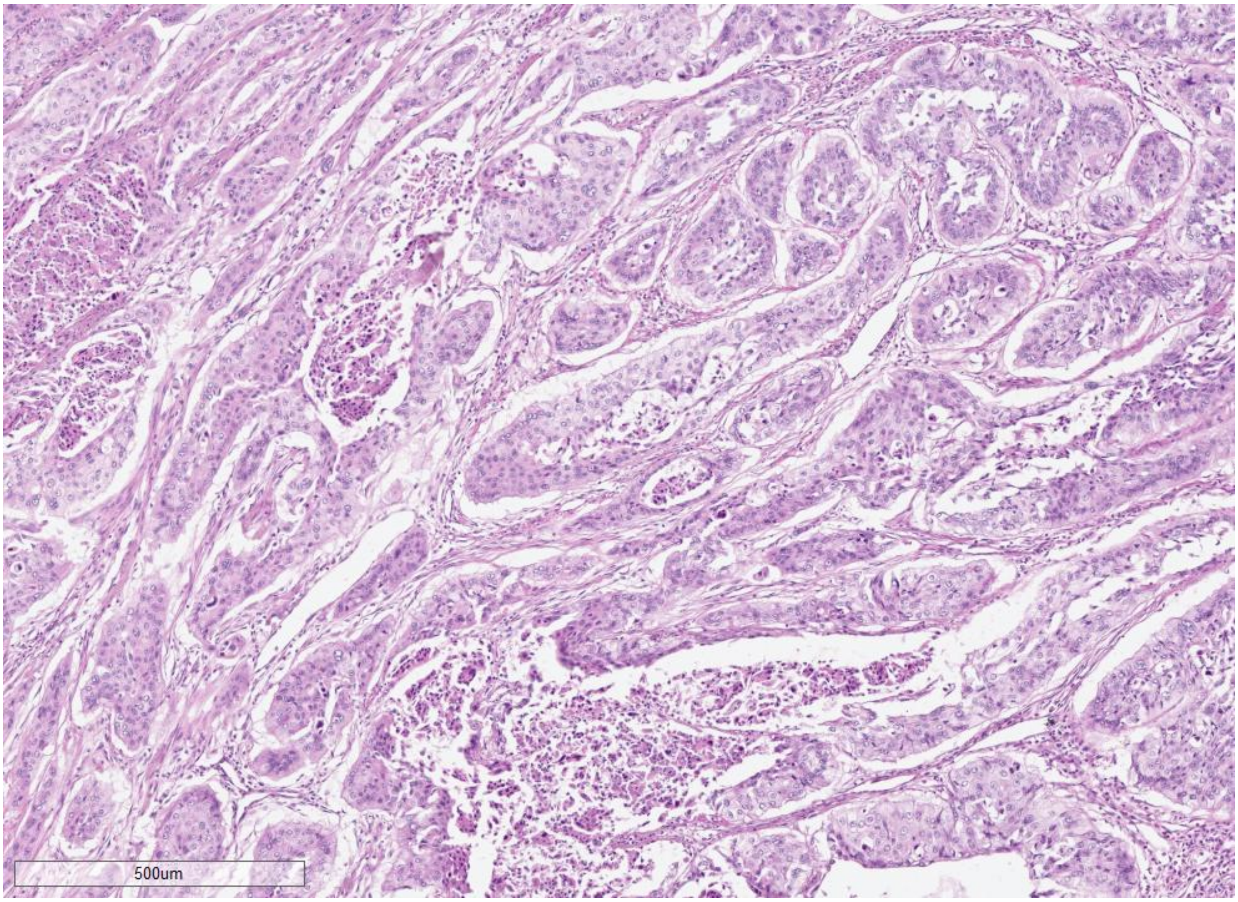

ASCP-2

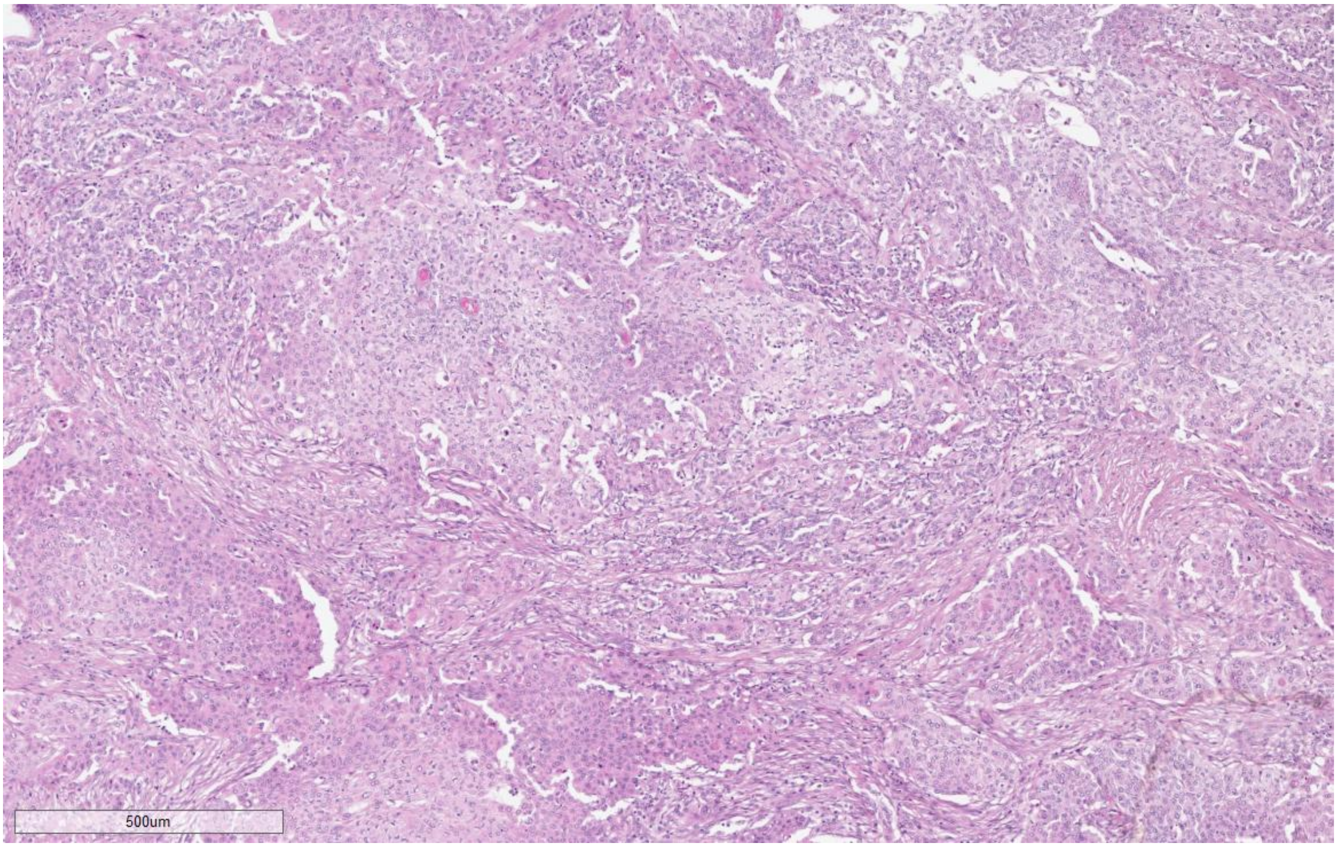

ASCP-3

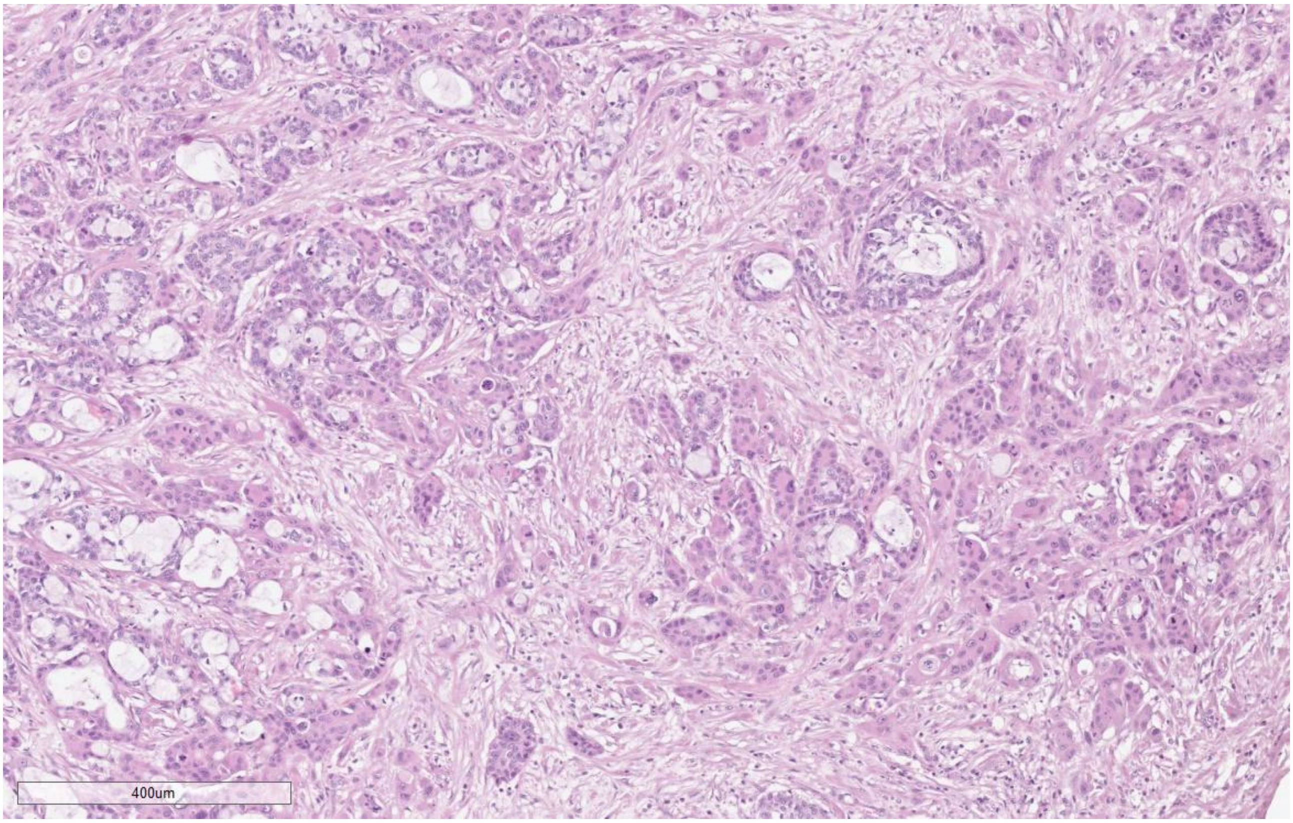

ASCP-4

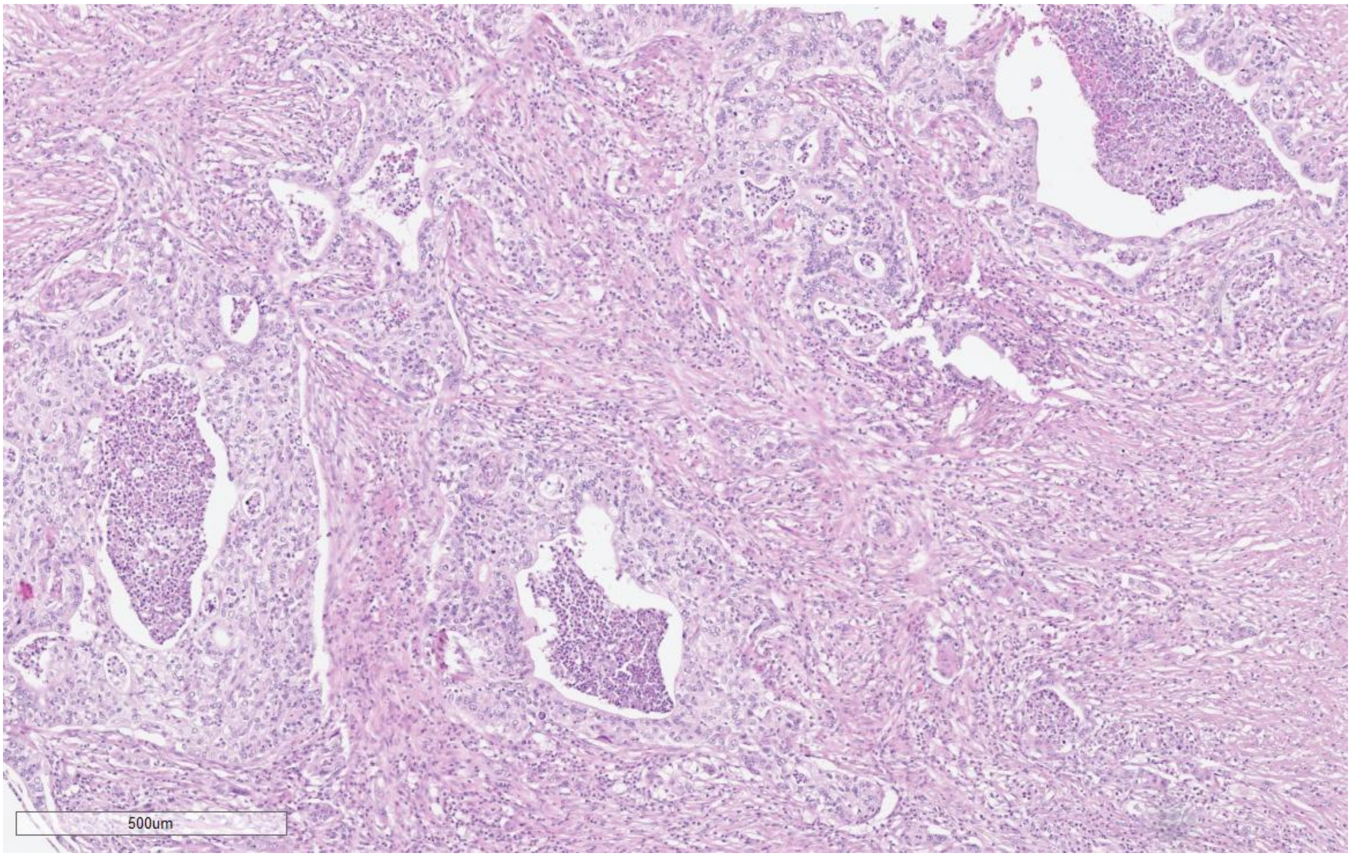

ASCP-5

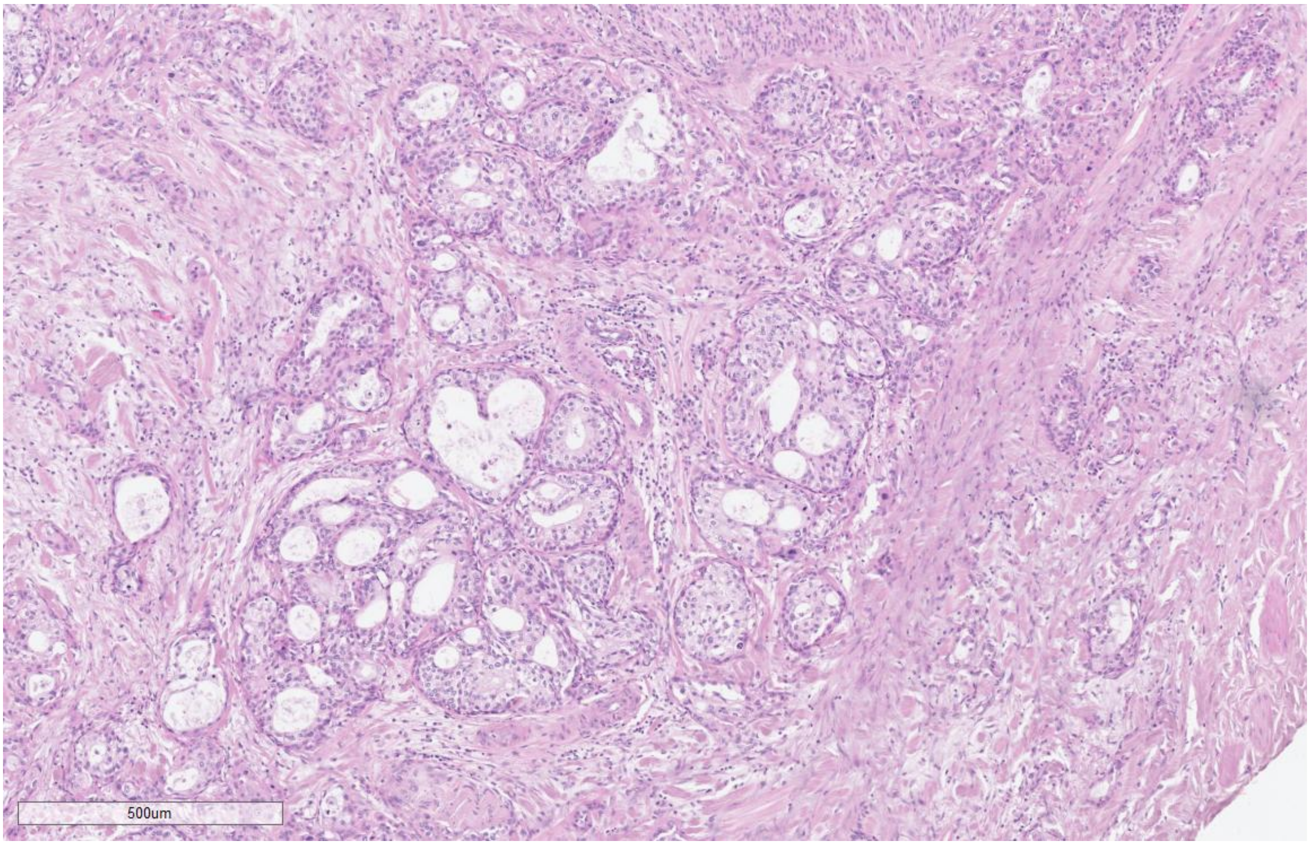

ASCP-6

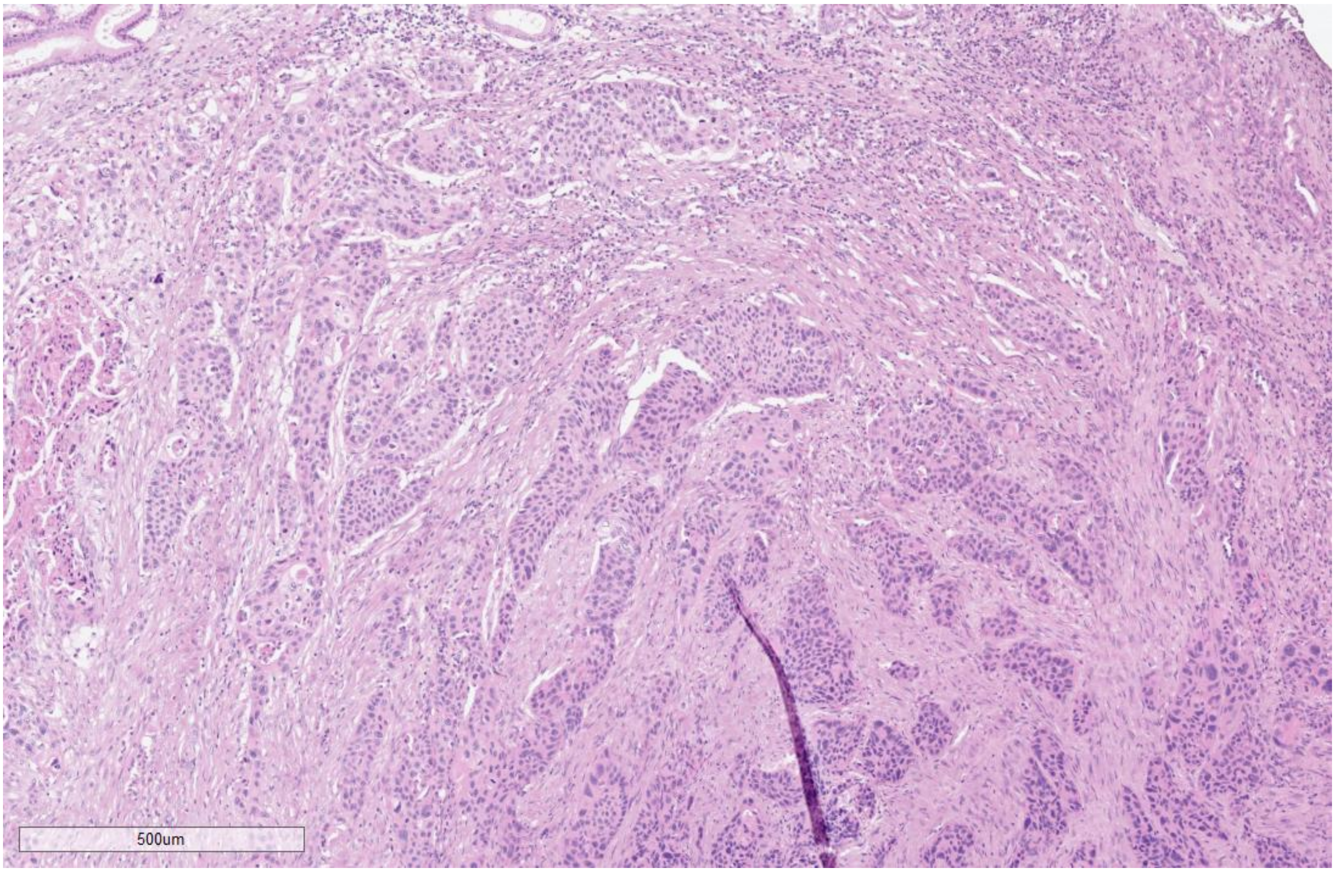

ASCP-7

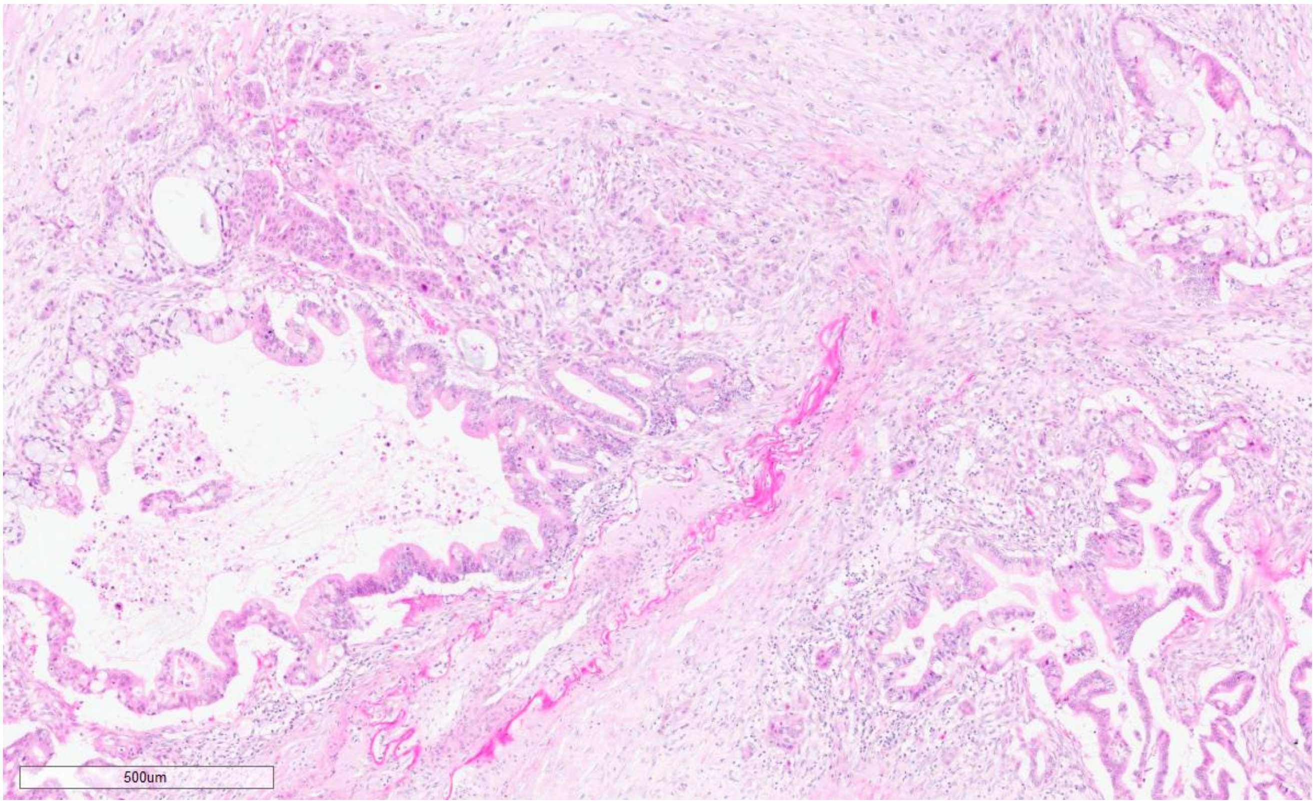

ASCP-8

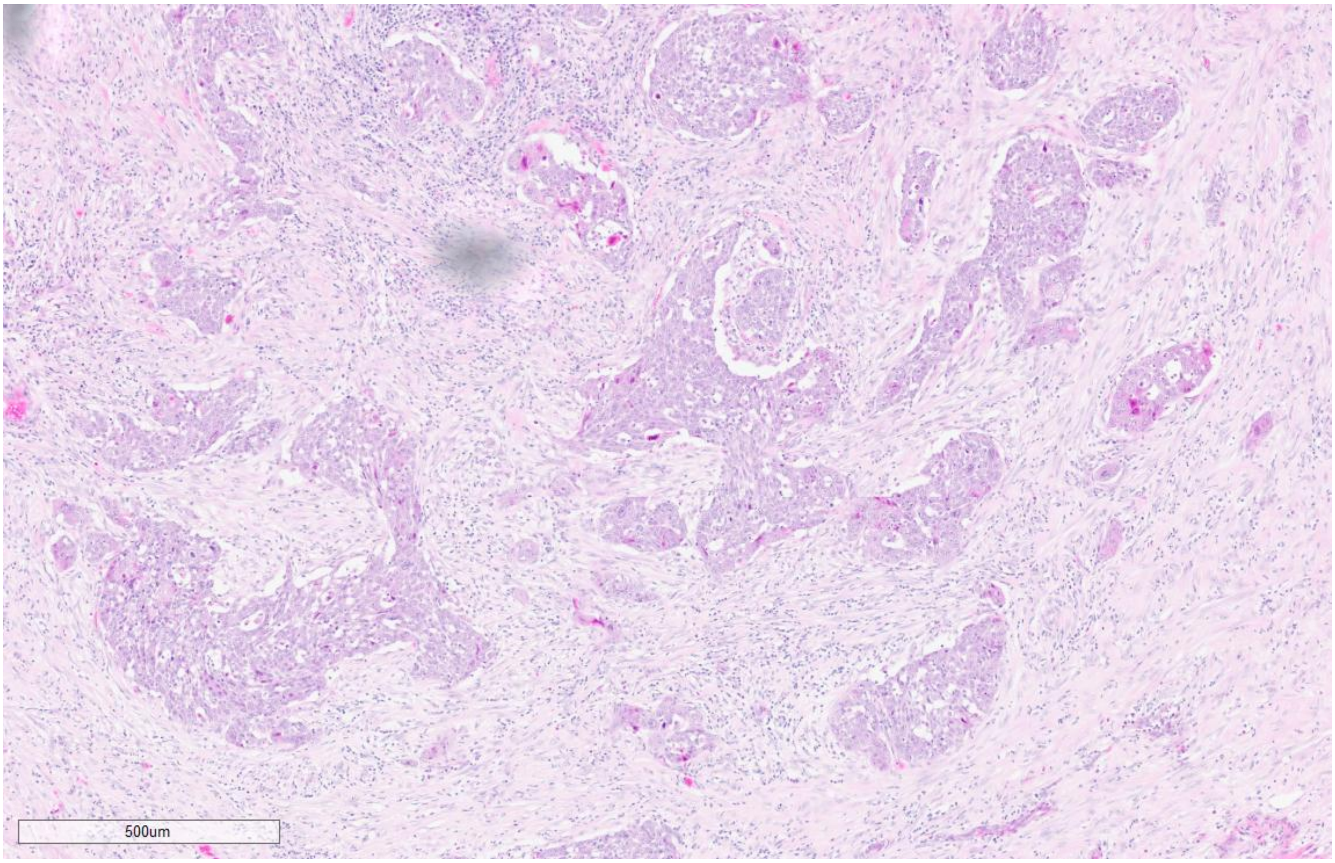

ASCP-9

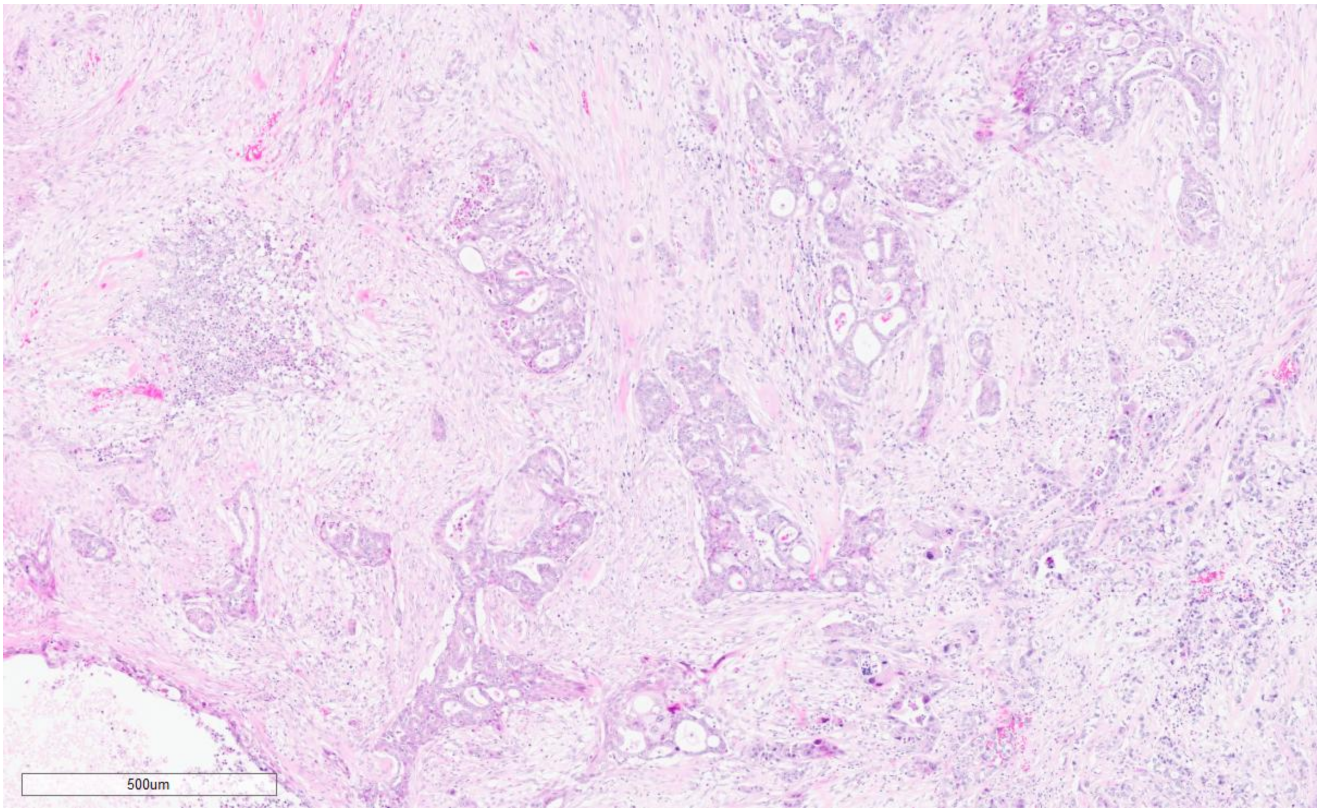

ASCP-10

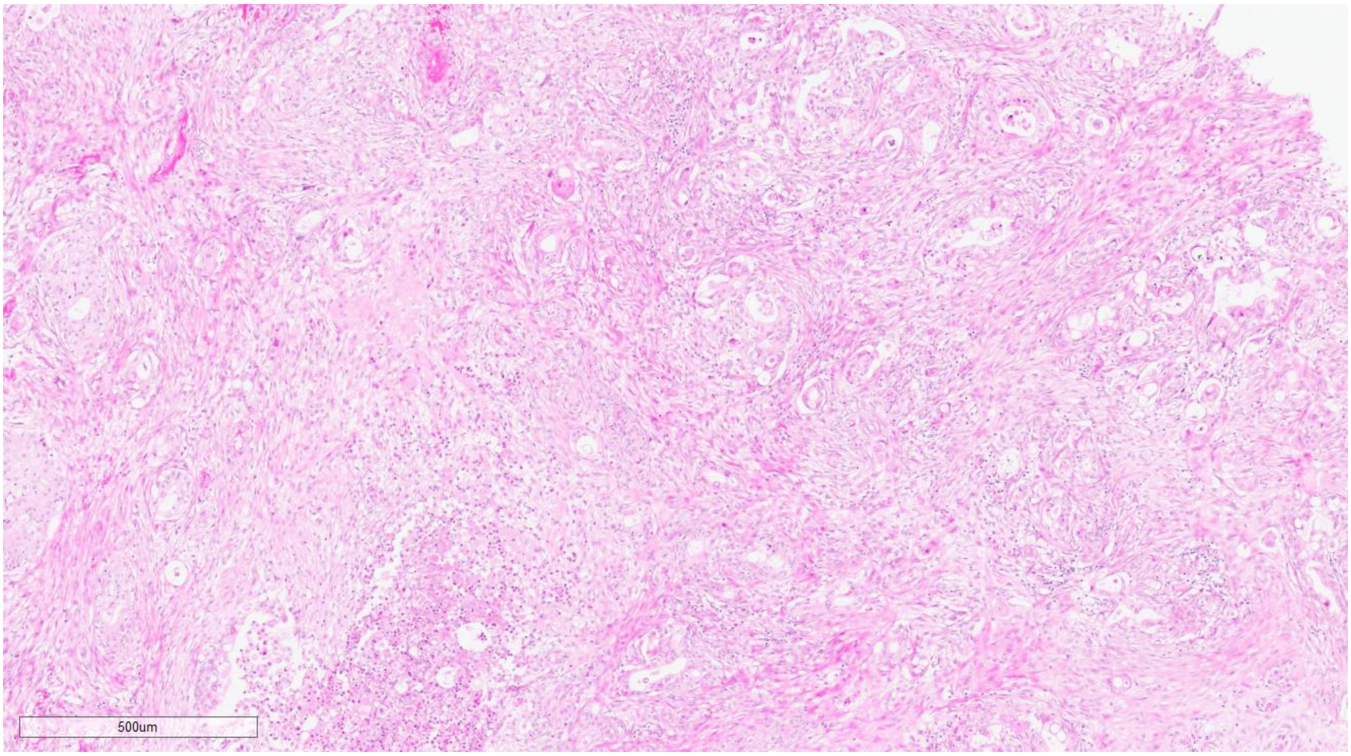

ASCP-11

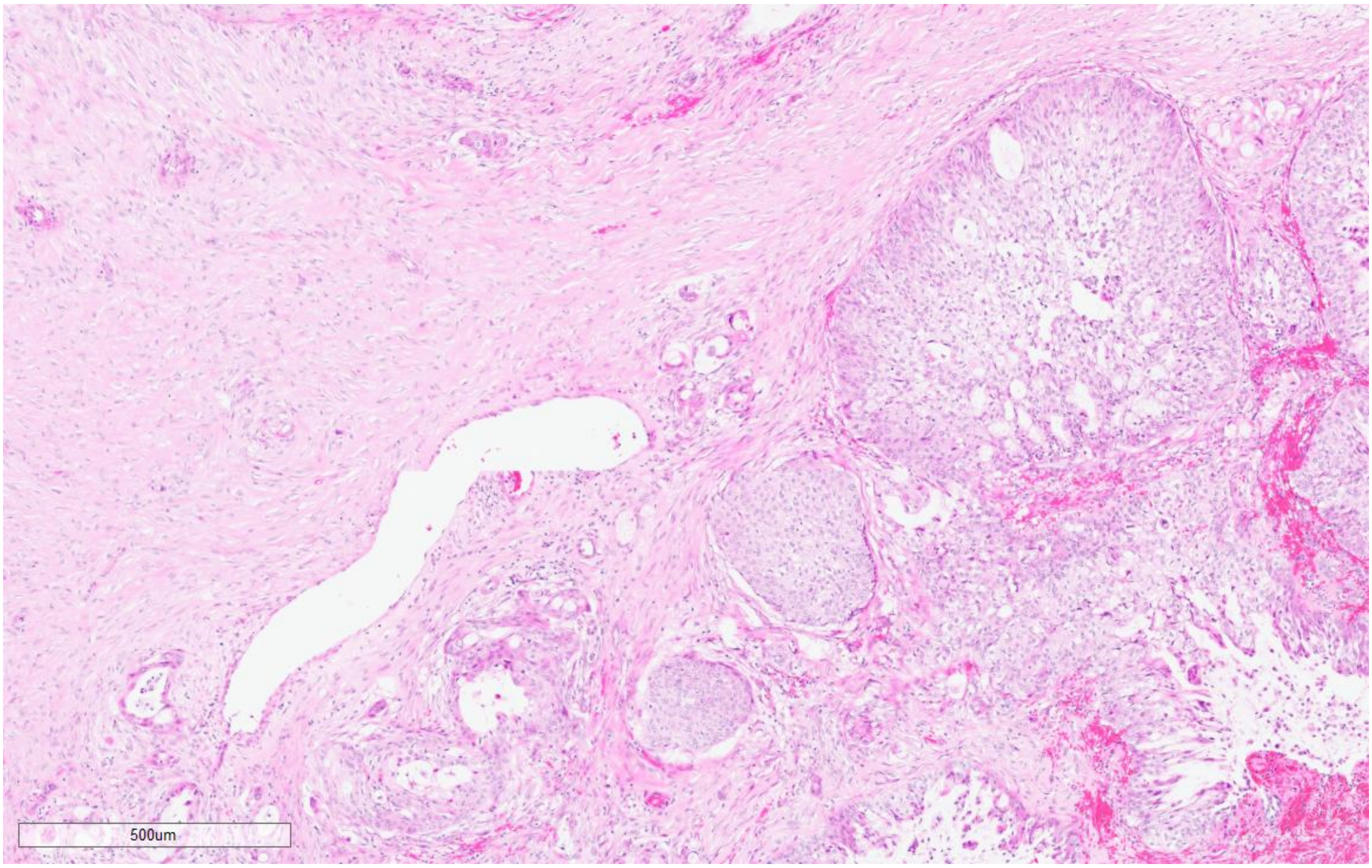

ASCP-12

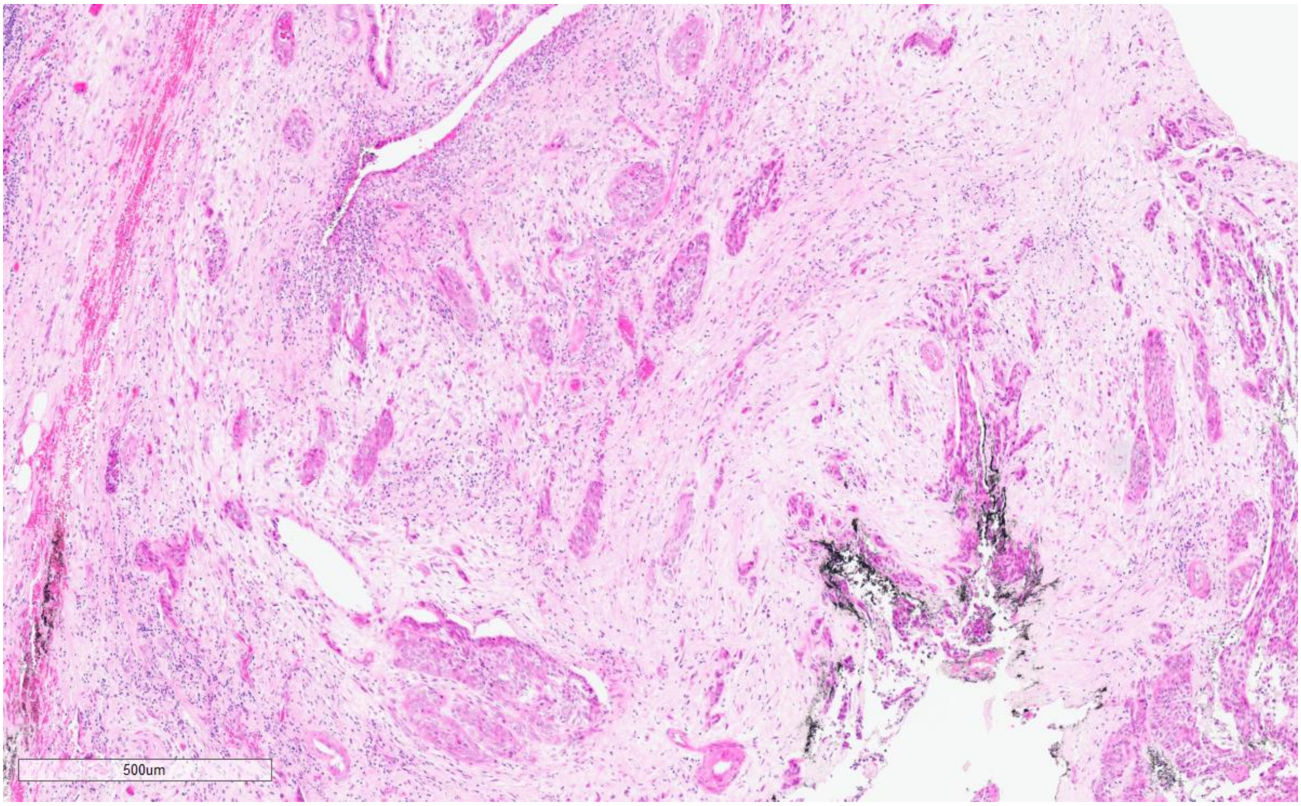

ASCP-13

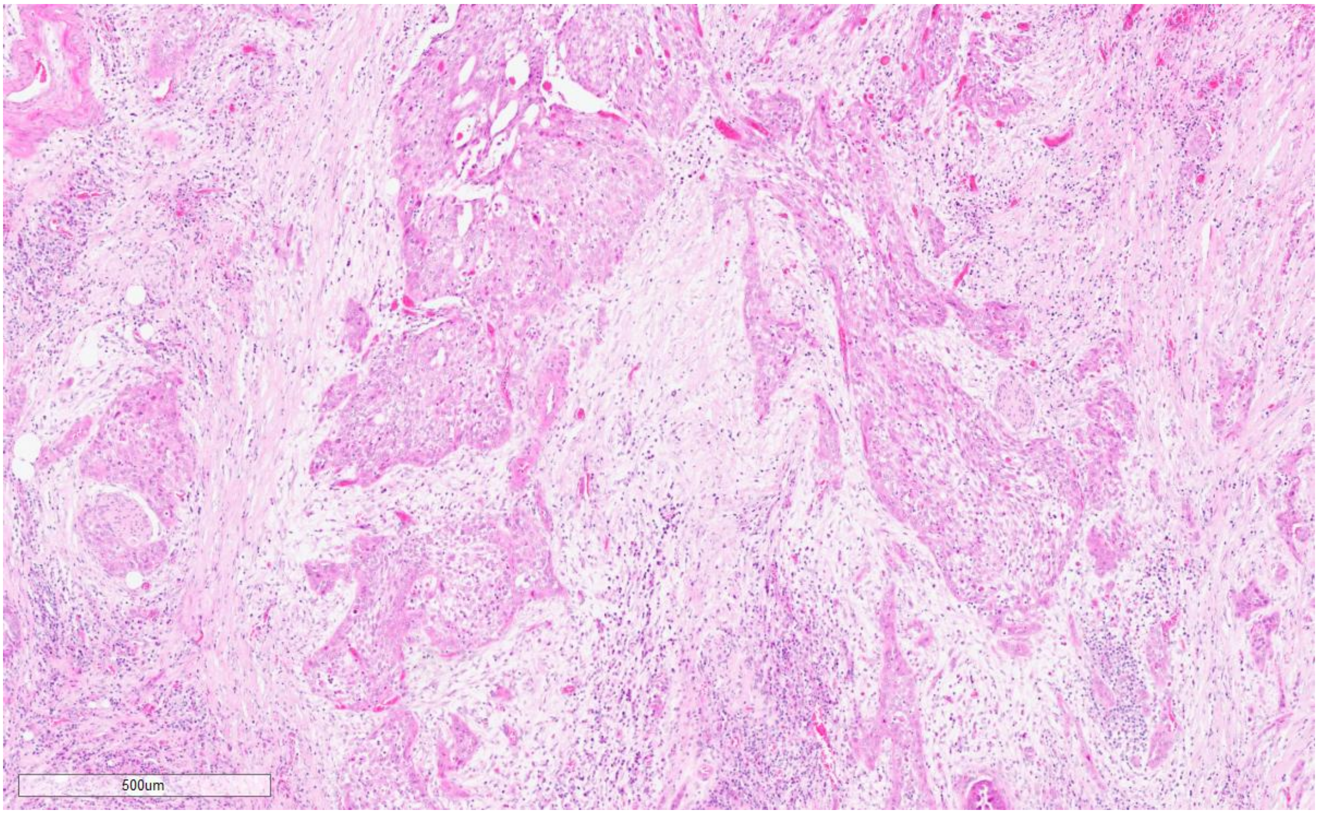

ASCP-14

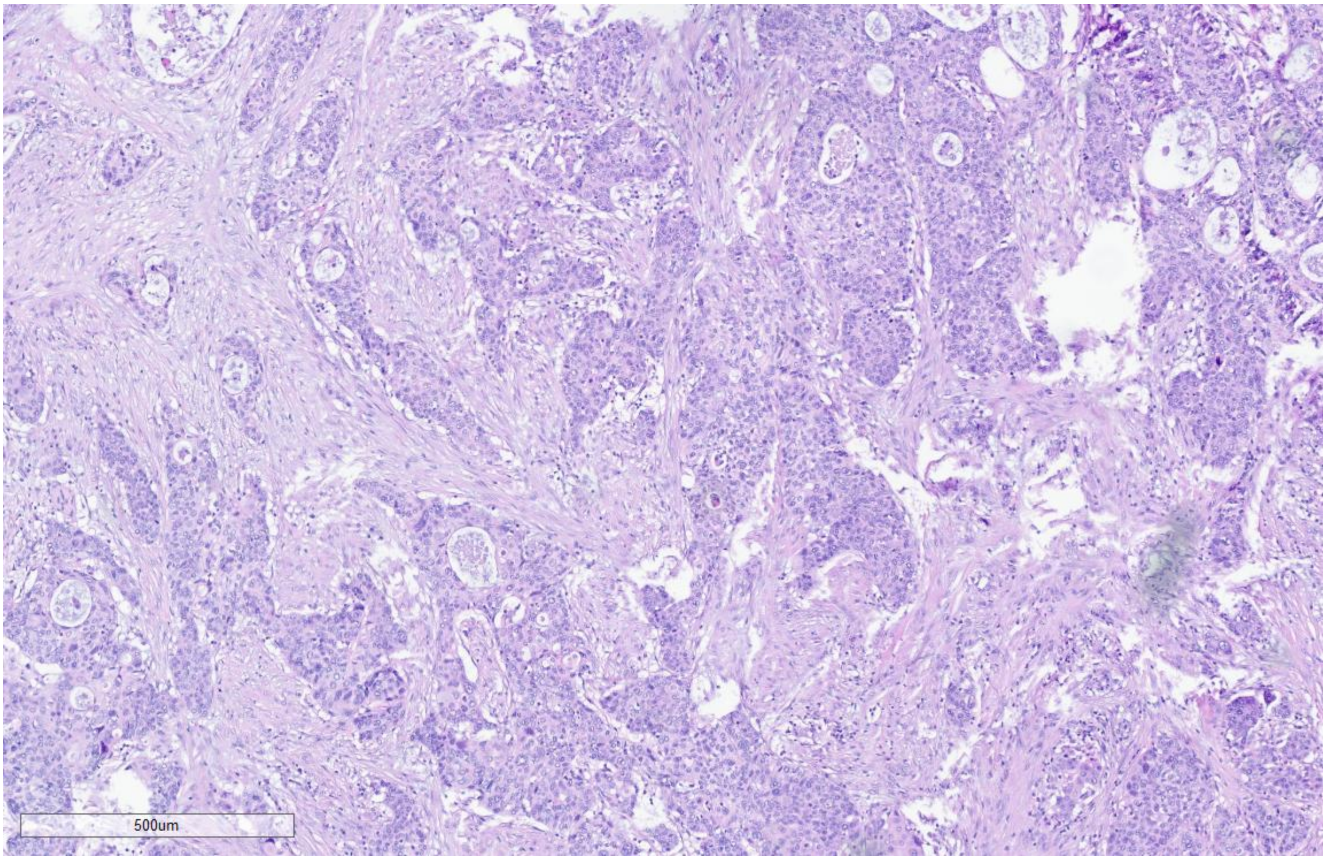

### Pancreatic Ductal Adenocarcinoma

PDAC 1 - BL

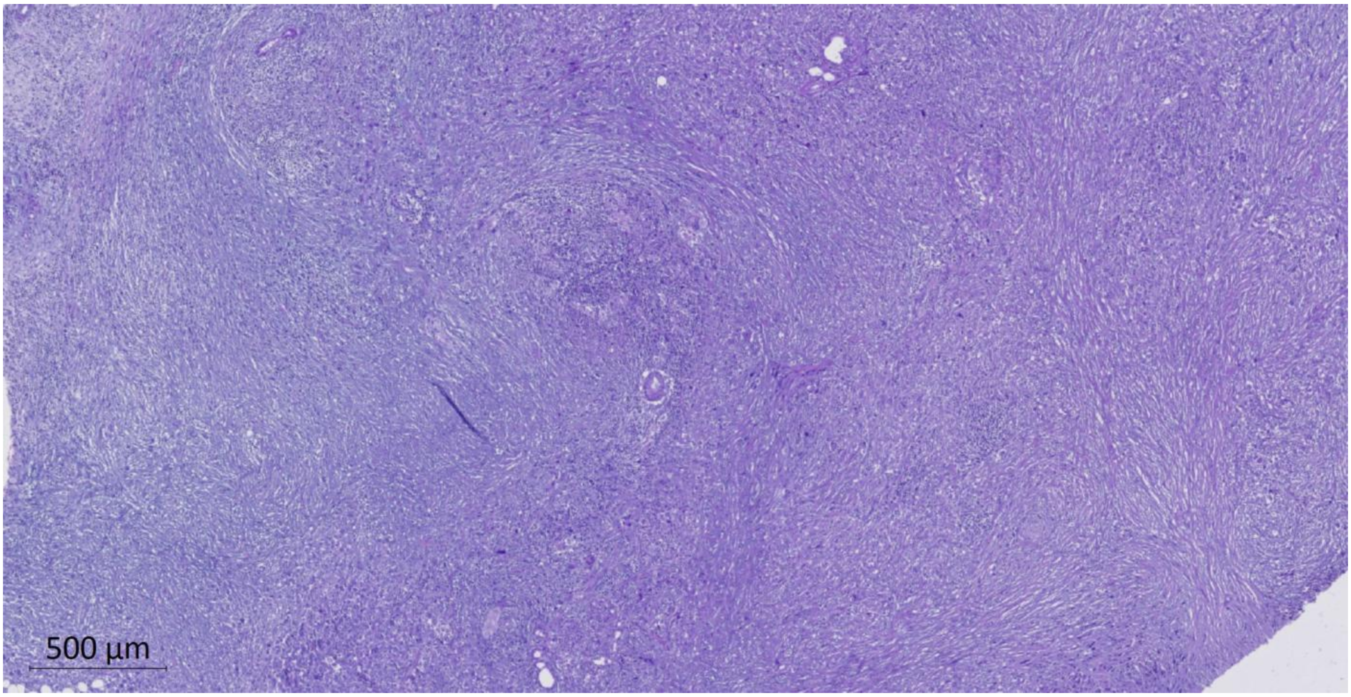

PDAC 2 - BL

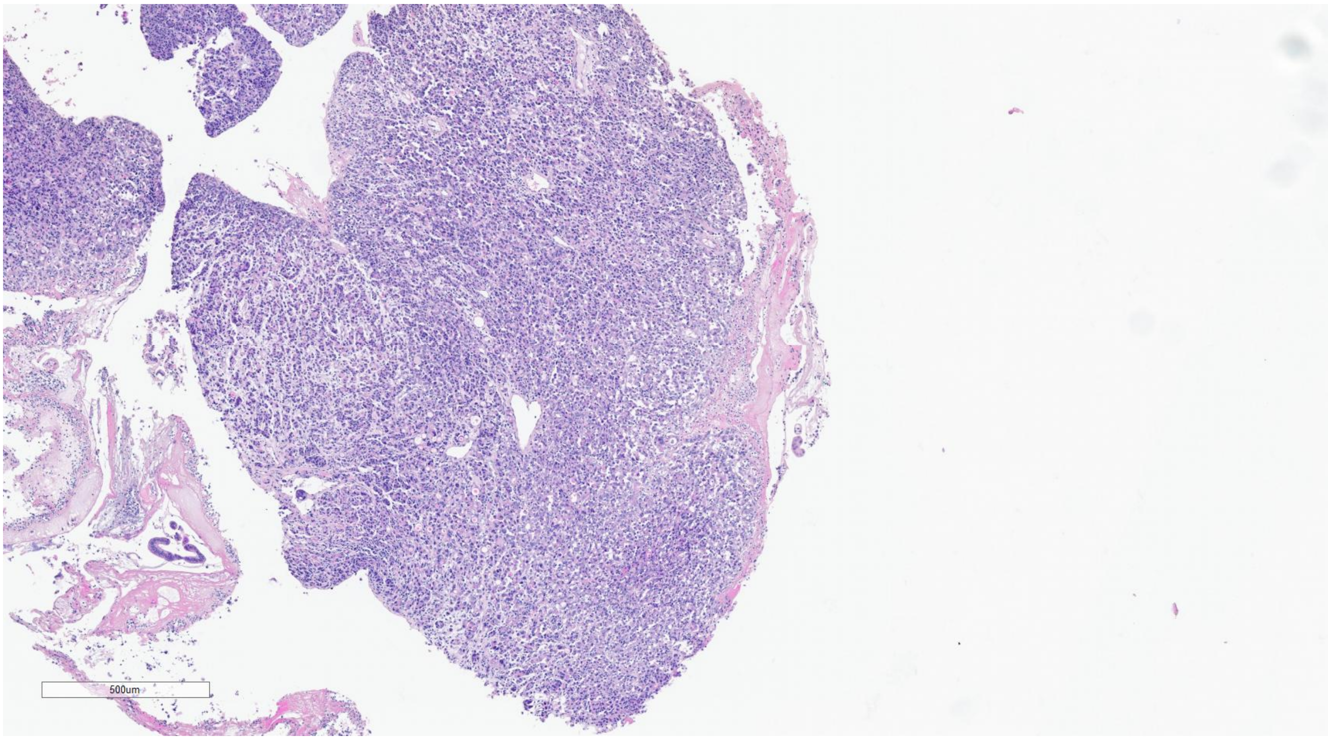

PDAC 3 - CL

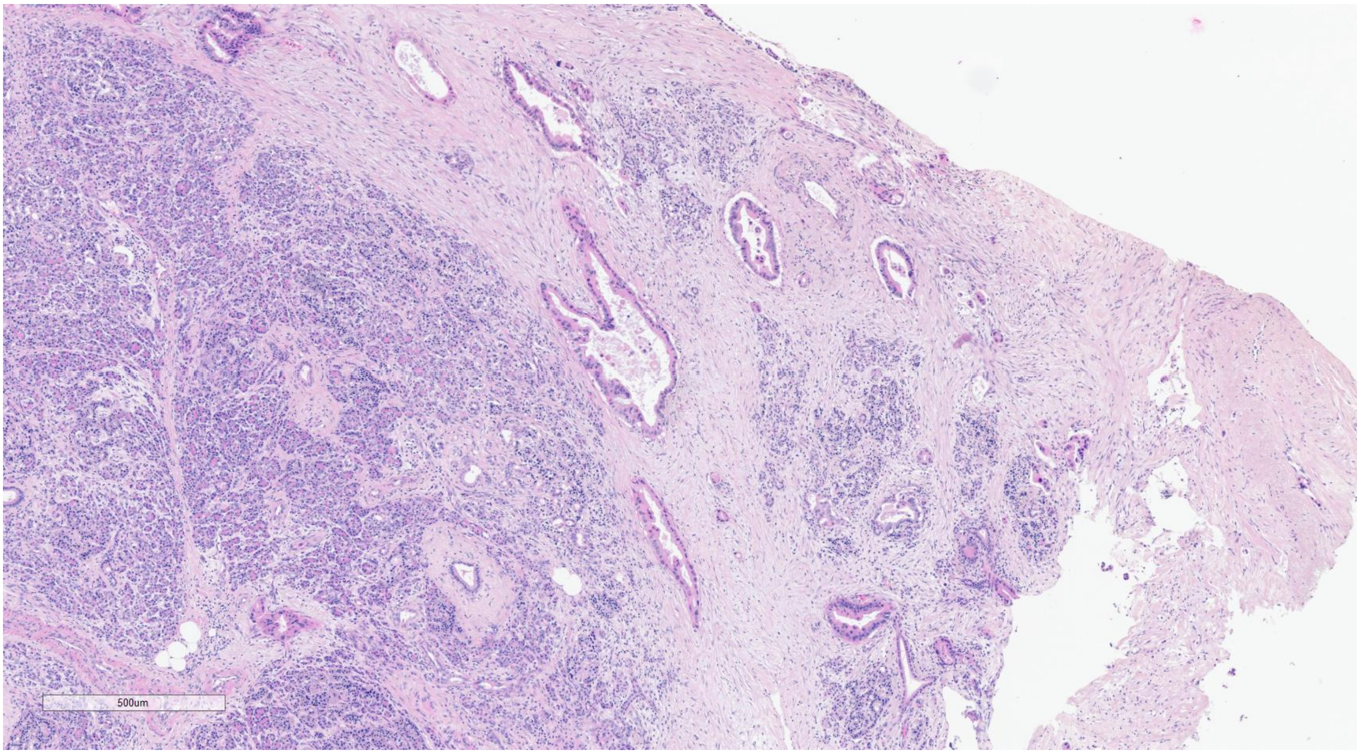

PDAC 4 - CL

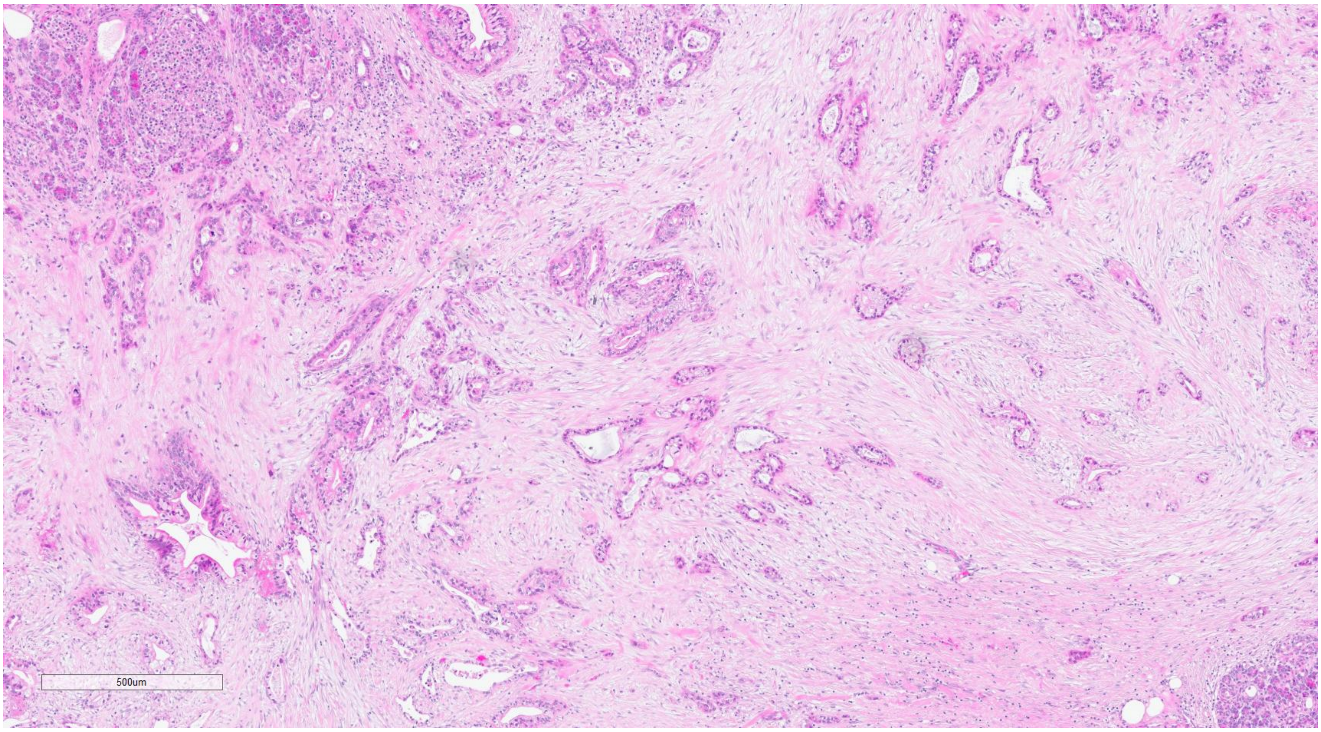

PDAC 5 - CL

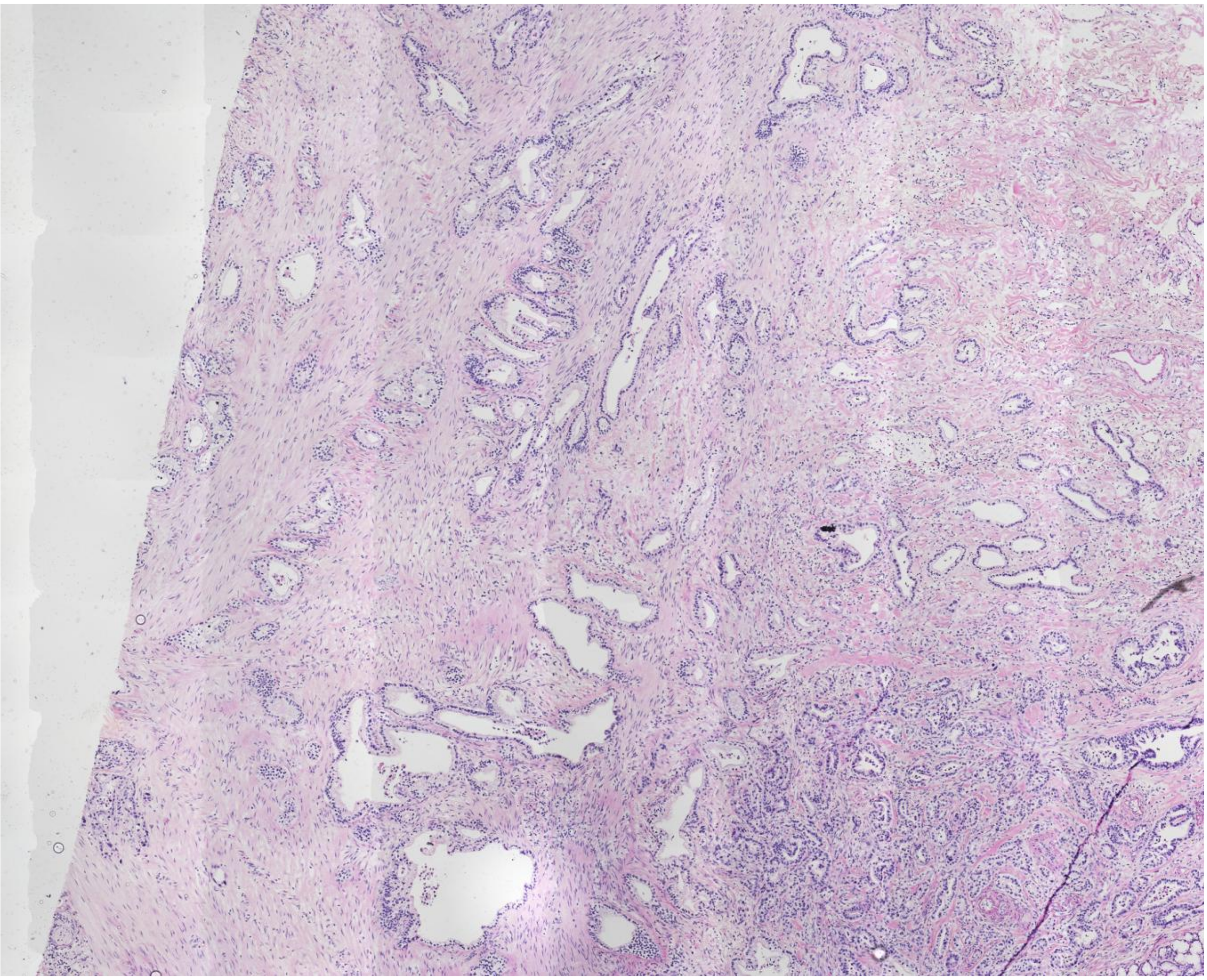

PDAC 6 - BL

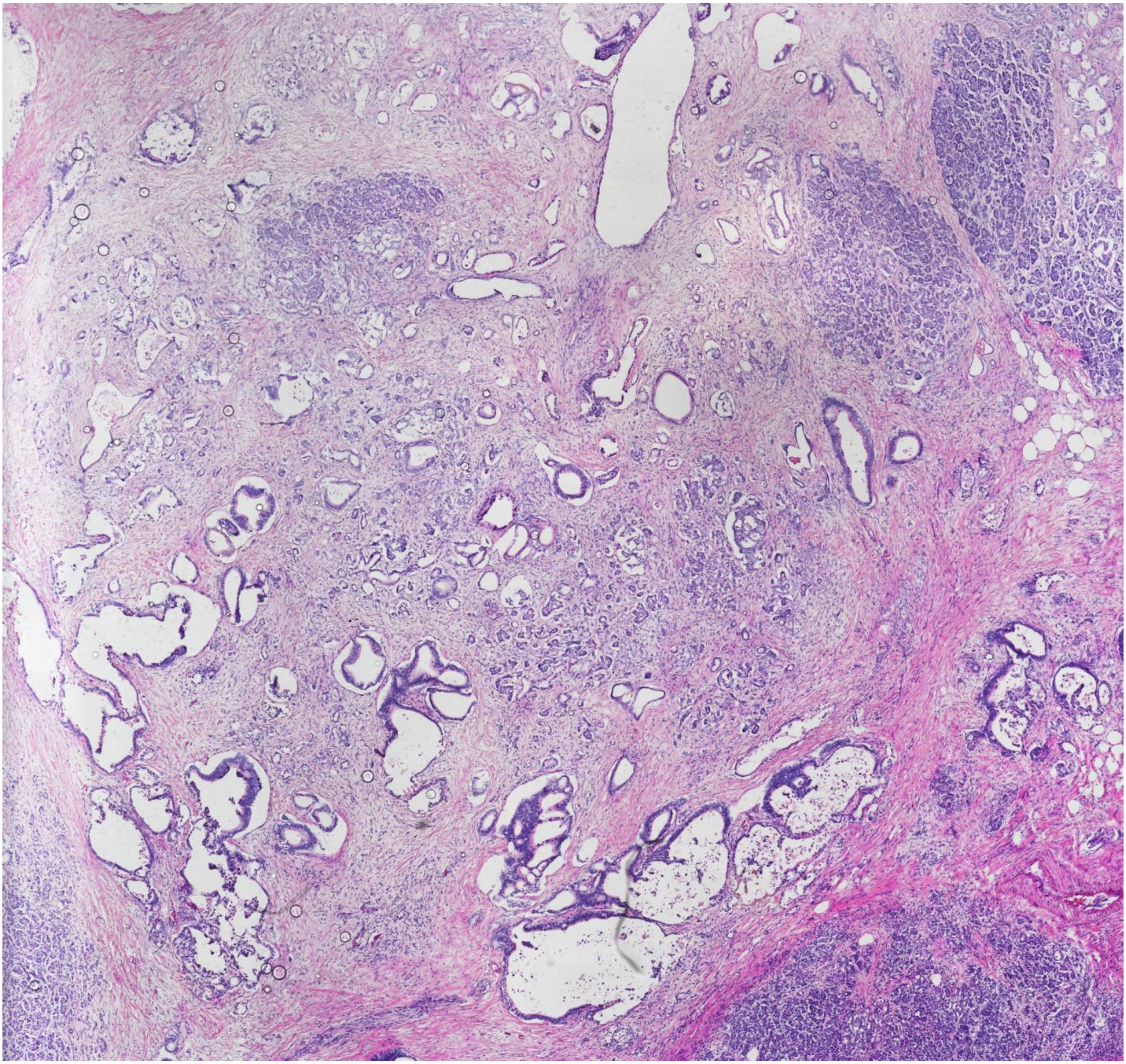

PDAC 7 - BL

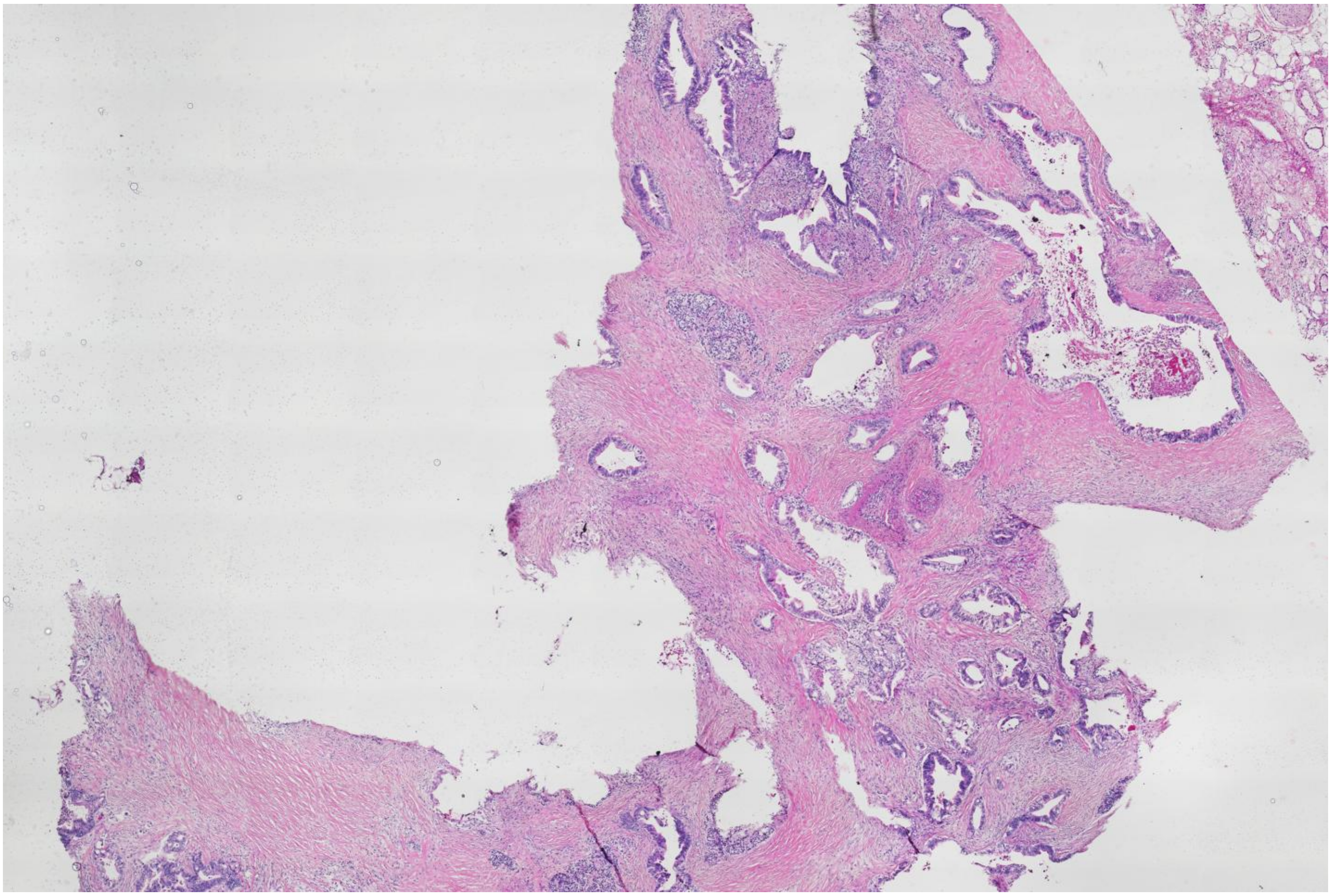
